## Supplementary Information for "Internetwork connectivity of molecular networks across species of life"

### 1 Methods

#### 1.1 Simulating multiplex degree-degree coupling and redundancy

Multiplexes with different degree-degree coupling and redundancy values were generated using a simulated annealing sampling approach [1]. This is achieved by randomly shuffling gene labels in the TRN network and the sampling is biased iteratively to match degree-degree coupling and redundancy with the desired values. Protein labels in the PPI network are kept fixed. The steps are described in Algorithm 1 below.

Algorithm 1:

1. Randomly shuffle gene labels in the TRN network, while protein labels in the PPI network are held constant. Calculate absolute difference between the shuffled and observed degree-degree coupling and redundancy. Let us call these differences $\mathrm{diffcor}^{\mathrm{cur}} = |\mathrm{cor}_{\mathrm{k,K}}^{\mathrm{cur}} - \mathrm{cor}_{\mathrm{k,K}}^{\mathrm{des}}|$ for degree-degree coupling and $\mathrm{diffRed}^{\mathrm{cur}} = |\mathrm{Redundancy}_{12}^{\mathrm{cur}} - \mathrm{Redundancy}_{12}^{\mathrm{des}}|$ for redundancy, where $\mathrm{cor}_{\mathrm{k,K}}^{\mathrm{cur}}$ is the correlation between k (degree) in the shuffled TRN network and K (degree) in the PPI network and $\mathrm{cor}_{\mathrm{k,K}}^{\mathrm{des}}$ is the desired correlation between k and K, $\mathrm{Redundancy}_{12}^{\mathrm{cur}}$ is the redundancy in the shuffled multiplex and $\mathrm{Redundancy}_{12}^{\mathrm{des}}$ is the desired Redundancy. Superscript 'cur' represents that these are the current values of the variables. Save the current gene labeling in the TRN network

2. Randomly select a subset of size N (we use N = 10) from the genes in the TRN network, and randomly shuffle their labels. Save this modified labeling of the TRN as a new variable. Calculate the new proposed difference variables $\mathrm{diffcor}^{\mathrm{prop}} = |\mathrm{cor}_{\mathrm{k,K}}^{\mathrm{cur}} - \mathrm{cor}_{\mathrm{k,K}}^{\mathrm{des}}|$ and $\mathrm{diffRed}^{\mathrm{prop}} = |\mathrm{Redundancy}_{12}^{\mathrm{cur}} - \mathrm{Redundancy}_{12}^{\mathrm{des}}|$as the absolute difference between the degree-degree coupling and redundancy in the new proposed multiplex and the desired values, respectively. Superscript 'prop' represents that these are the proposed values of the variables.

3. Calculate the difference $\Delta = \nu(\mathrm{diffcor}^{\mathrm{cur}} - \mathrm{diffcor}^{\mathrm{prop}}) + (\mathrm{diffRed}^{\mathrm{cur}} - \mathrm{diffRed}^{\mathrm{prop}})$, where $\nu$ is a scaling factor; we use $\nu = 10$. If $\Delta >= 0$, save the labeling proposed in step 2 as the current labeling for TRN. Otherwise, accept the labeling proposed in step 2 with probability $e^{\Delta/T}$, where $T = T_0 e^{-\lambda L}$ is a temperature variable, $T_0$ is the initial temperature (we use $T_0 = 1000$), $\lambda$ is a rate parameter (we use $\lambda = 0.01$) and L is the iteration number for the simulated annealing loop.

4. Repeat steps 2-3 until $\mathrm{diffcor}^{\mathrm{cur}}$ and $\mathrm{diffRed}^{\mathrm{cur}}$ reach below pre-defined thresholds (we use $\mathrm{diffcor}^{\mathrm{cur}} = 0.001$ and $\mathrm{diffRed}^{\mathrm{cur}} = 3/\sigma(\mathrm{Edges}_{12}^{\mathrm{null}})$ as the thresholds).

The node labeling for TRN achieved at the end of the simulated annealing loop has the desired degree-degree coupling and redundancy between the TRN and PPI layers. Since simulated annealing is a stochastic process, we repeat this procedure multiple times.

We can also match in-degree and out-degree couplings using this algorithm. For this purpose, $\mathrm{diffcor}^{\mathrm{cur}}$ and $\mathrm{diffcor}^{\mathrm{prop}}$ can be decomposed into $\mathrm{diffcor}^{\mathrm{cur}} = \mathrm{diffcor}_{\mathrm{in}}^{\mathrm{cur}} + \mathrm{diffcor}_{\mathrm{out}}^{\mathrm{cur}}$ and $\mathrm{diffcor}^{\mathrm{prop}} = \mathrm{diffcor}_{\mathrm{in}}^{\mathrm{prop}} + \mathrm{diffcor}_{\mathrm{out}}^{\mathrm{prop}}$ respectively, where $\mathrm{diffcor}_{\mathrm{in}}^{\mathrm{cur}} = |\mathrm{cor}_{\mathbf{k_{in}},\mathbf{K}}^{\mathrm{cur}} - \mathrm{cor}_{\mathbf{k_{in}},\mathbf{K}}^{\mathrm{des}}|$, $\mathrm{diffcor}_{\mathrm{out}}^{\mathrm{cur}} = |\mathrm{cor}_{\mathbf{k_{out}},\mathbf{K}}^{\mathrm{cur}} - \mathrm{cor}_{\mathbf{k_{out}},\mathbf{K}}^{\mathrm{des}}|$, $\mathrm{diffcor}_{\mathrm{in}}^{\mathrm{prop}} = |\mathrm{cor}_{\mathbf{k_{in}},\mathbf{K}}^{\mathrm{prop}} - \mathrm{cor}_{\mathbf{k_{in}},\mathbf{K}}^{\mathrm{des}}|$ and $\mathrm{diffcor}_{\mathrm{out}}^{\mathrm{prop}} = |\mathrm{cor}_{\mathbf{k_{out}},\mathbf{K}}^{\mathrm{prop}} - \mathrm{cor}_{\mathbf{k_{out}},\mathbf{K}}^{\mathrm{des}}|$, $\mathrm{cor}_{\mathbf{k_{in}},\mathbf{K}}^{\mathrm{cur}}$ ($\mathrm{cor}_{\mathbf{k_{in}},\mathbf{K}}^{\mathrm{prop}}$) and $\mathrm{cor}_{\mathbf{k_{out}},\mathbf{K}}^{\mathrm{cur}}$ ($\mathrm{cor}_{\mathbf{k_{out}},\mathbf{K}}^{\mathrm{prop}}$) are the correlations between $k_{in}$, $k_{out}$ in the shuffled TRN network and K in the PPI network respectively and, $\mathrm{cor}_{\mathbf{k_{in}},\mathbf{K}}^{\mathrm{des}}$ and $\mathrm{cor}_{\mathbf{k_{out}},\mathbf{K}}^{\mathrm{des}}$ are the desired correlations between $k_{in}$, $k_{out}$ and K respectively.

#### 1.2 Multiplex null model

We use a *Zero-Coupling-Zero-Redundancy* null model to assess the effect of degree-degree coupling and redundancy on robustness. This model is generated as follows.

*Zero-Coupling-Zero-Redundancy*

Under this null model, we generate null multiplexes which have no degree-degree coupling and no redundancy. We generate multiplexes from this model by setting $\mathrm{cor}_{\mathbf{k_{in}},\mathbf{K}}^{\mathrm{des}} = \mathrm{cor}_{\mathbf{k_{out}},\mathbf{K}}^{\mathrm{des}} = \mathrm{Redundancy}_{12}^{\mathrm{des}} = 0$ in Algorithm 1. This procedure is repeated 100 times to generate a distribution of multiplexes in the null model.

#### 1.3 Sampling a subset of multiplex with specific degree-degree coupling and redundancy.

Subsets of a multiplex with different degree-degree coupling and redundancy values were sampled using a simulated annealing sampling approach [1]. This is achieved by randomly sampling gene-protein pairs and the sampling is biased iteratively to match degree-degree coupling and redundancy with the desired values. The steps are described in Algorithm 2 below.

Algorithm 2:

1. Randomly sample gene-protein pairs from the multiplex. Calculate absolute difference between the sampled and observed degree-degree coupling and redundancy. Let us call these differences $\mathrm{diffcor}^{\mathrm{cur}} = |\mathrm{cor}_{\mathrm{k,K}}^{\mathrm{cur}} - \mathrm{cor}_{\mathrm{k,K}}^{\mathrm{des}}|$ for degree-degree coupling and $\mathrm{diffRed}^{\mathrm{cur}} = |\mathrm{Redundancy}_{12}^{\mathrm{cur}} - \mathrm{Redundancy}_{12}^{\mathrm{des}}|$ for redundancy, where $\mathrm{cor}_{\mathrm{k,K}}^{\mathrm{cur}}$ is the correlation between k (degree) in the sampled TRN network and K (degree) in the sampled PPI network and $\mathrm{cor}_{\mathrm{k,K}}^{\mathrm{des}}$ is the desired correlation between k and K, $\mathrm{Redundancy}_{12}^{\mathrm{cur}}$ is the redundancy in the sampled multiplex and $\mathrm{Redundancy}_{12}^{\mathrm{des}}$ is the desired Redundancy.

Superscript 'cur' represents that these are the current values of the variables. Save the current labels of the sampled gene-protein pairs in the multiplex.

2. Randomly select a subset of size N (we use N = 5) from the set of gene-proteins sampled in the previous step. Also, randomly select N gene-protein pairs from the set of gene-protein pairs not in the sampled set. Swap the N gene-proteins from the sampled set with the N genes in the not-sampled set. Save this sampled set as a new variable. Calculate the new proposed difference variables $\mathrm{diffcor}^{\mathrm{prop}} = |\mathrm{cor}_{\mathrm{k,K}}^{\mathrm{cur}} - \mathrm{cor}_{\mathrm{k,K}}^{\mathrm{des}}|$ and $\mathrm{diffRed}^{\mathrm{prop}} = |\mathrm{Redundancy}_{12}^{\mathrm{cur}} - \mathrm{Redundancy}_{12}^{\mathrm{des}}|$as the absolute difference between the degree-degree coupling and redundancy in the new proposed sampled subset of the multiplex and the desired values, respectively. Superscript 'prop' represents that these are the proposed values of the variables.

3. Calculate the difference $\Delta = \nu(\mathrm{diffcor}^{\mathrm{cur}} - \mathrm{diffcor}^{\mathrm{prop}}) + (\mathrm{diffRed}^{\mathrm{cur}} - \mathrm{diffRed}^{\mathrm{prop}})$, where $\nu$ is a scaling factor; we use $\nu = 10$. If $\Delta >= 0$, save the labeling proposed in step 2 as the current labeling for TRN. Otherwise, accept the labeling proposed in step 2 with probability $e^{\Delta/T}$, where $T = T_0 e^{-\lambda L}$ is a temperature variable, $T_0$ is the initial temperature (we use $T_0 = 1000$), $\lambda$ is a rate parameter (we use $\lambda = 0.01$) and L is the iteration number for the simulated annealing loop.

4. Repeat steps 2-3 until $\mathrm{diffcor}^{\mathrm{cur}}$ and $\mathrm{diffRed}^{\mathrm{cur}}$ reach below pre-defined thresholds (we use $\mathrm{diffcor}^{\mathrm{cur}} = 0.001$ and $\mathrm{diffRed}^{\mathrm{cur}} = 10$ as the thresholds).

The sampled set achieved at the end of the simulated annealing loop has the desired degree-degree coupling and redundancy between the TRN and PPI layers of the sampled subset of the multiplex. Since simulated annealing is a stochastic process, we repeat this procedure multiple times.

We can also match in-degree and out-degree couplings using this algorithm. For this purpose, $\mathrm{diffcor}^{\mathrm{cur}}$ and $\mathrm{diffcor}^{\mathrm{prop}}$ can be decomposed into $\mathrm{diffcor}^{\mathrm{cur}} = \mathrm{diffcor}_{\mathrm{in}}^{\mathrm{cur}} + \mathrm{diffcor}_{\mathrm{out}}^{\mathrm{cur}}$ and $\mathrm{diffcor}^{\mathrm{prop}} = \mathrm{diffcor}_{\mathrm{in}}^{\mathrm{prop}} + \mathrm{diffcor}_{\mathrm{out}}^{\mathrm{prop}}$ respectively, where $\mathrm{diffcor}_{\mathrm{in}}^{\mathrm{cur}} = |\mathrm{cor}_{\mathbf{k_{in},K}}^{\mathrm{cur}} - \mathrm{cor}_{\mathbf{k_{in},K}}^{\mathrm{des}}|$, $\mathrm{diffcor}_{\mathrm{out}}^{\mathrm{cur}} = |\mathrm{cor}_{\mathbf{k_{out},K}}^{\mathrm{cur}} - \mathrm{cor}_{\mathbf{k_{out},K}}^{\mathrm{des}}|$, $\mathrm{diffcor}_{\mathrm{in}}^{\mathrm{prop}} = |\mathrm{cor}_{\mathbf{k_{in},K}}^{\mathrm{prop}} - \mathrm{cor}_{\mathbf{k_{in},K}}^{\mathrm{des}}|$ and $\mathrm{diffcor}_{\mathrm{out}}^{\mathrm{prop}} = |\mathrm{cor}_{\mathbf{k_{out},K}}^{\mathrm{prop}} - \mathrm{cor}_{\mathbf{k_{out},K}}^{\mathrm{des}}|$, $\mathrm{cor}_{\mathbf{k_{in},K}}^{\mathrm{cur}}$ ($\mathrm{cor}_{\mathbf{k_{in},K}}^{\mathrm{prop}}$) and $\mathrm{cor}_{\mathbf{k_{out},K}}^{\mathrm{cur}}$ ($\mathrm{cor}_{\mathbf{k_{out},K}}^{\mathrm{prop}}$) are the correlations between $k_{in}$, $k_{out}$ in the sampled TRN network and K in the sampled PPI network respectively and, $\mathrm{cor}_{\mathbf{k_{in},K}}^{\mathrm{des}}$ and $\mathrm{cor}_{\mathbf{k_{out},K}}^{\mathrm{des}}$ are the desired correlations between $k_{in}$, $k_{out}$ and K respectively.

### 2 References


1. Kirkpatrick, Scott. "Optimization by simulated annealing: Quantitative studies." Journal of statistical physics 34.5-6 (1984): 975-986.

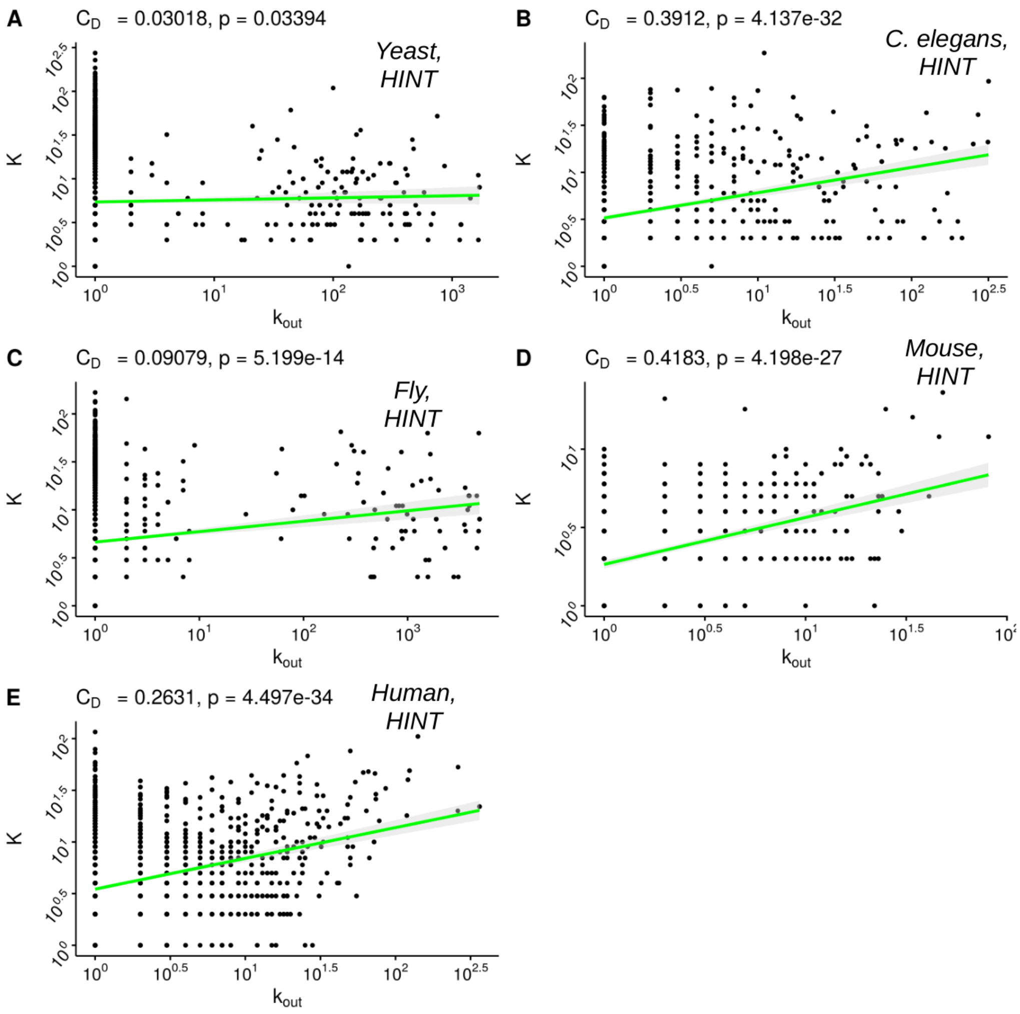


**Figure S1: Degree-degree coupling across species for HINT.** Scatter plot for degree ($K$) of proteins in the PPI network versus out-degree ($k_{out}$) of genes in the TRN for eukaryotes, A) yeast, B) *C. elegans*, C) fly, D) mouse and E) human. Both $K$ and $k_{out}$ values have been log-transformed after adding 1. The PPI networks are from the HINT database (Methods). Linear interpolated fits between $K$ and $k_{out}$ are also shown (green line) with 95% confidence region shaded in gray. Degree-degree coupling ($C_D$) values and corresponding p-values are also shown.

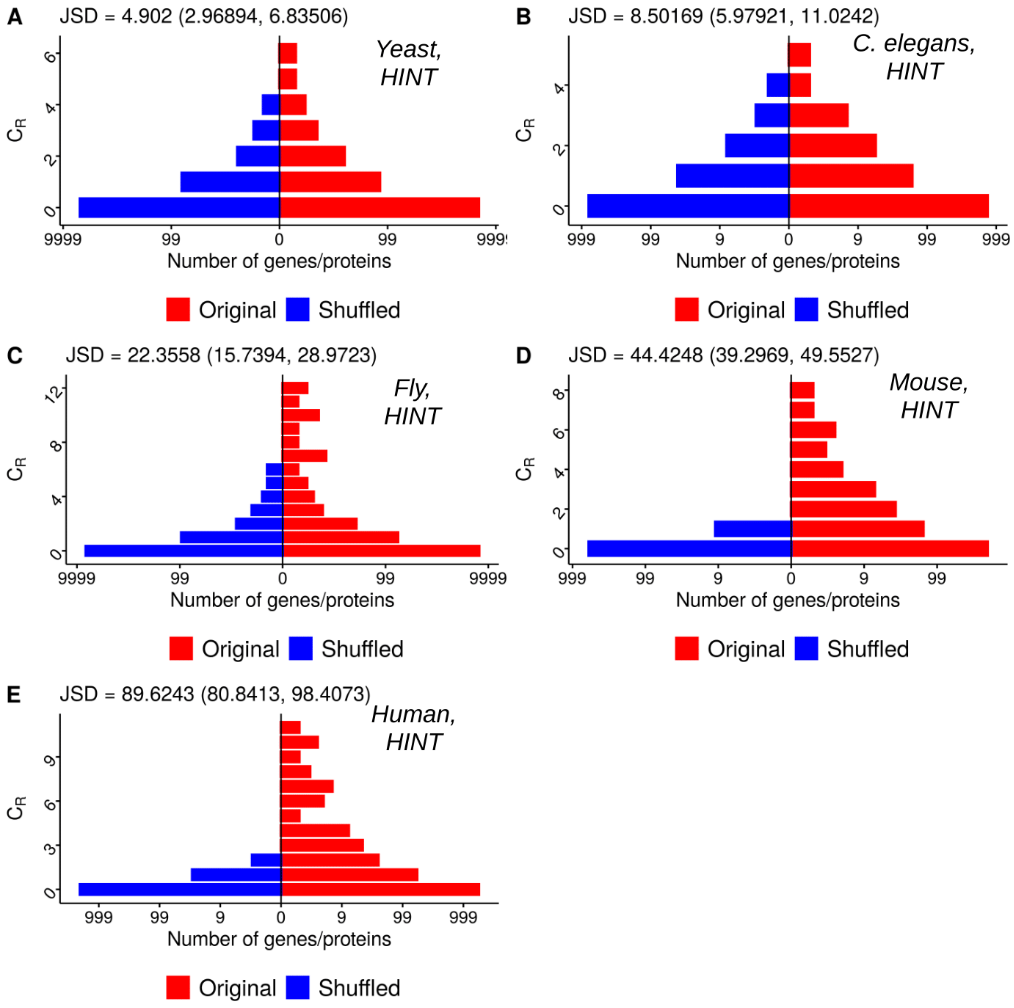


**Figure S2: Redundancy coupling across species for HINT.** Distribution of redundancy (CR) for gene-protein pairs for species multiplex and the randomly shuffled null model for eukaryotes, A) yeast, B) C. elegans, C) fly, D) mouse and E) human. For each gene-protein pair, CR is quantified by the number of redundant edges incident on that gene-protein pair. Shuffled null model is generated by randomly shuffling labels on genes in TRN, while keeping protein labels fixed in PPI. Jensen Shannon divergence (JSD) between distributions of CR in organismal and shuffled multiplexes is also shown.

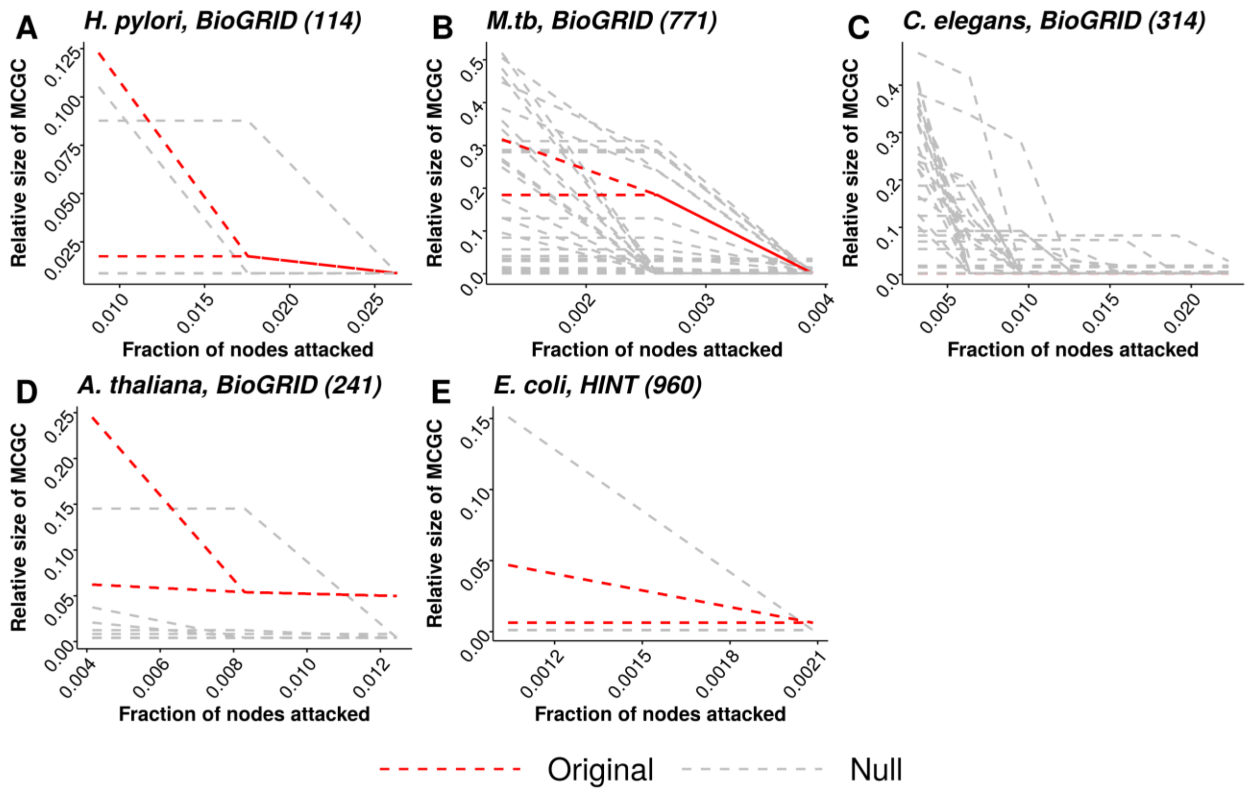


**Figure S3: Multiplex attack curves for different species.** Relative size of Mutually Connected Giant Component (MCGC) is plotted as a function of the fraction of gene-protein pairs attacked and removed from the multiplex (Methods). Attack curves are shown for five species using PPI networks from either BioGRID or HINT databases (Methods); database for a given panel are annotated next to the species name. Along with the attack curves for the species (red), attack curves for the *Zero-Coupling-Zero-Redundancy* null model are also shown (gray). Under this null model, multiplexes have no degree-degree coupling and no redundancy. On average, species attack curves are more robust than the null model. Robustness is quantified by area under the curve.

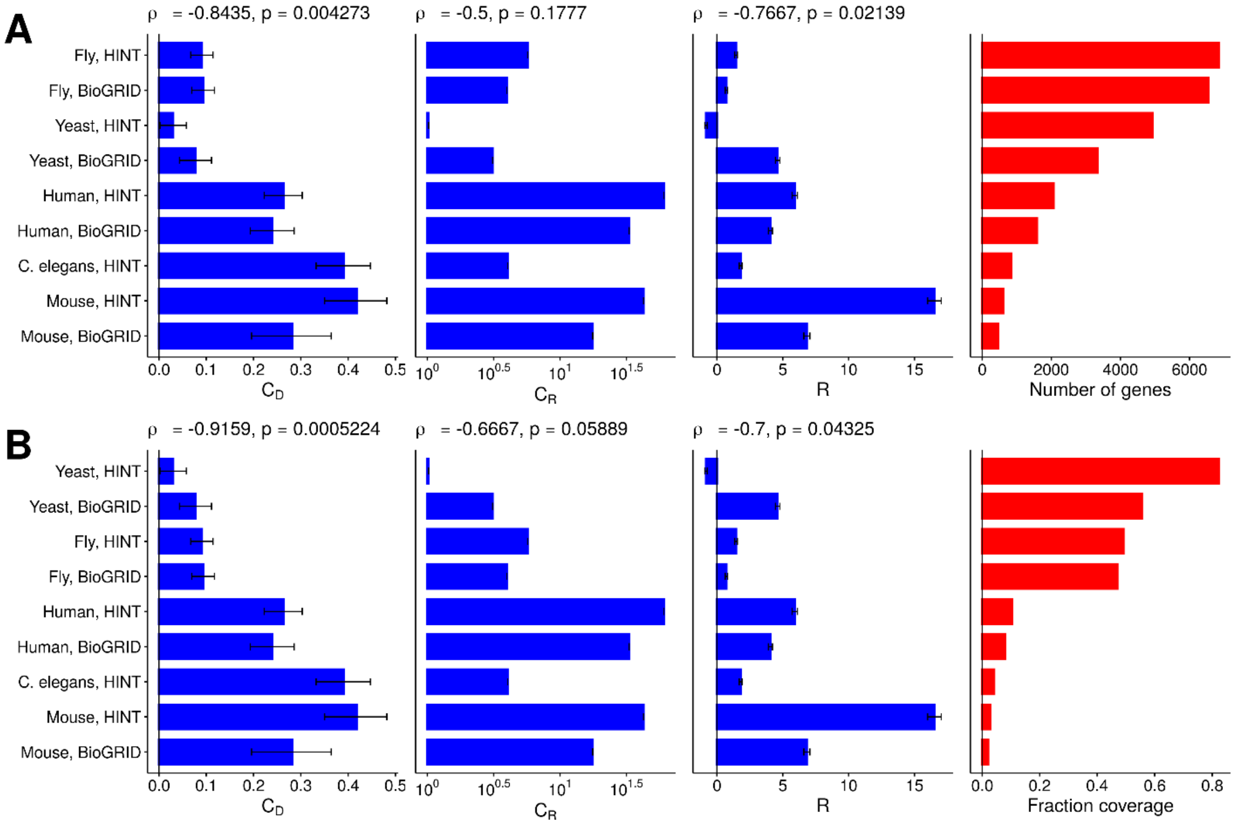


**Figure S4: Robustness, degree-degree coupling and redundancy coupling versus number of genes in the multiplex.** A) Degree-degree coupling ($C_D$), redundancy coupling ($C_R$) and robustness (R) versus number of genes in the multiplex. Spearman's correlation coefficient (with p-values) with number of genes is also mentioned. B) Degree-degree coupling ($C_D$), redundancy coupling ($C_R$) and robustness (R) versus number of fraction coverage (number of genes in the multiplex divided by the total number of protein-coding genes). Spearman's correlation coefficient (with p-values) with number of genes is also mentioned. In all the panels, error bars show 95% CI.

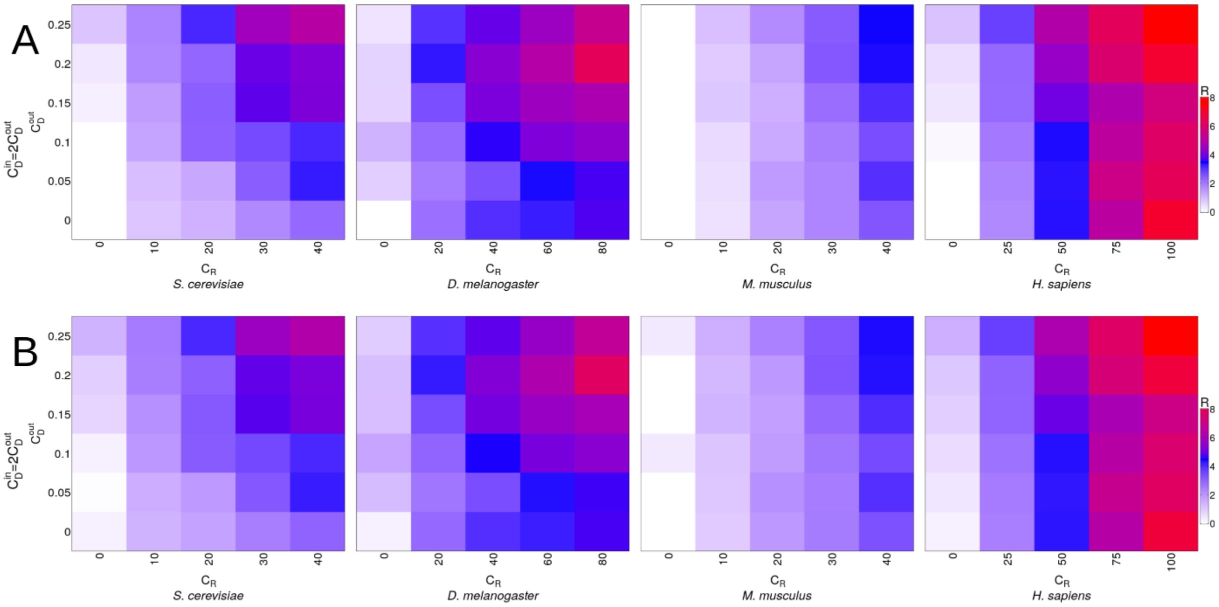


**Figure S5: 95% CI for robustness versus degree-degree and redundancy couplings plot in Figure 4B (main text).** A) Lower interval for the 95%CI for Figure 4B. Upper interval for the 95%CI for Figure 4B.

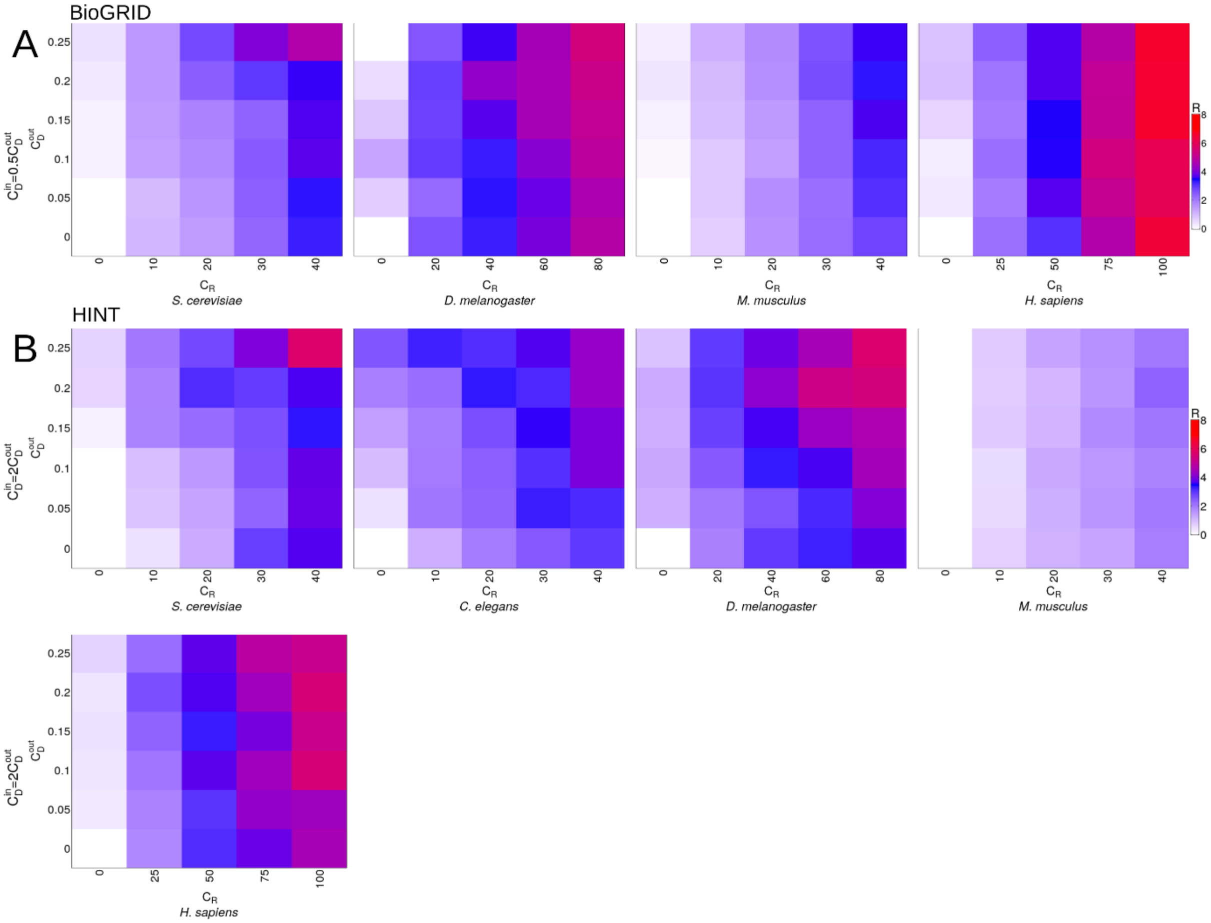


**Figure S6: Robustness vs degree-degree and redundancy couplings with different relationship between in-degree and out-degree based degree-degree couplings.** We sample a subset of gene-protein pairs from the species multiplexes (sizes for the subsets are: *S. cerevisiae*-1000, *C. elegans*-500, *D. melanogaster*-2000, *M. musculus*-300, *H. sapiens*-500) with specific $C_D$ and $C_R$ values. We repeat the sampling 100 times. We explore $C_D$ and $C_R$ over a grid. For each point over the 2D grid, the heatmap shows the robustness (R) value. R is computed by comparing *RobustArea* of any point over the grid against the lower-left point of the grid (with $C_D = 0$, $C_R = 0$). For each point over the grid, for each of the sampled subset, targeted attack is performed. Mean R values are shown here. A) $C_D^{in} = 0.5\ C_D^{out}$. BioGRID PPI networks are used. A) $C_D^{in} = 2\ C_D^{out}$. HINT PPI networks are used. $C_D^{out}$: degree-degree coupling between $k_{out}$ and K, $C_D^{in}$: degree-degree coupling between $k_{in}$ and K.

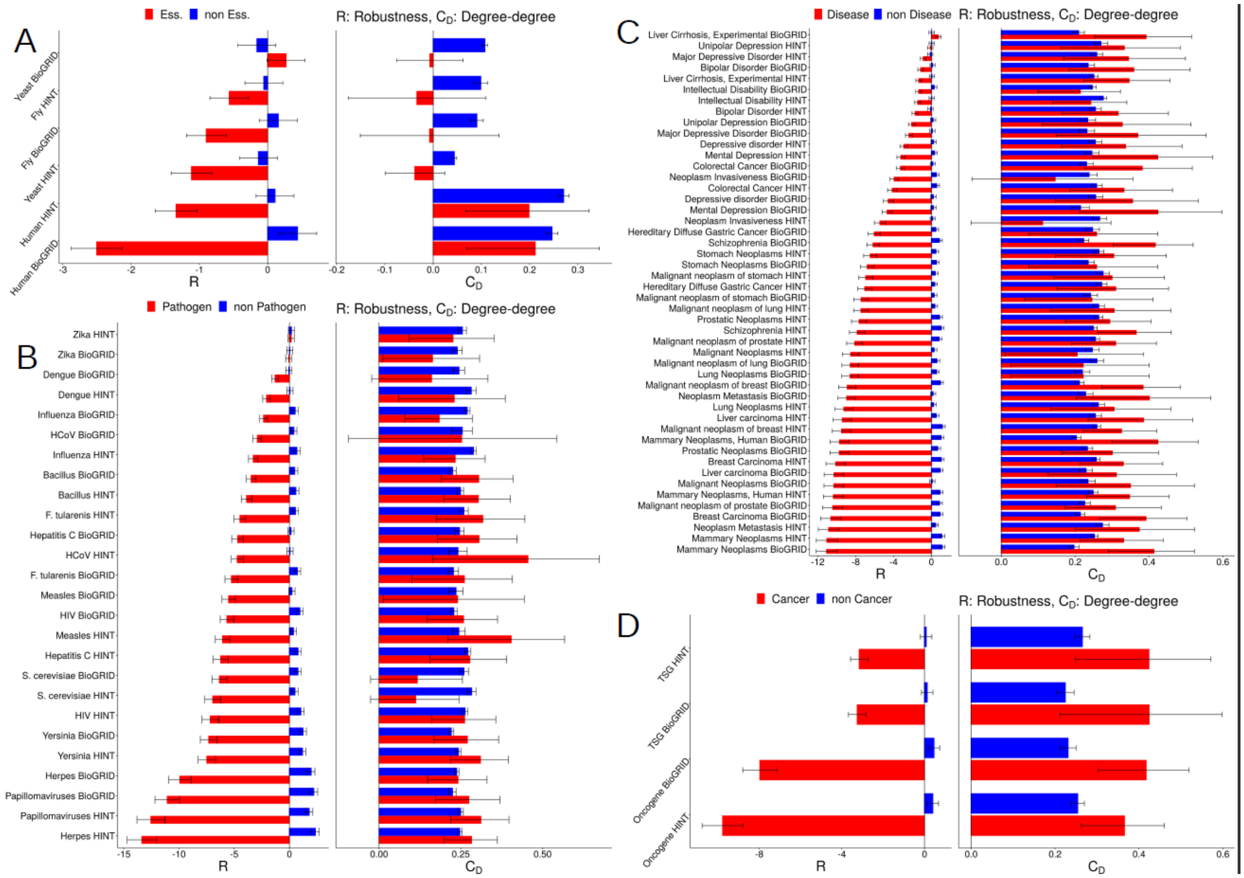

**Figure S7: Robustness to attack on functionally important genes and proteins is not dependent on degree-degree coupling in human multiplex.** A) (Left) Essential genes and proteins (red) have lower robustness to targeted attack across species compared to non-essential genes and proteins (blue). (Right) Expect for yeast, R and $C_D$ are similar for essential and non-essential genes B) (Left) Pathogen-related genes and proteins (red) have lower R compared to targeted attack across pathogens compared to non-pathogen genes and proteins (blue). (Right) R and $C_D$ are similar for pathogen and non-pathogen genes. C) (Left) Disease-related genes and proteins (red) have lower R to targeted attack across diseases compared to non-disease genes and proteins (blue). (Right) R and $C_D$ are similar for disease and non-disease genes. D) (Left) Oncogenes (tumor suppressor genes (TSGs)) and proteins (red) have lower R to targeted attack across pathogens compared to non-oncogenes (non-TSGs) and proteins (blue). (Right) R and $C_D$ are similar for oncogenes (TSGs) and non-oncogenes (non-TSGs). In all the panels, database used for PPI networks is annotated on the y-axis (Methods). In all the panels, relative robustness R is measured against a random set of genes and proteins used as the null (Methods), and $C_R$ values are calculated as the difference in redundancy against the random set of genes. Error bars show 95% CI.

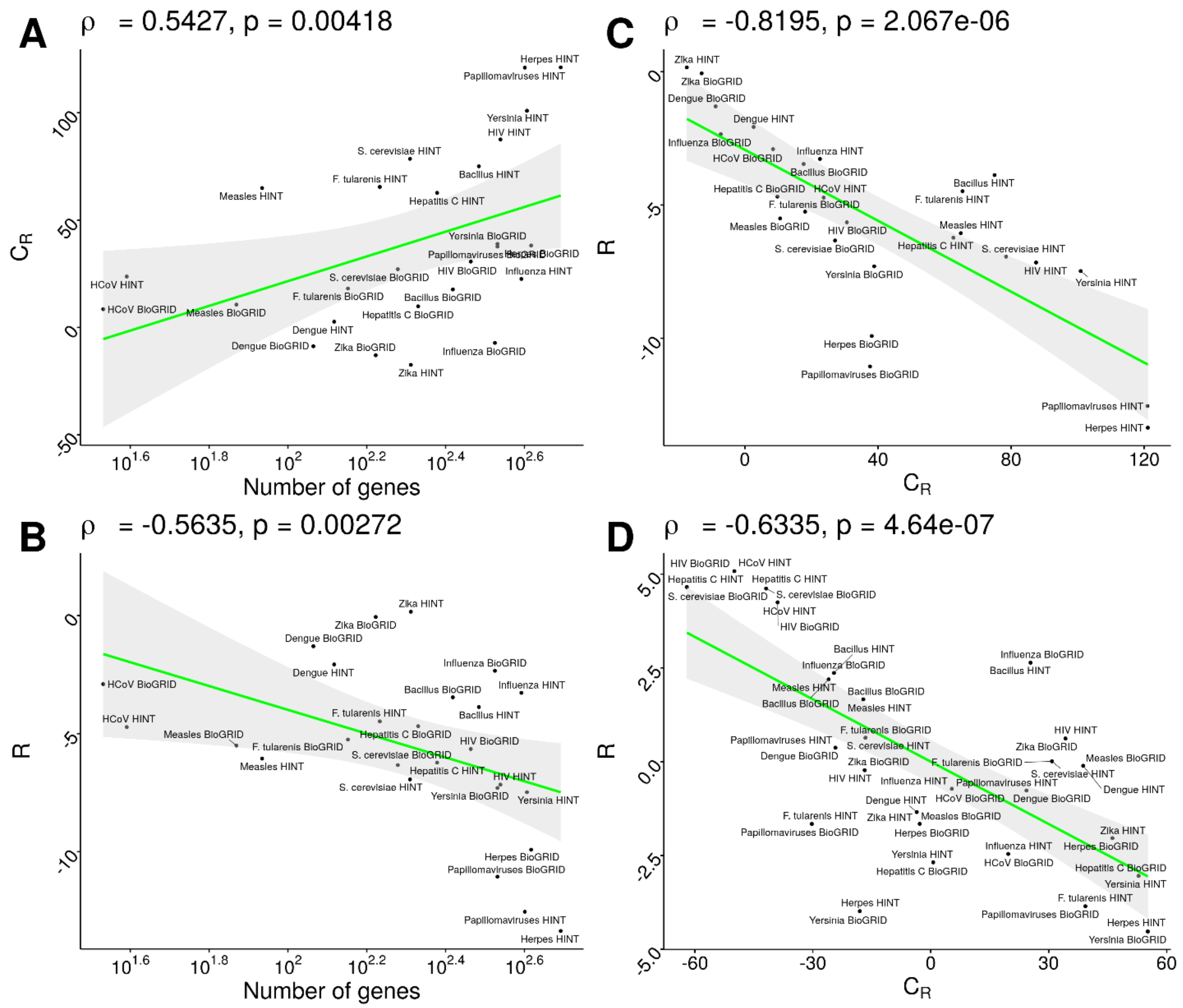


**Figure S8: Redundancy and robustness versus number of pathogen genes in the human multiplex.** A) $C_R$ for pathogen genes versus number of pathogen genes. B) R for pathogen genes versus number of pathogen genes. C) R versus $C_R$ for pathogen genes. D) R versus $C_R$ for pathogen genes while controlling for number of genes. R and $C_R$ are linearly regressed against number of genes and residuals are plotted. Pathogen names are annotated with the PPI database used for the analysis.

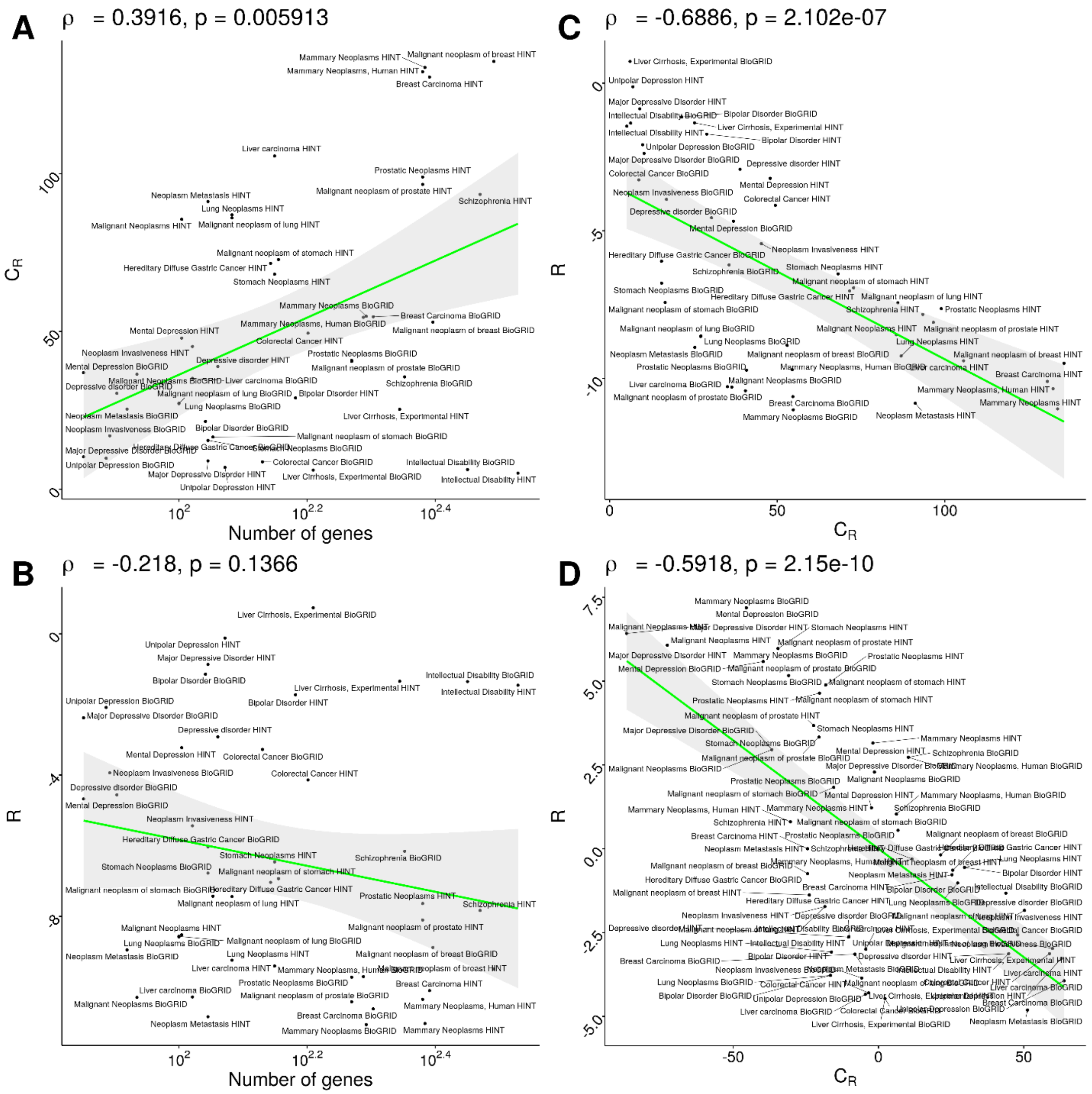


**Figure S9: Redundancy and robustness versus number of disease genes in the human multiplex.** A) $C_R$ for disease genes versus number of disease genes. B) R for disease genes versus number of disease genes. C) R versus $C_R$ for disease genes. D) R versus $C_R$ for disease genes while controlling for number of genes. R and $C_R$ are linearly regressed against number of genes and residuals are plotted. Disease names are annotated with the PPI database used for the analysis.

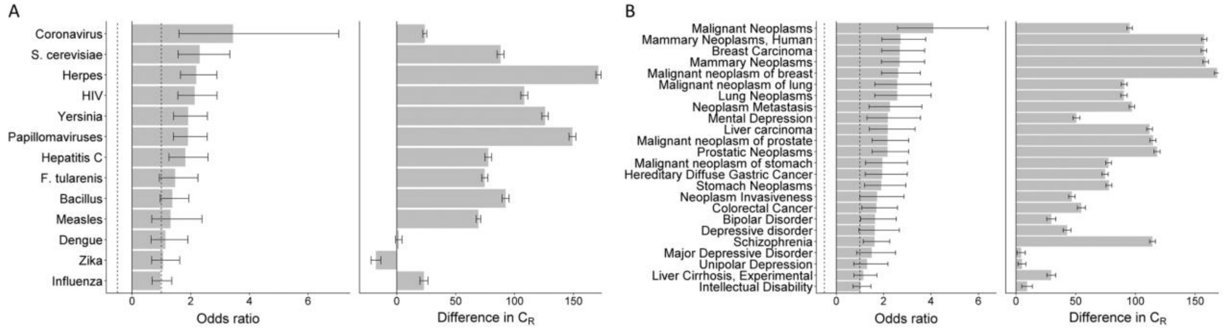


**Figure S10: Disease- and pathogen-related genes are enriched in the human multiplex:** A) (Left) Odds ratio between pathogen and non-pathogen genes for different pathogens. (Right) difference in redundancy coupling ($C_R$) between pathogen and non-pathogen genes. B) (Left) Odds ratio between disease and non- disease genes for different diseases. (Right) difference in redundancy coupling ($C_R$) between disease and non- disease genes. In all the panels, error bars show 95% confidence intervals.

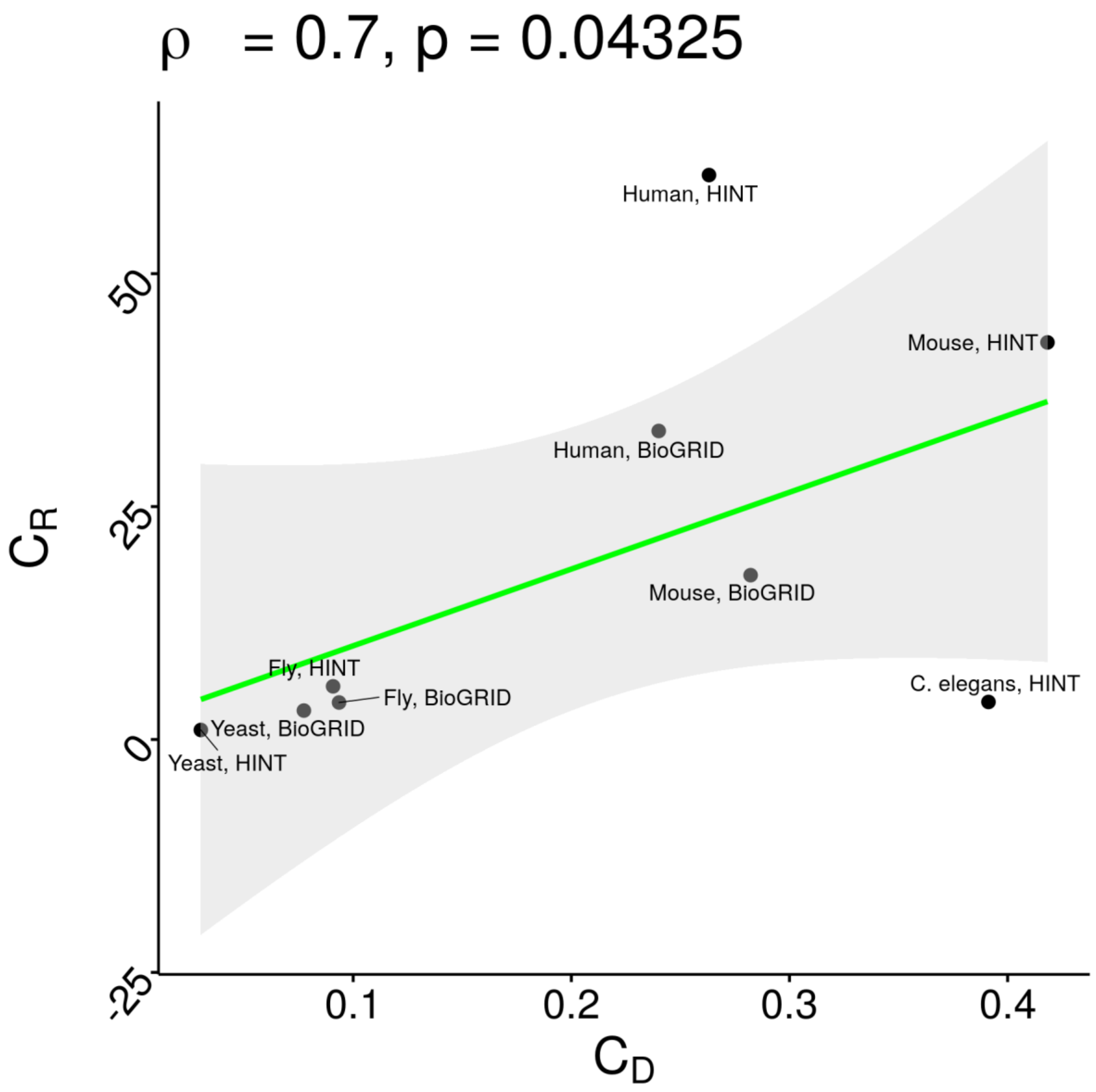


**Figure S11: Relationship between degree-degree and redundancy couplings across species:** Scatter plot between redundancy coupling ($C_R$) and degree-degree coupling ($C_D$). $C_R$ and $C_D$ are correlated with a Pearson correlation of 0.7 (p = 0.043).

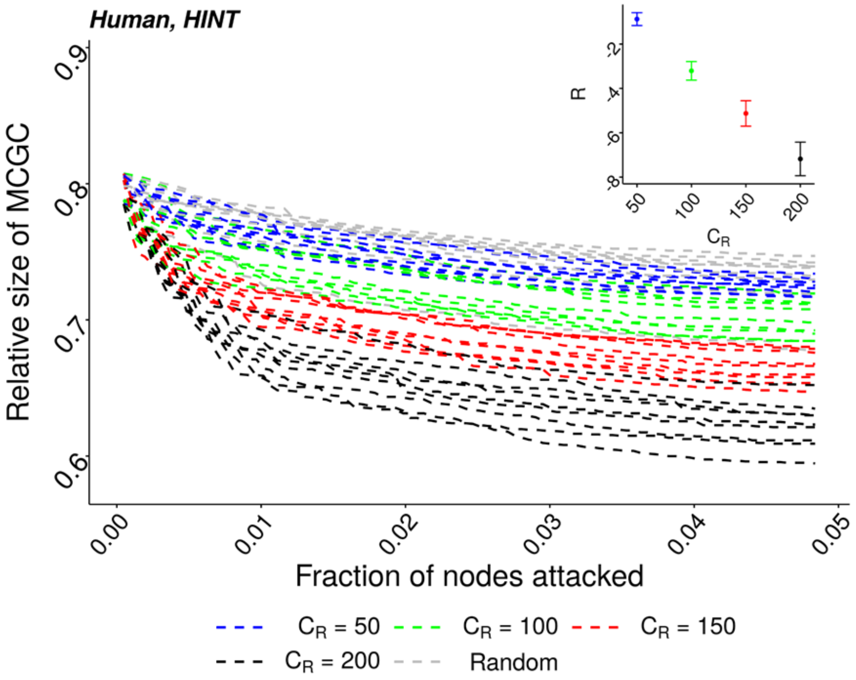


**Figure S12: Simulations showing dependence of robustness on redundancy coupling:** We sample 100 gene-protein pairs from the human multiplex. We performed the sampling experiment multiple times, each time generating a set with a different value of redundancy coupling ($C_R$). 'Random' refers to a set of randomly sampled gene-protein pairs. For each value of $C_R$, sampling was repeated 100 times. Partial attack curves for gene-protein pairs sampled from the human multiplex with different values of $C_R$ are shown. Increasing $C_R$ reduces area under the attack curve and also reduces robustness (inset). Error bars show 95% confidence intervals. HINT PPI network was used for the simulations.

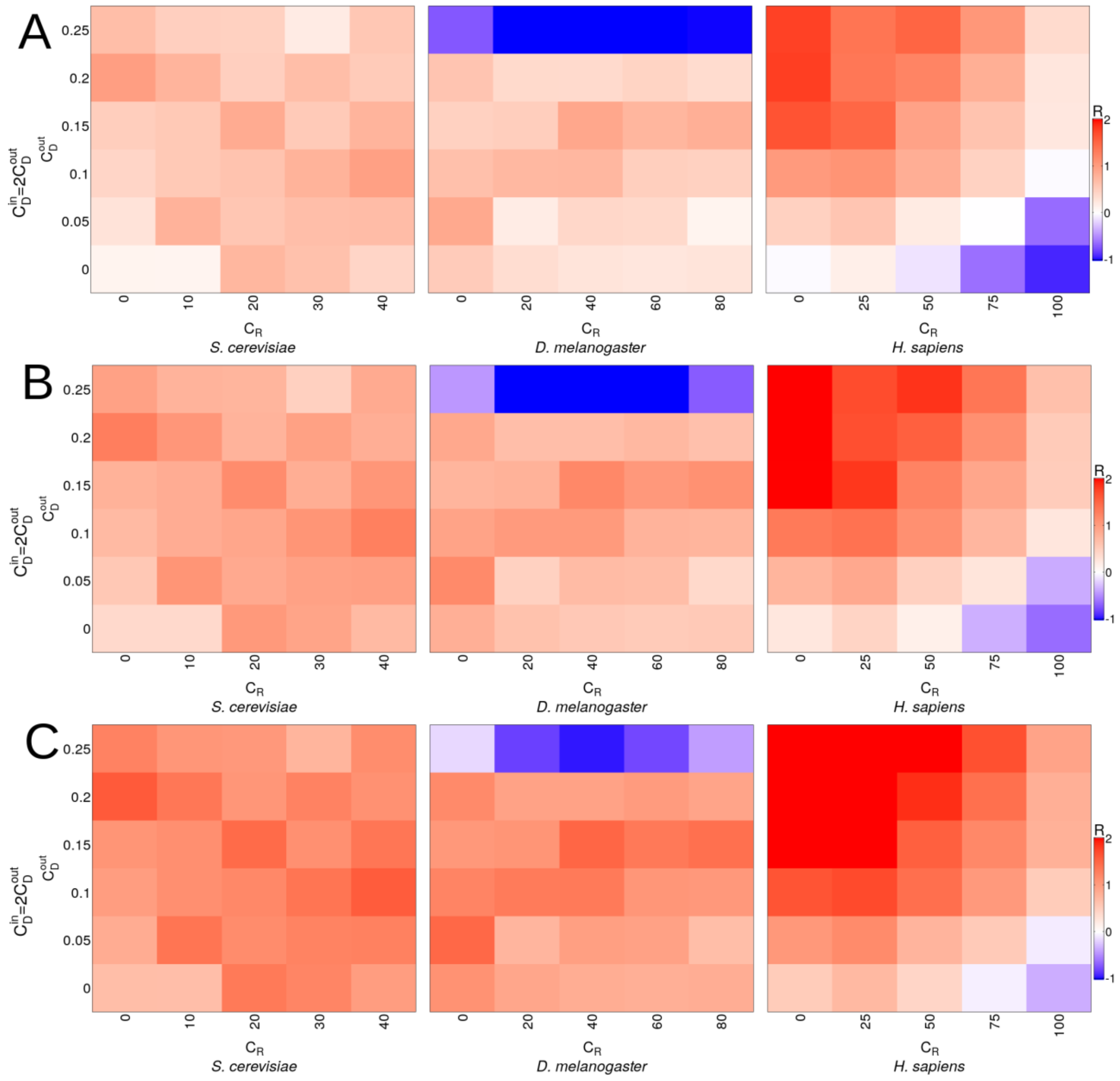


**Figure S13: Impact of size of multiplex on robustness:** We compared robustness between two subsets of gene-protein pairs, with different sizes, extracted from the same species multiplex over a grid of values for $C_D$ and $C_R$. For each pair of $C_D$ and $C_R$ values over the grid, we sample subsets of two different sizes for each species such that $C_D$ and $C_R$ values are the same for the two sampled subsets. Each cell in the heatmap shows relative robustness as the cohen's d difference between the RobustAreas of the larger and smaller subsets with the same $C_D$ and $C_R$ values. The sizes of the two subsets for each species are: *S. cerevisiae*-1000 and 1250, *D. melanogaster*-2000 and 2500, *H. sapiens*-500 and 750. For each subset, sampling was performed 100 times. A), B) and C) represent the mean, lower limit for the 95% CI and upper limit for the 95% CI for relative robustness between the two subsets. All panels use the BioGRID PPI.

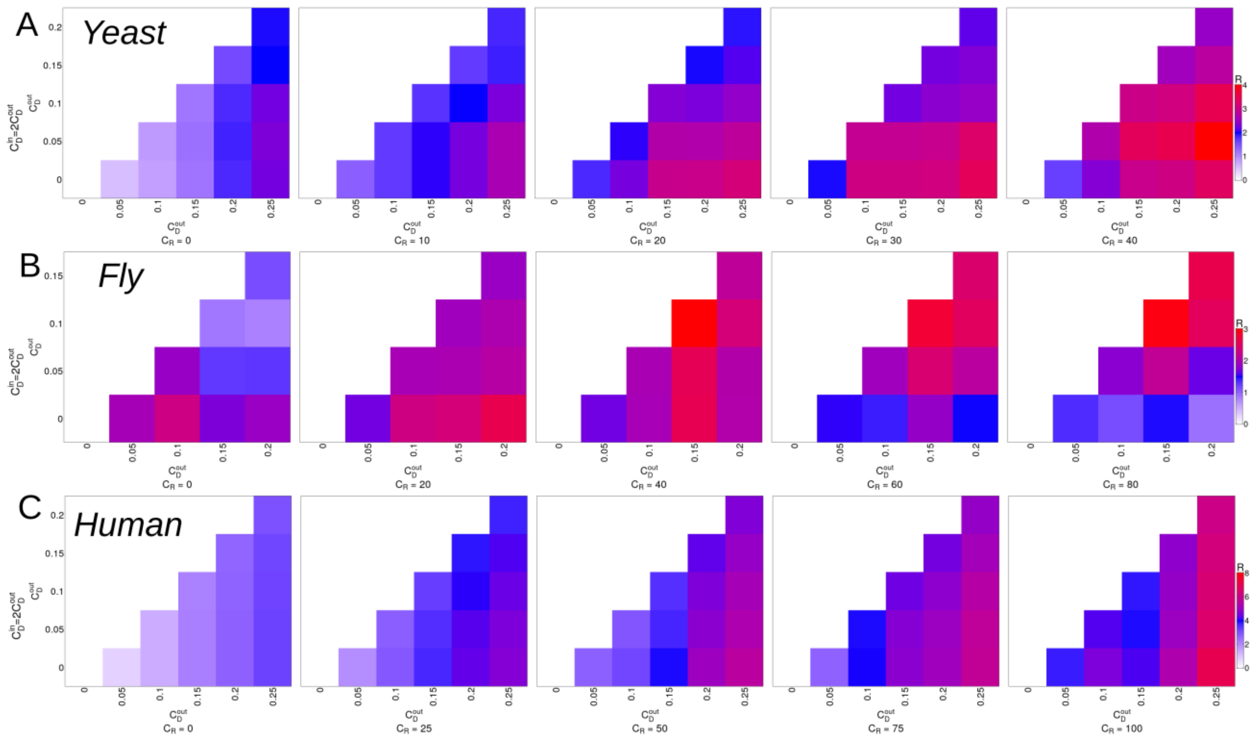


**Figure S14: Comparison between multiplexes of different sizes can reveal the impact of degree-degree coupling on robustness:** We compared robustness between two subsets of gene-protein pairs, with different sizes, extracted from the same species multiplex over a grid of values for $C_D$ for the two subsets. Values on the y-axis are the $C_D$ values for the smaller subset against which we compare robustness of the larger subset. $C_D$ values for the larger subset are shown on the x-axis. For each pair of $C_D$ values over the grid, the smaller sized subset has $C_D$ value shown on the y-axis, while the larger subset has $C_D$ value shown on the x-axis. Each heatmap corresponds to one value of $C_R$, which is same for both the subsets. Each cell in the heatmap shows relative robustness as the cohen's d difference between the RobustAreas of the larger and smaller subsets. The sizes of the two subsets for each species are: *S. cerevisiae*-1000 and 1250, *D. melanogaster*-2000 and 2500, *H. sapiens*-500 and 750. For each subset, sampling was performed 100 times. Moving along the x-axis while keeping the y-axis value fixed, we see that robustness (R) increases with increasing $C_D$ of the larger sized multiplex. For each row in a heatmap, R is computed by comparing the larger subset against the smaller subset. All panels use the BioGRID PPI. A), B) and C) show heatmaps for *S. cerevisiae*, *D. melanogaster* and *H. sapiens* respectively. Only the lower triangular part of the matrix is shown.

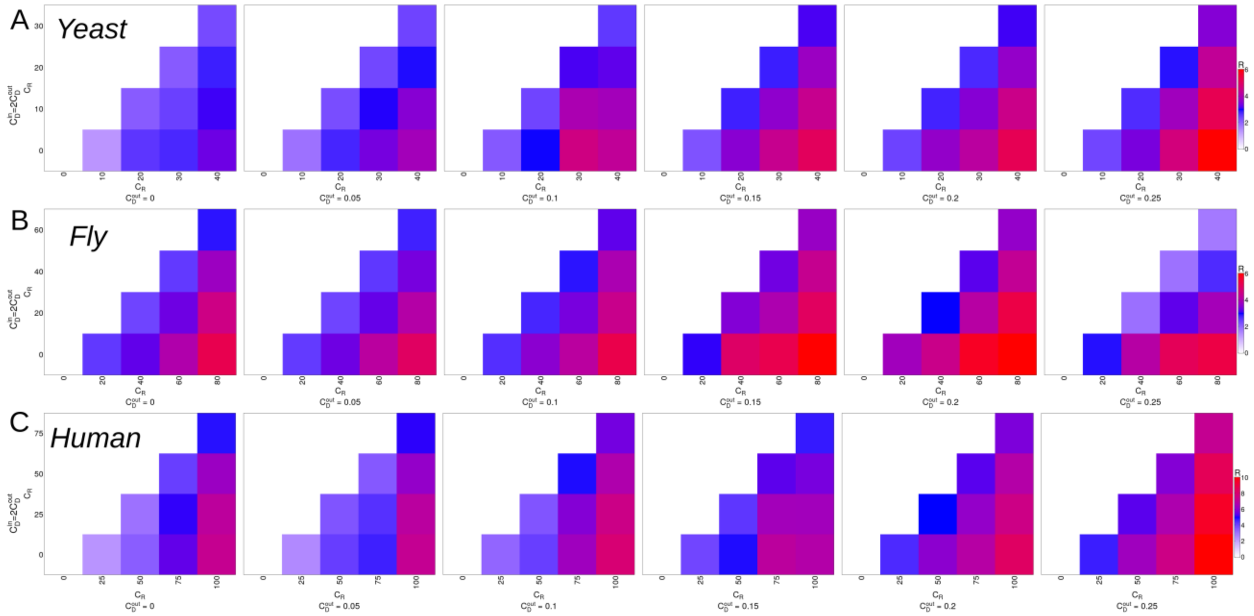


**Figure S15: Comparison between multiplexes of different sizes can reveal the impact of redundancy coupling on robustness:** We compared robustness between two subsets of gene-protein pairs, with different sizes, extracted from the same species multiplex over a grid of values for $C_R$ for the two subsets. Values on the y-axis are the $C_R$ values for the smaller subset against which we compare robustness of the larger subset. $C_R$ values for the larger subset are shown on the x-axis. For each pair of $C_R$ values over the grid, the smaller sized subset has $C_R$ value shown on the y-axis, while the larger subset has $C_R$ value shown on the x-axis. Each heatmap corresponds to one value of $C_D$, which is same for both the subsets. Each cell in the heatmap shows relative robustness as the cohen's d difference between the RobustAreas of the larger and smaller subsets. The sizes of the two subsets for each species are: *S. cerevisiae*-1000 and 1250, *D. melanogaster*-2000 and 2500, *H. sapiens*-500 and 750. For each subset, sampling was performed 100 times. Moving along the x-axis while keeping the y-axis value fixed, we see that robustness (R) increases with increasing $C_R$ of the larger sized multiplex. For each row in a heatmap, R is computed by comparing the larger subset against the smaller subset. All panels use the BioGRID PPI. A), B) and C) show heatmaps for *S. cerevisiae*, *D. melanogaster* and *H. sapiens* respectively. Only the lower triangular part of the matrix is shown.
