## Supplementary material for "Internetwork connectivity of molecular networks across species of life": Table S1

**Table S1: Transcriptional regulatory and protein-protein interaction networks**

| **Species** | **Number of Genes/Proteins** (TRN, PPI1, PPI2) | **%Overlapping Proteome Coverage** (TRN-PPI1, TRN-PPI2) | **Edges** (TRN, PPI1, PPI2) | **Average. Degree** ($k_{in}/k_{out}$, K1, K2) | **Size of LCC** (TRN, PPI1, PPI2) | **Reference** (TRN, PPI1, PPI2) |
|---|---|---|---|---|---|---|
| *H. pylori* | 1590, 789 | 7.64, NA | 718, 1687, NA | 2.94, 8.21, NA | 436, 731, NA | [1], [2], NA |
| *M. tuberculosis* | 1624, 2907 | 19.74, NA | 3212, 8007, NA | 1.98, 5.51, NA | 1604, 2895, NA | [3], [4], NA |
| *E. coli* | 1817, 1223, 2172 | 5.12, 22.64 | 3072, 1961, 3655 | 1.70, 3.19, 2.78 | 1695, 987, 1236 | [5], [6], [7] |
| *S. cerevisiae* | 3712, 4256, 5305 | 55.56, 82.3 | 9869, 14891, 23202 | 2.66, 7.0, 8.52 | 3707, 4133, 5210 | [8], [9], [7] |
| *C. elegans* | 3351, 3099, 4533 | 1.6, 4.16 | 26294, 5347, 12234 | 7.85, 3.45, 5.22 | 3351, 2860, 4298 | [10], [9], [7] |
| *D. melanogaster* | 12323, 6677, 7505 | 47.02, 49.14 | 157395, 19347, 30181 | 12.77, 5.80, 7.36 | 12323, 6540, 7378 | [11], [9], [7] |
| *A. thaliana* | 790, 5779, 5646 | 0.87, 0.56 | 1431, 14592, 23410 | 1.81, 5.05, 8.18 | 725, 5447, 5159 | [12], [9], [7] |
| *M. musculus* | 2456, 2056, 2429 | 2.08, 2.75 | 6490, 2595, 3542 | 2.64, 2.52, 2.59 | 2403, 1464, 1485 | [13], [9], [7] |
| *H. sapiens* | 2862, 11833, 12856 | 7.93, 10.33 | 8403, 48595, 62435 | 2.94, 8.21, 9.40 | 2804, 11731, 12117 | [13], [9], [7] |

Table showing the characteristics of TRN and PPI networks used for the nine different. For H, pylori and M. tuberculosis, there is one TRN and one PPI each. For all the other species, we have one TRN and two PPI each. The two PPIs are labeled as PPI1 and PPI2 respectively; protein degrees for these networks are labeled K1 and K2 respectively. We have also given the corresponding references for each network dataset used in this study.

**Abbreviations:**

TRN: Transcriptional Regulatory Network
PPI: Protein-Protein Interaction
PPI1: PPI network from the BioGRID database [9] or other species-specific publications [2,4,6].
PPP2: PPI network from the HINT database [7].
LCC: Largest Connected Component is the subset of genes/proteins in TRN/PPI network where every gene/protein is reachable from every other gene/protein.

**Definitions:**
%Overlapping Proteome Coverage: Percentage of the proteome included in the multiplexes consisting of TRN and PPI1 networks or TRN and PPI2 networks, respectively.
Edges: Number of edges in the TRN, PPI1 or PPI2 networks.

**References:**

1. Danielli A, Amore G, Scarlato V. Built shallow to maintain homeostasis and persistent infection: insight into the transcriptional regulatory network of the gastric human pathogen Helicobacter pylori. PLoS pathogens. 2010;6:e1000938.

2. Häuser R, Ceol A, Rajagopala SV, Mosca R, Siszler G, Wermke N, et al. A second-generation protein–protein interaction network of helicobacter pylori. Molecular & cellular proteomics. 2014;13:1318–1329.

3. Sanz J, Navarro J, Arbués A, Martín C, Marijuán PC, Moreno Y. The transcriptional regulatory network of Mycobacterium tuberculosis. PloS one. 2011;6:e22178.

4. Wang Y, Cui T, Zhang C, Yang M, Huang Y, Li W, et al. Global protein- protein interaction network in the human pathogen Mycobacterium tuberculosis H37Rv. Journal of proteome research. 2010;9:6665–6677.

5. Salgado H, Martínez-Flores I, Bustamante VH, Alquicira-Hernández K, García-Sotelo JS, García-Alonso D, et al. Using RegulonDB, the Escherichia coli K-12 Gene Regulatory Transcriptional Network Database. Current protocols in bioinformatics. 2018;61:1–32.

6. Rajagopala SV, Sikorski P, Kumar A, Mosca R, Vlasblom J, Arnold R, et al. The binary protein-protein interaction landscape of Escherichia coli. Nature biotechnology. 2014;32:285.

7. Das J, Yu H. HINT: High-quality protein interactomes and their applications in understanding human disease. BMC systems biology. 2012;6:92.

8. Teixeira MC, Monteiro PT, Palma M, Costa C, Godinho CP, Pais P, et al. YEASTRACT: an upgraded database for the analysis of transcription regulatory networks in Saccharomyces cerevisiae. Nucleic acids research. 2017;46:D348–D353.

9. Chatr-Aryamontri A, Oughtred R, Boucher L, Rust J, Chang C, Kolas NK, et al. The BioGRID interaction database: 2017 update. Nucleic acids research. 2017;45:D369–D379.

10. Bass JIF, Pons C, Kozlowski L, Reece-Hoyes JS, Shrestha S, Holdorf AD, et al. A gene-centered C. elegans protein–DNA interaction network provides a framework for functional predictions. Molecular systems biology. 2016;12.

11. Murali T, Pacifico S, Yu J, Guest S, Roberts GG, Finley RL. DroID 2011: a comprehensive, integrated resource for protein, transcription factor, RNA and gene interactions for Drosophila. Nucleic acids research. 2010;39:D736–D743.

12. Jin J, He K, Tang X, Li Z, Lv L, Zhao Y, et al. An Arabidopsis transcriptional regulatory map reveals distinct functional and evolutionary features of novel transcription factors. Molecular biology and evolution. 2015;32:1767–1773.

13. Han H, Cho J-W, Lee S, Yun A, Kim H, Bae D, et al. TRRUST v2: an expanded reference database of human and mouse transcriptional regulatory interactions. Nucleic acids research. 2017;46:D380–D386.
