## Supplementary material for "Internetwork connectivity of molecular networks across species of life": Code: Attack_functions_new_cpp.cpp

```
#include <RcppArmadillo.h>
#include <iostream>
#include <fstream>
#include <list>
#include <vector>
#include <omp.h>
#include <progress.hpp>
#include <progress_bar.hpp>
#define NIL -1 
// #include <otherpackage.h>
// #include "/igraph/igraph.h"
//[[Rcpp::depends(RcppProgress, RcppArmadillo)]]
using namespace Rcpp;
using namespace std;

// Graph class represents a undirected graph 
// using adjacency list representation 
class Graph 
{ 
  int V;    // No. of vertices 
  
  // Pointer to an array containing adjacency lists 
  list<int> *adj;
  
  // A function used by DFS 
  void DFSUtil(int v, bool visited[], int w);
  void bridgeUtil(int v, bool visited[], int disc[], int low[], 
                  int parent[]); 
public: 
  Graph(int V);   // Constructor 
  ~Graph();
  void printSizeNodeAdjList(int v);
  int SizeNodeAdjList(int v);
  void addEdge(int v, int w);
  void delEdge(int v, int w);
  void clearVertex(int v);
  void clearbridge(int NumNodes);
  void clearnonLCC( arma::vec comp );
  void delBridge( arma::vec comp );
  void bridge();
  void connectedComponents();
  int *component;
  list<int> *adj_bridge;
};

void Graph::printSizeNodeAdjList( int v){
  cout << adj[v].size() << endl;
}
int Graph::SizeNodeAdjList( int v){
  return adj[v].size();
}
// Method to print connected components in an 
// undirected graph 
void Graph::connectedComponents() 
{ 
  // Mark all the vertices as not visited 
  bool *visited = new bool[V]; 
  for(int v = 0; v < V; v++) 
    visited[v] = false; 
  
  for (int v=0; v<V; v++) 
  { 
    if (visited[v] == false) 
    { 
      // print all reachable vertices 
      // from v 
      DFSUtil(v, visited, v); 
      
      // cout << "\n"; 
    } 
  } 
  delete[] visited; 
} 


void Graph::DFSUtil(int v, bool visited[], int w) 
{ 
  // Mark the current node as visited and print it 
  visited[v] = true;
  component[v] = w;
  
  // cout << v << " "; 
  
  // Recur for all the vertices 
  // adjacent to this vertex 
  list<int>::iterator i; 
  for(i = adj[v].begin(); i != adj[v].end(); ++i) 
    if(!visited[*i]) 
      DFSUtil(*i, visited, w); 
}

// A recursive function that finds and prints bridges using 
// DFS traversal 
// u --> The vertex to be visited next 
// visited[] --> keeps tract of visited vertices 
// disc[] --> Stores discovery times of visited vertices 
// parent[] --> Stores parent vertices in DFS tree 
void Graph::bridgeUtil(int u, bool visited[], int disc[],  
                       int low[], int parent[]) 
{ 
  // A static variable is used for simplicity, we can  
  // avoid use of static variable by passing a pointer. 
  static int time = 0; 
  
  // Mark the current node as visited 
  visited[u] = true; 
  
  // Initialize discovery time and low value 
  disc[u] = low[u] = ++time; 
  
  // Go through all vertices aadjacent to this 
  list<int>::iterator i; 
  for (i = adj[u].begin(); i != adj[u].end(); ++i) 
  { 
    int v = *i;  // v is current adjacent of u 
    
    // If v is not visited yet, then recur for it 
    if (!visited[v]) 
    { 
      parent[v] = u; 
      bridgeUtil(v, visited, disc, low, parent); 
      
      // Check if the subtree rooted with v has a  
      // connection to one of the ancestors of u 
      low[u]  = min(low[u], low[v]); 
      
      // If the lowest vertex reachable from subtree  
      // under v is  below u in DFS tree, then u-v  
      // is a bridge 
      if (low[v] > disc[u]){
        cout << u <<" " << v << endl;
        adj_bridge[v].push_back(u); 
        adj_bridge[u].push_back(v);
      } 
      
    } 
    
    // Update low value of u for parent function calls. 
    else if (v != parent[u]) 
      low[u]  = min(low[u], disc[v]); 
  } 
} 

// DFS based function to find all bridges. It uses recursive  
// function bridgeUtil() 
void Graph::bridge() 
{ 
  // Mark all the vertices as not visited 
  bool *visited = new bool[V]; 
  int *disc = new int[V]; 
  int *low = new int[V]; 
  int *parent = new int[V]; 
  
  // Initialize parent and visited arrays 
  for (int i = 0; i < V; i++) 
  { 
    parent[i] = NIL; 
    visited[i] = false; 
  } 
  
  // Call the recursive helper function to find Bridges 
  // in DFS tree rooted with vertex 'i' 
  for (int i = 0; i < V; i++) 
    if (visited[i] == false) 
      bridgeUtil(i, visited, disc, low, parent); 
} 


void Graph::delBridge( arma::vec comp ){
  
  for( int v = 0; v < comp.size(); ++v ){
    if( adj[v].size() > 0 ){
      bool *state = new bool[comp.size()];
      for( int c = 0; c < comp.size(); ++c ){
        state[c] = false;
      }
      for (std::list<int>::iterator it=adj[v].begin(); it != adj[v].end(); ++it ){
        if( comp[v] != comp[*it] ){
          adj[*it].remove(v);
          state[*it] = true;
        }
      }
      for( int c = 0; c < comp.size(); ++c ){
        if( state[c] ){
          adj[v].remove(c);
        }
      }
      delete[] state;
    }
  }
  
  
  
}

void delBridgeInter ( Graph &gTRN, Graph &gPPI, arma::vec &TRN_comp, arma::vec &PPI_comp ){
  arma::vec TRN_un = arma::unique(TRN_comp);
  arma::vec PPI_un = arma::unique(PPI_comp);
  arma::vec TRN_count(TRN_un.size()), PPI_count(PPI_un.size());
  
  for( int i = 0; i < TRN_un.size(); ++i ){
    TRN_count(i) = 0;
  }
  for( int j = 0; j < TRN_comp.size(); ++j ){
    arma::uvec id = arma::find(TRN_un == TRN_comp(j));
    TRN_count(id) += 1;
  }
  
  for( int i = 0; i < PPI_un.size(); ++i ){
    PPI_count(i) = 0;
  }
  for( int j = 0; j < PPI_comp.size(); ++j ){
    arma::uvec id = arma::find(PPI_un == PPI_comp(j));
    PPI_count(id) += 1;
  }
  
  arma::uvec id_TRN = arma::find(TRN_count == max(TRN_count));
  arma::uvec id_PPI = arma::find(PPI_count == max(PPI_count));
  
  if( id_TRN.size() == 1 && id_PPI.size() == 1 ){
    arma::uvec id_TRN_comp = arma::find(TRN_comp == TRN_un[id_TRN(0)]);
    arma::uvec id_PPI_comp = arma::find(PPI_comp == PPI_un[id_PPI(0)]);
    
    arma::uvec id_inter = arma::intersect(id_TRN_comp, id_PPI_comp);
    
    bool *state = new bool[TRN_comp.size()];
    for( int c = 0; c < TRN_comp.size(); ++c ){
      state[c] = false;
    }
    for( int c = 0; c < id_inter.size(); ++c ){
      state[id_inter(c)] = true;
    }
    
    for( int v = 0; v < TRN_comp.size(); ++v ){
      if( !state[v] ){
        gTRN.clearVertex(v);
        gPPI.clearVertex(v);
      }
    }
    delete[] state;
    
    gTRN.connectedComponents();
    gPPI.connectedComponents();
    
    for( int i = 0; i < TRN_comp.size(); ++i ){
      TRN_comp(i) = gTRN.component[i];
      PPI_comp(i) = gPPI.component[i];
    }
  }else{
    arma::mat size_inter(id_TRN.size(), id_PPI.size());
    for( int i = 0; i < id_TRN.size(); ++i ){
      for( int j = 0; j < id_PPI.size(); ++j ){
        arma::uvec id_TRN_comp = arma::find(TRN_comp == TRN_un[id_TRN(i)]);
        arma::uvec id_PPI_comp = arma::find(PPI_comp == PPI_un[id_PPI(j)]);
        
        arma::uvec id_inter = arma::intersect(id_TRN_comp, id_PPI_comp);
        size_inter(i, j) = id_inter.size();
      }
    }
    
    arma::uvec max_inter = find(size_inter == size_inter.max());
    int j = int((max_inter(0) + 1)/size_inter.n_rows);
    int i = int(max_inter(0) - j*size_inter.n_rows);
    if( double(j*size_inter.n_rows) == double(max_inter(0) + 1) ){
      j -= 1;
      i = size_inter.n_rows - 1;
    }
    
    arma::uvec id_TRN_comp = arma::find(TRN_comp == TRN_un[id_TRN(i)]);
    arma::uvec id_PPI_comp = arma::find(PPI_comp == PPI_un[id_PPI(j)]);
    
    arma::uvec id_inter = arma::intersect(id_TRN_comp, id_PPI_comp);
    
    bool *state = new bool[TRN_comp.size()];
    for( int c = 0; c < TRN_comp.size(); ++c ){
      state[c] = false;
    }
    for( int c = 0; c < id_inter.size(); ++c ){
      state[id_inter(c)] = true;
    }
    
    for( int v = 0; v < TRN_comp.size(); ++v ){
      if( !state[v] ){
        gTRN.clearVertex(v);
        gPPI.clearVertex(v);
      }
    }
    delete[] state;
    
    gTRN.connectedComponents();
    gPPI.connectedComponents();
    
    for( int i = 0; i < TRN_comp.size(); ++i ){
      TRN_comp(i) = gTRN.component[i];
      PPI_comp(i) = gPPI.component[i];
    }
  }
  
  
  
}


void Graph::clearnonLCC( arma::vec comp ){
  arma::vec un = arma::unique(comp);
  arma::vec count(un.size());
  
  for( int i = 0; i < un.size(); ++i ){
    count(i) = 0;
  }
  for( int j = 0; j < comp.size(); ++j ){
    arma::uvec id = arma::find(un == comp(j));
    count(id) += 1;
  }
  
  arma::uvec id = arma::find(count == max(count));
  arma::uvec id_comp = arma::find(comp != un[id(0)]);
  
  for( int v = 0; v < id_comp.size(); ++v ){
    if( adj[int(id_comp(v))].size() != 0 ){
      adj[int(id_comp(v))].clear();
    }
  }
  
}

Graph::Graph(int V) 
{ 
  this->V = V; 
  adj = new list<int>[V];
  adj_bridge = new list<int>[V];
  component = new int [V];
}

Graph::~Graph() 
{ 
  delete[] adj;
  delete[] component;
}

// method to add an undirected edge 
void Graph::addEdge(int v, int w) 
{ 
  adj[v].push_back(w); 
  adj[w].push_back(v); 
}

void Graph::delEdge(int v, int w) 
{ 
  // Traversing through the second vector list 
  // and removing the first element from it 
  adj[v].remove(w);
  adj[w].remove(v);
} 

void Graph::clearVertex(int v) 
{ 
  // Traversing through the second vector list 
  // and removing the first element from it 
  if( adj[v].size() != 0 ){
    for (std::list<int>::iterator it=adj[v].begin(); it != adj[v].end(); ++it ){
      adj[*it].remove(v);
    }
    adj[v].clear();
  }
  
}

void Graph::clearbridge(int NumNodes) 
{ 
  // Traversing through the second vector list 
  // and removing the first element from it 
  for( int v = 0; v < NumNodes; ++v ){
    adj_bridge[v].clear();
  }
  
}


// Driver program to test above 
// int main() 
// { 
//   // Create a graph given in the above diagram 
//   Graph g(5); // 5 vertices numbered from 0 to 4 
//   g.addEdge(1, 0); 
//   g.addEdge(2, 3); 
//   g.addEdge(3, 4);
//   g.clearVertex(3);
//   
//   cout << "Following are connected components \n"; 
//   g.connectedComponents();
//   for( int i = 0; i < 5; i++ ){
//     cout << g.component[i] << endl;
//   }
//   
//   
//   return 0; 
// } 

int sizeTRNPPI( arma::vec TRN_comp, arma::vec PPI_comp ){
  arma::vec TRN_un = arma::unique(TRN_comp);
  arma::vec PPI_un = arma::unique(PPI_comp);
  arma::vec TRN_count(TRN_un.size()), PPI_count(PPI_un.size());
  
  for( int i = 0; i < TRN_un.size(); ++i ){
    TRN_count(i) = 0;
  }
  for( int j = 0; j < TRN_comp.size(); ++j ){
    arma::uvec id = arma::find(TRN_un == TRN_comp(j));
    TRN_count(id) += 1;
  }
  
  for( int i = 0; i < PPI_un.size(); ++i ){
    PPI_count(i) = 0;
  }
  for( int j = 0; j < PPI_comp.size(); ++j ){
    arma::uvec id = arma::find(PPI_un == PPI_comp(j));
    PPI_count(id) += 1;
  }
  
  if( max(TRN_count) > 1 && max(PPI_count) > 1 ){
    
    arma::uvec id_TRN = arma::find(TRN_count == max(TRN_count));
    arma::uvec id_PPI = arma::find(PPI_count == max(PPI_count));
    
    if( id_TRN.size() == 1 && id_PPI.size() == 1 ){
      arma::uvec id_TRN_comp = arma::find(TRN_comp == TRN_un[id_TRN(0)]);
      arma::uvec id_PPI_comp = arma::find(PPI_comp == PPI_un[id_PPI(0)]);
      
      arma::uvec id_inter = arma::intersect(id_TRN_comp, id_PPI_comp);
      
      if( id_TRN_comp.size() == id_PPI_comp.size() ){
        if( id_inter.size() == id_TRN_comp.size() ){
          return 1;
        }else{
          if( id_TRN_comp.size() == 1 ){
            return 1;
          }else{
            return 0;
          }
        }
      }else{
        return 0;
      }  
    }else{
      arma::mat size_inter(id_TRN.size(), id_PPI.size());
      for( int i = 0; i < id_TRN.size(); ++i ){
        for( int j = 0; j < id_PPI.size(); ++j ){
          arma::uvec id_TRN_comp = arma::find(TRN_comp == TRN_un[id_TRN(i)]);
          arma::uvec id_PPI_comp = arma::find(PPI_comp == PPI_un[id_PPI(j)]);
          
          arma::uvec id_inter = arma::intersect(id_TRN_comp, id_PPI_comp);
          size_inter(i, j) = id_inter.size();
        }
      }
      
      arma::uvec max_inter = find(size_inter == size_inter.max());
      int j = floor((max_inter(0) + 1)/size_inter.n_rows);
      int i = max_inter(0) - j*size_inter.n_rows;
      if( double(j*size_inter.n_rows) == double(max_inter(0) + 1) ){
        j -= 1;
        i = size_inter.n_rows - 1;
      }
      
      arma::uvec id_TRN_comp = arma::find(TRN_comp == TRN_un[id_TRN(i)]);
      arma::uvec id_PPI_comp = arma::find(PPI_comp == PPI_un[id_PPI(j)]);
      
      arma::uvec id_inter = arma::intersect(id_TRN_comp, id_PPI_comp);
      
      if( id_TRN_comp.size() == id_PPI_comp.size() ){
        if( id_inter.size() == id_TRN_comp.size() ){
          return 1;
        }else{
          if( id_TRN_comp.size() == 1 ){
            return 1;
          }else{
            return 0;
          }
        }
      }else{
        return 0;
      }  
    }
    
    
  }else{
    return(1);
  }
  
}


int sizeTRNPPIreturn( arma::vec TRN_comp, arma::vec PPI_comp ){
  arma::vec TRN_un = arma::unique(TRN_comp);
  arma::vec PPI_un = arma::unique(PPI_comp);
  arma::vec TRN_count(TRN_un.size()), PPI_count(PPI_un.size());
  
  for( int i = 0; i < TRN_un.size(); ++i ){
    TRN_count(i) = 0;
  }
  for( int j = 0; j < TRN_comp.size(); ++j ){
    arma::uvec id = arma::find(TRN_un == TRN_comp(j));
    TRN_count(id) += 1;
  }
  
  for( int i = 0; i < PPI_un.size(); ++i ){
    PPI_count(i) = 0;
  }
  for( int j = 0; j < PPI_comp.size(); ++j ){
    arma::uvec id = arma::find(PPI_un == PPI_comp(j));
    PPI_count(id) += 1;
  }
  
  if( max(TRN_count) > 1 && max(PPI_count) > 1 ){
    arma::uvec id_TRN = arma::find(TRN_count == max(TRN_count));
    arma::uvec id_PPI = arma::find(PPI_count == max(PPI_count));
    
    if( id_TRN.size() == 1 && id_PPI.size() == 1 ){
      arma::uvec id_TRN_comp = arma::find(TRN_comp == TRN_un[id_TRN(0)]);
      arma::uvec id_PPI_comp = arma::find(PPI_comp == PPI_un[id_PPI(0)]);
      
      arma::uvec id_inter = arma::intersect(id_TRN_comp, id_PPI_comp);
      
      return id_TRN_comp.size();
    }else{
      if( max(TRN_count) == 1 && max(PPI_count) == 1 ){
        return 1;
      }else{
        arma::mat size_inter(id_TRN.size(), id_PPI.size());
        for( int i = 0; i < id_TRN.size(); ++i ){
          for( int j = 0; j < id_PPI.size(); ++j ){
            arma::uvec id_TRN_comp = arma::find(TRN_comp == TRN_un[id_TRN(i)]);
            arma::uvec id_PPI_comp = arma::find(PPI_comp == PPI_un[id_PPI(j)]);
            
            arma::uvec id_inter = arma::intersect(id_TRN_comp, id_PPI_comp);
            size_inter(i, j) = id_inter.size();
          }
        }
        
        arma::uvec max_inter = find(size_inter == size_inter.max());
        int j = floor((max_inter(0) + 1)/size_inter.n_rows);
        int i = max_inter(0) - j*size_inter.n_rows;
        if( double(j*size_inter.n_rows) == double(max_inter(0) + 1) ){
          j -= 1;
          i = size_inter.n_rows - 1;
        }
        
        arma::uvec id_TRN_comp = arma::find(TRN_comp == TRN_un[id_TRN(i)]);
        arma::uvec id_PPI_comp = arma::find(PPI_comp == PPI_un[id_PPI(j)]);
        
        arma::uvec id_inter = arma::intersect(id_TRN_comp, id_PPI_comp);
        
        return id_TRN_comp.size();
      }
      
    }
    
    
  }else{
    return 1;
  }
  
}


/*Function to compute redundancy coupling for a single gene/protein in the multiplex*/
//[[Rcpp::export]]
Rcpp::List McgcCompleteGraph( arma::mat elTRN, arma::mat elPPI, int NumNodes ){
  Graph gTRN(NumNodes), gPPI(NumNodes);
  arma::vec TRN_comp(NumNodes), PPI_comp(NumNodes);
  
  for( int i = 0; i < elTRN.n_rows; ++i ){
    gTRN.addEdge(elTRN(i, 0), elTRN(i, 1));
  }
  
  for( int i = 0; i < elPPI.n_rows; ++i ){
    gPPI.addEdge(elPPI(i, 0), elPPI(i, 1));
  }
  
  gTRN.connectedComponents();
  gPPI.connectedComponents();
  
  for( int i = 0; i < NumNodes; ++i ){
    TRN_comp(i) = gTRN.component[i];
    PPI_comp(i) = gPPI.component[i];
  }
  
  // gTRN.clearnonLCC(TRN_comp);
  // gPPI.clearnonLCC(PPI_comp);
  // 
  // for( int i = 0; i < NumNodes; ++i ){
  //   TRN_comp(i) = gTRN.component[i];
  //   PPI_comp(i) = gPPI.component[i];
  // }
  
  int test = sizeTRNPPI(TRN_comp, PPI_comp);
  
  while( test == 0 ){
    delBridgeInter(gTRN, gPPI, TRN_comp, PPI_comp);
    test = sizeTRNPPI(TRN_comp, PPI_comp);
  }
  
  
  
  return Rcpp::List::create(Rcpp::Named("TRN")=TRN_comp,
                            Rcpp::Named("PPI")=PPI_comp);
}


void MCGCSize( Graph &gTRN, Graph &gPPI, arma::vec &TRN_comp, arma::vec &PPI_comp, int NumNodes ){
  int test = sizeTRNPPI(TRN_comp, PPI_comp);
  
  while( test == 0 ){
    delBridgeInter(gTRN, gPPI, TRN_comp, PPI_comp);
    test = sizeTRNPPI(TRN_comp, PPI_comp);
  }
}

/*Function to compute redundancy coupling for a single gene/protein in the multiplex*/
//[[Rcpp::export]]
Rcpp::List MCGCAttackCurve( arma::mat elTRN, arma::mat elPPI, int NumNodes, arma::vec NodeOrder ){
  Graph gTRN(NumNodes), gPPI(NumNodes);
  arma::vec TRN_comp(NumNodes), PPI_comp(NumNodes);
  arma::vec MCGC_curve = arma::ones<arma::vec>(NodeOrder.size());
  
  for( int i = 0; i < elTRN.n_rows; ++i ){
    gTRN.addEdge(elTRN(i, 0), elTRN(i, 1));
  }
  
  for( int i = 0; i < elPPI.n_rows; ++i ){
    gPPI.addEdge(elPPI(i, 0), elPPI(i, 1));
  }
  
  gTRN.connectedComponents();
  gPPI.connectedComponents();
  
  for( int i = 0; i < NumNodes; ++i ){
    TRN_comp(i) = gTRN.component[i];
    PPI_comp(i) = gPPI.component[i];
  }
  
  // gTRN.clearnonLCC(TRN_comp);
  // gPPI.clearnonLCC(PPI_comp);
  // 
  // for( int i = 0; i < NumNodes; ++i ){
  //   TRN_comp(i) = gTRN.component[i];
  //   PPI_comp(i) = gPPI.component[i];
  // }
  
  MCGCSize(gTRN, gPPI, TRN_comp, PPI_comp, NumNodes);
  // cout << "Size = " << sizeTRNPPIreturn(TRN_comp, PPI_comp) << endl;
  
  for( int v = 0; v < NodeOrder.size(); ++v ){
    // cout << "Attack Loop = " << v << " Node = " << NodeOrder(v) << endl;
    // arma::vec TRN_un = arma::unique(TRN_comp);
    // arma::vec PPI_un = arma::unique(PPI_comp);
    // cout << "TRN_un = " << TRN_un.size() << " PPI_un = " << PPI_un.size() << endl;
    
    if( gTRN.SizeNodeAdjList(NodeOrder(v)) == 0 && gPPI.SizeNodeAdjList(NodeOrder(v)) == 0 ){
      if( v != 0 ){
        MCGC_curve(v) = MCGC_curve(v - 1);
        continue;
      }
      
    }
    cout << "Attack Loop = " << v << " Node = " << NodeOrder(v) << endl;
    // gTRN.printSizeNodeAdjList(NodeOrder(v));
    // gPPI.printSizeNodeAdjList(NodeOrder(v));
    
    gTRN.clearVertex(NodeOrder(v));
    gPPI.clearVertex(NodeOrder(v));
    
    // gTRN.printSizeNodeAdjList(NodeOrder(v));
    // gPPI.printSizeNodeAdjList(NodeOrder(v));
    
    gTRN.connectedComponents();
    gPPI.connectedComponents();
    
    for( int i = 0; i < NumNodes; ++i ){
      TRN_comp(i) = gTRN.component[i];
      PPI_comp(i) = gPPI.component[i];
    }
    
    // gTRN.clearnonLCC(TRN_comp);
    // gPPI.clearnonLCC(PPI_comp);
    // 
    // cout << "Reached here 1" << endl;
    // for( int i = 0; i < NumNodes; ++i ){
    //   TRN_comp(i) = gTRN.component[i];
    //   PPI_comp(i) = gPPI.component[i];
    // }
    
    // cout << "Reached here 2" << endl;
    arma::vec node_size(NumNodes);
    for( int w = 0; w < NumNodes; ++w ){
      node_size(w) = gTRN.SizeNodeAdjList(w);
    }
    int max_size = node_size.max();
    if( max_size > 1 && sizeTRNPPIreturn(TRN_comp, PPI_comp) > 1 ){
      MCGCSize(gTRN, gPPI, TRN_comp, PPI_comp, NumNodes);
      // cout << "Size = " << sizeTRNPPIreturn(TRN_comp, PPI_comp) << endl;
      MCGC_curve(v) = sizeTRNPPIreturn(TRN_comp, PPI_comp);
    }
    
    
    // cout << "Reached here 2" << endl;
    
    if( MCGC_curve(v) == 1 ){
      break;
    }
  }
  
  
  // cout << "Reached here" << endl;
  return Rcpp::List::create(Rcpp::Named("TRN")=TRN_comp,
                            Rcpp::Named("PPI")=PPI_comp, 
                            Rcpp::Named("Attack_Curve")=MCGC_curve);
}

//[[Rcpp::export]]
arma::vec MultiplexDegree( arma::vec degTRN, arma::vec degPPI ){
  arma::vec NodeDeg(degTRN.size());
  for( int j = 0; j < degTRN.size(); ++j ){
    if( (degTRN(j) - degTRN.min())/(degTRN.max() - degTRN.min()) >= 
        (degPPI(j) - degPPI.min())/(degPPI.max() - degPPI.min()) ){
      NodeDeg(j) = (degTRN(j) - degTRN.min())/(degTRN.max() - degTRN.min());
    }else{
      NodeDeg(j) = (degPPI(j) - degPPI.min())/(degPPI.max() - degPPI.min());
    }
  }
  return NodeDeg;
}

//[[Rcpp::export]]
arma::vec MultiplexDegreeNew( arma::vec degTRN, arma::vec degPPI ){
  arma::vec NodeDeg(degTRN.size());
  for( int j = 0; j < degTRN.size(); ++j ){
    if( (degTRN(j) - degTRN.min())/(degTRN.max() - degTRN.min()) >= 
        (degPPI(j) - degPPI.min())/(degPPI.max() - degPPI.min()) ){
      NodeDeg(j) = (degTRN(j) - degTRN.min())/(degTRN.max() - degTRN.min());
    }else{
      NodeDeg(j) = (degPPI(j) - degPPI.min())/(degPPI.max() - degPPI.min());
    }
  }
  return NodeDeg;
}


//[[Rcpp::export]]
arma::vec shuffledTRN( string idfile, arma::mat &elTRN, arma::vec &degTRN, arma::vec degPPI, int b ){
  int NumNodes = degTRN.size();
  arma::uvec idTRN(NumNodes);
  arma::vec NodeDeg(degTRN.size());
  string readfile;
  readfile.append(idfile);
  readfile.append("/boot_");
  readfile.append(to_string(b + 1));
  ifstream myfile(readfile.c_str());
  if(myfile.is_open()){

    for( int j = 0; j < NumNodes; ++j ){
      string number;
      getline(myfile,number); //read number
      idTRN(j) = stoi(number.c_str()); //convert to integer
      // cout << "loop = " << j << " node = " << idTRN(j) << endl;
    }

  }
  
  arma::mat elTRNtemp = elTRN;
  for( int j = 0;j < idTRN.size(); ++j ){
    // cout << "succcess = " << idTRN(j) << endl;
    arma::uvec id1 = find(elTRNtemp.col(0) == idTRN(j));
    if( id1.size() != 0 ){
      for( int i = 0; i < id1.size(); ++i ){
        elTRN(id1(i), 0) = j;
      }
    }
    
    arma::uvec id2 = find(elTRNtemp.col(1) == idTRN(j));
    if( id2.size() != 0 ){
      for( int i = 0; i < id2.size(); ++i ){
        elTRN(id2(i), 1) = j;
      }
    }
  }
  
  degTRN = degTRN(idTRN);
  NodeDeg = MultiplexDegree(degTRN, degPPI);
  return NodeDeg;
  
}

//[[Rcpp::export]]
arma::mat shuffledTRNmat( string idfile, arma::mat elTRN, arma::vec degTRN, arma::vec degPPI, int b, 
                          arma::uvec idTRN){
  int NumNodes = degTRN.size();
  // arma::uvec idTRN(NumNodes);
  arma::vec NodeDeg(degTRN.size());
  // string readfile;
  // readfile.append(idfile);
  // readfile.append("/boot_");
  // readfile.append(to_string(b + 1));
  // ifstream myfile(readfile.c_str());
  // if(myfile.is_open()){
  //   
  //   for( int j = 0; j < NumNodes; ++j ){
  //     string number;
  //     getline(myfile,number); //read number
  //     idTRN(j) = stoi(number.c_str()); //convert to integer
  //     // cout << "loop = " << j << " node = " << idTRN(j) << endl;
  //   }
  //   
  // }
  
  arma::mat elTRNtemp = elTRN;
  for( int j = 0;j < idTRN.size(); ++j ){
    // cout << "succcess = " << idTRN(j) << endl;
    arma::uvec id1 = find(elTRNtemp.col(0) == idTRN(j));
    if( id1.size() != 0 ){
      for( int i = 0; i < id1.size(); ++i ){
        elTRN(id1(i), 0) = j;
      }
    }
    
    arma::uvec id2 = find(elTRNtemp.col(1) == idTRN(j));
    if( id2.size() != 0 ){
      for( int i = 0; i < id2.size(); ++i ){
        elTRN(id2(i), 1) = j;
      }
    }
  }
  
  degTRN = degTRN(idTRN);
  NodeDeg = MultiplexDegree(degTRN, degPPI);
  return elTRN;
  
}

//Parallel computation of MCGC attack curve for multiple repetitions of the attack
/*Input: 
 * elTRNorg - TRN edgelist
 * elPPI - PPI edgelist
 * NumNodes - number of genes/proteins in the multiplex
 * degTRNorg - outdegree for TRN
 * degPPI - degree for PPI
 * nboot - number of times to repeat the attack
 * nthreads - number of CPU cores for parallelization
 * outfile - location to write the size of MCGC at each step of the attack curve
 * seed - random seed
 * mode - whether to calculate attack curve for the original organism or shuffled multiplex. 0 (orginal multiplex). 1 (shuffled multiplex)
 * idfile - location where files with shuffled TRN ids are stored. used if mode = 1.
 */
/*Output:
 * success - success of the code
 */
//[[Rcpp::plugins(openmp)]]
//[[Rcpp::export]]
Rcpp::List MCGCAttackCurveParallel( arma::mat elTRNorg, arma::mat elPPI, int NumNodes, arma::vec degTRNorg, arma::vec degPPI, 
                                    int nboot, int nthreads, string outfile, int seed, int mode, string idfile){
  srand(seed);
  Progress p(nboot, true);
  omp_set_num_threads(nthreads);
#pragma omp parallel for

  for( int b = 0; b < nboot; ++b ){
    if ( ! Progress::check_abort() ) {
      arma::vec degTRN = degTRNorg;
      arma::mat elTRN = elTRNorg;
      arma::uvec idtemp(NumNodes);
      arma::vec NodeDeg(degTRN.size());
      int tid;
      if( mode == 1 ){
        arma::uvec idTRN(NumNodes);
        string readfile;
        readfile.append(idfile);
        readfile.append("/boot_");
        readfile.append(to_string(b + 1));
        ifstream myfile(readfile.c_str());
        if(myfile.is_open()){

          for( int j = 0; j < NumNodes; ++j ){
            string number;
            getline(myfile,number); //read number
            idTRN(j) = stoi(number.c_str()); //convert to integer
          }

        }
        arma::mat elTRNtemp = elTRN;
        for( int j = 0;j < idTRN.size(); ++j ){
          arma::uvec id1 = find(elTRNtemp.col(0) == idTRN(j));
          if( id1.size() != 0 ){
            for( int i = 0; i < id1.size(); ++i ){
              elTRN(id1(i), 0) = j;
            }
          }

          arma::uvec id2 = find(elTRNtemp.col(1) == idTRN(j));
          if( id2.size() != 0 ){
            for( int i = 0; i < id2.size(); ++i ){
              elTRN(id2(i), 1) = j;
            }
          }
        }

        degTRN = degTRN(idTRN);
        NodeDeg = MultiplexDegree(degTRN, degPPI);
      }else{
        NodeDeg = MultiplexDegree(degTRN, degPPI);
      }
      
      arma::uvec NodeOrder = sort_index(NodeDeg, "descend");
      arma::vec sort_deg = sort(NodeDeg, "descend");
      arma::vec NodeDeg_un = reverse(unique(NodeDeg));
      if( NodeDeg.size() != NodeDeg_un.size() ){
        for( int i = 0; i < NodeDeg_un.size(); ++i ){
          arma::uvec id_deg_unique = find(sort_deg == NodeDeg_un(i));
          if( id_deg_unique.size() > 1 ){
            arma::uvec rand_shuffle = shuffle(NodeOrder(id_deg_unique));
            for( int j = 0; j < id_deg_unique.size(); ++j ){
              NodeOrder(id_deg_unique(j)) = rand_shuffle(j);
            }
          }
        }
      }
      
      Graph gTRN(NumNodes), gPPI(NumNodes);
      arma::vec TRN_comp(NumNodes), PPI_comp(NumNodes);
      arma::vec MCGC_curve = arma::ones<arma::vec>(NodeOrder.size());
      
      for( int i = 0; i < elTRN.n_rows; ++i ){
        gTRN.addEdge(elTRN(i, 0), elTRN(i, 1));
      }
      
      for( int i = 0; i < elPPI.n_rows; ++i ){
        gPPI.addEdge(elPPI(i, 0), elPPI(i, 1));
      }
      
      gTRN.connectedComponents();
      gPPI.connectedComponents();
      
      for( int i = 0; i < NumNodes; ++i ){
        TRN_comp(i) = gTRN.component[i];
        PPI_comp(i) = gPPI.component[i];
      }
      
      MCGCSize(gTRN, gPPI, TRN_comp, PPI_comp, NumNodes);
      for( int v = 0; v < NumNodes; ++v ){
        if( gTRN.SizeNodeAdjList(NodeOrder(v)) == 0 && gPPI.SizeNodeAdjList(NodeOrder(v)) == 0 ){
          if( v != 0 ){
            MCGC_curve(v) = MCGC_curve(v - 1);
            continue;
          }
        }
        
        gTRN.clearVertex(NodeOrder(v));
        gPPI.clearVertex(NodeOrder(v));
        
        gTRN.connectedComponents();
        gPPI.connectedComponents();
        
        for( int i = 0; i < NumNodes; ++i ){
          TRN_comp(i) = gTRN.component[i];
          PPI_comp(i) = gPPI.component[i];
        }
        
        
        arma::vec node_size(NumNodes);
        for( int w = 0; w < NumNodes; ++w ){
          node_size(w) = gTRN.SizeNodeAdjList(w);
        }
        int max_size = node_size.max();
        if( max_size > 1 && sizeTRNPPIreturn(TRN_comp, PPI_comp) > 1 ){
          MCGCSize(gTRN, gPPI, TRN_comp, PPI_comp, NumNodes);
          
          MCGC_curve(v) = sizeTRNPPIreturn(TRN_comp, PPI_comp);
        }
        
        if( MCGC_curve(v) == 1 ){
          break;
        }
      }
      ofstream myfile;
      string writefile;
      writefile.append(outfile);
      writefile.append("/boot_");
      writefile.append(to_string(b + 1));
      myfile.open (writefile);
      for( int v = 0; v < NumNodes - 1; ++v ){
        myfile << MCGC_curve(v) << "\n";
      }
      myfile << MCGC_curve(NumNodes - 1);
      myfile.close();
      p.increment();
    }
    
  }
  return Rcpp::List::create(Rcpp::Named("Success")=1);
}

//[[Rcpp::export]]
arma::uvec testError( arma::vec degTRN, arma::vec degPPI, arma::uvec idnodes ){
  arma::vec NodeDeg = MultiplexDegree(degTRN, degPPI);
  
  arma::uvec NodeOrder = sort_index(NodeDeg(idnodes), "descend");
  arma::vec sort_deg = sort(NodeDeg(idnodes), "descend");
  arma::vec NodeDeg_un = reverse(unique(NodeDeg(idnodes)));
  if( idnodes.size() != NodeDeg_un.size() ){
    for( int i = 0; i < NodeDeg_un.size(); ++i ){
      arma::uvec id_deg_unique = find(sort_deg == NodeDeg_un(i));
      if( id_deg_unique.size() > 1 ){
        arma::uvec rand_shuffle = shuffle(NodeOrder(id_deg_unique));
        for( int j = 0; j < id_deg_unique.size(); ++j ){
          NodeOrder(id_deg_unique(j)) = rand_shuffle(j);
        }
      }
    }
  }
  return NodeOrder;
}


//[[Rcpp::plugins(openmp)]]
//[[Rcpp::export]]
Rcpp::List MCGCAttackCurveFunctionalGenesParallel( arma::mat elTRNorg, arma::mat elPPI, int NumNodes, arma::vec degTRNorg, arma::vec degPPI, 
                                    int nboot, int nthreads, string outfile, int seed, int mode, string idfile, 
                                    int NumFuncNodes, arma::uvec idnodesorg ){
  // arma::mat AttackCurves(NumNodes, nboot);
  srand(seed);
  Progress p(nboot, true);
  omp_set_num_threads(nthreads);
#pragma omp parallel for
  // {
  //   arma::mat AttackCurves_private(NumNodes, nboot);
  // #pragma omp for
  for( int b = 0; b < nboot; ++b ){
    // cout << "Boot Loop = " << b << endl;
    
    if ( ! Progress::check_abort() ) {
      arma::vec degTRN = degTRNorg;
      arma::mat elTRN = elTRNorg;
      arma::uvec idtemp(NumNodes);
      arma::vec NodeDeg(degTRN.size());
      
      int tid;
      
      arma::uvec idnodes(NumFuncNodes);
      if( mode == 1 ){
        string readfile;
        readfile.append(idfile);
        readfile.append("/boot_");
        readfile.append(to_string(b + 1));
        ifstream myinfile(readfile.c_str());
        if(myinfile.is_open()){
          
          for( int j = 0; j < NumFuncNodes; ++j ){
            string number;
            getline(myinfile,number); //read number
            idnodes(j) = stoi(number.c_str()); //convert to integer
            // cout << "loop = " << j << " node = " << idTRN(j) << endl;
          }
          
        }
      }else{
        idnodes = idnodesorg;
      }
      
      
      NodeDeg = MultiplexDegree(degTRN, degPPI);
      
      arma::uvec NodeOrder = sort_index(NodeDeg(idnodes), "descend");
      arma::vec sort_deg = sort(NodeDeg(idnodes), "descend");
      arma::vec NodeDeg_un = reverse(unique(NodeDeg(idnodes)));
      if( idnodes.size() != NodeDeg_un.size() ){
        for( int i = 0; i < NodeDeg_un.size(); ++i ){
          arma::uvec id_deg_unique = find(sort_deg == NodeDeg_un(i));
          if( id_deg_unique.size() > 1 ){
            arma::uvec rand_shuffle = shuffle(NodeOrder(id_deg_unique));
            for( int j = 0; j < id_deg_unique.size(); ++j ){
              NodeOrder(id_deg_unique(j)) = rand_shuffle(j);
            }
          }
        }
      }
      
      Graph gTRN(NumNodes), gPPI(NumNodes);
      arma::vec TRN_comp(NumNodes), PPI_comp(NumNodes);
      arma::vec MCGC_curve = arma::ones<arma::vec>(NodeOrder.size());
      
      for( int i = 0; i < elTRN.n_rows; ++i ){
        gTRN.addEdge(elTRN(i, 0), elTRN(i, 1));
      }
      
      for( int i = 0; i < elPPI.n_rows; ++i ){
        gPPI.addEdge(elPPI(i, 0), elPPI(i, 1));
      }
      
      gTRN.connectedComponents();
      gPPI.connectedComponents();
      
      for( int i = 0; i < NumNodes; ++i ){
        TRN_comp(i) = gTRN.component[i];
        PPI_comp(i) = gPPI.component[i];
      }
      
      // gTRN.clearnonLCC(TRN_comp);
      // gPPI.clearnonLCC(PPI_comp);
      // 
      // for( int i = 0; i < NumNodes; ++i ){
      //   TRN_comp(i) = gTRN.component[i];
      //   PPI_comp(i) = gPPI.component[i];
      // }
      
      MCGCSize(gTRN, gPPI, TRN_comp, PPI_comp, NumNodes);
      // cout << "Size = " << sizeTRNPPIreturn(TRN_comp, PPI_comp) << endl;
      // tid = omp_get_thread_num(); 
      // printf("Welcome to GFG from thread = %d\n", 
      //        tid);
      for( int v = 0; v < NumFuncNodes; ++v ){
        // cout << "Attack Loop = " << v << " Node = " << NodeOrder(v) << endl;
        // arma::vec TRN_un = arma::unique(TRN_comp);
        // arma::vec PPI_un = arma::unique(PPI_comp);
        // cout << "TRN_un = " << TRN_un.size() << " PPI_un = " << PPI_un.size() << endl;
        if( gTRN.SizeNodeAdjList(idnodes(NodeOrder(v))) == 0 && gPPI.SizeNodeAdjList(idnodes(NodeOrder(v))) == 0 ){
          if( v != 0 ){
            MCGC_curve(v) = MCGC_curve(v - 1);
            continue;
          }
        }
        
        // gTRN.printSizeNodeAdjList(NodeOrder(v));
        // gPPI.printSizeNodeAdjList(NodeOrder(v));
        
        gTRN.clearVertex(idnodes(NodeOrder(v)));
        gPPI.clearVertex(idnodes(NodeOrder(v)));
        
        // gTRN.printSizeNodeAdjList(NodeOrder(v));
        // gPPI.printSizeNodeAdjList(NodeOrder(v));
        
        gTRN.connectedComponents();
        gPPI.connectedComponents();
        
        for( int i = 0; i < NumNodes; ++i ){
          TRN_comp(i) = gTRN.component[i];
          PPI_comp(i) = gPPI.component[i];
        }
        
        // gTRN.clearnonLCC(TRN_comp);
        // gPPI.clearnonLCC(PPI_comp);
        // 
        // cout << "Reached here 1" << endl;
        // for( int i = 0; i < NumNodes; ++i ){
        //   TRN_comp(i) = gTRN.component[i];
        //   PPI_comp(i) = gPPI.component[i];
        // }
        
        
        arma::vec node_size(NumNodes);
        for( int w = 0; w < NumNodes; ++w ){
          node_size(w) = gTRN.SizeNodeAdjList(w);
        }
        int max_size = node_size.max();
        // cout << "Loop = " << v << " Reached here 1" << endl;
        if( max_size > 1 && sizeTRNPPIreturn(TRN_comp, PPI_comp) > 1 ){
          MCGCSize(gTRN, gPPI, TRN_comp, PPI_comp, NumNodes);
          
          MCGC_curve(v) = sizeTRNPPIreturn(TRN_comp, PPI_comp);
          // cout << "Size = " << sizeTRNPPIreturn(TRN_comp, PPI_comp) << endl;
        }
        
        
        // cout << "Loop = " << v << " Reached here 2" << endl;
        
        if( MCGC_curve(v) == 1 ){
          break;
        }
      }
      ofstream myfile;
      string writefile;
      writefile.append(outfile);
      writefile.append("/boot_");
      writefile.append(to_string(b + 1));
      myfile.open (writefile);
      for( int v = 0; v < NumFuncNodes - 1; ++v ){
        myfile << MCGC_curve(v) << "\n";
      }
      myfile << MCGC_curve(NumFuncNodes - 1);
      myfile.close();
      // tid = omp_get_thread_num(); 
      // printf("Welcome to GFG from thread = %d\n", 
      //        tid); 
      p.increment();
    }
    
    // for( int i = 0; i < NodeOrder.size(); ++i ){
    //   AttackCurves(b, i) = MCGC_curve(i);
    // }
    
  }
  
  // #pragma omp critical
  //   for( int b = 0; b < nboot; ++b ){
  //     for( int i = 0; i < NodeDeg.size(); ++i ){
  //       AttackCurves(b, i) = AttackCurves_private(b, i);
  //     }
  //   }
  // }
  
  
  // cout << "Reached here" << endl;
  return Rcpp::List::create(Rcpp::Named("Success")=1);
}

//[[Rcpp::plugins(openmp)]]
//[[Rcpp::export]]
Rcpp::List MCGCAttackCurveSampleParallel( arma::sp_mat adjTRN, arma::sp_mat adjPPI, int NumNodes, 
                                    int nboot, int nthreads, string outfile, int seed, int mode, string idfile, 
                                    arma::uvec idnodes ){
  // arma::mat AttackCurves(NumNodes, nboot);
  
 
  
  srand(seed);
  Progress p(nboot, true);
  // omp_set_num_threads(nthreads);
// #pragma omp parallel for
  // {
  //   arma::mat AttackCurves_private(NumNodes, nboot);
  // #pragma omp for
  for( int b = 0; b < nboot; ++b ){
    // cout << "Boot Loop = " << b << endl;
    
    // if ( ! Progress::check_abort() ) {
      arma::uvec idTRN(NumNodes);
      if( mode == 1 ){
        string readfile;
        readfile.append(idfile);
        readfile.append("/boot_");
        readfile.append(to_string(b + 1));
        ifstream myfile(readfile.c_str());
        if(myfile.is_open()){
          
          for( int j = 0; j < NumNodes; ++j ){
            string number;
            getline(myfile,number); //read number
            idTRN(j) = stoi(number.c_str()); //convert to integer
            // cout << "loop = " << j << " node = " << idTRN(j) << endl;
          }
          
        }
      }else{
        idTRN = idnodes;
      }
      
      arma::sp_mat adjtemp1(NumNodes, NumNodes);
      arma::sp_mat adjtemp2(NumNodes, NumNodes);
      arma::sp_mat::const_iterator it     = adjTRN.begin();
      arma::sp_mat::const_iterator it_end = adjTRN.end();
      for(; it != it_end; ++it){
        if( adjTRN(it.row(), it.col()) == 1 ){
          int id1 = 0;
          int id2 = 0;
          int test1 = 0;
          int test2 = 0;
          for( int j = 0; j < idTRN.size(); j++ ){
            if( int(it.row()) == int(idTRN(j)) ){
              test1 = 1;
              id1 = j;
            }
            if( int(it.col()) == int(idTRN(j)) ){
              test2 = 1;
              id2 = j;
            }
            if( test1 == 1 && test2 == 1 ){
              break;
            }
          }
          if( test1 == 1 && test2 == 1 ){
            adjtemp1(id1, id2) = 1;
          }
        }
        
      }
      
      it = adjPPI.begin();
      it_end = adjPPI.end();
      for(; it != it_end; ++it){
        if( adjPPI(it.row(), it.col()) == 1 ){
          int id1 = 0;
          int id2 = 0;
          int test1 = 0;
          int test2 = 0;
          for( int j = 0; j < idTRN.size(); j++ ){
            if( int(it.row()) == idTRN(j) ){
              test1 = 1;
              id1 = j;
            }
            if( int(it.col()) == idTRN(j) ){
              test2 = 1;
              id2 = j;
            }
            if( test1 == 1 && test2 == 1 ){
              break;
            }
          }
          if( test1 == 1 && test2 == 1 ){
            adjtemp2(id1, id2) = 1;
          }
        }
        
      }
      
      arma::vec degTRN(NumNodes);
      degTRN.fill(0);
      arma::vec degPPI(NumNodes);
      degPPI.fill(0);
      
      
      
      it = adjtemp1.begin();
      it_end = adjtemp1.end();
      for(; it != it_end; ++it){
        degTRN(it.row()) += 1;
      }
      
      
      it = adjtemp2.begin();
      it_end = adjtemp2.end();
      for(; it != it_end; ++it){
        degPPI(it.row()) += 1;
      }
      
      
      // arma::vec degTRN = degTRNorg;
      arma::uvec idtemp(NumNodes);
      arma::vec NodeDeg(degTRN.size());
      
      
      NodeDeg = MultiplexDegreeNew(degTRN, degPPI);
      
      int tid;
      
      arma::uvec NodeOrder = sort_index(NodeDeg, "descend");
      arma::vec sort_deg = sort(NodeDeg, "descend");
      arma::vec NodeDeg_un = reverse(unique(NodeDeg));
      if( NodeDeg.size() != NodeDeg_un.size() ){
        for( int i = 0; i < NodeDeg_un.size(); ++i ){
          arma::uvec id_deg_unique = find(sort_deg == NodeDeg_un(i));
          if( id_deg_unique.size() > 1 ){
            arma::uvec rand_shuffle = shuffle(NodeOrder(id_deg_unique));
            for( int j = 0; j < id_deg_unique.size(); ++j ){
              NodeOrder(id_deg_unique(j)) = rand_shuffle(j);
            }
          }
        }
      }
      
      Graph gTRN(NumNodes), gPPI(NumNodes);
      arma::vec TRN_comp(NumNodes), PPI_comp(NumNodes);
      arma::vec MCGC_curve = arma::ones<arma::vec>(NodeOrder.size());
      
      
      it  = adjtemp1.begin();
      it_end = adjtemp1.end();
      for(; it != it_end; ++it){
        if( adjtemp1(it.col(), it.row()) != 1 ){
          gTRN.addEdge(it.row(), it.col());
        }else{
          if( (it.col())*NumNodes + it.row() + 1 < (it.row())*NumNodes + it.col() ){
            gTRN.addEdge(it.row(), it.col());
          }
        }
        
      }
      
      it  = adjtemp2.begin();
      it_end = adjtemp2.end();
      for(; it != it_end; ++it){
        if( adjtemp2(it.col(), it.row()) != 1 ){
          gPPI.addEdge(it.row(), it.col());
        }else{
          if( (it.col())*NumNodes + it.row() + 1 < (it.row())*NumNodes + it.col() ){
            gPPI.addEdge(it.row(), it.col());
          }
        }
        
      }
      
     
      
      // for( int i = 0; i < elTRN.n_rows; ++i ){
      //   gTRN.addEdge(elTRN(i, 0), elTRN(i, 1));
      // }
      // 
      // for( int i = 0; i < elPPI.n_rows; ++i ){
      //   gPPI.addEdge(elPPI(i, 0), elPPI(i, 1));
      // }
      
      gTRN.connectedComponents();
      gPPI.connectedComponents();
      
      for( int i = 0; i < NumNodes; ++i ){
        TRN_comp(i) = gTRN.component[i];
        PPI_comp(i) = gPPI.component[i];
      }
      
      // gTRN.clearnonLCC(TRN_comp);
      // gPPI.clearnonLCC(PPI_comp);
      // 
      // for( int i = 0; i < NumNodes; ++i ){
      //   TRN_comp(i) = gTRN.component[i];
      //   PPI_comp(i) = gPPI.component[i];
      // }
      
      MCGCSize(gTRN, gPPI, TRN_comp, PPI_comp, NumNodes);
      // cout << "Size = " << sizeTRNPPIreturn(TRN_comp, PPI_comp) << endl;
      // tid = omp_get_thread_num(); 
      // printf("Welcome to GFG from thread = %d\n", 
      //        tid);
      for( int v = 0; v < NumNodes; ++v ){
        // cout << "Attack Loop = " << v << " Node = " << NodeOrder(v) << endl;
        // arma::vec TRN_un = arma::unique(TRN_comp);
        // arma::vec PPI_un = arma::unique(PPI_comp);
        // cout << "TRN_un = " << TRN_un.size() << " PPI_un = " << PPI_un.size() << endl;
        
        if( gTRN.SizeNodeAdjList(NodeOrder(v)) == 0 && gPPI.SizeNodeAdjList(NodeOrder(v)) == 0 ){
          if( v != 0 ){
            MCGC_curve(v) = MCGC_curve(v - 1);
            continue;
          }
        }
        
        // gTRN.printSizeNodeAdjList(NodeOrder(v));
        // gPPI.printSizeNodeAdjList(NodeOrder(v));
        
        gTRN.clearVertex(NodeOrder(v));
        gPPI.clearVertex(NodeOrder(v));
        
        // gTRN.printSizeNodeAdjList(NodeOrder(v));
        // gPPI.printSizeNodeAdjList(NodeOrder(v));
        
        gTRN.connectedComponents();
        gPPI.connectedComponents();
        
        for( int i = 0; i < NumNodes; ++i ){
          TRN_comp(i) = gTRN.component[i];
          PPI_comp(i) = gPPI.component[i];
        }
        
        // gTRN.clearnonLCC(TRN_comp);
        // gPPI.clearnonLCC(PPI_comp);
        // 
        // cout << "Reached here 1" << endl;
        // for( int i = 0; i < NumNodes; ++i ){
        //   TRN_comp(i) = gTRN.component[i];
        //   PPI_comp(i) = gPPI.component[i];
        // }
        
        
        arma::vec node_size(NumNodes);
        for( int w = 0; w < NumNodes; ++w ){
          node_size(w) = gTRN.SizeNodeAdjList(w);
        }
        int max_size = node_size.max();
        // cout << "Loop = " << v << " Reached here 1" << endl;
        if( max_size > 1 && sizeTRNPPIreturn(TRN_comp, PPI_comp) > 1 ){
          MCGCSize(gTRN, gPPI, TRN_comp, PPI_comp, NumNodes);
          
          MCGC_curve(v) = sizeTRNPPIreturn(TRN_comp, PPI_comp);
          // cout << "Size = " << sizeTRNPPIreturn(TRN_comp, PPI_comp) << endl;
        }
        
        
        // cout << "Loop = " << v << " Reached here 2" << endl;
        
        if( MCGC_curve(v) == 1 ){
          break;
        }
      }
      ofstream myfile;
      string writefile;
      writefile.append(outfile);
      writefile.append("/boot_");
      writefile.append(to_string(b + 1));
      myfile.open (writefile);
      for( int v = 0; v < NumNodes - 1; ++v ){
        myfile << MCGC_curve(v) << "\n";
      }
      myfile << MCGC_curve(NumNodes - 1);
      myfile.close();
      // tid = omp_get_thread_num(); 
      // printf("Welcome to GFG from thread = %d\n", 
      //        tid); 
      // p.increment();
    // }
    
    // for( int i = 0; i < NodeOrder.size(); ++i ){
    //   AttackCurves(b, i) = MCGC_curve(i);
    // }
    
  }
  
  // #pragma omp critical
  //   for( int b = 0; b < nboot; ++b ){
  //     for( int i = 0; i < NodeDeg.size(); ++i ){
  //       AttackCurves(b, i) = AttackCurves_private(b, i);
  //     }
  //   }
  // }
  
  
  // cout << "Reached here" << endl;
  return Rcpp::List::create(Rcpp::Named("Success")=1);
}


//[[Rcpp::export]]
arma::uvec Sort_ID( arma::vec vals ){
  arma::uvec id = sort_index(vals);
  return id;
}
```
