## Supplementary material for "Internetwork connectivity of molecular networks across species of life": Code: Figure_2_Paper.R

```
##set seed for reproducibility
set.seed(1)

##Load packages and functions
library(Rcpp)
library(Matrix)
library(igraph)
library(philentropy)
library(ggplot2)
library(ggpubr)
library(cowplot)
Rcpp::sourceCpp('utils.cpp')

##Load multiplex network data
load("NetworksMultiplex.RData")

##Name of species
Organisms <- c("H. pylori", "M.tb",        "E. coli",     "Yeast",       "C. elegans",  "Fly", 
               "A. thaliana", "Mouse", "Human")

#######################################################Fig. 2##################################################################
##For Fig. 2, BioGRID PPI networks are used.
##Define variables to store plots for degree-degree (Plot.deg) and redundancy (Plot.red) coupling.
Plot.deg <- Plot.red <- list()

##Define variables to store redundancy coupling for the organisms (influence.red) and the shuffled null model 
##(influence.red.shuffled).
influence.red <- influence.red.shuffled <- list()

##Number of times to compute redundancy coupling for the shuffled null model.
nboot <- 1000

##Compute redundancy coupling for each gene/protein for each organism.
for( i in c(4, 6, 8, 9) ){
  
  ##Print name of the organism.
  message(paste("Organism = ", Organisms[i], sep = ""))
  
  ##Get organism TRN and PPI networks from multiplex network object.
  adj.TRN <- NetworksMultiplex$BioGrid[[i]]$TRN
  adj.PPI <- NetworksMultiplex$BioGrid[[i]]$PPI
  
  ##Compute redundancy coupling for each gene/protein in the multiplex
  ##count_common_edges_among_nodes -- Function defined in 'utils.cpp'. Loaded by compiling 'utils.cpp' (line 12 above).
  influence.red[[i]] <- count_common_edges_among_nodes(adj.TRN*adj.PPI, c(1:ncol(adj.TRN)))
  
  
  ##Compute redundancy coupling for each gene/protein in the shuffled multiplex
  influence.red.shuffled[[i]] <- list()
  for( j in 1:nboot ){
    cat("Organism = ", Organisms[i], " Loop = ", j, "\n")
    id <- sample(1:ncol(adj.TRN))
    influence.red.shuffled[[i]][[j]] <- ((table(count_common_edges_among_nodes(adj.TRN[id, id]*adj.PPI, c(1:ncol(adj.TRN))))))
  }
  
  count <- count + 1
  save(influence.red, file = "Influence_red_Organisms_BioGRID_Paper.RData")
  save(influence.red.shuffled, file = "Influence_red_Shuffled_BioGRID_Paper.RData")
  
}


load("Influence_red_Organisms_BioGRID_Paper.RData")
load("Influence_red_Shuffled_BioGRID_Paper.RData")
count <- 1
nboot <- 1000
Plot.deg <- Plot.red <- list()
for( i in c(4, 6, 8, 9) ){
  cat("Organism = ", Organisms[i], "\n")
  adj.TRN <- NetworksMultiplex$BioGrid[[i]]$TRN
  adj.PPI <- NetworksMultiplex$BioGrid[[i]]$PPI
  
  
  ##Degree-degree coupling plot
  id <- 1:ncol(adj.TRN)
  dat.plot <- data.frame(x = 1 + rowSums(adj.TRN)[id],
                         y = 1 + colSums(adj.PPI)[id],
                         Type = rep(Organisms[i], ncol(adj.TRN[id, id])))
  data.cor.tmp <- data.frame(r = signif(cor(dat.plot$x, dat.plot$y, method = "spearman"), 4), 
                             p = signif(cor.test(dat.plot$x, dat.plot$y, method = "spearman")$p.value, 4), 
                             label = paste("C[D]", " = ", signif(cor(dat.plot$x, dat.plot$y, method = "spearman"), 4), 
                                           ", p = ", signif(cor.test(dat.plot$x, dat.plot$y, method = "spearman")$p.value, 4), sep = ""))
  
  data.cor <- paste(" = ", 
                    signif(cor(dat.plot$x, dat.plot$y, method = "spearman"), 4), 
                    ", p = ", signif(cor.test(dat.plot$x, dat.plot$y, 
                                              method = "spearman")$p.value, 4), sep = "")
  p <- ggscatter(dat.plot, x = "x", y = "y", 
                 fill = "black",          
                 color = "black",
                 # palette = c("red", "gray", "red", "black"),
                 sort.by.groups = FALSE,     
                 # x.text.angle = 50,          # Rotate vertically x axis texts
                 ylab = "K",
                 xlab = expression("k"["out"]),
                 # legend = "bottom",
                 add = "reg.line",
                 conf.int = TRUE,                       
                 add.params = list(color = "green",
                                   fill = "lightgray"),
                 legend.title = "",
                 y.text.angle = 50,
                 rotate = F,
                 font.x = 14, 
                 font.y = 14,
                 font.tickslab = 12,
                 size = 0.9,
                 point.size = 0.1,
                 font.label = 12,
                 theme = theme_pubr()
  )
  p <- p + xscale("log10", T) + yscale("log10", T) + 
    ggtitle(bquote(C[D] ~ .(data.cor))) +
    theme(title = element_text(size = 12, color = "black"), 
          axis.title = element_text(size = 16, colour = "black"), 
          text = element_text(size = 16, colour = "black"), 
          legend.text = element_text(size = 16, color = "black")
          , legend.position = "bottom")
  Plot.deg[[count]] <- p
  
  
  
  ##Redundancy coupling Organism
  tbl.inf.red <- table(influence.red[[i]])
  max.red <- max(as.numeric(names(tbl.inf.red)))
  max.red.shuffled <- rep(0, nboot)
  for( j in 1:nboot ){
    max.red.shuffled[j] <- max(as.numeric(names(influence.red.shuffled[[i]][[j]])))
  }
  max.red <- max(c(max.red, max.red.shuffled))
  red.dist <- rep(0, max.red + 1)
  red.dist[as.numeric(names(tbl.inf.red)) + 1] <- as.numeric(tbl.inf.red)
  
  ##Redundancy coupling shuffled multiplex
  red.dist.shuffled <- matrix(0, max.red + 1, nboot)
  for( j in 1:nboot ){
    red.dist.shuffled[as.numeric(names(influence.red.shuffled[[i]][[j]])) + 1, j] <-
      as.numeric(influence.red.shuffled[[i]][[j]])
  }
  red.dist.shuffled <- apply(red.dist.shuffled, 1, median)
  
  ##Jensen Shannon divergence between organism and shuffled multiplex redundancy coupling
  JSD.vec <- rep(0, nboot)
  for( j in 1:nboot ){
    cat("Loop = ", j, "\n")
    red.dist.tmp <- rep(0, max.red + 1)
    red.dist.tmp[as.numeric(names(influence.red.shuffled[[i]][[j]])) + 1] <- 
      as.numeric(influence.red.shuffled[[i]][[j]])
    JSD.vec[j] <- JSD(rbind(red.dist.tmp, red.dist))
  }
  
  id1 <- which(red.dist == 0)[1] - 1
  id2 <- which(red.dist.shuffled == 0)[1] - 1
  id <- max(c(id1, id2))
  
  ##Redundancy coupling plots  
  data.plot <- data.frame(x = rep(0:(id - 1), 2), 
                          y = log10(1 + c(red.dist[1:id], red.dist.shuffled[1:id])), 
                          Type = c(rep("Original", length(1:id)), 
                                   rep("Shuffled", length(1:id))))
  
  
  data.plot <- data.plot %>%
    mutate(y = ifelse(Type == "Original",
                      y,
                      -1*y))
  breaks_values_labs <- sign(pretty(data.plot$y))*(10^abs(pretty(data.plot$y)))
  breaks_values <- sign(pretty(data.plot$y))*log10(10^abs(pretty(data.plot$y)) + 1)
  data.JSD <- data.frame(x = signif(mean(JSD.vec), 6), y = signif(mean(JSD.vec) - sd(JSD.vec), 6), 
                         z = signif(mean(JSD.vec) + sd(JSD.vec), 6))
  p <- ggbarplot(data.plot, x = "x", y = "y", 
                 fill = "Type",         
                 color = "Type",
                 palette = c("red", "blue", "red", "black"),
                 sort.by.groups = FALSE,   
                 # x.text.angle = 50,      
                 ylab = "Number of genes/proteins",
                 xlab = expression("C"["R"]),
                 # legend = "bottom",
                 legend.title = "",
                 y.text.angle = 50,
                 rotate = T,
                 font.x = 14, 
                 font.y = 14,
                 font.tickslab = 12,
                 size = 0.9,
                 point.size = 0.1,
                 font.label = 12,
                 stat = "identity",
                 theme = theme_pubr()
  ) + rotate_x_text(angle = 45) +
    ggtitle(paste("JSD = ", data.JSD$x, " (", data.JSD$y, ", ", data.JSD$z, ")", sep = ""))+ geom_hline(yintercept = 0) +
    scale_y_continuous(breaks = breaks_values,
                       labels = abs(breaks_values_labs))
  p <- p + 
    theme(title = element_text(size = 12, color = "black"), 
          axis.title = element_text(size = 16, colour = "black"), 
          legend.text = element_text(size = 16, color = "black")
          , legend.position = "bottom")
  
  Plot.red[[count]] <- p
  count <- count + 1
}
plot.1 <- plot_grid(Plot.deg[[1]], Plot.deg[[2]], Plot.deg[[3]], Plot.deg[[4]], 
                    Plot.red[[1]], Plot.red[[2]], Plot.red[[3]], Plot.red[[4]], 
                    ncol = 4, labels = "AUTO", label_size = 16, 
                    align = "v")
save_plot(paste("Figure_2_degree_degree_redundancy.png", sep = ""), 
          plot.1, base_height = 8, base_width = 16)

###################HINT##############################
##Name of species
Organisms <- c("H. pylori", "M.tb",        "E. coli",     "Yeast",       "C. elegans",  "Fly", 
               "A. thaliana", "Mouse", "Human")
##For Fig. 2, HINT PPI networks are used.
##Define variables to store plots for degree-degree (Plot.deg) and redundancy (Plot.red) coupling.
Plot.deg <- Plot.red <- list()

##Define variables to store redundancy coupling for the organisms (influence.red) and the shuffled null model 
##(influence.red.shuffled).
influence.red <- influence.red.shuffled <- list()

##Number of times to compute redundancy coupling for the shuffled null model.
nboot <- 1000

##Compute redundancy coupling for each gene/protein for each organism.
for( i in c(4, 5, 6, 8, 9) ){
  
  ##Print name of the organism.
  message(paste("Organism = ", Organisms[i], sep = ""))
  
  ##Get organism TRN and PPI networks from multiplex network object.
  adj.TRN <- NetworksMultiplex$HINT[[i]]$TRN
  adj.PPI <- NetworksMultiplex$HINT[[i]]$PPI
  
  ##Compute redundancy coupling for each gene/protein in the multiplex
  ##count_common_edges_among_nodes -- Function defined in 'utils.cpp'. Loaded by compiling 'utils.cpp' (line 12 above).
  influence.red[[i]] <- count_common_edges_among_nodes(adj.TRN*adj.PPI, c(1:ncol(adj.TRN)))
  
  
  ##Compute redundancy coupling for each gene/protein in the shuffled multiplex
  influence.red.shuffled[[i]] <- list()
  for( j in 1:nboot ){
    cat("Organism = ", Organisms[i], " Loop = ", j, "\n")
    id <- sample(1:ncol(adj.TRN))
    influence.red.shuffled[[i]][[j]] <- ((table(count_common_edges_among_nodes(adj.TRN[id, id]*adj.PPI, c(1:ncol(adj.TRN))))))
  }
  
  count <- count + 1
  save(influence.red, file = "Influence_red_Organisms_HINT_Paper.RData")
  save(influence.red.shuffled, file = "Influence_red_Shuffled_HINT_Paper.RData")
  
}


load("Influence_red_Organisms_HINT_Paper.RData")
load("Influence_red_Shuffled_HINT_Paper.RData")
count <- 1
nboot <- 1000
Plot.deg <- Plot.red <- list()
for( i in c(4, 5, 6, 8, 9) ){
  cat("Organism = ", Organisms[i], "\n")
  adj.TRN <- NetworksMultiplex$HINT[[i]]$TRN
  adj.PPI <- NetworksMultiplex$HINT[[i]]$PPI
  
  
  ##Degree-degree coupling plot
  id <- 1:ncol(adj.TRN)
  dat.plot <- data.frame(x = 1 + rowSums(adj.TRN)[id],
                         y = 1 + colSums(adj.PPI)[id],
                         Type = rep(Organisms[i], ncol(adj.TRN[id, id])))
  data.cor.tmp <- data.frame(r = signif(cor(dat.plot$x, dat.plot$y, method = "spearman"), 4), 
                             p = signif(cor.test(dat.plot$x, dat.plot$y, method = "spearman")$p.value, 4), 
                             label = paste("C[D]", " = ", signif(cor(dat.plot$x, dat.plot$y, method = "spearman"), 4), 
                                           ", p = ", signif(cor.test(dat.plot$x, dat.plot$y, method = "spearman")$p.value, 4), sep = ""))
  
  data.cor <- paste(" = ", 
                    signif(cor(dat.plot$x, dat.plot$y, method = "spearman"), 4), 
                    ", p = ", signif(cor.test(dat.plot$x, dat.plot$y, 
                                              method = "spearman")$p.value, 4), sep = "")
  p <- ggscatter(dat.plot, x = "x", y = "y", 
                 fill = "black",          
                 color = "black",
                 # palette = c("red", "gray", "red", "black"),
                 sort.by.groups = FALSE,     
                 # x.text.angle = 50,          # Rotate vertically x axis texts
                 ylab = "K",
                 xlab = expression("k"["out"]),
                 # legend = "bottom",
                 add = "reg.line",
                 conf.int = TRUE,                       
                 add.params = list(color = "green",
                                   fill = "lightgray"),
                 legend.title = "",
                 y.text.angle = 50,
                 rotate = F,
                 font.x = 14, 
                 font.y = 14,
                 font.tickslab = 12,
                 size = 0.9,
                 point.size = 0.1,
                 font.label = 12,
                 theme = theme_pubr()
  )
  p <- p + xscale("log10", T) + yscale("log10", T) + 
    ggtitle(bquote(C[D] ~ .(data.cor))) +
    theme(title = element_text(size = 12, color = "black"), 
          axis.title = element_text(size = 16, colour = "black"), 
          text = element_text(size = 16, colour = "black"), 
          legend.text = element_text(size = 16, color = "black")
          , legend.position = "bottom")
  Plot.deg[[count]] <- p
  
  
  
  ##Redundancy coupling Organism
  tbl.inf.red <- table(influence.red[[i]])
  max.red <- max(as.numeric(names(tbl.inf.red)))
  max.red.shuffled <- rep(0, nboot)
  for( j in 1:nboot ){
    max.red.shuffled[j] <- max(as.numeric(names(influence.red.shuffled[[i]][[j]])))
  }
  max.red <- max(c(max.red, max.red.shuffled))
  red.dist <- rep(0, max.red + 1)
  red.dist[as.numeric(names(tbl.inf.red)) + 1] <- as.numeric(tbl.inf.red)
  
  ##Redundancy coupling shuffled multiplex
  red.dist.shuffled <- matrix(0, max.red + 1, nboot)
  for( j in 1:nboot ){
    red.dist.shuffled[as.numeric(names(influence.red.shuffled[[i]][[j]])) + 1, j] <-
      as.numeric(influence.red.shuffled[[i]][[j]])
  }
  red.dist.shuffled <- apply(red.dist.shuffled, 1, median)
  
  ##Jensen Shannon divergence between organism and shuffled multiplex redundancy coupling
  JSD.vec <- rep(0, nboot)
  for( j in 1:nboot ){
    cat("Loop = ", j, "\n")
    red.dist.tmp <- rep(0, max.red + 1)
    red.dist.tmp[as.numeric(names(influence.red.shuffled[[i]][[j]])) + 1] <- 
      as.numeric(influence.red.shuffled[[i]][[j]])
    JSD.vec[j] <- JSD(rbind(red.dist.tmp, red.dist))
  }
  
  id1 <- which(red.dist == 0)[1] - 1
  id2 <- which(red.dist.shuffled == 0)[1] - 1
  id <- max(c(id1, id2))
  
  ##Redundancy coupling plots  
  data.plot <- data.frame(x = rep(0:(id - 1), 2), 
                          y = log10(1 + c(red.dist[1:id], red.dist.shuffled[1:id])), 
                          Type = c(rep("Original", length(1:id)), 
                                   rep("Shuffled", length(1:id))))
  
  
  data.plot <- data.plot %>%
    mutate(y = ifelse(Type == "Original",
                      y,
                      -1*y))
  breaks_values_labs <- sign(pretty(data.plot$y))*(10^abs(pretty(data.plot$y)))
  breaks_values <- sign(pretty(data.plot$y))*log10(10^abs(pretty(data.plot$y)) + 1)
  data.JSD <- data.frame(x = signif(mean(JSD.vec), 6), y = signif(mean(JSD.vec) - sd(JSD.vec), 6), 
                         z = signif(mean(JSD.vec) + sd(JSD.vec), 6))
  p <- ggbarplot(data.plot, x = "x", y = "y", 
                 fill = "Type",         
                 color = "Type",
                 palette = c("red", "blue", "red", "black"),
                 sort.by.groups = FALSE,   
                 # x.text.angle = 50,      
                 ylab = "Number of genes/proteins",
                 xlab = expression("C"["R"]),
                 # legend = "bottom",
                 legend.title = "",
                 y.text.angle = 50,
                 rotate = T,
                 font.x = 14, 
                 font.y = 14,
                 font.tickslab = 12,
                 size = 0.9,
                 point.size = 0.1,
                 font.label = 12,
                 stat = "identity",
                 theme = theme_pubr()
  ) + rotate_x_text(angle = 45) +
    ggtitle(paste("JSD = ", data.JSD$x, " (", data.JSD$y, ", ", data.JSD$z, ")", sep = ""))+ geom_hline(yintercept = 0) +
    scale_y_continuous(breaks = breaks_values,
                       labels = abs(breaks_values_labs))
  p <- p + 
    theme(title = element_text(size = 12, color = "black"), 
          axis.title = element_text(size = 16, colour = "black"), 
          legend.text = element_text(size = 16, color = "black")
          , legend.position = "bottom")
  
  Plot.red[[count]] <- p
  count <- count + 1
}
plot.1 <- plot_grid(Plot.deg[[1]], Plot.deg[[2]], Plot.deg[[3]], Plot.deg[[4]], 
                    Plot.red[[1]], Plot.red[[2]], Plot.red[[3]], Plot.red[[4]], 
                    ncol = 4, labels = "AUTO", label_size = 16, 
                    align = "v")
save_plot(paste("Figure_2_degree_degree_redundancy_HINT.png", sep = ""), 
          plot.1, base_height = 8, base_width = 16)
```
