## Supplementary material for "Internetwork connectivity of molecular networks across species of life": Code: Figure_3_Paper.R

```
##Figure 3
##set seed for reproducibility
set.seed(1)

##Load packages and functions
library(Rcpp)
library(Matrix)
library(igraph)
library(ggplot2)
library(ggpubr)
library(cowplot)
Rcpp::sourceCpp('utils.cpp')
Rcpp::sourceCpp("Attack_functions_new_cpp.cpp")

##Load multiplex network data
load("NetworksMultiplex.RData")

##Name of species
Organisms <- c("H. pylori", "M.tb",        "E. coli",     "Yeast",       "C. elegans",  "Fly", 
               "A. thaliana", "Mouse", "Human")
Organisms_file <- c("H_pylori", "M_tb",        "E_coli",     "Yeast",       "C_elegans",  "Fly", 
                    "A_thaliana", "Mouse", "Human")

##Number of parallel threads
nthreads <- 10

##Number of times to repeat the attack on a multiplex
nboot <- 1000

##Random seed
seed <- 123


##Calculate MCGC attack curves for organisms with BioGRID PPI
for( i in c(4, 6, 8, 9) ){
  
  ##Print name of the organism.
  message(paste("Organism = ", Organisms[i], sep = ""))
  
  ##Get organism TRN and PPI networks from multiplex network object.
  adj.TRN <- NetworksMultiplex$BioGrid[[i]]$TRN
  adj.PPI <- NetworksMultiplex$BioGrid[[i]]$PPI
  id1 <- 1:ncol(adj.TRN)
  id2 <- 1:ncol(adj.TRN)
  adj1 <- adj.TRN[id1, id1]
  adj2 <- adj.PPI[id2, id2]
  
  ##TRN and PPI edgelist
  elTRN <- get.edgelist(graph_from_adjacency_matrix(adj1, "undirected")) - 1
  elPPI <- get.edgelist(graph_from_adjacency_matrix(adj2, "undirected")) - 1
  
  ##Make organism specific folder to store the attack curves
  dir.org <- paste(getwd(), "/", Organisms_file[i], "_BioGrid", sep = "")
  dir.create(dir.org, showWarnings = F)
  
  ##Compute attack curves
  system.time(test <- MCGCAttackCurveParallel(elTRN, elPPI, ncol(adj.TRN), rowSums(adj.TRN), 
                                                        colSums(adj.PPI), nboot, nthreads, dir.org, seed, 0, dir.org))
}

##MCGC attack curve list object
MCGC.organism <- list()

##Read MCGC attack curves for Organisms with BioGRID PPI
MCGC.organism$BioGRID <- list()
for( i in c(4, 6, 8, 9) ){
  message(paste("Organism = ", Organisms[i], sep = ""))
  dir.org <- paste(getwd(), "/", Organisms_file[i], "_BioGrid", sep = "")
  adj.TRN <- NetworksMultiplex$BioGrid[[i]]$TRN
  adj.PPI <- NetworksMultiplex$BioGrid[[i]]$PPI
  MCGC.organism$BioGRID[[i]] <- matrix(0, nboot, ncol(adj.TRN))
  for( b in 1:nboot ){
    MCGC.organism$BioGRID[[i]][b, ] <- c(unlist(read.table(paste(dir.org, "/boot_", b, sep = ""), sep = "\t", header = F)))
  }
}
save(MCGC.organism, file = "MCGC_Organism_attack_curves.RData")


##Calculate MCGC attack curves for organisms with HINT PPI
for( i in c(4, 5, 6, 8, 9) ){
  
  ##Print name of the organism.
  message(paste("Organism = ", Organisms[i], sep = ""))
  
  ##Get organism TRN and PPI networks from multiplex network object.
  adj.TRN <- NetworksMultiplex$HINT[[i]]$TRN
  adj.PPI <- NetworksMultiplex$HINT[[i]]$PPI
  id1 <- 1:ncol(adj.TRN)
  id2 <- 1:ncol(adj.TRN)
  adj1 <- adj.TRN[id1, id1]
  adj2 <- adj.PPI[id2, id2]
  
  ##TRN and PPI edgelist
  elTRN <- get.edgelist(graph_from_adjacency_matrix(adj1, "undirected")) - 1
  elPPI <- get.edgelist(graph_from_adjacency_matrix(adj2, "undirected")) - 1
  
  ##Make organism specific folder to store the attack curves
  dir.org <- paste(getwd(), "/", Organisms_file[i], "_HINT", sep = "")
  dir.create(dir.org, showWarnings = F)
  
  system.time(test <- MCGCAttackCurveParallel(elTRN, elPPI, ncol(adj.TRN), rowSums(adj.TRN), 
                                                        colSums(adj.PPI), nboot, nthreads, dir.org, seed, 0, dir.org))
}

##Read MCGC attack curves for Organisms with HINT PPI
MCGC.organism$HINT <- list()
for( i in c(4, 5, 6, 8, 9) ){
  ##Print name of the organism.
  message(paste("Organism = ", Organisms[i], sep = ""))
  dir.org <- paste(getwd(), "/", Organisms_file[i], "_HINT", sep = "")
  dir.create(dir.org, showWarnings = F)
  adj.TRN <- NetworksMultiplex$HINT[[i]]$TRN
  adj.PPI <- NetworksMultiplex$HINT[[i]]$PPI
  MCGC.organism$HINT[[i]] <- matrix(0, nboot, ncol(adj.TRN))
  for( b in 1:nboot ){
    MCGC.organism$HINT[[i]][b, ] <- c(unlist(read.table(paste(dir.org, "/boot_", b, sep = ""), sep = "\t", header = F)))
  }
}
save(MCGC.organism, file = "MCGC_Organism_attack_curves.RData")


#####################MCGC attack curves for Shuffled multiplexes for zero redundancy and zero degree-degree coupling##########################

##Number of CPU cores to use
nthreads <- 10

##Number of times to compute redundancy coupling for the shuffled null model.
nboot <- 1000

##Random seed
seed <- 123

##Compute TRN gene ids for shuffled multiplexes with zero degree-degree and redundancy couplings for organisms with BioGRID PPI
for( i in c(4, 6, 8, 9) ){
  
  ##Print name of the organism.
  message(paste("Organism = ", Organisms[i], sep = ""))
  
  ##Get organism TRN and PPI networks from multiplex network object.
  adj.TRN <- NetworksMultiplex$BioGrid[[i]]$TRN
  adj.PPI <- NetworksMultiplex$BioGrid[[i]]$PPI
  
  
  dir.org <- paste(getwd(), "/", Organisms_file[i], "_BioGrid_zero_red_zero_degdeg", sep = "")
  dir.create(dir.org, showWarnings = F)
  system.time(test <- SAShuffleMultiplexParallel(adj.TRN, adj.PPI, colSums(adj.TRN), rowSums(adj.TRN), 
                                                    colSums(adj.PPI), 0, 5, 1e8, 1e-9, 0.001, 0.001, 0, 1e7, 0, 
                                                    0, seed, 1000, 10, dir.org))
}


##Calculate MCGC attack curves for shuffled multiplexes with zero redundancy, zero degree-degree couplings for 
##organisms with BioGRID PPI
for( i in c(4, 6, 8, 9) ){
  
  ##Print name of the organism.
  message(paste("Organism = ", Organisms[i], sep = ""))
  
  ##Get organism TRN and PPI networks from multiplex network object.
  adj.TRN <- NetworksMultiplex$BioGrid[[i]]$TRN
  adj.PPI <- NetworksMultiplex$BioGrid[[i]]$PPI
  dir.read.id <- paste(getwd(), "/", Organisms_file[i], "_BioGrid_zero_red_zero_degdeg", sep = "")
  id.shuffle <- matrix(0, nboot, ncol(adj.TRN))
  for( b in 1:nboot ){
    tmp <- read.table(paste(dir.read.id, "/boot_", b, sep = ""), sep = "\t", header = F)
    id.shuffle[b, ] <- c(unlist(tmp))
  }
  # dir.create(dir.org, showWarnings = F)
  id1 <- 1:ncol(adj.TRN)
  id2 <- 1:ncol(adj.TRN)
  adj1 <- adj.TRN[id1, id1]
  adj2 <- adj.PPI[id2, id2]
  elTRN <- get.edgelist(graph_from_adjacency_matrix(adj1, "undirected")) - 1
  elPPI <- get.edgelist(graph_from_adjacency_matrix(adj2, "undirected")) - 1
  
  ##Make organism specific folder to store the attack curves
  dir.org <- paste(getwd(), "/", Organisms_file[i], "_BioGrid_zero_red_zero_degdeg_attack_curve", sep = "")
  dir.create(dir.org, showWarnings = F)
  
  system.time(test <- MCGCAttackCurveParallel(elTRN, elPPI, ncol(adj.TRN), rowSums(adj.TRN), 
                                                        colSums(adj.PPI), nboot, nthreads, dir.org, seed, 1, dir.read.id))
}

##Read MCGC attack curves for shuffled multiplexes with BioGRID PPI
MCGC.NULL <- list()
MCGC.NULL$BioGRID <- list()
for( i in c(1:9)[-c(1, 3, 7)] ){
  ##Print name of the organism.
  message(paste("Organism = ", Organisms[i], sep = ""))
  dir.org <- paste(getwd(), "/", Organisms_file[i], "_BioGrid_zero_red_zero_degdeg_attack_curve", sep = "")
  dir.create(dir.org, showWarnings = F)
  adj.TRN <- NetworksMultiplex$BioGrid[[i]]$TRN
  adj.PPI <- NetworksMultiplex$BioGrid[[i]]$PPI
  MCGC.NULL$BioGRID[[i]] <- matrix(0, nboot, ncol(adj.TRN))
  for( b in 1:nboot ){
    MCGC.NULL$BioGRID[[i]][b, ] <- c(unlist(read.table(paste(dir.org, "/boot_", b, sep = ""), sep = "\t", header = F)))
  }
}
save(MCGC.NULL, file = "MCGC_NULL_attack_curves.RData")


##Compute TRN gene ids for shuffled multiplexes with zero degree-degree and redundancy couplings for organisms with HINT PPI
for( i in c(4, 5, 6, 8, 9) ){
  
  ##Print name of the organism.
  message(paste("Organism = ", Organisms[i], sep = ""))
  
  ##Get organism TRN and PPI networks from multiplex network object.
  adj.TRN <- NetworksMultiplex$HINT[[i]]$TRN
  adj.PPI <- NetworksMultiplex$HINT[[i]]$PPI
  dir.org <- paste(getwd(), "/", Organisms_file[i], "_HINT_zero_red_zero_degdeg", sep = "")
  dir.create(dir.org, showWarnings = F)
  system.time(id.rand <- SAShuffleMultiplexParallel(adj.TRN, adj.PPI, colSums(adj.TRN), rowSums(adj.TRN), 
                                                    colSums(adj.PPI), 0, 5, 1e8, 1e-9, 0.001, 0.001, 0, 1e7, 0, 
                                                    0, seed, 1000, 10, dir.org))
}

##Calculate MCGC attack curves for shuffled multiplexes with zero redundancy, zero degree-degree couplings for 
##organisms with HINT PPI
for( i in c(4, 5, 6, 8, 9) ){
  
  ##Print name of the organism.
  message(paste("Organism = ", Organisms[i], sep = ""))
  
  ##Get organism TRN and PPI networks from multiplex network object.
  adj.TRN <- NetworksMultiplex$HINT[[i]]$TRN
  adj.PPI <- NetworksMultiplex$HINT[[i]]$PPI
  dir.read.id <- paste(getwd(), "/", Organisms_file[i], "_HINT_zero_red_zero_degdeg", sep = "")
  id1 <- 1:ncol(adj.TRN)
  id2 <- 1:ncol(adj.TRN)
  adj1 <- adj.TRN[id1, id1]
  adj2 <- adj.PPI[id2, id2]
  elTRN <- get.edgelist(graph_from_adjacency_matrix(adj1, "undirected")) - 1
  elPPI <- get.edgelist(graph_from_adjacency_matrix(adj2, "undirected")) - 1
  
  ##Make organism specific folder to store the attack curves
  dir.org <- paste(getwd(), "/", Organisms_file[i], "_HINT_zero_red_zero_degdeg_attack_curve", sep = "")
  dir.create(dir.org, showWarnings = F)
  
  system.time(tmp.curve.rand <- MCGCAttackCurveParallel(elTRN, elPPI, ncol(adj.TRN), rowSums(adj.TRN), 
                                                        colSums(adj.PPI), nboot, nthreads, dir.org, seed, 1, dir.read.id))
}

##Read MCGC attack curves for shuffled multiplexes with HINT PPI
MCGC.NULL$HINT <- list()
for( i in c(1:9)[-c(1, 2, 7)] ){
  ##Print name of the organism.
  message(paste("Organism = ", Organisms[i], sep = ""))
  dir.org <- paste(getwd(), "/", Organisms_file[i], "_HINT_zero_red_zero_degdeg_attack_curve", sep = "")
  dir.create(dir.org, showWarnings = F)
  adj.TRN <- NetworksMultiplex$HINT[[i]]$TRN
  adj.PPI <- NetworksMultiplex$HINT[[i]]$PPI
  MCGC.NULL$HINT[[i]] <- matrix(0, nboot, ncol(adj.TRN))
  for( b in 1:nboot ){
    MCGC.NULL$HINT[[i]][b, ] <- c(unlist(read.table(paste(dir.org, "/boot_", b, sep = ""), sep = "\t", header = F)))
  }
}
save(MCGC.NULL, file = "MCGC_NULL_attack_curves.RData")


##Plots for BioGRID Figs. 4A-4D
len.genes.Bio <- MCC.size.Bio <- rep(0, length(Organisms))
Plot.list.1 <- list()
database.type <- rep("BioGRID", length(Organisms))
for( i in c(4, 6, 8, 9) ){
  cat("Organism = ", Organisms[i], "\n")
  adj.TRN <- NetworksMultiplex$BioGrid[[i]]$TRN
  adj.PPI <- NetworksMultiplex$BioGrid[[i]]$PPI
  id.inter <- NetworksMultiplex$BioGrid[[i]]$genes
  MCC.size.Bio[i] <- MCC.complete.graph(adj.TRN, adj.PPI)$size
  len.genes.Bio[i] <- ncol(adj.TRN)

  id.sample <- 1:nboot
  size.SN <- matrix(0, nboot, 2)
  for( b in id.sample ){
    if( i != 1 ){
      size.SN[b, 1] <- which(MCGC.organism$BioGRID[[i]][b, ] <= sqrt(ncol(adj.TRN)))[1]
      size.SN[b, 2] <- which(MCGC.NULL$BioGRID[[i]][b, ] <= sqrt(ncol(adj.TRN)))[1]
    }else{
      size.SN[b, 1] <- which(MCGC.organism$BioGRID[[i]][b, ] <= 1)[1]
      size.SN[b, 2] <- which(MCGC.NULL$BioGRID[[i]][b, ] <= 1)[1]
    }
  }
  size.plot <- 1:max(c(size.SN))
  b <- id.sample[1]
  
  dat.plot <- data.frame(x = rep((1:length(size.plot))/len.genes.Bio[i], 2), 
                         y = c(MCGC.organism$BioGRID[[i]][b, 1:length(size.plot)]/len.genes.Bio[i], 
                               MCGC.NULL$BioGRID[[i]][b, 1:length(size.plot)]/len.genes.Bio[i]), 
                         Type = c(rep("Original", length(size.plot)), 
                                  rep("Null", length(size.plot))))
  # database.type[i] <- "BioGRID"
  
  dat.plot$Type <- factor(dat.plot$Type, levels = c("Original", "Null"))
  p <- ggline(dat.plot, x = "x", y = "y", 
              numeric.x.axis = T, 
              linetype = "dashed",
              plot_type = c("l"),
              fill = "Type",           # change fill color by mpg_level
              color = "Type",# jco journal color palett. see ?ggpar         # Sort the value in descending order
              palette = c("red", "gray", "red", "black"),
              sort.by.groups = FALSE,     # Don't sort inside each group
              x.text.angle = 50,          # Rotate vertically x axis texts
              ylab = "Relative size of MCGC",
              xlab = "Fraction of nodes attacked",
              # legend = "bottom",
              legend.title = "",
              y.text.angle = 50,
              rotate = F,
              font.x = 14, 
              font.y = 14,
              font.tickslab = 15,
              size = 0.9,
              point.size = 0.1,
              # font.label = 14,
              theme = theme_pubr()
  ) 
  # + 
  #   geom_ribbon(aes(ymin=y.lower, ymax=y.upper, color = Type), linetype=2, alpha=0.1)
  # p <- p + geom_hline(yintercept = 0, linetype = 1) 
  # +   geom_hline(yintercept = -0.2, linetype = 3)
  # p <- p + geom_errorbar(aes(x = x, ymin = y.lower, ymax = y.upper, color = Type))
  
  for( b in id.sample[-1] ){
    dat.plot <- data.frame(x = rep((1:length(size.plot))/len.genes.Bio[i], 2), 
                           y = c(MCGC.organism$BioGRID[[i]][b, 1:length(size.plot)]/len.genes.Bio[i], 
                                 MCGC.NULL$BioGRID[[i]][b, 1:length(size.plot)]/len.genes.Bio[i]), 
                           Type = c(rep("Original", length(size.plot)), 
                                    rep("Null", length(size.plot))))
    dat.plot$Type <- factor(dat.plot$Type, levels = c("Original", "Null"))
    p <- p + geom_line(data = dat.plot, mapping = aes(x = x, y = y, color = Type), 
                       linetype = 2, size = 0.9)
  }
  p <- p + 
    theme(plot.title = element_text(size = 20, color = "black", face="bold.italic"), 
          axis.title.x = element_text(size = 16, colour = "black", face = "bold"), 
          axis.title.y = element_text(size = 16, colour = "black", face = "bold"), 
          legend.text = element_text(size = 16, color = "black", face = "bold")
          , legend.position = "none")
  p <- p +  guides(fill=guide_legend(ncol=2,byrow=F)) + ggtitle(paste(Organisms[i], ", ", database.type[i], 
                                                                      " (", len.genes.Bio[i], ")", sep = ""))
  Plot.list.1[[i]] <- p
  
}

####HINT#####
len.hint <- matrix(0, length(Organisms), 2)
len.genes.HINT <- MCC.size.HINT <- rep(0, length(Organisms))
Plot.list.2 <- list()
database.type <- rep("HINT", 9)
for( i in (1:length(Organisms))[-c(1, 2, 7)] ){
  cat("Organism = ", Organisms[i], "\n")
  adj.TRN <- NetworksMultiplex$HINT[[i]]$TRN
  adj.PPI <- NetworksMultiplex$HINT[[i]]$PPI
  id.inter <- NetworksMultiplex$HINT[[i]]$genes
  MCC.size.HINT[i] <- MCC.complete.graph(adj.TRN, adj.PPI)$size
  len.genes.HINT[i] <- ncol(adj.TRN)
  id.sample <- 1:nboot
  size.SN <- matrix(0, nboot, 2)
  for( b in id.sample ){
    if( i != 1 ){
      size.SN[b, 1] <- which(MCGC.organism$HINT[[i]][b, ] <= sqrt(ncol(adj.TRN)))[1]
      size.SN[b, 2] <- which(MCGC.NULL$HINT[[i]][b, ] <= sqrt(ncol(adj.TRN)))[1]
    }else{
      size.SN[b, 1] <- which(MCGC.organism$HINT[[i]][b, ] <= 1)[1]
      size.SN[b, 2] <- which(MCGC.NULL$HINT[[i]][b, ] <= 1)[1]
    }
  }
  size.plot <- 1:max(c(size.SN))
  b <- id.sample[1]
  
  dat.plot <- data.frame(x = rep((1:length(size.plot))/len.genes.HINT[i], 2), 
                         y = c(MCGC.organism$HINT[[i]][b, 1:length(size.plot)]/len.genes.HINT[i], 
                               MCGC.NULL$HINT[[i]][b, 1:length(size.plot)]/len.genes.HINT[i]), 
                         Type = c(rep("Original", length(size.plot)), 
                                  rep("Null", length(size.plot))))
  
  dat.plot$Type <- factor(dat.plot$Type, levels = c("Original", "Null"))
  p <- ggline(dat.plot, x = "x", y = "y", 
              numeric.x.axis = T, 
              linetype = "dashed",
              plot_type = c("l"),
              fill = "Type",           # change fill color by mpg_level
              color = "Type",# jco journal color palett. see ?ggpar         # Sort the value in descending order
              palette = c("red", "gray", "red", "black"),
              sort.by.groups = FALSE,     # Don't sort inside each group
              x.text.angle = 50,          # Rotate vertically x axis texts
              ylab = "Relative size of MCGC",
              xlab = "Fraction of nodes attacked",
              # legend = "bottom",
              legend.title = "",
              y.text.angle = 50,
              rotate = F,
              font.x = 14, 
              font.y = 14,
              font.tickslab = 15,
              size = 0.9,
              point.size = 0.1,
              # font.label = 12,
              theme = theme_pubr()
  ) 
  # + 
  #   geom_ribbon(aes(ymin=y.lower, ymax=y.upper, color = Type), linetype=2, alpha=0.1)
  # p <- p + geom_hline(yintercept = 0, linetype = 1) 
  # +   geom_hline(yintercept = -0.2, linetype = 3)
  # p <- p + geom_errorbar(aes(x = x, ymin = y.lower, ymax = y.upper, color = Type))
  p <- p + 
    theme(plot.title = element_text(size = 20, color = "black", face="bold.italic"), 
          axis.title.x = element_text(size = 16, colour = "black", face = "bold"), 
          axis.title.y = element_text(size = 16, colour = "black", face = "bold"), 
          legend.text = element_text(size = 16, color = "black", face = "bold")
          , legend.position = "none")
  for( b in id.sample[-1] ){
    dat.plot <- data.frame(x = rep((1:length(size.plot))/len.genes.HINT[i], 2), 
                           y = c(MCGC.organism$HINT[[i]][b, 1:length(size.plot)]/len.genes.HINT[i], 
                                 MCGC.NULL$HINT[[i]][b, 1:length(size.plot)]/len.genes.HINT[i]), 
                           Type = c(rep("Original", length(size.plot)), 
                                    rep("Null", length(size.plot))))
    dat.plot$Type <- factor(dat.plot$Type, levels = c("Original", "Null"))
    p <- p + geom_line(data = dat.plot, mapping = aes(x = x, y = y, color = Type), 
                       linetype = 2, size = 0.9)
  }
  p <- p +  guides(fill=guide_legend(ncol=2,byrow=F)) + ggtitle(paste(Organisms[i], ", ", database.type[i], 
                                                                      " (", len.genes.HINT[i], ")", sep = ""))
  Plot.list.2[[i]] <- p
  
}
dat.plot <- data.frame(x = rep((1:length(size.plot))/len.genes.HINT[i], 2), 
                       y = c(MCGC.organism$HINT[[i]][b, 1:length(size.plot)]/len.genes.HINT[i], 
                             MCGC.NULL$HINT[[i]][b, 1:length(size.plot)]/len.genes.HINT[i]), 
                       Type = c(rep("Original", length(size.plot)), 
                                rep("Null", length(size.plot))))

dat.plot$Type <- factor(dat.plot$Type, levels = c("Original", "Null"))
p <- ggline(dat.plot, x = "x", y = "y", 
            numeric.x.axis = T, 
            linetype = "dashed",
            plot_type = c("l"),
            fill = "Type",           # change fill color by mpg_level
            color = "Type",# jco journal color palett. see ?ggpar         # Sort the value in descending order
            palette = c("red", "gray", "red", "black"),
            sort.by.groups = FALSE,     # Don't sort inside each group
            x.text.angle = 50,          # Rotate vertically x axis texts
            ylab = "Relative size of MCGC",
            xlab = "Fraction of nodes attacked",
            legend = "bottom",
            legend.title = "",
            y.text.angle = 50,
            rotate = F,
            font.x = 16, 
            font.y = 16,
            font.tickslab = 16,
            size = 0.9,
            point.size = 0.1,
            font.label = 16,
            theme = theme_pubr()
)
p <- p + 
  theme(title = element_text(size = 16, color = "black"), 
        axis.title = element_text(size = 16, colour = "black"), 
        legend.text = element_text(size = 25, color = "black"), 
        legend.key.width = unit(5,"cm")
        , legend.position = "bottom") + guides(linetype = guide_legend(override.aes = list(size = 12)))
lgnd <- get_legend(p)

database.type <- rep(0, length(Organisms))
Plot.list <- list()
count <- 1
tmp <- MCC.size.Bio/len.genes.Bio
tmp[is.nan(tmp)] <- 0
for( j in 1:length(Organisms) ){
  if( tmp[j] >= 0.4 ){
    Plot.list[[count]] <- Plot.list.1[[j]]
    count <- count + 1
  }
}

tmp <- MCC.size.HINT/len.genes.HINT
tmp[is.nan(tmp)] <- 0
for( j in 1:length(Organisms) ){
  if( tmp[j] >= 0.4 ){
    Plot.list[[count]] <- Plot.list.2[[j]]
    count <- count + 1
  }
}
plot.1 <- plot_grid(Plot.list[[1]], Plot.list[[2]], Plot.list[[3]], 
                    Plot.list[[4]], Plot.list[[5]], Plot.list[[6]], 
                    Plot.list[[7]], Plot.list[[8]], Plot.list[[9]], ncol = 3, labels = "AUTO", label_size = 30)
plot.2 <- plot_grid(plot.1, lgnd, nrow = 2, rel_heights = c(1, 0.1))
save_plot("Figure_4_paper.png", plot.2, base_height = 14, base_width = 18)
```
