## Supplementary material for "Internetwork connectivity of molecular networks across species of life": Code: Figure_4_Paper.R

```
#Figure 4
##set seed for reproducibility
set.seed(1)

##Load packages and functions
library(Rcpp)
library(Matrix)
library(igraph)
library(ggplot2)
library(ggpubr)
library(cowplot)
Rcpp::sourceCpp('utils.cpp')
Rcpp::sourceCpp("Attack_functions_new_cpp.cpp")
Rcpp::sourceCpp('Simulated_Annealing_Functions.cpp')

####################################Panel 4A####################################################
Plot.fig.4 <- list()
###BioGRID
load("MCGC_Organism_attack_curves.RData")
load("MCGC_NULL_attack_curves.RData")
len.genes.Bio <- MCC.size.Bio <- rep(0, length(Organisms))
Plot.list.1 <- list()
database.type <- rep("BioGRID", 9)

nboot <- 1000
MCC.redundancy <- array(0, c(length(Organisms), 1, nboot))
MCC.corr <- array(0, c(length(Organisms), 1, nboot, 3))
MCC.areas.slopes <- MCC.siz <- array(0, c(2, length(Organisms), 1, nboot))

dat.plot.red.Bio <- dat.plot.red.Bio.size <- dat.plot.corr.Bio <- dat.plot.corr.Bio.size <- list()
for( i in (1:length(Organisms))[c(4, 6, 8, 9)] ){
  message(paste("Organism = ", Organisms[i], sep = ""))
  adj.TRN <- NetworksMultiplex$BioGrid[[i]]$TRN
  adj.PPI <- NetworksMultiplex$BioGrid[[i]]$PPI
  MCC.size.Bio[i] <- MCC.complete.graph(adj.TRN, adj.PPI)$size

  len.genes.Bio[i] <- ncol(adj.TRN)
  
  len <- ncol(adj.TRN)
  for( b in 1:(nboot) ){
    b.id <- b
    j <- 1
    id1 <- 1:ncol(adj.TRN)
    id2 <- 1:ncol(adj.TRN)
    MCC.redundancy[i, j, b.id] <- nnzero(adj.TRN[id1, id1]*adj.PPI[id2, id2])
    MCC.corr[i, j, b.id, ] <- SpearmanRho(rowSums(adj.TRN[id1, id1]), colSums(adj.PPI[id2, id2]), conf.level = 0.95)

    id.stop <- which(MCGC.organism$BioGRID[[i]][b, ] <= sqrt(ncol(adj.TRN)))[1]
    
    if( id.stop >= 2 ){
      MCC.areas.slopes[1, i, j, b.id] <- abs(sintegral((1:(id.stop[length(id.stop)]))/len,
                                                       MCGC.organism$BioGRID[[i]][b, 1:(id.stop[length(id.stop)])]/len)$value)
    }else{
      MCC.areas.slopes[1, i, j, b.id] <- 0
    }
    MCC.siz[1, i, j, b.id] <- id.stop
  }
  
  MCC.rand <- red.rand <- MCC.size.rand <- rep(0, nboot)
  for( b in 1:nboot ){
    id.stop <- which(MCGC.NULL$BioGRID[[i]][b, ] <= sqrt(ncol(adj.TRN)))[1]
    
    if( id.stop >= 2 ){
      MCC.rand[b] <- abs(sintegral((1:(id.stop[length(id.stop)]))/len, 
                                   MCGC.NULL$BioGRID[[i]][b, 1:(id.stop[length(id.stop)])]/len)$value)
    }else{
      MCC.rand[b] <- 0
    }
    MCC.size.rand[b] <- id.stop
    id <- sample(1:ncol(adj.TRN))
    red.rand[b] <- nnzero(adj.TRN[id, id]*adj.PPI)
  }
  
  tmp.MCC <- tmp.red <- tmp.MCC.size <- tmp.corr <- matrix(0, 9, 3)
  for( j in 1 ){
    tmp <- orddom(MCC.rand, MCC.areas.slopes[1, i, j, ])
    tmp.MCC[j, ] <- c(as.numeric(tmp[23, 2]), as.numeric(tmp[24, 2]), 
                      as.numeric(tmp[25, 2]))
    tmp <- (MCC.redundancy[i, 1, 1] - mean(red.rand))/sd(red.rand)
    tmp.red[j, ] <- rep(tmp, 3)
    tmp <- orddom(MCC.size.rand, MCC.siz[1, i, j, ])
    tmp.MCC.size[j, ] <- c(as.numeric(tmp[23, 2]), as.numeric(tmp[24, 2]), 
                           as.numeric(tmp[25, 2]))
  }
  
  tmp.corr[1, ] <- MCC.corr[i, 1, 1, ]
  
  
  dat.plot.red.Bio[[i]] <- data.frame(x = c(tmp.red[1, 1]),
                                      x.lower = tmp.red[1, 2],
                                      x.upper = tmp.red[1, 3],
                                      y = rep(tmp.MCC[1, 1], 1),
                                      y.lower = tmp.MCC[1, 2], 
                                      y.upper = tmp.MCC[1, 3],
                                      Type = unlist(lapply(c(rep(Organisms[i], 1)), function(x){
                                        paste(x, ",BioGRID", sep = "")
                                      })))
  dat.plot.red.Bio.size[[i]] <- data.frame(x = c(tmp.red[1, 1]),
                                           x.lower = tmp.red[1, 2],
                                           x.upper = tmp.red[1, 3],
                                           y = rep(tmp.MCC.size[1, 1], 1),
                                           y.lower = tmp.MCC.size[1, 2], 
                                           y.upper = tmp.MCC.size[1, 3],
                                           Type = unlist(lapply(c(rep(Organisms[i], 1)), function(x){
                                             paste(x, ",BioGRID", sep = "")
                                           })))
  
  dat.plot.corr.Bio[[i]] <- data.frame(x = c(tmp.corr[1, 1]),
                                       x.lower = tmp.corr[1, 2],
                                       x.upper = tmp.corr[1, 3],
                                       y = rep(tmp.MCC[1, 1], 1),
                                       y.lower = tmp.MCC[1, 2], 
                                       y.upper = tmp.MCC[1, 3],
                                       Type = unlist(lapply(c(rep(Organisms[i])), function(x){
                                         paste(x, ", BioGRID", sep = "")
                                       })))
  
  dat.plot.corr.Bio.size[[i]] <- data.frame(x = c(tmp.corr[1, 1]),
                                            x.lower = tmp.corr[1, 2],
                                            x.upper = tmp.corr[1, 3],
                                            y = rep(tmp.MCC.size[1, 1], 1),
                                            y.lower = tmp.MCC.size[1, 2], 
                                            y.upper = tmp.MCC.size[1, 3],
                                            Type = unlist(lapply(c(rep(Organisms[i])), function(x){
                                              paste(x, ", BioGRID", sep = "")
                                            })))
  
}
dat.plot.corr.Bio.org <- dat.plot.corr.Bio
dat.plot.corr.Bio.org.size <- dat.plot.corr.Bio.size
dat.plot.red.Bio.org <- dat.plot.red.Bio
dat.plot.red.Bio.org.size <- dat.plot.red.Bio.size


####HINT#####
MCC.redundancy <- array(0, c(length(Organisms), 1, nboot))
MCC.corr <- array(0, c(length(Organisms), 1, nboot, 3))
MCC.areas.slopes <- MCC.siz <- array(0, c(2, length(Organisms), 1, nboot))

len.genes.HINT <- MCC.size.HINT <- rep(0, length(Organisms))
Plot.list.2 <- list()
dat.plot.red.HINT <- dat.plot.corr.HINT <- dat.plot.red.HINT.size <- dat.plot.corr.HINT.size <- list()
database.type <- rep("HINT", 9)
for( i in (1:length(Organisms))[c(4, 5, 6, 8, 9)] ){
  message(paste("Organism = ", Organisms[i], sep = ""))
  adj.TRN <- NetworksMultiplex$HINT[[i]]$TRN
  adj.PPI <- NetworksMultiplex$HINT[[i]]$PPI
  MCC.size.HINT[i] <- MCC.complete.graph(adj.TRN, adj.PPI)$size
  len <- ncol(adj.TRN)
  for( b in 1:(nboot) ){
    j <- 1
    b.id <- b
    id1 <- 1:ncol(adj.TRN)
    id2 <- 1:ncol(adj.TRN)
    MCC.redundancy[i, j, b.id] <- nnzero(adj.TRN[id1, id1]*adj.PPI[id2, id2])
    MCC.corr[i, j, b.id, ] <- SpearmanRho(rowSums(adj.TRN[id1, id1]), colSums(adj.PPI[id2, id2]), conf.level = 0.95)
    id.stop <- which(MCGC.organism$HINT[[i]][b, ] <= sqrt(ncol(adj.TRN)))[1]
    
    if( id.stop >= 2 ){
      MCC.areas.slopes[1, i, j, b.id] <- abs(sintegral((1:(id.stop[length(id.stop)]))/len,
                                                       MCGC.organism$HINT[[i]][b, 1:(id.stop[length(id.stop)])]/len)$value)
    }else{
      MCC.areas.slopes[1, i, j, b.id] <- 0
    }
    MCC.siz[1, i, j, b.id] <- id.stop
  }
  
  MCC.rand <- MCC.size.rand <- red.rand <- rep(0, nboot)
  for( b in 1:nboot ){
    id.stop <- which(MCGC.NULL$HINT[[i]][b, ] <= sqrt(ncol(adj.TRN)))[1]
    
    if( id.stop >= 2 ){
      MCC.rand[b] <- abs(sintegral((1:(id.stop[length(id.stop)]))/len, 
                                   MCGC.NULL$HINT[[i]][b, 1:(id.stop[length(id.stop)])]/len)$value)
    }else{
      MCC.rand[b] <- 0
    }
    MCC.size.rand[b] <- id.stop
    id <- sample(1:ncol(adj.TRN))
    red.rand[b] <- nnzero(adj.TRN[id, id]*adj.PPI)
  }
  
  tmp.MCC <- tmp.MCC.size <- tmp.red <- tmp.corr <- matrix(0, 9, 3)
  for( j in 1 ){
    tmp <- orddom(MCC.rand, MCC.areas.slopes[1, i, j, ])
    tmp.MCC[j, ] <- c(as.numeric(tmp[23, 2]), as.numeric(tmp[24, 2]), 
                      as.numeric(tmp[25, 2]))
    tmp <- orddom(MCC.size.rand, MCC.siz[1, i, j, ])
    tmp.MCC.size[j, ] <- c(as.numeric(tmp[23, 2]), as.numeric(tmp[24, 2]), 
                           as.numeric(tmp[25, 2]))
    tmp <- (MCC.redundancy[i, 1, 1] - mean(red.rand))/sd(red.rand)
    tmp.red[j, ] <- rep(tmp, 3)
  }
  
  tmp.corr[1, ] <- MCC.corr[i, 1, 1, ]
  
  len.genes.HINT[i] <- ncol(adj.TRN)
  id.sample <- 1:nboot
  dat.plot.red.HINT[[i]] <- data.frame(x = c(tmp.red[1, 1]),
                                       x.lower = tmp.red[1, 2],
                                       x.upper = tmp.red[1, 3],
                                       y = rep(tmp.MCC[1, 1], 1),
                                       y.lower = tmp.MCC[1, 2], 
                                       y.upper = tmp.MCC[1, 3],
                                       Type = unlist(lapply(c(rep(Organisms[i], 1)), function(x){
                                         paste(x, ", HINT", sep = "")
                                       })))
  
  dat.plot.corr.HINT[[i]] <- data.frame(x = c(tmp.corr[1, 1]),
                                        x.lower = tmp.corr[1, 2],
                                        x.upper = tmp.corr[1, 3],
                                        y = rep(tmp.MCC[1, 1], 1),
                                        y.lower = tmp.MCC[1, 2], 
                                        y.upper = tmp.MCC[1, 3],
                                        Type = unlist(lapply(c(rep(Organisms[i], 1)), function(x){
                                          paste(x, ", HINT", sep = "")
                                        })))
  
  dat.plot.red.HINT.size[[i]] <- data.frame(x = c(tmp.red[1, 1]),
                                            x.lower = tmp.red[1, 2],
                                            x.upper = tmp.red[1, 3],
                                            y = rep(tmp.MCC.size[1, 1], 1),
                                            y.lower = tmp.MCC.size[1, 2], 
                                            y.upper = tmp.MCC.size[1, 3],
                                            Type = unlist(lapply(c(rep(Organisms[i], 1)), function(x){
                                              paste(x, ", HINT", sep = "")
                                            })))
  
  dat.plot.corr.HINT.size[[i]] <- data.frame(x = c(tmp.corr[1, 1]),
                                             x.lower = tmp.corr[1, 2],
                                             x.upper = tmp.corr[1, 3],
                                             y = rep(tmp.MCC.size[1, 1], 1),
                                             y.lower = tmp.MCC.size[1, 2], 
                                             y.upper = tmp.MCC.size[1, 3],
                                             Type = unlist(lapply(c(rep(Organisms[i], 1)), function(x){
                                               paste(x, ", HINT", sep = "")
                                             })))
  
}


count <- 1
tmp <- MCC.size.Bio/len.genes.Bio
tmp[is.nan(tmp)] <- 0
for( j in c(4, 6, 8, 9) ){
  if( tmp[j] >= 0.4 ){
    
    if( count == 1 ){
      data.plot <- dat.plot.corr.Bio[[j]]
    }else{
      data.plot <- rbind(data.plot, dat.plot.corr.Bio[[j]])
    }
    
    count <- count + 1
  }
  
}

tmp <- MCC.size.HINT/len.genes.HINT
tmp[is.nan(tmp)] <- 0
for( j in c(4, 5, 6, 8, 9) ){
  if( tmp[j] >= 0.3 ){
    data.plot <- rbind(data.plot, dat.plot.corr.HINT[[j]])
    count <- count + 1
  }
}
Org.order <- as.character(data.plot$Type[order(data.plot$y, decreasing = F)])
data.cor <- paste(" = ", 
                  signif(cor(data.plot$x, data.plot$y, method = "pearson"), 4), 
                  ", p = ", signif(cor.test(data.plot$x, data.plot$y, 
                                            method = "pearson")$p.value, 4), sep = "")
data.plot$Type <- factor(data.plot$Type, levels = Org.order)
p <- ggbarplot(data.plot, x = "Type", y = "x", 
               fill = "blue",       
               color = "blue",
               # palette = c("red", "blue"),
               sort.by.groups = FALSE,    
               # x.text.angle = 50,         
               xlab = "",
               ylab = expression("C"["D"]),
               # legend = "bottom",
               legend.title = "",
               y.text.angle = 50,
               rotate = T,
               font.x = 16, 
               font.y = 16,
               font.tickslab = 14,
               size = 0.9,
               point.size = 0.1,
               font.label = 12,
               stat = "identity",
               theme = theme_pubr()
) + 
  ggtitle(bquote("r" ~ .(data.cor)))+ geom_hline(yintercept = 0) 
p <- p + geom_errorbar(aes(ymin = x.lower, ymax = x.upper), width = 0.2) +
  theme(title = element_text(size = 18, color = "black"), 
        plot.title = element_text(size = 18, color = "black"), 
        axis.title = element_text(size = 16, colour = "black"), 
        legend.text = element_text(size = 16, color = "black"))
p1 <- p


data.cor <- paste(" = ", 
                  signif(cor(data.plot$x, data.plot$y, method = "spearman"), 4), 
                  ", p = ", signif(cor.test(data.plot$x, data.plot$y, 
                                            method = "spearman")$p.value, 4), sep = "")
data.plot$Type <- factor(data.plot$Type, levels = Org.order)
p <- ggbarplot(data.plot, x = "Type", y = "y", 
               fill = "red",           # change fill color by mpg_level
               color = "red",# jco journal color palett. see ?ggpar         # Sort the value in descending order
               # palette = c("red", "blue"),
               sort.by.groups = FALSE,     # Don't sort inside each group
               # x.text.angle = 50,          # Rotate vertically x axis texts
               xlab = "",
               ylab = expression("R"),
               # legend = "bottom",
               legend.title = "",
               y.text.angle = 50,
               rotate = T,
               font.x = 16, 
               font.y = 16,
               font.tickslab = 14,
               size = 0.9,
               point.size = 0.1,
               font.label = 12,
               stat = "identity",
               theme = theme_pubr()
)
p2 <- p + geom_hline(yintercept = 0) + geom_errorbar(aes(ymin = y.lower, ymax = y.upper), width = 0.2)

count <- 1
tmp <- MCC.size.Bio/len.genes.Bio
tmp[is.nan(tmp)] <- 0
for( j in (1:length(Organisms))[-3] ){
  if( tmp[j] >= 0.4 ){
    
    if( count == 1 ){
      data.plot <- dat.plot.red.Bio[[j]]
    }else{
      data.plot <- rbind(data.plot, dat.plot.red.Bio[[j]])
    }
    
    count <- count + 1
  }
  
}

tmp <- MCC.size.HINT/len.genes.HINT
tmp[is.nan(tmp)] <- 0
for( j in (1:length(Organisms))[-c(1, 2, 7)] ){
  if( tmp[j] >= 0.4 ){
    data.plot <- rbind(data.plot, dat.plot.red.HINT[[j]])
    count <- count + 1
  }
}

Org.order <- as.character(data.plot$Type[order(data.plot$y, decreasing = F)])
# Org.order <- data.plot$Type[order(data.plot$y, decreasing = F)]
# dat <- data.frame(x = c(data.plot$Type, data.plot$Type), 
#                   y = c(data.plot$x, data.plot$y), 
#                   y.lower = c(data.plot$x.lower, data.plot$y.lower), 
#                   y.upper = c(data.plot$x.upper, data.plot$y.upper), 
#                   Type = c(rep("CD", nrow(data.plot)), rep("Rb", nrow(data.plot))))
data.cor <- paste(" = ", 
                  signif(cor(data.plot$x, data.plot$y, method = "spearman"), 4), 
                  ", p = ", signif(cor.test(data.plot$x, data.plot$y, 
                                            method = "spearman")$p.value, 4), sep = "")
data.plot$Type <- factor(data.plot$Type, levels = Org.order)
p <- ggbarplot(data.plot, x = "Type", y = "x", 
               fill = "blue",           # change fill color by mpg_level
               color = "blue",# jco journal color palett. see ?ggpar         # Sort the value in descending order
               # palette = c("red", "blue"),
               sort.by.groups = FALSE,     # Don't sort inside each group
               # x.text.angle = 50,          # Rotate vertically x axis texts
               xlab = "",
               ylab = expression("C"["R"]),
               # legend = "bottom",
               legend.title = "",
               y.text.angle = 50,
               rotate = T,
               font.x = 16, 
               font.y = 16,
               font.tickslab = 14,
               size = 0.9,
               point.size = 0.1,
               font.label = 12,
               stat = "identity",
               theme = theme_pubr()
) + 
  ggtitle(bquote("r" ~ .(data.cor)))+ geom_hline(yintercept = 0) 
p <- p + yscale("log10", T) + geom_errorbar(aes(ymin = x.lower, ymax = x.upper), width = 0.2) 
p3 <- p

Plot.fig.4[[1]] <- plot_grid(p1, 
                             p3 + 
                               theme(axis.text.y = element_blank(),
                                     # axis.line.y = element_blank(),
                                     axis.title.y= element_blank(),
                                     axis.ticks.y= element_blank(), 
                                     axis.title.x = element_text(size = 16, colour = "black")), 
                             p2 + 
                               theme(axis.text.y = element_blank(),
                                     # axis.line.y = element_blank(),
                                     axis.title.y= element_blank(),
                                     axis.ticks.y= element_blank(), 
                                     axis.title.x = element_text(size = 16, colour = "black")),
                             nrow = 1,
                             rel_widths = c(1.2, 1, 1),
                             align = 'h', axis = 'tb')


#############################################Figure 4B############################################################3
#sample genes and proteins from organism multiplex with specific degdeg and redundancy coupling
#degdeg and redundancy coupling explored over a grid. And also generate the MCGC attack curves
###redundancy values
edge.vec <- list()
edge.vec[[4]] <- seq(0, 40, by = 10)
edge.vec[[6]] <- seq(0, 80, length = 5)
edge.vec[[8]] <- seq(0, 40, length = 5)
edge.vec[[9]] <- seq(0, 100, length = 5)

###number of sampled genes
NumGenes <- rep(500, 9)
NumGenes[4] <- 1000
NumGenes[6] <- 2000
NumGenes[8] <- 300
NumGenes[9] <- 500

###degree-degree coupling
cor.req <- cor.vec <- seq(0, 0.5, by = 0.1)

###number of repititions for MCGC attack
nboot <- 100
for( i in c(4, 6, 8, 9) ){
  # Start the clock!
  
  for( k in (1:length(edge.vec[[i]])) ){
    for( j in 1:length(cor.vec) ){
      ##Print name of the organism.
      message(paste("Organism = ", Organisms[i], " edge = ", edge.vec[[i]][[k]], " cor = ", cor.vec[j], sep = ""))
      
      ##Get organism TRN and PPI networks from multiplex network object.
      adj.TRN <- NetworksMultiplex$BioGrid[[i]]$TRN
      adj.PPI <- NetworksMultiplex$BioGrid[[i]]$PPI
      rin <- cor(colSums(adj.TRN), colSums(adj.PPI), method = "spearman")
      rout <- cor(rowSums(adj.TRN), colSums(adj.PPI), method = "spearman")
      dir.org <- paste(getwd(), "/", Organisms_file[i], "_BioGrid_sample_degdeg_", cor.vec[j], "_red_", edge.vec[[i]][k], 
                       sep = "")
      dir.create(dir.org, showWarnings = F)
      ptm <- proc.time()
      for( b in 1:nboot ){
        cat("boot = ", b, "\n")
        id <- c(SASampleMultiplex(adj.TRN, adj.PPI, colSums(adj.TRN), rowSums(adj.TRN), 
                                  colSums(adj.PPI), edge.vec[[i]][k], 5, 1e8, 1e-9, 
                                  0.01, 0.001, 10, 1e7, cor.vec[j], 
                                  cor.vec[j]/2, NumGenes[i]))
        file <- paste(dir.org, "/boot_", b, sep = "")
        write.table(cbind(id), file, sep = "\t", row.names = F, 
                    col.names = F, quote = F)
      }
      # Stop the clock
      proc.time() - ptm
      
      
      dir.write <- paste(getwd(), "/", Organisms_file[i], "_BioGrid_sample_degdeg_", cor.vec[j], "_red_", 
                         edge.vec[[i]][k], "_Attack_curves", sep = "")
      dir.create(dir.write, showWarnings = F)
      system.time(id.rand <- MCGCAttackCurveSampleParallel(adj.TRN, adj.PPI, NumGenes[i], nboot, nthreads, 
                                                           dir.write, seed, 1, dir.org, 1:ncol(adj.TRN)))
    }
  }
}

###Plot heatmap for Figure 4B mean
ht_list <- list()
nboot <- 100
for( i in c(4, 6, 8, 9) ){
  len <- NumGenes[i]
  MCGC.mat.mean <- MCGC.mat.lower <- MCGC.mat.upper <- matrix(0, length(cor.vec), length(edge.vec[[i]]))
  MCGC.rand <- rep(0, nboot)
  dir.org <- paste(getwd(), "/", Organisms_file[i], "_BioGrid_sample_degdeg_", cor.vec[1], "_red_", edge.vec[[i]][1], 
                   sep = "")
  dir.write <- paste(getwd(), "/", Organisms_file[i], "_BioGrid_sample_degdeg_", cor.vec[1], "_red_", 
                     edge.vec[[i]][1], "_Attack_curves", sep = "")
  for( b in 1:nboot ){
    MCGC.temp <- c(unlist(read.table(paste(dir.write, "/boot_", b, sep = ""), sep = "\t", header = F)))
    id.stop <- which(MCGC.temp <= 1)[1]
    if( id.stop >= 2 ){
      MCGC.rand[b] <- abs(sintegral((1:id.stop)/len, 
                                    MCGC.temp[1:id.stop]/len)$value)
    }else{
      MCGC.rand[b] <- 0
    }
  }
  for( k in 1:length(edge.vec[[i]]) ){
    for( j in 1:length(cor.vec) ){
      message(paste("Organism = ", Organisms[i], " edge = ", edge.vec[[i]][[k]], " cor = ", cor.vec[j], sep = ""))
      
      
      MCGC.sample <- rep(0, nboot)
      dir.org <- paste(getwd(), "/", Organisms_file[i], "_BioGrid_sample_degdeg_", cor.vec[j], "_red_", edge.vec[[i]][k], 
                       sep = "")
      dir.write <- paste(getwd(), "/", Organisms_file[i], "_BioGrid_sample_degdeg_", cor.vec[j], "_red_", 
                         edge.vec[[i]][k], "_Attack_curves", sep = "")
      for( b in 1:nboot ){
        MCGC.temp <- c(unlist(read.table(paste(dir.write, "/boot_", b, sep = ""), sep = "\t", header = F)))
        id.stop <- which(MCGC.temp <= 1)[1]
        if( id.stop >= 2 ){
          MCGC.sample[b] <- abs(sintegral((1:id.stop)/len, 
                                          MCGC.temp[1:id.stop]/len)$value)
        }else{
          MCGC.sample[b] <- 0
        }
      }
      tmp <- orddom(MCGC.rand, MCGC.sample)
      MCGC.mat.mean[j, k] <- as.numeric(tmp[23, 2])
      MCGC.mat.lower[j, k] <- as.numeric(tmp[24, 2])
      MCGC.mat.upper[j, k] <- as.numeric(tmp[25, 2])
    }
  }
  MCGC.mat.mean[is.nan(MCGC.mat.mean)] <- 0
  MCGC.mat.mean <- MCGC.mat.mean[rev(1:nrow(MCGC.mat.mean)), ]
  
  library(ComplexHeatmap)
  library(circlize)
  
  if( i == 4 ){
    rownames(MCGC.mat.lower) <- cor.vec/2
    rownames(MCGC.mat.mean) <- rev(cor.vec/2)
  }
  colnames(MCGC.mat.mean) <- colnames(MCGC.mat.lower) <- colnames(MCGC.mat.upper) <- edge.vec[[i]]
  col_fun = colorRamp2(c(0, 4, 8), c("white", "blue", "red"))
  
  if( i == 4 ){
    ht1 <- Heatmap(MCGC.mat.mean, cluster_rows = F, cluster_columns = F, col = col_fun, 
                   column_title = expression(atop("C"["R"],
                                                  italic("S. cerevisiae"))), 
                   row_title = expression(paste("C"["D"]^{"out"}, sep = "")), 
                   heatmap_legend_param = list(title = "R, mean - se", title_gp = gpar(fontsize = 30), 
                                               labels_gp = gpar(fontsize = 15)), 
                   use_raster = T, raster_quality = 4,
                   raster_device = "png", column_title_gp = gpar(fontsize = 30), column_title_side = "bottom", 
                   row_title_side = "left", row_names_side = "left", row_title_gp = gpar(fontsize = 30), 
                   row_names_gp = gpar(fontsize = 28), column_names_gp = gpar(fontsize = 28)) 
    ht1@matrix_param$border <- T
    ht1@heatmap_param$show_heatmap_legend <- F
    ht1@matrix_param$border <- T
    ht_list[[1]] <- ht1
  }else{
    if( i == 6 ){
      ht2 <- Heatmap(MCGC.mat.mean, cluster_rows = F, cluster_columns = F, col = col_fun, 
                     column_title = expression(atop("C"["R"],
                                                    italic("D. melanogaster"))), 
                     heatmap_legend_param = list(title = "R", title_gp = gpar(fontsize = 30), 
                                                 labels_gp = gpar(fontsize = 24)), 
                     use_raster = T, raster_quality = 4,
                     raster_device = "png", column_title_gp = gpar(fontsize = 30), column_title_side = "bottom", 
                     column_names_gp = gpar(fontsize = 28)) 
      ht2@heatmap_param$show_heatmap_legend <- F
      ht2@matrix_param$border <- T
      ht2@matrix_legend_param$border <- T
      ht_list[[2]] <- ht2
      
    }
    if( i == 8 ){
      ht3 <- Heatmap(MCGC.mat.mean, cluster_rows = F, cluster_columns = F, col = col_fun, 
                     column_title = expression(atop("C"["R"],
                                                    italic("M. musculus"))), 
                     heatmap_legend_param = list(title = "R, mean + se", title_gp = gpar(fontsize = 30), 
                                                 labels_gp = gpar(fontsize = 28)), 
                     use_raster = T, raster_quality = 4,
                     raster_device = "png", column_title_gp = gpar(fontsize = 30), column_title_side = "bottom", 
                     column_names_gp = gpar(fontsize = 28)) 
      ht3@heatmap_param$show_heatmap_legend <- F
      ht3@matrix_param$border <- T
      ht_list[[3]] <- ht3
    }
    if( i == 9 ){
      ht4 <- Heatmap(MCGC.mat.mean, cluster_rows = F, cluster_columns = F, col = col_fun, 
                     column_title = expression(atop("C"["R"],
                                                    italic("H. sapiens"))), 
                     heatmap_legend_param = list(title = "R", title_gp = gpar(fontsize = 30), 
                                                 labels_gp = gpar(fontsize = 24)), 
                     use_raster = T, raster_quality = 4,
                     raster_device = "png", column_title_gp = gpar(fontsize = 30), column_title_side = "bottom", 
                     column_names_gp = gpar(fontsize = 28)) 
      ht4@heatmap_param$show_heatmap_legend <- T
      ht4@matrix_param$border <- T
      ht4@matrix_legend_param$border <- T
      ht4@matrix_legend_param$legend_height <- unit(10, "cm")
      ht4@matrix_legend_param$grid_width <- unit(7, "mm")
      ht4@matrix_legend_param$gap <- unit(1, "cm")
      ht_list[[4]] <- ht4
    }
  }
}

ht_list1 <- ht1 + ht2 + ht3 + ht4
png(paste("Figure_4B.png", sep = ""), width = 720*4 + 80, height = 720)
draw(ht_list1, row_title = expression(paste("C"["D"]^{"in"}, "=2", "C"["D"]^{"out"}, sep = "")), 
     row_title_gp = gpar(fontsize = 34, fontface = "bold"), ht_gap = unit(1, "cm"))
dev.off()


##Figure 4B Lower CI
ht_list <- list()
nboot <- 100
for( i in c(4, 6, 8, 9) ){
  len <- NumGenes[i]
  MCGC.mat.mean <- MCGC.mat.lower <- MCGC.mat.upper <- matrix(0, length(cor.vec), length(edge.vec[[i]]))
  MCGC.rand <- rep(0, nboot)
  dir.org <- paste(getwd(), "/", Organisms_file[i], "_BioGrid_sample_degdeg_", cor.vec[1], "_red_", edge.vec[[i]][1], 
                   sep = "")
  dir.write <- paste(getwd(), "/", Organisms_file[i], "_BioGrid_sample_degdeg_", cor.vec[1], "_red_", 
                     edge.vec[[i]][1], "_Attack_curves", sep = "")
  for( b in 1:nboot ){
    MCGC.temp <- c(unlist(read.table(paste(dir.write, "/boot_", b, sep = ""), sep = "\t", header = F)))
    id.stop <- which(MCGC.temp <= 1)[1]
    if( id.stop >= 2 ){
      MCGC.rand[b] <- abs(sintegral((1:id.stop)/len, 
                                    MCGC.temp[1:id.stop]/len)$value)
    }else{
      MCGC.rand[b] <- 0
    }
  }
  for( k in 1:length(edge.vec[[i]]) ){
    for( j in 1:length(cor.vec) ){
      message(paste("Organism = ", Organisms[i], " edge = ", edge.vec[[i]][[k]], " cor = ", cor.vec[j], sep = ""))
      
      
      MCGC.sample <- rep(0, nboot)
      dir.org <- paste(getwd(), "/", Organisms_file[i], "_BioGrid_sample_degdeg_", cor.vec[j], "_red_", edge.vec[[i]][k], 
                       sep = "")
      dir.write <- paste(getwd(), "/", Organisms_file[i], "_BioGrid_sample_degdeg_", cor.vec[j], "_red_", 
                         edge.vec[[i]][k], "_Attack_curves", sep = "")
      for( b in 1:nboot ){
        MCGC.temp <- c(unlist(read.table(paste(dir.write, "/boot_", b, sep = ""), sep = "\t", header = F)))
        id.stop <- which(MCGC.temp <= 1)[1]
        if( id.stop >= 2 ){
          MCGC.sample[b] <- abs(sintegral((1:id.stop)/len, 
                                          MCGC.temp[1:id.stop]/len)$value)
        }else{
          MCGC.sample[b] <- 0
        }
      }
      tmp <- orddom(MCGC.rand, MCGC.sample)
      MCGC.mat.mean[j, k] <- as.numeric(tmp[23, 2])
      MCGC.mat.lower[j, k] <- as.numeric(tmp[24, 2])
      MCGC.mat.upper[j, k] <- as.numeric(tmp[25, 2])
    }
  }
  MCGC.mat.lower[is.nan(MCGC.mat.lower)] <- 0
  MCGC.mat.lower <- MCGC.mat.lower[rev(1:nrow(MCGC.mat.lower)), ]
  
  library(ComplexHeatmap)
  library(circlize)
  
  if( i == 4 ){
    rownames(MCGC.mat.lower) <- rev(cor.vec/2)
    # rownames(MCGC.mat.mean) <- rev(cor.vec/2)
  }
  colnames(MCGC.mat.mean) <- colnames(MCGC.mat.lower) <- colnames(MCGC.mat.upper) <- edge.vec[[i]]
  col_fun = colorRamp2(c(0, 3.5, 7.5), c("white", "blue", "red"))
  
  if( i == 4 ){
    ht1 <- Heatmap(MCGC.mat.lower, cluster_rows = F, cluster_columns = F, col = col_fun, 
                   column_title = expression(atop("C"["R"],
                                                  italic("S. cerevisiae"))), 
                   row_title = expression(paste("C"["D"]^{"out"}, sep = "")), 
                   heatmap_legend_param = list(title = "R, mean - se", title_gp = gpar(fontsize = 30), 
                                               labels_gp = gpar(fontsize = 15)), 
                   use_raster = T, raster_quality = 4,
                   raster_device = "png", column_title_gp = gpar(fontsize = 30), column_title_side = "bottom", 
                   row_title_side = "left", row_names_side = "left", row_title_gp = gpar(fontsize = 30), 
                   row_names_gp = gpar(fontsize = 28), column_names_gp = gpar(fontsize = 28)) 
    ht1@matrix_param$border <- T
    ht1@heatmap_param$show_heatmap_legend <- F
    ht1@matrix_param$border <- T
    ht_list[[1]] <- ht1
  }else{
    if( i == 6 ){
      ht2 <- Heatmap(MCGC.mat.lower, cluster_rows = F, cluster_columns = F, col = col_fun, 
                     column_title = expression(atop("C"["R"],
                                                    italic("D. melanogaster"))), 
                     heatmap_legend_param = list(title = "R", title_gp = gpar(fontsize = 30), 
                                                 labels_gp = gpar(fontsize = 24)), 
                     use_raster = T, raster_quality = 4,
                     raster_device = "png", column_title_gp = gpar(fontsize = 30), column_title_side = "bottom", 
                     column_names_gp = gpar(fontsize = 28)) 
      ht2@heatmap_param$show_heatmap_legend <- F
      ht2@matrix_param$border <- T
      ht2@matrix_legend_param$border <- T
      ht_list[[2]] <- ht2
      
    }
    if( i == 8 ){
      ht3 <- Heatmap(MCGC.mat.lower, cluster_rows = F, cluster_columns = F, col = col_fun, 
                     column_title = expression(atop("C"["R"],
                                                    italic("M. musculus"))), 
                     heatmap_legend_param = list(title = "R, mean + se", title_gp = gpar(fontsize = 30), 
                                                 labels_gp = gpar(fontsize = 28)), 
                     use_raster = T, raster_quality = 4,
                     raster_device = "png", column_title_gp = gpar(fontsize = 30), column_title_side = "bottom", 
                     column_names_gp = gpar(fontsize = 28)) 
      ht3@heatmap_param$show_heatmap_legend <- F
      ht3@matrix_param$border <- T
      ht_list[[3]] <- ht3
    }
    if( i == 9 ){
      ht4 <- Heatmap(MCGC.mat.lower, cluster_rows = F, cluster_columns = F, col = col_fun, 
                     column_title = expression(atop("C"["R"],
                                                    italic("H. sapiens"))), 
                     heatmap_legend_param = list(title = "R", title_gp = gpar(fontsize = 30), 
                                                 labels_gp = gpar(fontsize = 24)), 
                     use_raster = T, raster_quality = 4,
                     raster_device = "png", column_title_gp = gpar(fontsize = 30), column_title_side = "bottom", 
                     column_names_gp = gpar(fontsize = 28)) 
      ht4@heatmap_param$show_heatmap_legend <- T
      ht4@matrix_param$border <- T
      ht4@matrix_legend_param$border <- T
      ht4@matrix_legend_param$legend_height <- unit(10, "cm")
      ht4@matrix_legend_param$grid_width <- unit(7, "mm")
      ht4@matrix_legend_param$gap <- unit(1, "cm")
      ht_list[[4]] <- ht4
    }
  }
}

ht_list1 <- ht1 + ht2 + ht3 + ht4
png(paste("Figure_4B_lower_CI.png", sep = ""), width = 720*4 + 80, height = 720)
draw(ht_list1, row_title = expression(paste("C"["D"]^{"in"}, "=2", "C"["D"]^{"out"}, sep = "")), 
     row_title_gp = gpar(fontsize = 34, fontface = "bold"), ht_gap = unit(1, "cm"))
dev.off()


##Figure 4B Upper CI
ht_list <- list()
nboot <- 100
for( i in c(4, 6, 8, 9) ){
  len <- NumGenes[i]
  MCGC.mat.mean <- MCGC.mat.lower <- MCGC.mat.upper <- matrix(0, length(cor.vec), length(edge.vec[[i]]))
  MCGC.rand <- rep(0, nboot)
  dir.org <- paste(getwd(), "/", Organisms_file[i], "_BioGrid_sample_degdeg_", cor.vec[1], "_red_", edge.vec[[i]][1], 
                   sep = "")
  dir.write <- paste(getwd(), "/", Organisms_file[i], "_BioGrid_sample_degdeg_", cor.vec[1], "_red_", 
                     edge.vec[[i]][1], "_Attack_curves", sep = "")
  for( b in 1:nboot ){
    MCGC.temp <- c(unlist(read.table(paste(dir.write, "/boot_", b, sep = ""), sep = "\t", header = F)))
    id.stop <- which(MCGC.temp <= 1)[1]
    if( id.stop >= 2 ){
      MCGC.rand[b] <- abs(sintegral((1:id.stop)/len, 
                                    MCGC.temp[1:id.stop]/len)$value)
    }else{
      MCGC.rand[b] <- 0
    }
  }
  for( k in 1:length(edge.vec[[i]]) ){
    for( j in 1:length(cor.vec) ){
      message(paste("Organism = ", Organisms[i], " edge = ", edge.vec[[i]][[k]], " cor = ", cor.vec[j], sep = ""))
      
      
      MCGC.sample <- rep(0, nboot)
      dir.org <- paste(getwd(), "/", Organisms_file[i], "_BioGrid_sample_degdeg_", cor.vec[j], "_red_", edge.vec[[i]][k], 
                       sep = "")
      dir.write <- paste(getwd(), "/", Organisms_file[i], "_BioGrid_sample_degdeg_", cor.vec[j], "_red_", 
                         edge.vec[[i]][k], "_Attack_curves", sep = "")
      for( b in 1:nboot ){
        MCGC.temp <- c(unlist(read.table(paste(dir.write, "/boot_", b, sep = ""), sep = "\t", header = F)))
        id.stop <- which(MCGC.temp <= 1)[1]
        if( id.stop >= 2 ){
          MCGC.sample[b] <- abs(sintegral((1:id.stop)/len, 
                                          MCGC.temp[1:id.stop]/len)$value)
        }else{
          MCGC.sample[b] <- 0
        }
      }
      tmp <- orddom(MCGC.rand, MCGC.sample)
      MCGC.mat.mean[j, k] <- as.numeric(tmp[23, 2])
      MCGC.mat.lower[j, k] <- as.numeric(tmp[24, 2])
      MCGC.mat.upper[j, k] <- as.numeric(tmp[25, 2])
    }
  }
  MCGC.mat.upper[is.nan(MCGC.mat.upper)] <- 0
  MCGC.mat.upper <- MCGC.mat.upper[rev(1:nrow(MCGC.mat.upper)), ]
  
  library(ComplexHeatmap)
  library(circlize)
  
  if( i == 4 ){
    rownames(MCGC.mat.upper) <- rev(cor.vec/2)
    # rownames(MCGC.mat.mean) <- rev(cor.vec/2)
  }
  colnames(MCGC.mat.mean) <- colnames(MCGC.mat.lower) <- colnames(MCGC.mat.upper) <- edge.vec[[i]]
  col_fun = colorRamp2(c(0, 4.5, 9.5), c("white", "blue", "red"))
  
  if( i == 4 ){
    ht1 <- Heatmap(MCGC.mat.upper, cluster_rows = F, cluster_columns = F, col = col_fun, 
                   column_title = expression(atop("C"["R"],
                                                  italic("S. cerevisiae"))), 
                   row_title = expression(paste("C"["D"]^{"out"}, sep = "")), 
                   heatmap_legend_param = list(title = "R, mean - se", title_gp = gpar(fontsize = 30), 
                                               labels_gp = gpar(fontsize = 15)), 
                   use_raster = T, raster_quality = 4,
                   raster_device = "png", column_title_gp = gpar(fontsize = 30), column_title_side = "bottom", 
                   row_title_side = "left", row_names_side = "left", row_title_gp = gpar(fontsize = 30), 
                   row_names_gp = gpar(fontsize = 28), column_names_gp = gpar(fontsize = 28)) 
    ht1@matrix_param$border <- T
    ht1@heatmap_param$show_heatmap_legend <- F
    ht1@matrix_param$border <- T
    ht_list[[1]] <- ht1
  }else{
    if( i == 6 ){
      ht2 <- Heatmap(MCGC.mat.upper, cluster_rows = F, cluster_columns = F, col = col_fun, 
                     column_title = expression(atop("C"["R"],
                                                    italic("D. melanogaster"))), 
                     heatmap_legend_param = list(title = "R", title_gp = gpar(fontsize = 30), 
                                                 labels_gp = gpar(fontsize = 24)), 
                     use_raster = T, raster_quality = 4,
                     raster_device = "png", column_title_gp = gpar(fontsize = 30), column_title_side = "bottom", 
                     column_names_gp = gpar(fontsize = 28)) 
      ht2@heatmap_param$show_heatmap_legend <- F
      ht2@matrix_param$border <- T
      ht2@matrix_legend_param$border <- T
      ht_list[[2]] <- ht2
      
    }
    if( i == 8 ){
      ht3 <- Heatmap(MCGC.mat.upper, cluster_rows = F, cluster_columns = F, col = col_fun, 
                     column_title = expression(atop("C"["R"],
                                                    italic("M. musculus"))), 
                     heatmap_legend_param = list(title = "R, mean + se", title_gp = gpar(fontsize = 30), 
                                                 labels_gp = gpar(fontsize = 28)), 
                     use_raster = T, raster_quality = 4,
                     raster_device = "png", column_title_gp = gpar(fontsize = 30), column_title_side = "bottom", 
                     column_names_gp = gpar(fontsize = 28)) 
      ht3@heatmap_param$show_heatmap_legend <- F
      ht3@matrix_param$border <- T
      ht_list[[3]] <- ht3
    }
    if( i == 9 ){
      ht4 <- Heatmap(MCGC.mat.upper, cluster_rows = F, cluster_columns = F, col = col_fun, 
                     column_title = expression(atop("C"["R"],
                                                    italic("H. sapiens"))), 
                     heatmap_legend_param = list(title = "R", title_gp = gpar(fontsize = 30), 
                                                 labels_gp = gpar(fontsize = 24)), 
                     use_raster = T, raster_quality = 4,
                     raster_device = "png", column_title_gp = gpar(fontsize = 30), column_title_side = "bottom", 
                     column_names_gp = gpar(fontsize = 28)) 
      ht4@heatmap_param$show_heatmap_legend <- T
      ht4@matrix_param$border <- T
      ht4@matrix_legend_param$border <- T
      ht4@matrix_legend_param$legend_height <- unit(10, "cm")
      ht4@matrix_legend_param$grid_width <- unit(7, "mm")
      ht4@matrix_legend_param$gap <- unit(1, "cm")
      ht_list[[4]] <- ht4
    }
  }
}

ht_list1 <- ht1 + ht2 + ht3 + ht4
png(paste("Figure_4B_upper_CI.png", sep = ""), width = 720*4 + 80, height = 720)
draw(ht_list1, row_title = expression(paste("C"["D"]^{"in"}, "=2", "C"["D"]^{"out"}, sep = "")), 
     row_title_gp = gpar(fontsize = 34, fontface = "bold"), ht_gap = unit(1, "cm"))
dev.off()

##Figure 4C
edges.id <- rep(1, 9)
edges.id[4] <- 4
edges.id[6] <- 1
edges.id[8] <- 3
edges.id[9] <- 3
NumGenes <- rep(500, 9)
NumGenes[4] <- 1000
NumGenes[6] <- 2000
NumGenes[8] <- 300
NumGenes[9] <- 500
len.genes.Bio <- MCC.size.Bio <- rep(0, length(Organisms))
Plot.list.1 <- list()
database.type <- rep("BioGRID", 9)

cor.req <- cor.vec <- seq(0, 0.5, by = 0.1)
# cor.req <- seq(-0.8, 0.8, by = 0.2)


nboot <- 1000
dat.plot.red.Bio <- dat.plot.corr.Bio <- list()
for( i in (1:length(Organisms))[c(4, 6, 8, 9)] ){
  message(paste("Organism = ", Organisms[i], sep = ""))
  adj.TRN <- NetworksMultiplex$BioGrid[[i]]$TRN
  adj.PPI <- NetworksMultiplex$BioGrid[[i]]$PPI
  MCC.size.Bio[i] <- MCC.complete.graph(adj.TRN, adj.PPI)$size

  len.genes.Bio[i] <- ncol(adj.TRN)
  
  len <- ncol(adj.TRN)
  len <- NumGenes[i]
  
  id.cor <- 1:(length(cor.vec))
  
  MCC.redundancy <- array(0, c(length(id.cor), nboot))
  MCC.corr <- array(0, c(length(id.cor), nboot, 3))
  MCC.areas.slopes <- array(0, c(2, length(id.cor), nboot))
  
  if( i < 10 ){
    for( j in id.cor ){
      dir.org <- paste(getwd(), "/", Organisms_file[i], "_BioGrid_sample_degdeg_", cor.vec[j], "_red_", 
                       edge.vec[[i]][edges.id[i]], 
                       sep = "")
      dir.write <- paste(getwd(), "/", Organisms_file[i], "_BioGrid_sample_degdeg_", cor.vec[j], "_red_", 
                         edge.vec[[i]][edges.id[i]], "_Attack_curves", sep = "")
      for( b in 1:(1*nboot) ){
        b.id <- b
        MCGC.temp <- c(unlist(read.table(paste(dir.write, "/boot_", b, sep = ""), sep = "\t", header = F)))
        id1 <- c(unlist(read.table(paste(dir.org, "/boot_", b, sep = ""), sep = "\t", header = F))) + 1
        id2 <- id1
        MCC.redundancy[j, b.id] <- nnzero(adj.TRN[id1, id1]*adj.PPI[id2, id2])
        MCC.corr[j, b.id, ] <- SpearmanRho(rowSums(adj.TRN[id1, id1]), colSums(adj.PPI[id2, id2]), conf.level = 0.95)
        
        id.stop <- which(MCGC.temp <= 1)[1]
        if( id.stop >= 2 ){
          MCC.areas.slopes[1, j, b.id] <- abs(sintegral((1:id.stop)/len,
                                                        MCGC.temp[1:id.stop]/len)$value)
        }else{
          MCC.areas.slopes[1, j, b.id] <- 0
        }
      }
    }
  }
  
  MCC.rand <- red.rand <- rep(0, nboot)
  dir.org <- paste(getwd(), "/", Organisms_file[i], "_BioGrid_sample_degdeg_", cor.vec[1], "_red_", 
                   edge.vec[[i]][edges.id[i]], 
                   sep = "")
  dir.write <- paste(getwd(), "/", Organisms_file[i], "_BioGrid_sample_degdeg_", cor.vec[1], "_red_", 
                     edge.vec[[i]][edges.id[i]], "_Attack_curves", sep = "")
  for( b in 1:(1*nboot) ){
    MCGC.temp <- c(unlist(read.table(paste(dir.write, "/boot_", b, sep = ""), sep = "\t", header = F)))
    id.stop <- which(MCGC.temp <= 1)[1]
    if( id.stop >= 2 ){
      MCC.rand[b] <- abs(sintegral((1:id.stop)/len, 
                                   MCGC.temp[1:id.stop]/len)$value)
    }else{
      MCC.rand[b] <- 0
    }
  }
  
  tmp.MCC <- tmp.red <- tmp.corr <- matrix(0, length(id.cor) + 1, 3)
  if( i < 10 ){
    for( j in id.cor ){
      tmp <- orddom(MCC.rand, MCC.areas.slopes[1, j, ])
      tmp.MCC[j, ] <- c(as.numeric(tmp[23, 2]), as.numeric(tmp[24, 2]), 
                        as.numeric(tmp[25, 2]))
    }
  }
  tmp.corr[-c(length(id.cor) + 1), 1] <- apply(MCC.corr[, , 1], 1, function(x){
    return(t.test(x)$estimate)
  })
  
  tmp.corr[-c(length(id.cor) + 1), 2] <- apply(MCC.corr[, , 1], 1, function(x){
    return(t.test(x)$conf.int[1])
  })
  
  tmp.corr[-c(length(id.cor) + 1), 3] <- apply(MCC.corr[, , 1], 1, function(x){
    return(t.test(x)$conf.int[2])
  })
  
  if( i == 4 ){
    data.full.plot <- data.frame(x = rep(cor.vec/2, nboot), 
                                 y = c(MCC.areas.slopes[1, , ]), 
                                 Organism = rep(Organisms[i], length(cor.vec)*nboot))
  }else{
    data.tmp <- data.frame(x = rep(cor.vec/2, nboot), 
                           y = c(MCC.areas.slopes[1, , ]), 
                           Organism = rep(Organisms[i], length(cor.vec)*nboot))
    data.full.plot <- rbind(data.full.plot, data.tmp)
  }
  dat.plot.corr.Bio[[i]] <- data.frame(x = c(tmp.corr[-c(length(id.cor) + 1), 1]),
                                       x.lower = tmp.corr[-c(length(id.cor) + 1), 2],
                                       x.upper = tmp.corr[-c(length(id.cor) + 1), 3],
                                       y = rep(tmp.MCC[-c(length(id.cor) + 1), 1], 1),
                                       y.lower = tmp.MCC[-c(length(id.cor) + 1), 2], 
                                       y.upper = tmp.MCC[-c(length(id.cor) + 1), 3],
                                       Type = unlist(lapply(c(rep(Organisms[i], length(id.cor))), function(x){
                                         paste(x, "", sep = "")
                                       })))
}

data.full.plot$y <- data.full.plot$y + 1
p <- ggboxplot(data.full.plot, x = "x", y = "y",
               fill = "Organism", palette = c("red", "green", "blue", "gray")) + 
  yscale("log2", T)

###Plots
#Plots
count <- 1
tmp <- MCC.size.Bio/len.genes.Bio
tmp[is.nan(tmp)] <- 0
for( j in (1:length(Organisms))[c(4, 6, 8, 9)] ){
  if( tmp[j] >= 0.4 ){
    if( count == 1 ){
      data.plot <- dat.plot.corr.Bio[[j]]
    }else{
      data.plot <- rbind(data.plot, dat.plot.corr.Bio[[j]])
    }
    
    count <- count + 1
  }
}

data.plot$x <- c(matrix(rep(cor.vec, 4), length(cor.vec)))
data.tmp <- cbind(data.plot, data.frame(sim.type = c(rep("Simulation", 24))))
data.tmp$sim.type <- factor(data.tmp$sim.type, levels = c("Simulation"))
p <- ggline(data = data.tmp[1:24, ], x = "x", y = "y", 
            numeric.x.axis = T, 
            linetype = "dashed",
            plot_type = c("l"),
            shape = 16,
            fill = "Type",           # change fill color by mpg_level
            color = "Type",# jco journal color palett. see ?ggpar         # Sort the value in descending order
            # palette = c("red", "blue", "green", "gray"),
            sort.by.groups = FALSE,     # Don't sort inside each group
            x.text.angle = 50,          # Rotate vertically x axis texts
            ylab = "R",
            xlab = expression("C"["D"]),
            legend = "bottom",
            legend.title = "",
            y.text.angle = 50,
            rotate = F,
            font.x = 16, 
            font.y = 16,
            font.tickslab = 14,
            size = 0.9,
            point.size = 4,
            # font.label = 12,
            theme = theme_pubr()
) + ggtitle("BioGRID") +
  geom_errorbar(aes(ymin = y.lower, ymax = y.upper, color = Type), width = 0.01) 
p <- p + geom_point(data = data.tmp[1:24, ], mapping = aes(x = x, y = y, color = Type, fill = Type), shape = 16, 
                    size = 4) + 
  geom_errorbar(data = data.tmp[1:24, ], aes(ymin = y.lower, ymax = y.upper, color = Type), width = 0.01) + 
  scale_colour_manual(values = c(c("red", "blue", "green", "gray"))) + 
  geom_hline(yintercept = 0, linetype = 3)
p <- p +
  theme(title = element_text(size = 20, color = "black"), 
        plot.title = element_text(size = 18, color = "black"), 
        axis.title = element_text(size = 18, colour = "black"), 
        legend.text = element_text(size = 18, color = "black"))
Plot.fig.4[[2]] <- p


##Panel 4C, right degree-degree coupling fixed to org, redundancy increasing
edge.vec <- list()
edge.vec[[4]] <- seq(0, 40, by = 10)
edge.vec[[6]] <- seq(0, 80, length = 5)
edge.vec[[8]] <- seq(0, 40, length = 5)
edge.vec[[9]] <- seq(0, 100, length = 5)
NumGenes <- rep(500, 9)
NumGenes[4] <- 1000
NumGenes[6] <- 2000
NumGenes[8] <- 300
NumGenes[9] <- 500
nboot <- 1000
for( i in c(4, 6, 8, 9) ){
  # Start the clock!
  
  for( j in 1:length(edge.vec[[i]]) ){
    ##Print name of the organism.
    message(paste("Organism = ", Organisms[i], " edges = ", edge.vec[[i]][j], sep = ""))
    
    ##Get organism TRN and PPI networks from multiplex network object.
    adj.TRN <- NetworksMultiplex$BioGrid[[i]]$TRN
    adj.PPI <- NetworksMultiplex$BioGrid[[i]]$PPI
    rin <- cor(colSums(adj.TRN), colSums(adj.PPI), method = "spearman")
    rout <- cor(rowSums(adj.TRN), colSums(adj.PPI), method = "spearman")
    dir.org <- paste(getwd(), "/", Organisms_file[i], "_BioGrid_sample_degdeg_org", "_red_", edge.vec[[i]][j], 
                     sep = "")
    dir.create(dir.org, showWarnings = F)
    ptm <- proc.time()
    for( b in 1:nboot ){
      cat("boot = ", b, "\n")
      id <- c(SASampleMultiplex(adj.TRN, adj.PPI, colSums(adj.TRN), rowSums(adj.TRN), 
                                colSums(adj.PPI), edge.vec[[i]][j], 5, 1e8, 1e-9, 
                                0.01, 0.001, 10, 1e7, rin, 
                                rout, NumGenes[i]))
      file <- paste(dir.org, "/boot_", b, sep = "")
      write.table(cbind(id), file, sep = "\t", row.names = F, 
                  col.names = F, quote = F)
    }
    # Stop the clock
    proc.time() - ptm
    
    
    dir.write <- paste(getwd(), "/", Organisms_file[i], "_BioGrid_sample_degdeg_org", "_red_", 
                       edge.vec[[i]][j], "_Attack_curves", sep = "")
    dir.create(dir.write, showWarnings = F)
    system.time(id.rand <- MCGCAttackCurveSampleParallel(adj.TRN, adj.PPI, NumGenes[i], nboot, nthreads, 
                                                         dir.write, seed, 1, dir.org, 1:ncol(adj.TRN)))
  }
}


nboot <- 1000
dat.plot.red.Bio <- dat.plot.corr.Bio <- list()
for( i in (1:length(Organisms))[c(4, 6, 8, 9)] ){
  message(paste("Organism = ", Organisms[i], sep = ""))
  adj.TRN <- NetworksMultiplex$BioGrid[[i]]$TRN
  adj.PPI <- NetworksMultiplex$BioGrid[[i]]$PPI
  MCC.size.Bio[i] <- MCC.complete.graph(adj.TRN, adj.PPI)$size
  len.genes.Bio[i] <- ncol(adj.TRN)
  
  len <- ncol(adj.TRN)
  len <- NumGenes[i]
  
  id.red <- 1:(length(edge.vec[[i]]))
  
  MCC.redundancy <- array(0, c(length(id.red), nboot))
  MCC.corr <- array(0, c(length(id.red), nboot, 3))
  MCC.areas.slopes <- array(0, c(2, length(id.red), nboot))
  
  if( i < 10 ){
    for( j in id.red ){
      dir.org <- paste(getwd(), "/", Organisms_file[i], "_BioGrid_sample_degdeg_org", "_red_", 
                       edge.vec[[i]][j], sep = "")
      dir.write <- paste(getwd(), "/", Organisms_file[i], "_BioGrid_sample_degdeg_org", "_red_", 
                         edge.vec[[i]][j], "_Attack_curves", sep = "")
      for( b in 1:(1*nboot) ){
        b.id <- b
        MCGC.temp <- c(unlist(read.table(paste(dir.write, "/boot_", b, sep = ""), sep = "\t", header = F)))
        id1 <- c(unlist(read.table(paste(dir.org, "/boot_", b, sep = ""), sep = "\t", header = F))) + 1
        id2 <- id1
        MCC.redundancy[j, b.id] <- nnzero(adj.TRN[id1, id1]*adj.PPI[id2, id2])
        MCC.corr[j, b.id, ] <- SpearmanRho(rowSums(adj.TRN[id1, id1]), colSums(adj.PPI[id2, id2]), conf.level = 0.95)
        
        id.stop <- which(MCGC.temp <= 1)[1]
        if( id.stop >= 2 ){
          MCC.areas.slopes[1, j, b.id] <- abs(sintegral((1:id.stop)/len,
                                                        MCGC.temp[1:id.stop]/len)$value)
        }else{
          MCC.areas.slopes[1, j, b.id] <- 0
        }
        # id <- which(MCC.attack.curves.Graphlet.org.slopes.DA[[i]][[b]]$MCC.curve <= 1)[1]
      }
    }
  }
  
  MCC.rand <- red.rand <- rep(0, nboot)
  dir.org <- paste(getwd(), "/", Organisms_file[i], "_BioGrid_sample_degdeg_org", "_red_", 
                   edge.vec[[i]][1], sep = "")
  dir.write <- paste(getwd(), "/", Organisms_file[i], "_BioGrid_sample_degdeg_org", "_red_", 
                     edge.vec[[i]][1], "_Attack_curves", sep = "")
  for( b in 1:(1*nboot) ){
    MCGC.temp <- c(unlist(read.table(paste(dir.write, "/boot_", b, sep = ""), sep = "\t", header = F)))
    id.stop <- which(MCGC.temp <= 1)[1]
    if( id.stop >= 2 ){
      MCC.rand[b] <- abs(sintegral((1:id.stop)/len, 
                                   MCGC.temp[1:id.stop]/len)$value)
    }else{
      MCC.rand[b] <- 0
    }
  }
  
  tmp.MCC <- tmp.red <- tmp.corr <- matrix(0, length(id.red) + 1, 3)
  if( i < 10 ){
    for( j in id.red ){
      tmp <- orddom(MCC.rand, MCC.areas.slopes[1, j, ])
      tmp.MCC[j, ] <- c(as.numeric(tmp[23, 2]), as.numeric(tmp[24, 2]), 
                        as.numeric(tmp[25, 2]))
    }
  }
  tmp.corr[-c(length(id.red) + 1), 1] <- apply(MCC.corr[, , 1], 1, function(x){
    return(t.test(x)$estimate)
  })
  
  tmp.corr[-c(length(id.red) + 1), 2] <- apply(MCC.corr[, , 1], 1, function(x){
    return(t.test(x)$conf.int[1])
  })
  
  tmp.corr[-c(length(id.red) + 1), 3] <- apply(MCC.corr[, , 1], 1, function(x){
    return(t.test(x)$conf.int[2])
  })
  
  if( i == 4 ){
    data.full.plot <- data.frame(x = rep(edge.vec[[i]], nboot), 
                                 y = c(MCC.areas.slopes[1, , ]), 
                                 Organism = rep(Organisms[i], length(edge.vec[[i]])*nboot))
  }else{
    data.tmp <- data.frame(x = rep(edge.vec[[i]], nboot), 
                           y = c(MCC.areas.slopes[1, , ]), 
                           Organism = rep(Organisms[i], length(edge.vec[[i]])*nboot))
    data.full.plot <- rbind(data.full.plot, data.tmp)
  }
  dat.plot.corr.Bio[[i]] <- data.frame(x = edge.vec[[i]],
                                       x.lower = edge.vec[[i]],
                                       x.upper = edge.vec[[i]],
                                       y = rep(tmp.MCC[-c(length(id.red) + 1), 1], 1),
                                       y.lower = tmp.MCC[-c(length(id.red) + 1), 2], 
                                       y.upper = tmp.MCC[-c(length(id.red) + 1), 3],
                                       Type = unlist(lapply(c(rep(Organisms[i], length(id.red))), function(x){
                                         paste(x, "", sep = "")
                                       })))
  
}

count <- 1
tmp <- MCC.size.Bio/len.genes.Bio
tmp[is.nan(tmp)] <- 0
for( j in (1:length(Organisms))[c(4, 6, 8, 9)] ){
  if( tmp[j] >= 0.4 ){
    if( count == 1 ){
      data.plot <- dat.plot.corr.Bio[[j]]
    }else{
      data.plot <- rbind(data.plot, dat.plot.corr.Bio[[j]])
    }
    
    count <- count + 1
  }
}

data.tmp <- cbind(data.plot, data.frame(sim.type = c(rep("Simulation", 20))))
data.tmp$sim.type <- factor(data.tmp$sim.type, levels = c("Simulation"))
p <- ggline(data = data.tmp[1:20, ], x = "x", y = "y", 
            numeric.x.axis = T, 
            linetype = "dashed",
            plot_type = c("l"),
            shape = 16,
            fill = "Type",           # change fill color by mpg_level
            color = "Type",# jco journal color palett. see ?ggpar         # Sort the value in descending order
            # palette = c("red", "blue", "green", "gray"),
            sort.by.groups = FALSE,     # Don't sort inside each group
            x.text.angle = 50,          # Rotate vertically x axis texts
            ylab = "R",
            xlab = expression("C"["R"]),
            legend = "bottom",
            legend.title = "",
            y.text.angle = 50,
            rotate = F,
            font.x = 16, 
            font.y = 16,
            font.tickslab = 14,
            size = 0.9,
            point.size = 4,
            # font.label = 12,
            theme = theme_pubr()
) + ggtitle("BioGRID") +
  geom_errorbar(aes(ymin = y.lower, ymax = y.upper, color = Type), width = 3) 
p <- p + geom_point(data = data.tmp[1:20, ], mapping = aes(x = x, y = y, color = Type, fill = Type), shape = 16, 
                    size = 4) + 
  geom_errorbar(data = data.tmp[1:20, ], aes(ymin = y.lower, ymax = y.upper, color = Type), width = 0.01) + 
  scale_colour_manual(values = c(c("red", "blue", "green", "gray"))) + 
  geom_hline(yintercept = 0, linetype = 3)
p <- p + 
  theme(title = element_text(size = 20, color = "black"), 
        plot.title = element_text(size = 18, color = "black"), 
        axis.title = element_text(size = 18, colour = "black"), 
        legend.text = element_text(size = 16, color = "black"))
Plot.fig.4[[3]] <- p

##Figures. 4A and 4C
plot.2 <- plot_grid(Plot.fig.4[[1]], Plot.fig.4[[2]], Plot.fig.4[[3]], nrow = 1, labels = "", label_size = 35, 
                    align = "h", axis = "left", rel_widths = c(2, 1, 1))
save_plot(paste("Figure_4A_4C.png", sep = ""), 
          plot.2, base_height = 6, base_width = 6*4)
```
