## Supplementary material for "Internetwork connectivity of molecular networks across species of life": Code: Figure_5_Paper.R

```
#Figure 5
##set seed for reproducibility
set.seed(1)

##Load packages and functions
library(Rcpp)
library(Matrix)
library(igraph)
library(ggplot2)
library(ggpubr)
library(cowplot)
Rcpp::sourceCpp('utils.cpp')
Rcpp::sourceCpp("Attack_functions_new_cpp.cpp")
Rcpp::sourceCpp('Simulated_Annealing_Functions.cpp')

##Generate data for Figure 5A
##Essential (Lethal) genes HINT
load("Lethality_HINT.RData")

Organisms <- c("H. pylori", "M.tb",        "E. coli",     "Yeast",       "C. elegans",  "Fly", 
               "A. thaliana", "Mouse", "Human")
Organisms_file <- c("H_pylori", "M_tb",        "E_coli",     "Yeast",       "C_elegans",  "Fly", 
                    "A_thaliana", "Mouse", "Human")
nthreads <- 10
nboot <- 100
seed <- 123
for( i in c(4, 6, 8, 9) ){
  
  ##Print name of the organism.
  message(paste("Organism = ", Organisms[i], sep = ""))
  
  ##Get organism TRN and PPI networks from multiplex network object.
  adj.TRN <- NetworksMultiplex$HINT[[i]]$TRN
  adj.PPI <- NetworksMultiplex$HINT[[i]]$PPI
  id.inter <- NetworksMultiplex$HINT[[i]]$genes
  
  genes.lethal <- toupper(Lethality.HINT$genes[[i]])[Lethality.HINT$gene.lethality[[i]] == 0]
  id.match <- match(toupper(id.inter), toupper(genes.lethal))
  id.lethal <- which(!is.na(id.match))
  
  genes.nonlethal <- toupper(Lethality.HINT$genes[[i]])[Lethality.HINT$gene.lethality[[i]] != 0]
  id.match <- match(toupper(id.inter), toupper(genes.nonlethal))
  id.nonlethal <- which(!is.na(id.match))
  
  ###Attack curves attack on disease genes and proteins
  id1 <- 1:ncol(adj.TRN)
  id2 <- 1:ncol(adj.TRN)
  adj1 <- adj.TRN[id1, id1]
  adj2 <- adj.PPI[id2, id2]
  elTRN <- get.edgelist(graph_from_adjacency_matrix(adj1, "undirected")) - 1
  elPPI <- get.edgelist(graph_from_adjacency_matrix(adj2, "undirected")) - 1

  ##Make organism specific folder to store the attack curves
  dir.write <- paste(getwd(), "/", Organisms_file[i], "_HINT_", 
                     "Lethal_attack_curve", sep = "")
  dir.create(dir.write, showWarnings = F)
  
  system.time(tmp.curve.rand <- MCGCAttackCurveFunctionalGenesParallel(elTRN, elPPI, ncol(adj.TRN), rowSums(adj.TRN), 
                                                                       colSums(adj.PPI), nboot, nthreads, dir.write, seed, 0, dir.write, 
                                                                       length(id.lethal), id.lethal - 1))
}

for( i in c(4, 6, 8, 9) ){
  
  ##Print name of the organism.
  message(paste("Organism = ", Organisms[i], sep = ""))
  
  ##Get organism TRN and PPI networks from multiplex network object.
  adj.TRN <- NetworksMultiplex$HINT[[i]]$TRN
  adj.PPI <- NetworksMultiplex$HINT[[i]]$PPI
  id.inter <- NetworksMultiplex$HINT[[i]]$genes
  
  genes.lethal <- toupper(Lethality.HINT$genes[[i]])[Lethality.HINT$gene.lethality[[i]] == 0]
  id.match <- match(toupper(id.inter), toupper(genes.lethal))
  id.lethal <- which(!is.na(id.match))
  
  genes.nonlethal <- toupper(Lethality.HINT$genes[[i]])[Lethality.HINT$gene.lethality[[i]] != 0]
  id.match <- match(toupper(id.inter), toupper(genes.nonlethal))
  id.nonlethal <- which(!is.na(id.match))
  
  dir.id <- paste(getwd(), "/", Organisms_file[i], "_HINT_", 
                  "nonLethal_ids", sep = "")
  dir.create(dir.id, showWarnings = F)
  for( b in 1:nboot ){
    # cat("boot = ", b, "\n")
    id <- sample(id.nonlethal, length(id.lethal)) - 1
    file <- paste(dir.id, "/boot_", b, sep = "")
    write.table(cbind(id), file, sep = "\t", row.names = F, 
                col.names = F, quote = F)
  }
  
  ###Attack curves attack on disease genes and proteins
  id1 <- 1:ncol(adj.TRN)
  id2 <- 1:ncol(adj.TRN)
  adj1 <- adj.TRN[id1, id1]
  adj2 <- adj.PPI[id2, id2]
  elTRN <- get.edgelist(graph_from_adjacency_matrix(adj1, "undirected")) - 1
  elPPI <- get.edgelist(graph_from_adjacency_matrix(adj2, "undirected")) - 1
  
  ##Make organism specific folder to store the attack curves
  dir.write <- paste(getwd(), "/", Organisms_file[i], "_HINT_", 
                     "nonLethal_attack_curve", sep = "")
  dir.create(dir.write, showWarnings = F)
  
  system.time(tmp.curve.rand <- MCGCAttackCurveFunctionalGenesParallel(elTRN, elPPI, ncol(adj.TRN), rowSums(adj.TRN), 
                                                                       colSums(adj.PPI), nboot, nthreads, dir.write, 
                                                                       seed, 1, dir.id, 
                                                                       length(id.lethal), id.lethal - 1))
}

for( i in c(4, 6, 8, 9) ){
  
  ##Print name of the organism.
  message(paste("Organism = ", Organisms[i], sep = ""))
  
  ##Get organism TRN and PPI networks from multiplex network object.
  adj.TRN <- NetworksMultiplex$HINT[[i]]$TRN
  adj.PPI <- NetworksMultiplex$HINT[[i]]$PPI
  id.inter <- NetworksMultiplex$HINT[[i]]$genes
  
  genes.lethal <- toupper(Lethality.HINT$genes[[i]])[Lethality.HINT$gene.lethality[[i]] == 0]
  id.match <- match(toupper(id.inter), toupper(genes.lethal))
  id.lethal <- which(!is.na(id.match))
  
  genes.nonlethal <- toupper(Lethality.HINT$genes[[i]])[Lethality.HINT$gene.lethality[[i]] != 0]
  id.match <- match(toupper(id.inter), toupper(genes.nonlethal))
  id.nonlethal <- which(!is.na(id.match))
  
  dir.id <- paste(getwd(), "/", Organisms_file[i], "_HINT_",
                  "randomLethal_ids", sep = "")
  dir.create(dir.id, showWarnings = F)
  for( b in 1:nboot ){
    # cat("boot = ", b, "\n")
    id <- sample(c(id.lethal, id.nonlethal), length(id.lethal)) - 1
    file <- paste(dir.id, "/boot_", b, sep = "")
    write.table(cbind(id), file, sep = "\t", row.names = F, 
                col.names = F, quote = F)
  }
  
  ###Attack curves attack on disease genes and proteins
  id1 <- 1:ncol(adj.TRN)
  id2 <- 1:ncol(adj.TRN)
  adj1 <- adj.TRN[id1, id1]
  adj2 <- adj.PPI[id2, id2]
  elTRN <- get.edgelist(graph_from_adjacency_matrix(adj1, "undirected")) - 1
  elPPI <- get.edgelist(graph_from_adjacency_matrix(adj2, "undirected")) - 1
  
  ##Make organism specific folder to store the attack curves
  dir.write <- paste(getwd(), "/", Organisms_file[i], "_HINT_",
                     "randomLethal_attack_curve", sep = "")
  dir.create(dir.write, showWarnings = F)
  
  system.time(tmp.curve.rand <- MCGCAttackCurveFunctionalGenesParallel(elTRN, elPPI, ncol(adj.TRN), rowSums(adj.TRN), 
                                                                       colSums(adj.PPI), nboot, nthreads, dir.write, 
                                                                       seed, 1, dir.id, 
                                                                       length(id.lethal), id.lethal - 1))
}


##Essential (Lethal) genes BioGRID
load("Lethality_BioGRID.RData")
Organisms <- c("H. pylori", "M.tb",        "E. coli",     "Yeast",       "C. elegans",  "Fly", 
               "A. thaliana", "Mouse", "Human")
Organisms_file <- c("H_pylori", "M_tb",        "E_coli",     "Yeast",       "C_elegans",  "Fly", 
                    "A_thaliana", "Mouse", "Human")
nthreads <- 10
nboot <- 100
seed <- 123
i <- 9
for( i in c(4, 6, 8, 9) ){
  
  ##Print name of the organism.
  message(paste("Organism = ", Organisms[i], sep = ""))
  
  ##Get organism TRN and PPI networks from multiplex network object.
  adj.TRN <- NetworksMultiplex$BioGrid[[i]]$TRN
  adj.PPI <- NetworksMultiplex$BioGrid[[i]]$PPI
  id.inter <- NetworksMultiplex$BioGrid[[i]]$genes
  
  genes.lethal <- toupper(Lethality.BioGRID$genes[[i]])[Lethality.BioGRID$gene.lethality[[i]] == 0]
  id.match <- match(toupper(id.inter), toupper(genes.lethal))
  id.lethal <- which(!is.na(id.match))
  
  genes.nonlethal <- toupper(Lethality.BioGRID$genes[[i]])[Lethality.BioGRID$gene.lethality[[i]] != 0]
  id.match <- match(toupper(id.inter), toupper(genes.nonlethal))
  id.nonlethal <- which(!is.na(id.match))
  
  ###Attack curves attack on disease genes and proteins
  id1 <- 1:ncol(adj.TRN)
  id2 <- 1:ncol(adj.TRN)
  adj1 <- adj.TRN[id1, id1]
  adj2 <- adj.PPI[id2, id2]
  elTRN <- get.edgelist(graph_from_adjacency_matrix(adj1, "undirected")) - 1
  elPPI <- get.edgelist(graph_from_adjacency_matrix(adj2, "undirected")) - 1
  
  ##Make organism specific folder to store the attack curves
  dir.write <- paste(getwd(), "/", Organisms_file[i], "_BioGrid_", 
                     "Lethal_attack_curve", sep = "")
  dir.create(dir.write, showWarnings = F)
  
  system.time(tmp.curve.rand <- MCGCAttackCurveFunctionalGenesParallel(elTRN, elPPI, ncol(adj.TRN), rowSums(adj.TRN), 
                                                                       colSums(adj.PPI), nboot, nthreads, dir.write, seed, 0, dir.write, 
                                                                       length(id.lethal), id.lethal - 1))
}


for( i in c(4, 6, 8, 9) ){
  
  ##Print name of the organism.
  message(paste("Organism = ", Organisms[i], sep = ""))
  
  ##Get organism TRN and PPI networks from multiplex network object.
  adj.TRN <- NetworksMultiplex$BioGrid[[i]]$TRN
  adj.PPI <- NetworksMultiplex$BioGrid[[i]]$PPI
  id.inter <- NetworksMultiplex$BioGrid[[i]]$genes
  
  genes.lethal <- toupper(Lethality.BioGRID$genes[[i]])[Lethality.BioGRID$gene.lethality[[i]] == 0]
  id.match <- match(toupper(id.inter), toupper(genes.lethal))
  id.lethal <- which(!is.na(id.match))
  
  genes.nonlethal <- toupper(Lethality.BioGRID$genes[[i]])[Lethality.BioGRID$gene.lethality[[i]] != 0]
  id.match <- match(toupper(id.inter), toupper(genes.nonlethal))
  id.nonlethal <- which(!is.na(id.match))
  
  dir.id <- paste(getwd(), "/", Organisms_file[i], "_BioGrid_",
                  "nonLethal_ids", sep = "")
  dir.create(dir.id, showWarnings = F)
  for( b in 1:nboot ){
    # cat("boot = ", b, "\n")
    id <- sample(id.nonlethal, length(id.lethal)) - 1
    file <- paste(dir.id, "/boot_", b, sep = "")
    write.table(cbind(id), file, sep = "\t", row.names = F, 
                col.names = F, quote = F)
  }
  
  ###Attack curves attack on disease genes and proteins
  id1 <- 1:ncol(adj.TRN)
  id2 <- 1:ncol(adj.TRN)
  adj1 <- adj.TRN[id1, id1]
  adj2 <- adj.PPI[id2, id2]
  elTRN <- get.edgelist(graph_from_adjacency_matrix(adj1, "undirected")) - 1
  elPPI <- get.edgelist(graph_from_adjacency_matrix(adj2, "undirected")) - 1
  
  ##Make organism specific folder to store the attack curves
  dir.write <- paste(getwd(), "/", Organisms_file[i], "_BioGrid_", 
                     "nonLethal_attack_curve", sep = "")
  dir.create(dir.write, showWarnings = F)
  
  system.time(tmp.curve.rand <- MCGCAttackCurveFunctionalGenesParallel(elTRN, elPPI, ncol(adj.TRN), rowSums(adj.TRN), 
                                                                       colSums(adj.PPI), nboot, nthreads, dir.write, 
                                                                       seed, 1, dir.id, 
                                                                       length(id.lethal), id.lethal - 1))
}


for( i in c(4, 6, 8, 9) ){
  
  ##Print name of the organism.
  message(paste("Organism = ", Organisms[i], sep = ""))
  
  ##Get organism TRN and PPI networks from multiplex network object.
  adj.TRN <- NetworksMultiplex$BioGrid[[i]]$TRN
  adj.PPI <- NetworksMultiplex$BioGrid[[i]]$PPI
  id.inter <- NetworksMultiplex$BioGrid[[i]]$genes
  
  genes.lethal <- toupper(Lethality.BioGRID$genes[[i]])[Lethality.BioGRID$gene.lethality[[i]] == 0]
  id.match <- match(toupper(id.inter), toupper(genes.lethal))
  id.lethal <- which(!is.na(id.match))
  
  genes.nonlethal <- toupper(Lethality.BioGRID$genes[[i]])[Lethality.BioGRID$gene.lethality[[i]] != 0]
  id.match <- match(toupper(id.inter), toupper(genes.nonlethal))
  id.nonlethal <- which(!is.na(id.match))
  
  dir.id <- paste(getwd(), "/", Organisms_file[i], "_BioGrid_",
                  "randomLethal_ids", sep = "")
  dir.create(dir.id, showWarnings = F)
  for( b in 1:nboot ){
    # cat("boot = ", b, "\n")
    id <- sample(c(id.lethal, id.nonlethal), length(id.lethal)) - 1
    file <- paste(dir.id, "/boot_", b, sep = "")
    write.table(cbind(id), file, sep = "\t", row.names = F, 
                col.names = F, quote = F)
  }
  
  ###Attack curves attack on disease genes and proteins
  id1 <- 1:ncol(adj.TRN)
  id2 <- 1:ncol(adj.TRN)
  adj1 <- adj.TRN[id1, id1]
  adj2 <- adj.PPI[id2, id2]
  elTRN <- get.edgelist(graph_from_adjacency_matrix(adj1, "undirected")) - 1
  elPPI <- get.edgelist(graph_from_adjacency_matrix(adj2, "undirected")) - 1
  
  ##Make organism specific folder to store the attack curves
  dir.write <- paste(getwd(), "/", Organisms_file[i], "_BioGrid_",
                     "randomLethal_attack_curve", sep = "")
  dir.create(dir.write, showWarnings = F)
  
  system.time(tmp.curve.rand <- MCGCAttackCurveFunctionalGenesParallel(elTRN, elPPI, ncol(adj.TRN), rowSums(adj.TRN), 
                                                                       colSums(adj.PPI), nboot, nthreads, dir.write, 
                                                                       seed, 1, dir.id, 
                                                                       length(id.lethal), id.lethal - 1))
}

## Generate data for Figure 5C
load("Genes_Diseases.RData")
load("Diseases.RData")

##Diseases HINT
i <- 9
for( j in c(1:length(Diseases)) ){
  
  ##Print name of the organism.
  message(paste("Disease = ", Diseases[j], sep = ""))
  
  ##Get organism TRN and PPI networks from multiplex network object.
  adj.TRN <- NetworksMultiplex$HINT[[i]]$TRN
  adj.PPI <- NetworksMultiplex$HINT[[i]]$PPI
  id.inter <- NetworksMultiplex$HINT[[i]]$genes
  
  id.match <- match(toupper(id.inter), toupper(Genes.Diseases[[j]]))
  id.disease <- which(!is.na(id.match))
  id.non.disease <- setdiff(1:length(id.inter), id.disease)
  
  ###Attack curves attack on disease genes and proteins
  id1 <- 1:ncol(adj.TRN)
  id2 <- 1:ncol(adj.TRN)
  adj1 <- adj.TRN[id1, id1]
  adj2 <- adj.PPI[id2, id2]
  elTRN <- get.edgelist(graph_from_adjacency_matrix(adj1, "undirected")) - 1
  elPPI <- get.edgelist(graph_from_adjacency_matrix(adj2, "undirected")) - 1
  
  ##Make organism specific folder to store the attack curves
  dir.write <- paste(getwd(), "/", Organisms_file[i], "_HINT_", sub(" ", "", paste(Diseases[j], sep = "_")), 
                     "_disease_attack_curve", sep = "")
  dir.create(dir.write, showWarnings = F)
  
  system.time(tmp.curve.rand <- MCGCAttackCurveFunctionalGenesParallel(elTRN, elPPI, ncol(adj.TRN), rowSums(adj.TRN), 
                                                                       colSums(adj.PPI), nboot, nthreads, dir.write, seed, 0, dir.write, 
                                                                       length(id.disease), id.disease - 1))
}


for( j in c(1:length(Diseases)) ){
  
  ##Print name of the organism.
  message(paste("Disease = ", Diseases[j], sep = ""))
  
  ##Get organism TRN and PPI networks from multiplex network object.
  adj.TRN <- NetworksMultiplex$HINT[[i]]$TRN
  adj.PPI <- NetworksMultiplex$HINT[[i]]$PPI
  id.inter <- NetworksMultiplex$HINT[[i]]$genes
  
  id.match <- match(toupper(id.inter), toupper(Genes.Diseases[[j]]))
  id.disease <- which(!is.na(id.match))
  id.non.disease <- setdiff(1:length(id.inter), id.disease)
  dir.id <- paste(getwd(), "/", Organisms_file[i], "_HINT_", sub(" ", "", paste(Diseases[j], sep = "_")), 
                  "_nondisease_ids", sep = "")
  dir.create(dir.id, showWarnings = F)
  for( b in 1:nboot ){
    # cat("boot = ", b, "\n")
    id <- sample(id.non.disease, length(id.disease)) - 1
    file <- paste(dir.id, "/boot_", b, sep = "")
    write.table(cbind(id), file, sep = "\t", row.names = F, 
                col.names = F, quote = F)
  }
  
  ###Attack curves attack on disease genes and proteins
  id1 <- 1:ncol(adj.TRN)
  id2 <- 1:ncol(adj.TRN)
  adj1 <- adj.TRN[id1, id1]
  adj2 <- adj.PPI[id2, id2]
  elTRN <- get.edgelist(graph_from_adjacency_matrix(adj1, "undirected")) - 1
  elPPI <- get.edgelist(graph_from_adjacency_matrix(adj2, "undirected")) - 1
  
  ##Make organism specific folder to store the attack curves
  dir.write <- paste(getwd(), "/", Organisms_file[i], "_HINT_", sub(" ", "", paste(Diseases[j], sep = "_")), 
                     "_nondisease_attack_curve", sep = "")
  dir.create(dir.write, showWarnings = F)
  
  system.time(tmp.curve.rand <- MCGCAttackCurveFunctionalGenesParallel(elTRN, elPPI, ncol(adj.TRN), rowSums(adj.TRN), 
                                                                       colSums(adj.PPI), nboot, nthreads, dir.write, 
                                                                       seed, 1, dir.id, 
                                                                       length(id.disease), id.disease - 1))
}


for( j in c(1:length(Diseases)) ){
  
  ##Print name of the organism.
  message(paste("Disease = ", Diseases[j], sep = ""))
  
  ##Get organism TRN and PPI networks from multiplex network object.
  adj.TRN <- NetworksMultiplex$HINT[[i]]$TRN
  adj.PPI <- NetworksMultiplex$HINT[[i]]$PPI
  id.inter <- NetworksMultiplex$HINT[[i]]$genes
  
  id.match <- match(toupper(id.inter), toupper(Genes.Diseases[[j]]))
  id.disease <- which(!is.na(id.match))
  id.non.disease <- setdiff(1:length(id.inter), id.disease)
  dir.id <- paste(getwd(), "/", Organisms_file[i], "_HINT_", sub(" ", "", paste(Diseases[j], sep = "_")), 
                  "_random_ids", sep = "")
  dir.create(dir.id, showWarnings = F)
  for( b in 1:nboot ){
    # cat("boot = ", b, "\n")
    id <- sample(c(id.disease, id.non.disease), length(id.disease)) - 1
    file <- paste(dir.id, "/boot_", b, sep = "")
    write.table(cbind(id), file, sep = "\t", row.names = F, 
                col.names = F, quote = F)
  }
  
  ###Attack curves attack on disease genes and proteins
  id1 <- 1:ncol(adj.TRN)
  id2 <- 1:ncol(adj.TRN)
  adj1 <- adj.TRN[id1, id1]
  adj2 <- adj.PPI[id2, id2]
  elTRN <- get.edgelist(graph_from_adjacency_matrix(adj1, "undirected")) - 1
  elPPI <- get.edgelist(graph_from_adjacency_matrix(adj2, "undirected")) - 1
  
  ##Make organism specific folder to store the attack curves
  dir.write <- paste(getwd(), "/", Organisms_file[i], "_HINT_", sub(" ", "", paste(Diseases[j], sep = "_")), 
                     "_random_attack_curve", sep = "")
  dir.create(dir.write, showWarnings = F)
  
  system.time(tmp.curve.rand <- MCGCAttackCurveFunctionalGenesParallel(elTRN, elPPI, ncol(adj.TRN), rowSums(adj.TRN), 
                                                                       colSums(adj.PPI), nboot, nthreads, dir.write, 
                                                                       seed, 1, dir.id, 
                                                                       length(id.disease), id.disease - 1))
}


##Diseases BioGRID
i <- 9
for( j in c(1:length(Diseases)) ){
  
  ##Print name of the organism.
  message(paste("Disease = ", Diseases[j], sep = ""))
  
  ##Get organism TRN and PPI networks from multiplex network object.
  adj.TRN <- NetworksMultiplex$BioGrid[[i]]$TRN
  adj.PPI <- NetworksMultiplex$BioGrid[[i]]$PPI
  id.inter <- NetworksMultiplex$BioGrid[[i]]$genes
  
  id.match <- match(toupper(id.inter), toupper(Genes.Diseases[[j]]))
  id.disease <- which(!is.na(id.match))
  id.non.disease <- setdiff(1:length(id.inter), id.disease)
  
  ###Attack curves attack on disease genes and proteins
  id1 <- 1:ncol(adj.TRN)
  id2 <- 1:ncol(adj.TRN)
  adj1 <- adj.TRN[id1, id1]
  adj2 <- adj.PPI[id2, id2]
  elTRN <- get.edgelist(graph_from_adjacency_matrix(adj1, "undirected")) - 1
  elPPI <- get.edgelist(graph_from_adjacency_matrix(adj2, "undirected")) - 1
  
  ##Make organism specific folder to store the attack curves
  dir.write <- paste(getwd(), "/", Organisms_file[i], "_BioGrid_", sub(" ", "", paste(Diseases[j], sep = "_")), 
                     "_disease_attack_curve", sep = "")
  dir.create(dir.write, showWarnings = F)
  
  system.time(tmp.curve.rand <- MCGCAttackCurveFunctionalGenesParallel(elTRN, elPPI, ncol(adj.TRN), rowSums(adj.TRN), 
                                                                       colSums(adj.PPI), nboot, nthreads, dir.write, seed, 0, dir.write, 
                                                                       length(id.disease), id.disease - 1))
}


for( j in c(1:length(Diseases)) ){
  
  ##Print name of the organism.
  message(paste("Disease = ", Diseases[j], sep = ""))
  
  ##Get organism TRN and PPI networks from multiplex network object.
  adj.TRN <- NetworksMultiplex$BioGrid[[i]]$TRN
  adj.PPI <- NetworksMultiplex$BioGrid[[i]]$PPI
  id.inter <- NetworksMultiplex$BioGrid[[i]]$genes
  
  id.match <- match(toupper(id.inter), toupper(Genes.Diseases[[j]]))
  id.disease <- which(!is.na(id.match))
  id.non.disease <- setdiff(1:length(id.inter), id.disease)
  dir.id <- paste(getwd(), "/", Organisms_file[i], "_BioGrid_", sub(" ", "", paste(Diseases[j], sep = "_")), 
                  "_nondisease_ids", sep = "")
  dir.create(dir.id, showWarnings = F)
  for( b in 1:nboot ){
    # cat("boot = ", b, "\n")
    id <- sample(id.non.disease, length(id.disease)) - 1
    file <- paste(dir.id, "/boot_", b, sep = "")
    write.table(cbind(id), file, sep = "\t", row.names = F, 
                col.names = F, quote = F)
  }
  
  ###Attack curves attack on disease genes and proteins
  id1 <- 1:ncol(adj.TRN)
  id2 <- 1:ncol(adj.TRN)
  adj1 <- adj.TRN[id1, id1]
  adj2 <- adj.PPI[id2, id2]
  elTRN <- get.edgelist(graph_from_adjacency_matrix(adj1, "undirected")) - 1
  elPPI <- get.edgelist(graph_from_adjacency_matrix(adj2, "undirected")) - 1
  
  ##Make organism specific folder to store the attack curves
  dir.write <- paste(getwd(), "/", Organisms_file[i], "_BioGrid_", sub(" ", "", paste(Diseases[j], sep = "_")), 
                     "_nondisease_attack_curve", sep = "")
  dir.create(dir.write, showWarnings = F)
  
  system.time(tmp.curve.rand <- MCGCAttackCurveFunctionalGenesParallel(elTRN, elPPI, ncol(adj.TRN), rowSums(adj.TRN), 
                                                                       colSums(adj.PPI), nboot, nthreads, dir.write, 
                                                                       seed, 1, dir.id, 
                                                                       length(id.disease), id.disease - 1))
}


for( j in c(1:length(Diseases)) ){
  
  ##Print name of the organism.
  message(paste("Disease = ", Diseases[j], sep = ""))
  
  ##Get organism TRN and PPI networks from multiplex network object.
  adj.TRN <- NetworksMultiplex$BioGrid[[i]]$TRN
  adj.PPI <- NetworksMultiplex$BioGrid[[i]]$PPI
  id.inter <- NetworksMultiplex$BioGrid[[i]]$genes
  
  id.match <- match(toupper(id.inter), toupper(Genes.Diseases[[j]]))
  id.disease <- which(!is.na(id.match))
  id.non.disease <- setdiff(1:length(id.inter), id.disease)
  dir.id <- paste(getwd(), "/", Organisms_file[i], "_BioGrid_", sub(" ", "", paste(Diseases[j], sep = "_")), 
                  "_random_ids", sep = "")
  dir.create(dir.id, showWarnings = F)
  for( b in 1:nboot ){
    # cat("boot = ", b, "\n")
    id <- sample(c(id.disease, id.non.disease), length(id.disease)) - 1
    file <- paste(dir.id, "/boot_", b, sep = "")
    write.table(cbind(id), file, sep = "\t", row.names = F, 
                col.names = F, quote = F)
  }
  
  ###Attack curves attack on disease genes and proteins
  id1 <- 1:ncol(adj.TRN)
  id2 <- 1:ncol(adj.TRN)
  adj1 <- adj.TRN[id1, id1]
  adj2 <- adj.PPI[id2, id2]
  elTRN <- get.edgelist(graph_from_adjacency_matrix(adj1, "undirected")) - 1
  elPPI <- get.edgelist(graph_from_adjacency_matrix(adj2, "undirected")) - 1
  
  ##Make organism specific folder to store the attack curves
  dir.write <- paste(getwd(), "/", Organisms_file[i], "_BioGrid_", sub(" ", "", paste(Diseases[j], sep = "_")), 
                     "_random_attack_curve", sep = "")
  dir.create(dir.write, showWarnings = F)
  
  system.time(tmp.curve.rand <- MCGCAttackCurveFunctionalGenesParallel(elTRN, elPPI, ncol(adj.TRN), rowSums(adj.TRN), 
                                                                       colSums(adj.PPI), nboot, nthreads, dir.write, 
                                                                       seed, 1, dir.id, 
                                                                       length(id.disease), id.disease - 1))
}


##Pathogens
## Generate data for Figure 5B
load("Genes_Pathogen.RData")
load("Pathogens.RData")

##Pathogen HINT
for( j in c(1:length(Pathogens)) ){
  
  ##Print name of the organism.
  message(paste("Pathogen = ", Pathogens[j], sep = ""))
  
  ##Get organism TRN and PPI networks from multiplex network object.
  adj.TRN <- NetworksMultiplex$HINT[[i]]$TRN
  adj.PPI <- NetworksMultiplex$HINT[[i]]$PPI
  id.inter <- NetworksMultiplex$HINT[[i]]$genes
  
  id.match <- match(toupper(id.inter), toupper(Genes.HPI[[j]]))
  id.disease <- which(!is.na(id.match))
  id.non.disease <- setdiff(1:length(id.inter), id.disease)
  
  ###Attack curves attack on disease genes and proteins
  id1 <- 1:ncol(adj.TRN)
  id2 <- 1:ncol(adj.TRN)
  adj1 <- adj.TRN[id1, id1]
  adj2 <- adj.PPI[id2, id2]
  elTRN <- get.edgelist(graph_from_adjacency_matrix(adj1, "undirected")) - 1
  elPPI <- get.edgelist(graph_from_adjacency_matrix(adj2, "undirected")) - 1
  
  ##Make organism specific folder to store the attack curves
  dir.write <- paste(getwd(), "/", Organisms_file[i], "_HINT_", sub("\\.", "", 
                                                                    sub(" ", "", paste(Pathogens[j], sep = "_"))), 
                     "_pathogen_attack_curve", sep = "")
  dir.create(dir.write, showWarnings = F)
  
  system.time(tmp.curve.rand <- MCGCAttackCurveFunctionalGenesParallel(elTRN, elPPI, ncol(adj.TRN), rowSums(adj.TRN), 
                                                                       colSums(adj.PPI), nboot, nthreads, dir.write, seed, 0, dir.write, 
                                                                       length(id.disease), id.disease - 1))
}


for( j in c(1:length(Pathogens)) ){
  
  ##Print name of the organism.
  message(paste("Pathogen = ", Pathogens[j], sep = ""))
  
  ##Get organism TRN and PPI networks from multiplex network object.
  adj.TRN <- NetworksMultiplex$HINT[[i]]$TRN
  adj.PPI <- NetworksMultiplex$HINT[[i]]$PPI
  id.inter <- NetworksMultiplex$HINT[[i]]$genes
  
  id.match <- match(toupper(id.inter), toupper(Genes.HPI[[j]]))
  id.disease <- which(!is.na(id.match))
  id.non.disease <- setdiff(1:length(id.inter), id.disease)
  # dir.id <- paste(getwd(), "/", Organisms_file[i], "_HINT_", sub(" ", "", paste(Diseases[j], sep = "_")), 
  #                 "_nondisease_ids", sep = "")
  dir.id <- paste(getwd(), "/", Organisms_file[i], "_HINT_", sub("\\.", "", 
                                                                 sub(" ", "", paste(Pathogens[j], sep = "_"))), 
                  "_nonpathogen_ids", sep = "")
  dir.create(dir.id, showWarnings = F)
  for( b in 1:nboot ){
    # cat("boot = ", b, "\n")
    id <- sample(id.non.disease, length(id.disease)) - 1
    file <- paste(dir.id, "/boot_", b, sep = "")
    write.table(cbind(id), file, sep = "\t", row.names = F, 
                col.names = F, quote = F)
  }
  
  ###Attack curves attack on disease genes and proteins
  id1 <- 1:ncol(adj.TRN)
  id2 <- 1:ncol(adj.TRN)
  adj1 <- adj.TRN[id1, id1]
  adj2 <- adj.PPI[id2, id2]
  elTRN <- get.edgelist(graph_from_adjacency_matrix(adj1, "undirected")) - 1
  elPPI <- get.edgelist(graph_from_adjacency_matrix(adj2, "undirected")) - 1
  
  ##Make organism specific folder to store the attack curves
  # dir.write <- paste(getwd(), "/", Organisms_file[i], "_HINT_", sub(" ", "", paste(Diseases[j], sep = "_")), 
  #                    "_nondisease_attack_curve", sep = "")
  dir.write <- paste(getwd(), "/", Organisms_file[i], "_HINT_", sub("\\.", "", 
                                                                    sub(" ", "", paste(Pathogens[j], sep = "_"))), 
                     "_nonpathogen_attack_curve", sep = "")
  dir.create(dir.write, showWarnings = F)
  
  system.time(tmp.curve.rand <- MCGCAttackCurveFunctionalGenesParallel(elTRN, elPPI, ncol(adj.TRN), rowSums(adj.TRN), 
                                                                       colSums(adj.PPI), nboot, nthreads, dir.write, 
                                                                       seed, 1, dir.id, 
                                                                       length(id.disease), id.disease - 1))
}


for( j in c(1:length(Pathogens)) ){
  
  ##Print name of the organism.
  message(paste("Pathogen = ", Pathogens[j], sep = ""))
  
  ##Get organism TRN and PPI networks from multiplex network object.
  adj.TRN <- NetworksMultiplex$HINT[[i]]$TRN
  adj.PPI <- NetworksMultiplex$HINT[[i]]$PPI
  id.inter <- NetworksMultiplex$HINT[[i]]$genes
  
  id.match <- match(toupper(id.inter), toupper(Genes.HPI[[j]]))
  id.disease <- which(!is.na(id.match))
  id.non.disease <- setdiff(1:length(id.inter), id.disease)
  # dir.id <- paste(getwd(), "/", Organisms_file[i], "_HINT_", sub(" ", "", paste(Diseases[j], sep = "_")), 
  #                 "_random_ids", sep = "")
  dir.id <- paste(getwd(), "/", Organisms_file[i], "_HINT_", sub("\\.", "", 
                                                                 sub(" ", "", paste(Pathogens[j], sep = "_"))), 
                  "_random_ids", sep = "")
  dir.create(dir.id, showWarnings = F)
  for( b in 1:nboot ){
    # cat("boot = ", b, "\n")
    id <- sample(c(id.disease, id.non.disease), length(id.disease)) - 1
    file <- paste(dir.id, "/boot_", b, sep = "")
    write.table(cbind(id), file, sep = "\t", row.names = F, 
                col.names = F, quote = F)
  }
  
  ###Attack curves attack on disease genes and proteins
  id1 <- 1:ncol(adj.TRN)
  id2 <- 1:ncol(adj.TRN)
  adj1 <- adj.TRN[id1, id1]
  adj2 <- adj.PPI[id2, id2]
  elTRN <- get.edgelist(graph_from_adjacency_matrix(adj1, "undirected")) - 1
  elPPI <- get.edgelist(graph_from_adjacency_matrix(adj2, "undirected")) - 1
  
  ##Make organism specific folder to store the attack curves
  # dir.write <- paste(getwd(), "/", Organisms_file[i], "_HINT_", sub(" ", "", paste(Diseases[j], sep = "_")), 
  #                    "_random_attack_curve", sep = "")
  dir.write <- paste(getwd(), "/", Organisms_file[i], "_HINT_", sub("\\.", "", 
                                                                    sub(" ", "", paste(Pathogens[j], sep = "_"))), 
                     "_random_attack_curve", sep = "")
  dir.create(dir.write, showWarnings = F)
  
  system.time(tmp.curve.rand <- MCGCAttackCurveFunctionalGenesParallel(elTRN, elPPI, ncol(adj.TRN), rowSums(adj.TRN), 
                                                                       colSums(adj.PPI), nboot, nthreads, dir.write, 
                                                                       seed, 1, dir.id, 
                                                                       length(id.disease), id.disease - 1))
}

##Pathogen BioGRID
i <- 9
for( j in c(1:length(Pathogens)) ){
  
  ##Print name of the organism.
  message(paste("Pathogen = ", Pathogens[j], sep = ""))
  
  ##Get organism TRN and PPI networks from multiplex network object.
  adj.TRN <- NetworksMultiplex$BioGrid[[i]]$TRN
  adj.PPI <- NetworksMultiplex$BioGrid[[i]]$PPI
  id.inter <- NetworksMultiplex$BioGrid[[i]]$genes
  
  id.match <- match(toupper(id.inter), toupper(Genes.HPI[[j]]))
  id.disease <- which(!is.na(id.match))
  id.non.disease <- setdiff(1:length(id.inter), id.disease)
  
  ###Attack curves attack on disease genes and proteins
  id1 <- 1:ncol(adj.TRN)
  id2 <- 1:ncol(adj.TRN)
  adj1 <- adj.TRN[id1, id1]
  adj2 <- adj.PPI[id2, id2]
  elTRN <- get.edgelist(graph_from_adjacency_matrix(adj1, "undirected")) - 1
  elPPI <- get.edgelist(graph_from_adjacency_matrix(adj2, "undirected")) - 1

  ##Make organism specific folder to store the attack curves
  # dir.write <- paste(getwd(), "/", Organisms_file[i], "_BioGrid_", sub(" ", "", paste(Diseases[j], sep = "_")), 
  #                    "_disease_attack_curve", sep = "")
  dir.write <- paste(getwd(), "/", Organisms_file[i], "_BioGrid_", sub("\\.", "", 
                                                                       sub(" ", "", paste(Pathogens[j], sep = "_"))), 
                     "_disease_attack_curve", sep = "")
  dir.create(dir.write, showWarnings = F)
  
  system.time(tmp.curve.rand <- MCGCAttackCurveFunctionalGenesParallel(elTRN, elPPI, ncol(adj.TRN), rowSums(adj.TRN), 
                                                                       colSums(adj.PPI), nboot, nthreads, dir.write, seed, 0, dir.write, 
                                                                       length(id.disease), id.disease - 1))
}


for( j in c(1:length(Pathogens)) ){
  
  ##Print name of the organism.
  message(paste("Pathogen = ", Pathogens[j], sep = ""))
  
  ##Get organism TRN and PPI networks from multiplex network object.
  adj.TRN <- NetworksMultiplex$BioGrid[[i]]$TRN
  adj.PPI <- NetworksMultiplex$BioGrid[[i]]$PPI
  id.inter <- NetworksMultiplex$BioGrid[[i]]$genes
  
  id.match <- match(toupper(id.inter), toupper(Genes.HPI[[j]]))
  id.disease <- which(!is.na(id.match))
  id.non.disease <- setdiff(1:length(id.inter), id.disease)
  # dir.id <- paste(getwd(), "/", Organisms_file[i], "_BioGrid_", sub(" ", "", paste(Diseases[j], sep = "_")), 
  #                 "_nondisease_ids", sep = "")
  dir.id <- paste(getwd(), "/", Organisms_file[i], "_BioGrid_", sub("\\.", "", 
                                                                    sub(" ", "", paste(Pathogens[j], sep = "_"))), 
                  "_nondisease_ids", sep = "")
  dir.create(dir.id, showWarnings = F)
  for( b in 1:nboot ){
    # cat("boot = ", b, "\n")
    id <- sample(id.non.disease, length(id.disease)) - 1
    file <- paste(dir.id, "/boot_", b, sep = "")
    write.table(cbind(id), file, sep = "\t", row.names = F, 
                col.names = F, quote = F)
  }
  
  ###Attack curves attack on disease genes and proteins
  id1 <- 1:ncol(adj.TRN)
  id2 <- 1:ncol(adj.TRN)
  adj1 <- adj.TRN[id1, id1]
  adj2 <- adj.PPI[id2, id2]
  elTRN <- get.edgelist(graph_from_adjacency_matrix(adj1, "undirected")) - 1
  elPPI <- get.edgelist(graph_from_adjacency_matrix(adj2, "undirected")) - 1
  
  ##Make organism specific folder to store the attack curves
  # dir.write <- paste(getwd(), "/", Organisms_file[i], "_BioGrid_", sub(" ", "", paste(Diseases[j], sep = "_")), 
  #                    "_nondisease_attack_curve", sep = "")
  dir.write <- paste(getwd(), "/", Organisms_file[i], "_BioGrid_", sub("\\.", "", 
                                                                       sub(" ", "", paste(Pathogens[j], sep = "_"))), 
                     "_nondisease_attack_curve", sep = "")
  dir.create(dir.write, showWarnings = F)
  
  system.time(tmp.curve.rand <- MCGCAttackCurveFunctionalGenesParallel(elTRN, elPPI, ncol(adj.TRN), rowSums(adj.TRN), 
                                                                       colSums(adj.PPI), nboot, nthreads, dir.write, 
                                                                       seed, 1, dir.id, 
                                                                       length(id.disease), id.disease - 1))
}


for( j in c(1:length(Pathogens)) ){
  
  ##Print name of the organism.
  message(paste("Pathogen = ", Pathogens[j], sep = ""))
  
  ##Get organism TRN and PPI networks from multiplex network object.
  adj.TRN <- NetworksMultiplex$BioGrid[[i]]$TRN
  adj.PPI <- NetworksMultiplex$BioGrid[[i]]$PPI
  id.inter <- NetworksMultiplex$BioGrid[[i]]$genes
  
  id.match <- match(toupper(id.inter), toupper(Genes.HPI[[j]]))
  id.disease <- which(!is.na(id.match))
  id.non.disease <- setdiff(1:length(id.inter), id.disease)
  # dir.id <- paste(getwd(), "/", Organisms_file[i], "_BioGrid_", sub(" ", "", paste(Diseases[j], sep = "_")), 
  #                 "_random_ids", sep = "")
  dir.id <- paste(getwd(), "/", Organisms_file[i], "_BioGrid_", sub("\\.", "", 
                                                                    sub(" ", "", paste(Pathogens[j], sep = "_"))), 
                  "_random_ids", sep = "")
  dir.create(dir.id, showWarnings = F)
  for( b in 1:nboot ){
    # cat("boot = ", b, "\n")
    id <- sample(c(id.disease, id.non.disease), length(id.disease)) - 1
    file <- paste(dir.id, "/boot_", b, sep = "")
    write.table(cbind(id), file, sep = "\t", row.names = F, 
                col.names = F, quote = F)
  }
  
  ###Attack curves attack on disease genes and proteins
  id1 <- 1:ncol(adj.TRN)
  id2 <- 1:ncol(adj.TRN)
  adj1 <- adj.TRN[id1, id1]
  adj2 <- adj.PPI[id2, id2]
  elTRN <- get.edgelist(graph_from_adjacency_matrix(adj1, "undirected")) - 1
  elPPI <- get.edgelist(graph_from_adjacency_matrix(adj2, "undirected")) - 1

  ##Make organism specific folder to store the attack curves
  # dir.write <- paste(getwd(), "/", Organisms_file[i], "_BioGrid_", sub(" ", "", paste(Diseases[j], sep = "_")), 
  #                    "_random_attack_curve", sep = "")
  dir.write <- paste(getwd(), "/", Organisms_file[i], "_BioGrid_", sub("\\.", "", 
                                                                       sub(" ", "", paste(Pathogens[j], sep = "_"))), 
                     "_random_attack_curve", sep = "")
  dir.create(dir.write, showWarnings = F)
  
  system.time(tmp.curve.rand <- MCGCAttackCurveFunctionalGenesParallel(elTRN, elPPI, ncol(adj.TRN), rowSums(adj.TRN), 
                                                                       colSums(adj.PPI), nboot, nthreads, dir.write, 
                                                                       seed, 1, dir.id, 
                                                                       length(id.disease), id.disease - 1))
}


##Cancer
##Generate Data for Figure 5D
dir.results <- getwd()
file <- paste(dir.results, "/NCG6_tsgoncogene.tsv", sep = "")
cancer.genes <- read.table(file, sep = "\t", header = T, 
                           blank.lines.skip = F, quote = "")

##Cancer HINT
for( i in c(9) ){
  
  ##Print name of the organism.
  message(paste("Organism = ", Organisms[i], sep = ""))
  
  ##Get organism TRN and PPI networks from multiplex network object.
  adj.TRN <- NetworksMultiplex$HINT[[i]]$TRN
  adj.PPI <- NetworksMultiplex$HINT[[i]]$PPI
  Genes <- id.inter <- NetworksMultiplex$HINT[[i]]$genes
  
  id.cancer <- match(id.inter, toupper(as.character(cancer.genes[, 2])))
  id.cancer.genes <- which(!is.na(id.cancer))
  id.non.cancer.genes <- which(is.na(id.cancer))
  
  id.cancer <- match(Genes, toupper(as.character(cancer.genes[, 2])))
  cancer.genes.new <- cancer.genes[id.cancer[!is.na(id.cancer)], ]
  id <- which(cancer.genes.new[, 5] == 1)
  id.onco <- match(Genes, toupper(as.character(cancer.genes.new[id, 2])))
  id <- which(cancer.genes.new[, 6] == 1)
  id.suppressor <- match(Genes, toupper(as.character(cancer.genes.new[id, 2])))
  id.onco.genes <- which(!is.na(id.onco))
  id.suppressor.genes <- which(!is.na(id.suppressor))
  
  ##Onco
  id1 <- 1:ncol(adj.TRN)
  id2 <- 1:ncol(adj.TRN)
  adj1 <- adj.TRN[id1, id1]
  adj2 <- adj.PPI[id2, id2]
  elTRN <- get.edgelist(graph_from_adjacency_matrix(adj1, "undirected")) - 1
  elPPI <- get.edgelist(graph_from_adjacency_matrix(adj2, "undirected")) - 1
  
  ##Make organism specific folder to store the attack curves
  dir.write <- paste(getwd(), "/", Organisms_file[i], "_HINT_", 
                     "onco_attack_curve", sep = "")
  dir.create(dir.write, showWarnings = F)
  
  system.time(tmp.curve.rand <- MCGCAttackCurveFunctionalGenesParallel(elTRN, elPPI, ncol(adj.TRN), rowSums(adj.TRN), 
                                                                       colSums(adj.PPI), nboot, nthreads, dir.write, seed, 0, dir.write, 
                                                                       length(id.onco.genes), id.onco.genes - 1))
  
  ##TSG
  id1 <- 1:ncol(adj.TRN)
  id2 <- 1:ncol(adj.TRN)
  adj1 <- adj.TRN[id1, id1]
  adj2 <- adj.PPI[id2, id2]
  elTRN <- get.edgelist(graph_from_adjacency_matrix(adj1, "undirected")) - 1
  elPPI <- get.edgelist(graph_from_adjacency_matrix(adj2, "undirected")) - 1
  
  ##Make organism specific folder to store the attack curves
  dir.write <- paste(getwd(), "/", Organisms_file[i], "_HINT_", 
                     "TSG_attack_curve", sep = "")
  dir.create(dir.write, showWarnings = F)
  
  system.time(tmp.curve.rand <- MCGCAttackCurveFunctionalGenesParallel(elTRN, elPPI, ncol(adj.TRN), rowSums(adj.TRN), 
                                                                       colSums(adj.PPI), nboot, nthreads, dir.write, seed, 0, dir.write, 
                                                                       length(id.suppressor.genes), id.suppressor.genes - 1))
}


for( i in c(9) ){
  
  ##Print name of the organism.
  message(paste("Organism = ", Organisms[i], sep = ""))
  
  ##Get organism TRN and PPI networks from multiplex network object.
  adj.TRN <- NetworksMultiplex$HINT[[i]]$TRN
  adj.PPI <- NetworksMultiplex$HINT[[i]]$PPI
  Genes <- id.inter <- NetworksMultiplex$HINT[[i]]$genes
  
  id.cancer <- match(id.inter, toupper(as.character(cancer.genes[, 2])))
  id.cancer.genes <- which(!is.na(id.cancer))
  id.non.cancer.genes <- which(is.na(id.cancer))
  
  id.cancer <- match(Genes, toupper(as.character(cancer.genes[, 2])))
  cancer.genes.new <- cancer.genes[id.cancer[!is.na(id.cancer)], ]
  id <- which(cancer.genes.new[, 5] == 1)
  id.onco <- match(Genes, toupper(as.character(cancer.genes.new[id, 2])))
  id <- which(cancer.genes.new[, 6] == 1)
  id.suppressor <- match(Genes, toupper(as.character(cancer.genes.new[id, 2])))
  id.onco.genes <- which(!is.na(id.onco))
  id.suppressor.genes <- which(!is.na(id.suppressor))
  
  dir.id <- paste(getwd(), "/", Organisms_file[i], "_HINT_", 
                  "nononco_ids", sep = "")
  dir.create(dir.id, showWarnings = F)
  for( b in 1:nboot ){
    # cat("boot = ", b, "\n")
    id <- sample(id.non.cancer.genes, length(id.onco.genes)) - 1
    file <- paste(dir.id, "/boot_", b, sep = "")
    write.table(cbind(id), file, sep = "\t", row.names = F, 
                col.names = F, quote = F)
  }
  
  
  ###Onco
  id1 <- 1:ncol(adj.TRN)
  id2 <- 1:ncol(adj.TRN)
  adj1 <- adj.TRN[id1, id1]
  adj2 <- adj.PPI[id2, id2]
  elTRN <- get.edgelist(graph_from_adjacency_matrix(adj1, "undirected")) - 1
  elPPI <- get.edgelist(graph_from_adjacency_matrix(adj2, "undirected")) - 1
  
  ##Make organism specific folder to store the attack curves
  dir.write <- paste(getwd(), "/", Organisms_file[i], "_HINT_", 
                     "nononco_attack_curve", sep = "")
  dir.create(dir.write, showWarnings = F)
  
  system.time(tmp.curve.rand <- MCGCAttackCurveFunctionalGenesParallel(elTRN, elPPI, ncol(adj.TRN), rowSums(adj.TRN), 
                                                                       colSums(adj.PPI), nboot, nthreads, dir.write, 
                                                                       seed, 1, dir.id, 
                                                                       length(id.onco.genes), id.onco.genes - 1))
  
  
  dir.id <- paste(getwd(), "/", Organisms_file[i], "_HINT_", 
                  "nonTSG_ids", sep = "")
  dir.create(dir.id, showWarnings = F)
  for( b in 1:nboot ){
    # cat("boot = ", b, "\n")
    id <- sample(id.non.cancer.genes, length(id.suppressor.genes)) - 1
    file <- paste(dir.id, "/boot_", b, sep = "")
    write.table(cbind(id), file, sep = "\t", row.names = F, 
                col.names = F, quote = F)
  }
  
  
  ###TSG
  id1 <- 1:ncol(adj.TRN)
  id2 <- 1:ncol(adj.TRN)
  adj1 <- adj.TRN[id1, id1]
  adj2 <- adj.PPI[id2, id2]
  elTRN <- get.edgelist(graph_from_adjacency_matrix(adj1, "undirected")) - 1
  elPPI <- get.edgelist(graph_from_adjacency_matrix(adj2, "undirected")) - 1

  ##Make organism specific folder to store the attack curves
  dir.write <- paste(getwd(), "/", Organisms_file[i], "_HINT_", 
                     "nonTSG_attack_curve", sep = "")
  dir.create(dir.write, showWarnings = F)
  
  system.time(tmp.curve.rand <- MCGCAttackCurveFunctionalGenesParallel(elTRN, elPPI, ncol(adj.TRN), rowSums(adj.TRN), 
                                                                       colSums(adj.PPI), nboot, nthreads, dir.write, 
                                                                       seed, 1, dir.id, 
                                                                       length(id.suppressor.genes), id.suppressor.genes - 1))
}


for( i in c(9) ){
  
  ##Print name of the organism.
  message(paste("Organism = ", Organisms[i], sep = ""))
  
  ##Get organism TRN and PPI networks from multiplex network object.
  adj.TRN <- NetworksMultiplex$HINT[[i]]$TRN
  adj.PPI <- NetworksMultiplex$HINT[[i]]$PPI
  Genes <- id.inter <- NetworksMultiplex$HINT[[i]]$genes
  
  id.cancer <- match(id.inter, toupper(as.character(cancer.genes[, 2])))
  id.cancer.genes <- which(!is.na(id.cancer))
  id.non.cancer.genes <- which(is.na(id.cancer))
  
  id.cancer <- match(Genes, toupper(as.character(cancer.genes[, 2])))
  cancer.genes.new <- cancer.genes[id.cancer[!is.na(id.cancer)], ]
  id <- which(cancer.genes.new[, 5] == 1)
  id.onco <- match(Genes, toupper(as.character(cancer.genes.new[id, 2])))
  id <- which(cancer.genes.new[, 6] == 1)
  id.suppressor <- match(Genes, toupper(as.character(cancer.genes.new[id, 2])))
  id.onco.genes <- which(!is.na(id.onco))
  id.suppressor.genes <- which(!is.na(id.suppressor))
  
  dir.id <- paste(getwd(), "/", Organisms_file[i], "_HINT_",
                  "randomonco_ids", sep = "")
  dir.create(dir.id, showWarnings = F)
  for( b in 1:nboot ){
    # cat("boot = ", b, "\n")
    id <- sample(c(id.onco.genes, id.non.cancer.genes), length(id.onco.genes)) - 1
    file <- paste(dir.id, "/boot_", b, sep = "")
    write.table(cbind(id), file, sep = "\t", row.names = F, 
                col.names = F, quote = F)
  }
  
  
  
  ###Onco
  id1 <- 1:ncol(adj.TRN)
  id2 <- 1:ncol(adj.TRN)
  adj1 <- adj.TRN[id1, id1]
  adj2 <- adj.PPI[id2, id2]
  elTRN <- get.edgelist(graph_from_adjacency_matrix(adj1, "undirected")) - 1
  elPPI <- get.edgelist(graph_from_adjacency_matrix(adj2, "undirected")) - 1
  
  ##Make organism specific folder to store the attack curves
  dir.write <- paste(getwd(), "/", Organisms_file[i], "_HINT_",
                     "randomonco_attack_curve", sep = "")
  dir.create(dir.write, showWarnings = F)
  
  system.time(tmp.curve.rand <- MCGCAttackCurveFunctionalGenesParallel(elTRN, elPPI, ncol(adj.TRN), rowSums(adj.TRN), 
                                                                       colSums(adj.PPI), nboot, nthreads, dir.write, 
                                                                       seed, 1, dir.id, 
                                                                       length(id.onco.genes), id.onco.genes - 1))
  
  
  dir.id <- paste(getwd(), "/", Organisms_file[i], "_HINT_",
                  "randomTSG_ids", sep = "")
  dir.create(dir.id, showWarnings = F)
  for( b in 1:nboot ){
    # cat("boot = ", b, "\n")
    id <- sample(c(id.suppressor.genes, id.non.cancer.genes), length(id.suppressor.genes)) - 1
    file <- paste(dir.id, "/boot_", b, sep = "")
    write.table(cbind(id), file, sep = "\t", row.names = F, 
                col.names = F, quote = F)
  }
  
  ###TSG
  id1 <- 1:ncol(adj.TRN)
  id2 <- 1:ncol(adj.TRN)
  adj1 <- adj.TRN[id1, id1]
  adj2 <- adj.PPI[id2, id2]
  elTRN <- get.edgelist(graph_from_adjacency_matrix(adj1, "undirected")) - 1
  elPPI <- get.edgelist(graph_from_adjacency_matrix(adj2, "undirected")) - 1
  
  ##Make organism specific folder to store the attack curves
  dir.write <- paste(getwd(), "/", Organisms_file[i], "_HINT_",
                     "randomTSG_attack_curve", sep = "")
  dir.create(dir.write, showWarnings = F)
  
  system.time(tmp.curve.rand <- MCGCAttackCurveFunctionalGenesParallel(elTRN, elPPI, ncol(adj.TRN), rowSums(adj.TRN), 
                                                                       colSums(adj.PPI), nboot, nthreads, dir.write, 
                                                                       seed, 1, dir.id, 
                                                                       length(id.suppressor.genes), id.suppressor.genes - 1))
}


##Cancer BioGRID
for( i in c(9) ){
  
  ##Print name of the organism.
  message(paste("Organism = ", Organisms[i], sep = ""))
  
  ##Get organism TRN and PPI networks from multiplex network object.
  adj.TRN <- NetworksMultiplex$BioGrid[[i]]$TRN
  adj.PPI <- NetworksMultiplex$BioGrid[[i]]$PPI
  Genes <- id.inter <- NetworksMultiplex$BioGrid[[i]]$genes
  
  id.cancer <- match(id.inter, toupper(as.character(cancer.genes[, 2])))
  id.cancer.genes <- which(!is.na(id.cancer))
  id.non.cancer.genes <- which(is.na(id.cancer))
  
  id.cancer <- match(Genes, toupper(as.character(cancer.genes[, 2])))
  cancer.genes.new <- cancer.genes[id.cancer[!is.na(id.cancer)], ]
  id <- which(cancer.genes.new[, 5] == 1)
  id.onco <- match(Genes, toupper(as.character(cancer.genes.new[id, 2])))
  id <- which(cancer.genes.new[, 6] == 1)
  id.suppressor <- match(Genes, toupper(as.character(cancer.genes.new[id, 2])))
  id.onco.genes <- which(!is.na(id.onco))
  id.suppressor.genes <- which(!is.na(id.suppressor))
  
  ###Onco
  id1 <- 1:ncol(adj.TRN)
  id2 <- 1:ncol(adj.TRN)
  adj1 <- adj.TRN[id1, id1]
  adj2 <- adj.PPI[id2, id2]
  elTRN <- get.edgelist(graph_from_adjacency_matrix(adj1, "undirected")) - 1
  elPPI <- get.edgelist(graph_from_adjacency_matrix(adj2, "undirected")) - 1
  
  ##Make organism specific folder to store the attack curves
  dir.write <- paste(getwd(), "/", Organisms_file[i], "_BioGrid_", 
                     "onco_attack_curve", sep = "")
  dir.create(dir.write, showWarnings = F)
  
  system.time(tmp.curve.rand <- MCGCAttackCurveFunctionalGenesParallel(elTRN, elPPI, ncol(adj.TRN), rowSums(adj.TRN), 
                                                                       colSums(adj.PPI), nboot, nthreads, dir.write, seed, 0, dir.write, 
                                                                       length(id.onco.genes), id.onco.genes - 1))
  
  dir.write <- paste(getwd(), "/", Organisms_file[i], "_BioGrid_", 
                     "TSG_attack_curve", sep = "")
  dir.create(dir.write, showWarnings = F)
  
  system.time(tmp.curve.rand <- MCGCAttackCurveFunctionalGenesParallel(elTRN, elPPI, ncol(adj.TRN), rowSums(adj.TRN), 
                                                                       colSums(adj.PPI), nboot, nthreads, dir.write, seed, 0, dir.write, 
                                                                       length(id.suppressor.genes), id.suppressor.genes - 1))
}


for( i in c(9) ){
  
  ##Print name of the organism.
  message(paste("Organism = ", Organisms[i], sep = ""))
  
  ##Get organism TRN and PPI networks from multiplex network object.
  adj.TRN <- NetworksMultiplex$BioGrid[[i]]$TRN
  adj.PPI <- NetworksMultiplex$BioGrid[[i]]$PPI
  Genes <- id.inter <- NetworksMultiplex$BioGrid[[i]]$genes
  
  id.cancer <- match(id.inter, toupper(as.character(cancer.genes[, 2])))
  id.cancer.genes <- which(!is.na(id.cancer))
  id.non.cancer.genes <- which(is.na(id.cancer))
  
  id.cancer <- match(Genes, toupper(as.character(cancer.genes[, 2])))
  cancer.genes.new <- cancer.genes[id.cancer[!is.na(id.cancer)], ]
  id <- which(cancer.genes.new[, 5] == 1)
  id.onco <- match(Genes, toupper(as.character(cancer.genes.new[id, 2])))
  id <- which(cancer.genes.new[, 6] == 1)
  id.suppressor <- match(Genes, toupper(as.character(cancer.genes.new[id, 2])))
  id.onco.genes <- which(!is.na(id.onco))
  id.suppressor.genes <- which(!is.na(id.suppressor))
  
  dir.id <- paste(getwd(), "/", Organisms_file[i], "_BioGrid_",
                  "nononco_ids", sep = "")
  dir.create(dir.id, showWarnings = F)
  for( b in 1:nboot ){
    # cat("boot = ", b, "\n")
    id <- sample(id.non.cancer.genes, length(id.onco.genes)) - 1
    file <- paste(dir.id, "/boot_", b, sep = "")
    write.table(cbind(id), file, sep = "\t", row.names = F, 
                col.names = F, quote = F)
  }
  
  ###Attack curves attack on disease genes and proteins
  id1 <- 1:ncol(adj.TRN)
  id2 <- 1:ncol(adj.TRN)
  adj1 <- adj.TRN[id1, id1]
  adj2 <- adj.PPI[id2, id2]
  elTRN <- get.edgelist(graph_from_adjacency_matrix(adj1, "undirected")) - 1
  elPPI <- get.edgelist(graph_from_adjacency_matrix(adj2, "undirected")) - 1

  ##Make organism specific folder to store the attack curves
  dir.write <- paste(getwd(), "/", Organisms_file[i], "_BioGrid_", 
                     "nononco_attack_curve", sep = "")
  dir.create(dir.write, showWarnings = F)
  
  system.time(tmp.curve.rand <- MCGCAttackCurveFunctionalGenesParallel(elTRN, elPPI, ncol(adj.TRN), rowSums(adj.TRN), 
                                                                       colSums(adj.PPI), nboot, nthreads, dir.write, 
                                                                       seed, 1, dir.id, 
                                                                       length(id.onco.genes), id.onco.genes - 1))
  
  
  ##TSG
  dir.id <- paste(getwd(), "/", Organisms_file[i], "_BioGrid_",
                  "nonTSG_ids", sep = "")
  dir.create(dir.id, showWarnings = F)
  for( b in 1:nboot ){
    # cat("boot = ", b, "\n")
    id <- sample(id.non.cancer.genes, length(id.suppressor.genes)) - 1
    file <- paste(dir.id, "/boot_", b, sep = "")
    write.table(cbind(id), file, sep = "\t", row.names = F, 
                col.names = F, quote = F)
  }
  
  ###Attack curves attack on disease genes and proteins
  id1 <- 1:ncol(adj.TRN)
  id2 <- 1:ncol(adj.TRN)
  adj1 <- adj.TRN[id1, id1]
  adj2 <- adj.PPI[id2, id2]
  elTRN <- get.edgelist(graph_from_adjacency_matrix(adj1, "undirected")) - 1
  elPPI <- get.edgelist(graph_from_adjacency_matrix(adj2, "undirected")) - 1

  ##Make organism specific folder to store the attack curves
  dir.write <- paste(getwd(), "/", Organisms_file[i], "_BioGrid_", 
                     "nonTSG_attack_curve", sep = "")
  dir.create(dir.write, showWarnings = F)
  
  system.time(tmp.curve.rand <- MCGCAttackCurveFunctionalGenesParallel(elTRN, elPPI, ncol(adj.TRN), rowSums(adj.TRN), 
                                                                       colSums(adj.PPI), nboot, nthreads, dir.write, 
                                                                       seed, 1, dir.id, 
                                                                       length(id.suppressor.genes), id.suppressor.genes - 1))
}


for( i in c(9) ){
  
  ##Print name of the organism.
  message(paste("Organism = ", Organisms[i], sep = ""))
  
  ##Get organism TRN and PPI networks from multiplex network object.
  adj.TRN <- NetworksMultiplex$BioGrid[[i]]$TRN
  adj.PPI <- NetworksMultiplex$BioGrid[[i]]$PPI
  Genes <- id.inter <- NetworksMultiplex$BioGrid[[i]]$genes
  
  id.cancer <- match(id.inter, toupper(as.character(cancer.genes[, 2])))
  id.cancer.genes <- which(!is.na(id.cancer))
  id.non.cancer.genes <- which(is.na(id.cancer))
  
  id.cancer <- match(Genes, toupper(as.character(cancer.genes[, 2])))
  cancer.genes.new <- cancer.genes[id.cancer[!is.na(id.cancer)], ]
  id <- which(cancer.genes.new[, 5] == 1)
  id.onco <- match(Genes, toupper(as.character(cancer.genes.new[id, 2])))
  id <- which(cancer.genes.new[, 6] == 1)
  id.suppressor <- match(Genes, toupper(as.character(cancer.genes.new[id, 2])))
  id.onco.genes <- which(!is.na(id.onco))
  id.suppressor.genes <- which(!is.na(id.suppressor))
  
  dir.id <- paste(getwd(), "/", Organisms_file[i], "_BioGrid_",
                  "randomonco_ids", sep = "")
  dir.create(dir.id, showWarnings = F)
  for( b in 1:nboot ){
    # cat("boot = ", b, "\n")
    id <- sample(c(id.onco.genes, id.non.cancer.genes), length(id.onco.genes)) - 1
    file <- paste(dir.id, "/boot_", b, sep = "")
    write.table(cbind(id), file, sep = "\t", row.names = F, 
                col.names = F, quote = F)
  }
  
  ###Attack curves attack on disease genes and proteins
  id1 <- 1:ncol(adj.TRN)
  id2 <- 1:ncol(adj.TRN)
  adj1 <- adj.TRN[id1, id1]
  adj2 <- adj.PPI[id2, id2]
  elTRN <- get.edgelist(graph_from_adjacency_matrix(adj1, "undirected")) - 1
  elPPI <- get.edgelist(graph_from_adjacency_matrix(adj2, "undirected")) - 1

  ##Make organism specific folder to store the attack curves
  dir.write <- paste(getwd(), "/", Organisms_file[i], "_BioGrid_",
                     "randomonco_attack_curve", sep = "")
  dir.create(dir.write, showWarnings = F)
  
  system.time(tmp.curve.rand <- MCGCAttackCurveFunctionalGenesParallel(elTRN, elPPI, ncol(adj.TRN), rowSums(adj.TRN), 
                                                                       colSums(adj.PPI), nboot, nthreads, dir.write, 
                                                                       seed, 1, dir.id, 
                                                                       length(id.onco.genes), id.onco.genes - 1))
  
  ##TSG
  dir.id <- paste(getwd(), "/", Organisms_file[i], "_BioGrid_",
                  "randomTSG_ids", sep = "")
  dir.create(dir.id, showWarnings = F)
  for( b in 1:nboot ){
    # cat("boot = ", b, "\n")
    id <- sample(c(id.suppressor.genes, id.non.cancer.genes), length(id.suppressor.genes)) - 1
    file <- paste(dir.id, "/boot_", b, sep = "")
    write.table(cbind(id), file, sep = "\t", row.names = F, 
                col.names = F, quote = F)
  }
  
  ###Attack curves attack on disease genes and proteins
  id1 <- 1:ncol(adj.TRN)
  id2 <- 1:ncol(adj.TRN)
  adj1 <- adj.TRN[id1, id1]
  adj2 <- adj.PPI[id2, id2]
  elTRN <- get.edgelist(graph_from_adjacency_matrix(adj1, "undirected")) - 1
  elPPI <- get.edgelist(graph_from_adjacency_matrix(adj2, "undirected")) - 1
  
  ##Make organism specific folder to store the attack curves
  dir.write <- paste(getwd(), "/", Organisms_file[i], "_BioGrid_",
                     "randomTSG_attack_curve", sep = "")
  dir.create(dir.write, showWarnings = F)
  
  system.time(tmp.curve.rand <- MCGCAttackCurveFunctionalGenesParallel(elTRN, elPPI, ncol(adj.TRN), rowSums(adj.TRN), 
                                                                       colSums(adj.PPI), nboot, nthreads, dir.write, 
                                                                       seed, 1, dir.id, 
                                                                       length(id.suppressor.genes), id.suppressor.genes - 1))
}

##Object for Figure 5 plot
Plot.5 <- list()

##Plot Figure 5A

for( i in c(4, 6, 9) ){
  
  ##Print name of the organism.
  message(paste("Organism = ", Organisms[i], sep = ""))
  
  ##Get organism TRN and PPI networks from multiplex network object.
  adj.TRN <- NetworksMultiplex$HINT[[i]]$TRN
  adj.PPI <- NetworksMultiplex$HINT[[i]]$PPI
  id.inter <- NetworksMultiplex$HINT[[i]]$genes
  
  genes.lethal <- toupper(Lethality.HINT$genes[[i]])[Lethality.HINT$gene.lethality[[i]] == 0]
  id.match <- match(toupper(id.inter), toupper(genes.lethal))
  id.lethal <- which(!is.na(id.match))
  
  genes.nonlethal <- toupper(Lethality.HINT$genes[[i]])[Lethality.HINT$gene.lethality[[i]] != 0]
  id.match <- match(toupper(id.inter), toupper(genes.nonlethal))
  id.nonlethal <- which(!is.na(id.match))
  
  dir.write <- paste(getwd(), "/", Organisms_file[i], "_HINT_", 
                     "Lethal_attack_curve", sep = "")
  tmp.ess <- rep(0, nboot)
  for( b in 1:nboot ){
    tmp.MCGC <- c(unlist(read.table(paste(dir.write, "/boot_", b, sep = ""), sep = "\t", header = F)))
    id <- which(tmp.MCGC <= 1)[1]
    tmp.ess[b] <- abs(sintegral((1:length(tmp.MCGC)/ncol(adj.TRN)), tmp.MCGC/ncol(adj.TRN))$value)
  }
  edge.ess <- count_common_edges_among_nodes_complete_iterator(adj.TRN, adj.PPI, id.lethal - 1)
  
  dir.write <- paste(getwd(), "/", Organisms_file[i], "_HINT_", 
                     "randomLethal_attack_curve", sep = "")
  tmp.rand <- rep(0, nboot)
  for( b in 1:nboot ){
    tmp.MCGC <- c(unlist(read.table(paste(dir.write, "/boot_", b, sep = ""), sep = "\t", header = F)))
    id <- which(tmp.MCGC <= 1)[1]
    tmp.rand[b] <- abs(sintegral((1:length(tmp.MCGC)/ncol(adj.TRN)), tmp.MCGC/ncol(adj.TRN))$value)
  }
  dir.write <- paste(getwd(), "/", Organisms_file[i], "_HINT_", 
                     "randomLethal_ids", sep = "")
  edge.rand <- cor.rand <- rep(0, nboot)
  for( b in 1:nboot ){
    id <- c(unlist(read.table(paste(dir.write, "/boot_", b, sep = ""), sep = "\t", header = F)))
    edge.rand[b] <- count_common_edges_among_nodes_complete_iterator(adj.TRN, adj.PPI, id)
    cor.rand[b] <- cor(rowSums(adj.TRN)[id + 1], colSums(adj.PPI)[id + 1], method = "spearman")
  }
  
  dir.write <- paste(getwd(), "/", Organisms_file[i], "_HINT_", 
                     "nonLethal_attack_curve", sep = "")
  tmp.noness <- rep(0, nboot)
  for( b in 1:nboot ){
    tmp.MCGC <- c(unlist(read.table(paste(dir.write, "/boot_", b, sep = ""), sep = "\t", header = F)))
    id <- which(tmp.MCGC <= 1)[1]
    tmp.noness[b] <- abs(sintegral((1:length(tmp.MCGC)/ncol(adj.TRN)), tmp.MCGC/ncol(adj.TRN))$value)
  }
  dir.write <- paste(getwd(), "/", Organisms_file[i], "_HINT_", 
                     "nonLethal_ids", sep = "")
  edge.noness <- cor.noness <- rep(0, nboot)
  for( b in 1:nboot ){
    id <- c(unlist(read.table(paste(dir.write, "/boot_", b, sep = ""), sep = "\t", header = F)))
    edge.noness[b] <- count_common_edges_among_nodes_complete_iterator(adj.TRN, adj.PPI, id)
    cor.noness[b] <- cor(rowSums(adj.TRN)[id + 1], colSums(adj.PPI)[id + 1], method = "spearman")
  }
  
  tmp <- orddom(tmp.rand, tmp.ess)
  R.ess <- as.numeric(tmp[c(23:25), 2])
  
  tmp <- orddom(tmp.rand, tmp.noness)
  R.noness <- as.numeric(tmp[c(23:25), 2])
  
  CR.ess <- c(edge.ess - mean(edge.rand), 
              edge.ess - mean(edge.rand) - 2*sd(edge.rand)/10, 
              edge.ess - mean(edge.rand) + 2*sd(edge.rand)/10)
  CD.ess <- SpearmanRho(rowSums(adj.TRN)[id.lethal], colSums(adj.PPI)[id.lethal], conf.level = 0.95)
  
  CR.noness <- c(mean(edge.noness) - mean(edge.rand), 
                 mean(edge.noness) - mean(edge.rand) - 2*sqrt(var(edge.rand) + var(edge.noness))/10, 
                 mean(edge.noness) - mean(edge.rand) + 2*sqrt(var(edge.rand) + var(edge.noness))/10)
  CD.noness <- mean(cor.noness, na.rm = T) + c(0, - 2*sd(cor.noness, na.rm = T)/10, + 2*sd(cor.noness, na.rm = T)/10)
  
  if( i == 4 ){
    data.plot <- data.frame(Org = rep(paste(Organisms[i], " HINT", sep = ""), 2), 
                            CR = c(CR.ess[1], CR.noness[1]),
                            CR.lower = c(CR.ess[2], CR.noness[2]),
                            CR.upper = c(CR.ess[3], CR.noness[3]),
                            CD = c(CD.ess[1], CD.noness[1]),
                            CD.lower = c(CD.ess[2], CD.noness[2]),
                            CD.upper = c(CD.ess[3], CD.noness[3]),
                            R = c(R.ess[1], R.noness[1]), 
                            R.lower = c(R.ess[2], R.noness[2]), 
                            R.upper = c(R.ess[3], R.noness[3]), 
                            Type = c("Ess.", "non Ess."), 
                            size = c(length(id.lethal), length(id.nonlethal)))
  }else{
    data.tmp <- data.frame(Org = rep(paste(Organisms[i], " HINT", sep = ""), 2), 
                           CR = c(CR.ess[1], CR.noness[1]),
                           CR.lower = c(CR.ess[2], CR.noness[2]),
                           CR.upper = c(CR.ess[3], CR.noness[3]),
                           CD = c(CD.ess[1], CD.noness[1]),
                           CD.lower = c(CD.ess[2], CD.noness[2]),
                           CD.upper = c(CD.ess[3], CD.noness[3]),
                           R = c(R.ess[1], R.noness[1]), 
                           R.lower = c(R.ess[2], R.noness[2]), 
                           R.upper = c(R.ess[3], R.noness[3]), 
                           Type = c("Ess.", "non Ess."), size = c(length(id.lethal), length(id.nonlethal)))
    data.plot <- rbind(data.plot, data.tmp)
  }
  
  ##BioGRID
  adj.TRN <- NetworksMultiplex$BioGrid[[i]]$TRN
  adj.PPI <- NetworksMultiplex$BioGrid[[i]]$PPI
  id.inter <- NetworksMultiplex$BioGrid[[i]]$genes
  
  genes.lethal <- toupper(Lethality.BioGRID$genes[[i]])[Lethality.BioGRID$gene.lethality[[i]] == 0]
  id.match <- match(toupper(id.inter), toupper(genes.lethal))
  id.lethal <- which(!is.na(id.match))
  
  genes.nonlethal <- toupper(Lethality.BioGRID$genes[[i]])[Lethality.BioGRID$gene.lethality[[i]] != 0]
  id.match <- match(toupper(id.inter), toupper(genes.nonlethal))
  id.nonlethal <- which(!is.na(id.match))
  
  dir.write <- paste(getwd(), "/", Organisms_file[i], "_BioGrid_", 
                     "Lethal_attack_curve", sep = "")
  tmp.ess <- rep(0, nboot)
  for( b in 1:nboot ){
    tmp.MCGC <- c(unlist(read.table(paste(dir.write, "/boot_", b, sep = ""), sep = "\t", header = F)))
    id <- which(tmp.MCGC <= 1)[1]
    tmp.ess[b] <- abs(sintegral((1:length(tmp.MCGC)/ncol(adj.TRN)), tmp.MCGC/ncol(adj.TRN))$value)
  }
  edge.ess <- count_common_edges_among_nodes_complete_iterator(adj.TRN, adj.PPI, id.lethal - 1)
  
  dir.write <- paste(getwd(), "/", Organisms_file[i], "_BioGrid_", 
                     "randomLethal_attack_curve", sep = "")
  tmp.rand <- rep(0, nboot)
  for( b in 1:nboot ){
    tmp.MCGC <- c(unlist(read.table(paste(dir.write, "/boot_", b, sep = ""), sep = "\t", header = F)))
    id <- which(tmp.MCGC <= 1)[1]
    tmp.rand[b] <- abs(sintegral((1:length(tmp.MCGC)/ncol(adj.TRN)), tmp.MCGC/ncol(adj.TRN))$value)
  }
  dir.write <- paste(getwd(), "/", Organisms_file[i], "_BioGrid_", 
                     "randomLethal_ids", sep = "")
  edge.rand <- rep(0, nboot)
  for( b in 1:nboot ){
    id <- c(unlist(read.table(paste(dir.write, "/boot_", b, sep = ""), sep = "\t", header = F)))
    edge.rand[b] <- count_common_edges_among_nodes_complete_iterator(adj.TRN, adj.PPI, id)
  }
  
  dir.write <- paste(getwd(), "/", Organisms_file[i], "_BioGrid_", 
                     "nonLethal_attack_curve", sep = "")
  tmp.noness <- rep(0, nboot)
  for( b in 1:nboot ){
    tmp.MCGC <- c(unlist(read.table(paste(dir.write, "/boot_", b, sep = ""), sep = "\t", header = F)))
    id <- which(tmp.MCGC <= 1)[1]
    tmp.noness[b] <- abs(sintegral((1:length(tmp.MCGC)/ncol(adj.TRN)), tmp.MCGC/ncol(adj.TRN))$value)
  }
  dir.write <- paste(getwd(), "/", Organisms_file[i], "_BioGrid_", 
                     "nonLethal_ids", sep = "")
  edge.noness <- cor.noness <- rep(0, nboot)
  for( b in 1:nboot ){
    id <- c(unlist(read.table(paste(dir.write, "/boot_", b, sep = ""), sep = "\t", header = F)))
    edge.noness[b] <- count_common_edges_among_nodes_complete_iterator(adj.TRN, adj.PPI, id)
    cor.noness[b] <- cor(rowSums(adj.TRN)[id + 1], colSums(adj.PPI)[id + 1], method = "spearman")
  }
  
  
  tmp <- orddom(tmp.rand, tmp.ess)
  R.ess <- as.numeric(tmp[c(23:25), 2])
  
  tmp <- orddom(tmp.rand, tmp.noness)
  R.noness <- as.numeric(tmp[c(23:25), 2])
  
  CR.ess <- c(edge.ess - mean(edge.rand), 
              edge.ess - mean(edge.rand) - 2*sd(edge.rand)/10, 
              edge.ess - mean(edge.rand) + 2*sd(edge.rand)/10)
  
  CD.ess <- SpearmanRho(rowSums(adj.TRN)[id.lethal], colSums(adj.PPI)[id.lethal], conf.level = 0.95)
  
  CR.noness <- c(mean(edge.noness) - mean(edge.rand), 
                 mean(edge.noness) - mean(edge.rand) - 2*sqrt(var(edge.rand) + var(edge.noness))/10, 
                 mean(edge.noness) - mean(edge.rand) + 2*sqrt(var(edge.rand) + var(edge.noness))/10)
  CD.noness <- mean(cor.noness, na.rm = T) + c(0, - 2*sd(cor.noness, na.rm = T)/10, + 2*sd(cor.noness, na.rm = T)/10)
  
  data.tmp <- data.frame(Org = rep(paste(Organisms[i], " BioGRID", sep = ""), 2), 
                         CR = c(CR.ess[1], CR.noness[1]),
                         CR.lower = c(CR.ess[2], CR.noness[2]),
                         CR.upper = c(CR.ess[3], CR.noness[3]),
                         CD = c(CD.ess[1], CD.noness[1]),
                         CD.lower = c(CD.ess[2], CD.noness[2]),
                         CD.upper = c(CD.ess[3], CD.noness[3]),
                         R = c(R.ess[1], R.noness[1]), 
                         R.lower = c(R.ess[2], R.noness[2]), 
                         R.upper = c(R.ess[3], R.noness[3]), 
                         Type = c("Ess.", "non Ess."), 
                         size = c(length(id.lethal), length(id.nonlethal)))
  data.plot <- rbind(data.plot, data.tmp)
  
}
data.plot.list <- list()
data.plot.list[[1]] <- data.plot
id <- order(data.plot[data.plot$Type == "Ess.", 8], decreasing = F)
data.plot$Org <- factor(data.plot$Org, levels = as.character(data.plot[data.plot$Type == "Ess.", 1][id]))
p <- ggbarplot(data.plot, x = "Org", y = "R", 
               fill = "Type",           # change fill color by mpg_level
               group = "Type",
               color = "Type",# jco journal color palett. see ?ggpar         # Sort the value in descending order
               palette = c("red", "blue"),
               sort.by.groups = F,     # Don't sort inside each group
               # x.text.angle = 50,          # Rotate vertically x axis texts
               xlab = "",
               sort.val = "none",
               ylab = expression("R"),
               # legend = "bottom",
               legend.title = "",
               y.text.angle = 50,
               rotate = T,
               font.x = 30, 
               font.y = 20,
               font.tickslab = 20,
               size = 0.9,
               point.size = 0.1,
               font.label = 12,
               stat = "identity",
               theme = theme_pubr(), 
               position = position_dodge(0.8),
)
p1 <- p + geom_hline(yintercept = 0)+ geom_errorbar(aes(ymin = R.lower, ymax = R.upper, group = Type), width = 0.2, 
                                                    position = position_dodge(0.8)) + 
  theme(title = element_text(size = 30, color = "black"), 
        plot.title = element_text(size = 20, color = "black"), 
        axis.title = element_text(size = 30, colour = "black"), 
        legend.text = element_text(size = 25, color = "black"))


id <- order(data.plot[data.plot$Type == "Ess.", 8], decreasing = F)
data.plot$Org <- factor(data.plot$Org, levels = as.character(data.plot[data.plot$Type == "Ess.", 1][id]))
p <- ggbarplot(data.plot, x = "Org", y = "CR", 
               fill = "Type",           # change fill color by mpg_level
               group = "Type",
               color = "Type",# jco journal color palett. see ?ggpar         # Sort the value in descending order
               palette = c("red", "blue"),
               sort.by.groups = F,     # Don't sort inside each group
               # x.text.angle = 50,          # Rotate vertically x axis texts
               xlab = "",
               sort.val = "none",
               ylab = expression("C"["R"]),
               legend = "none",
               legend.title = "",
               y.text.angle = 50,
               rotate = T,
               font.x = 20, 
               font.y = 20,
               font.tickslab = 20,
               size = 0.9,
               point.size = 0.1,
               font.label = 12,
               stat = "identity",
               theme = theme_pubr(), 
               position = position_dodge(0.8),
)
p2 <- p + geom_hline(yintercept = 0)+ geom_errorbar(aes(ymin = CR.lower, ymax = CR.upper, group = Type), width = 0.2, 
                                                    position = position_dodge(0.8)) + 
  ggtitle(expression(paste("R: Robustness, ", "C"["R"], ": Redundancy", sep = "")))

Plot.5[[1]] <- plot_grid(p1, 
                         p2 + 
                           theme(axis.text.y = element_blank(),
                                 # axis.line.y = element_blank(),
                                 plot.title = element_text(size = 30, color = "black"),
                                 axis.title.y= element_blank(),
                                 axis.ticks.y= element_blank(), 
                                 axis.title.x = element_text(size = 30, colour = "black")),
                         nrow = 1,
                         rel_widths = c(1.2, 1),
                         align = 'h', axis = 'tb')


##Plot Figure 5B
i <- 9
for( j in 1:length(Pathogens) ){
  
  ##Print name of the organism.
  message(paste("Organism = ", Organisms[i], sep = ""))
  
  ##Get organism TRN and PPI networks from multiplex network object.
  adj.TRN <- NetworksMultiplex$HINT[[i]]$TRN
  adj.PPI <- NetworksMultiplex$HINT[[i]]$PPI
  id.inter <- NetworksMultiplex$HINT[[i]]$genes
  
  id.match <- match(toupper(id.inter), toupper(Genes.HPI[[j]]))
  id.disease <- which(!is.na(id.match))
  id.non.disease <- setdiff(1:length(id.inter), id.disease)
  
  
  ##Make organism specific folder to store the attack curves
  dir.write <- paste(getwd(), "/", Organisms_file[i], "_HINT_", sub("\\.", "", 
                                                                    sub(" ", "", paste(Pathogens[j], sep = "_"))), 
                     "_pathogen_attack_curve", sep = "")
  tmp.path <- rep(0, nboot)
  for( b in 1:nboot ){
    tmp.MCGC <- c(unlist(read.table(paste(dir.write, "/boot_", b, sep = ""), sep = "\t", header = F)))
    id <- which(tmp.MCGC <= 1)[1]
    tmp.path[b] <- abs(sintegral((1:length(tmp.MCGC)/ncol(adj.TRN)), tmp.MCGC/ncol(adj.TRN))$value)
  }
  edge.path <- count_common_edges_among_nodes_complete_iterator(adj.TRN, adj.PPI, id.disease - 1)
  
  dir.write <- paste(getwd(), "/", Organisms_file[i], "_HINT_", sub("\\.", "", 
                                                                    sub(" ", "", paste(Pathogens[j], sep = "_"))), 
                     "_random_attack_curve", sep = "")
  tmp.rand <- rep(0, nboot)
  for( b in 1:nboot ){
    tmp.MCGC <- c(unlist(read.table(paste(dir.write, "/boot_", b, sep = ""), sep = "\t", header = F)))
    id <- which(tmp.MCGC <= 1)[1]
    tmp.rand[b] <- abs(sintegral((1:length(tmp.MCGC)/ncol(adj.TRN)), tmp.MCGC/ncol(adj.TRN))$value)
  }
  dir.write <- paste(getwd(), "/", Organisms_file[i], "_HINT_", sub("\\.", "", 
                                                                    sub(" ", "", paste(Pathogens[j], sep = "_"))), 
                     "_random_ids", sep = "")
  edge.rand <- cor.rand <- rep(0, nboot)
  for( b in 1:nboot ){
    id <- c(unlist(read.table(paste(dir.write, "/boot_", b, sep = ""), sep = "\t", header = F)))
    edge.rand[b] <- count_common_edges_among_nodes_complete_iterator(adj.TRN, adj.PPI, id)
    cor.rand[b] <- cor(rowSums(adj.TRN)[id + 1], colSums(adj.PPI)[id + 1], method = "spearman")
  }
  
  dir.write <- paste(getwd(), "/", Organisms_file[i], "_HINT_", sub("\\.", "", 
                                                                    sub(" ", "", paste(Pathogens[j], sep = "_"))), 
                     "_nonpathogen_attack_curve", sep = "")
  tmp.nonpath <- rep(0, nboot)
  for( b in 1:nboot ){
    tmp.MCGC <- c(unlist(read.table(paste(dir.write, "/boot_", b, sep = ""), sep = "\t", header = F)))
    id <- which(tmp.MCGC <= 1)[1]
    tmp.nonpath[b] <- abs(sintegral((1:length(tmp.MCGC)/ncol(adj.TRN)), tmp.MCGC/ncol(adj.TRN))$value)
  }
  dir.write <- paste(getwd(), "/", Organisms_file[i], "_HINT_", sub("\\.", "", 
                                                                    sub(" ", "", paste(Pathogens[j], sep = "_"))), 
                     "_nonpathogen_ids", sep = "")
  edge.nonpath <- cor.nonpath <- rep(0, nboot)
  for( b in 1:nboot ){
    id <- c(unlist(read.table(paste(dir.write, "/boot_", b, sep = ""), sep = "\t", header = F)))
    edge.nonpath[b] <- count_common_edges_among_nodes_complete_iterator(adj.TRN, adj.PPI, id)
    cor.nonpath[b] <- cor(rowSums(adj.TRN)[id + 1], colSums(adj.PPI)[id + 1], method = "spearman")
  }
  
  tmp <- orddom(tmp.rand, tmp.path)
  R.path <- as.numeric(tmp[c(23:25), 2])
  
  tmp <- orddom(tmp.rand, tmp.nonpath)
  R.nonpath <- as.numeric(tmp[c(23:25), 2])
  
  CR.path <- c(edge.path - mean(edge.rand), 
               edge.path - mean(edge.rand) - 2*sd(edge.rand)/10, 
               edge.path - mean(edge.rand) + 2*sd(edge.rand)/10)
  CD.path <- SpearmanRho(rowSums(adj.TRN)[id.disease], colSums(adj.PPI)[id.disease], conf.level = 0.95)
  
  CR.nonpath <- c(mean(edge.nonpath) - mean(edge.rand), 
                  mean(edge.nonpath) - mean(edge.rand) - 2*sqrt(var(edge.rand) + var(edge.nonpath))/10, 
                  mean(edge.nonpath) - mean(edge.rand) + 2*sqrt(var(edge.rand) + var(edge.nonpath))/10)
  CD.nonpath <- mean(cor.nonpath, na.rm = T) + c(0, - 2*sd(cor.nonpath, na.rm = T)/10, + 2*sd(cor.nonpath, na.rm = T)/10)
  
  if( j == 1 ){
    data.plot <- data.frame(Org = rep(paste(Pathogens[j], " HINT", sep = ""), 2), 
                            CR = c(CR.path[1], CR.nonpath[1]),
                            CR.lower = c(CR.path[2], CR.nonpath[2]),
                            CR.upper = c(CR.path[3], CR.nonpath[3]),
                            CD = c(CD.path[1], CD.nonpath[1]),
                            CD.lower = c(CD.path[2], CD.nonpath[2]),
                            CD.upper = c(CD.path[3], CD.nonpath[3]),
                            R = c(R.path[1], R.nonpath[1]), 
                            R.lower = c(R.path[2], R.nonpath[2]), 
                            R.upper = c(R.path[3], R.nonpath[3]), 
                            Type = c("Pathogen", "non Pathogen"), 
                            size = c(length(id.disease), length(id.non.disease)))
  }else{
    data.tmp <- data.frame(Org = rep(paste(Pathogens[j], " HINT", sep = ""), 2), 
                           CR = c(CR.path[1], CR.nonpath[1]),
                           CR.lower = c(CR.path[2], CR.nonpath[2]),
                           CR.upper = c(CR.path[3], CR.nonpath[3]),
                           CD = c(CD.path[1], CD.nonpath[1]),
                           CD.lower = c(CD.path[2], CD.nonpath[2]),
                           CD.upper = c(CD.path[3], CD.nonpath[3]),
                           R = c(R.path[1], R.nonpath[1]), 
                           R.lower = c(R.path[2], R.nonpath[2]), 
                           R.upper = c(R.path[3], R.nonpath[3]), 
                           Type = c("Pathogen", "non Pathogen"), 
                           size = c(length(id.disease), length(id.non.disease)))
    data.plot <- rbind(data.plot, data.tmp)
  }
  
  ##BioGRID
  adj.TRN <- NetworksMultiplex$BioGrid[[i]]$TRN
  adj.PPI <- NetworksMultiplex$BioGrid[[i]]$PPI
  id.inter <- NetworksMultiplex$BioGrid[[i]]$genes
  
  id.match <- match(toupper(id.inter), toupper(Genes.HPI[[j]]))
  id.disease <- which(!is.na(id.match))
  id.non.disease <- setdiff(1:length(id.inter), id.disease)
  
  dir.write <- paste(getwd(), "/", Organisms_file[i], "_BioGrid_", sub("\\.", "", 
                                                                       sub(" ", "", paste(Pathogens[j], sep = "_"))), 
                     "_disease_attack_curve", sep = "")
  tmp.path <- rep(0, nboot)
  for( b in 1:nboot ){
    tmp.MCGC <- c(unlist(read.table(paste(dir.write, "/boot_", b, sep = ""), sep = "\t", header = F)))
    id <- which(tmp.MCGC <= 1)[1]
    tmp.path[b] <- abs(sintegral((1:length(tmp.MCGC)/ncol(adj.TRN)), tmp.MCGC/ncol(adj.TRN))$value)
  }
  edge.path <- count_common_edges_among_nodes_complete_iterator(adj.TRN, adj.PPI, id.disease - 1)
  
  dir.write <- paste(getwd(), "/", Organisms_file[i], "_BioGrid_", sub("\\.", "", 
                                                                       sub(" ", "", paste(Pathogens[j], sep = "_"))), 
                     "_random_attack_curve", sep = "")
  tmp.rand <- rep(0, nboot)
  for( b in 1:nboot ){
    tmp.MCGC <- c(unlist(read.table(paste(dir.write, "/boot_", b, sep = ""), sep = "\t", header = F)))
    id <- which(tmp.MCGC <= 1)[1]
    tmp.rand[b] <- abs(sintegral((1:length(tmp.MCGC)/ncol(adj.TRN)), tmp.MCGC/ncol(adj.TRN))$value)
  }
  dir.write <- paste(getwd(), "/", Organisms_file[i], "_BioGrid_", sub("\\.", "", 
                                                                       sub(" ", "", paste(Pathogens[j], sep = "_"))), 
                     "_random_ids", sep = "")
  edge.rand <- rep(0, nboot)
  for( b in 1:nboot ){
    id <- c(unlist(read.table(paste(dir.write, "/boot_", b, sep = ""), sep = "\t", header = F)))
    edge.rand[b] <- count_common_edges_among_nodes_complete_iterator(adj.TRN, adj.PPI, id)
  }
  
  dir.write <- paste(getwd(), "/", Organisms_file[i], "_BioGrid_", sub("\\.", "", 
                                                                       sub(" ", "", paste(Pathogens[j], sep = "_"))), 
                     "_nondisease_attack_curve", sep = "")
  tmp.nonpath <- rep(0, nboot)
  for( b in 1:nboot ){
    tmp.MCGC <- c(unlist(read.table(paste(dir.write, "/boot_", b, sep = ""), sep = "\t", header = F)))
    id <- which(tmp.MCGC <= 1)[1]
    tmp.nonpath[b] <- abs(sintegral((1:length(tmp.MCGC)/ncol(adj.TRN)), tmp.MCGC/ncol(adj.TRN))$value)
  }
  dir.write <- paste(getwd(), "/", Organisms_file[i], "_BioGrid_", sub("\\.", "", 
                                                                       sub(" ", "", paste(Pathogens[j], sep = "_"))), 
                     "_nondisease_ids", sep = "")
  edge.nonpath <- cor.nonpath <- rep(0, nboot)
  for( b in 1:nboot ){
    id <- c(unlist(read.table(paste(dir.write, "/boot_", b, sep = ""), sep = "\t", header = F)))
    edge.nonpath[b] <- count_common_edges_among_nodes_complete_iterator(adj.TRN, adj.PPI, id)
    cor.nonpath[b] <- cor(rowSums(adj.TRN)[id + 1], colSums(adj.PPI)[id + 1], method = "spearman")
  }
  
  
  tmp <- orddom(tmp.rand, tmp.path)
  R.path <- as.numeric(tmp[c(23:25), 2])
  
  tmp <- orddom(tmp.rand, tmp.nonpath)
  R.nonpath <- as.numeric(tmp[c(23:25), 2])
  
  CR.path <- c(edge.path - mean(edge.rand), 
               edge.path - mean(edge.rand) - 2*sd(edge.rand)/10, 
               edge.path - mean(edge.rand) + 2*sd(edge.rand)/10)
  
  CD.path <- SpearmanRho(rowSums(adj.TRN)[id.disease], colSums(adj.PPI)[id.disease], conf.level = 0.95)
  
  CR.nonpath <- c(mean(edge.nonpath) - mean(edge.rand), 
                  mean(edge.nonpath) - mean(edge.rand) - 2*sqrt(var(edge.rand) + var(edge.nonpath))/10, 
                  mean(edge.nonpath) - mean(edge.rand) + 2*sqrt(var(edge.rand) + var(edge.nonpath))/10)
  CD.nonpath <- mean(cor.nonpath, na.rm = T) + c(0, - 2*sd(cor.nonpath, na.rm = T)/10, + 2*sd(cor.nonpath, na.rm = T)/10)
  
  data.tmp <- data.frame(Org = rep(paste(Pathogens[j], " BioGRID", sep = ""), 2), 
                         CR = c(CR.path[1], CR.nonpath[1]),
                         CR.lower = c(CR.path[2], CR.nonpath[2]),
                         CR.upper = c(CR.path[3], CR.nonpath[3]),
                         CD = c(CD.path[1], CD.nonpath[1]),
                         CD.lower = c(CD.path[2], CD.nonpath[2]),
                         CD.upper = c(CD.path[3], CD.nonpath[3]),
                         R = c(R.path[1], R.nonpath[1]), 
                         R.lower = c(R.path[2], R.nonpath[2]), 
                         R.upper = c(R.path[3], R.nonpath[3]), 
                         Type = c("Pathogen", "non Pathogen"), 
                         size = c(length(id.disease), length(id.non.disease)))
  data.plot <- rbind(data.plot, data.tmp)
  
}
data.plot.list[[2]] <- data.plot

id <- order(data.plot[data.plot$Type == "Pathogen", 8], decreasing = F)
data.plot$Org <- factor(data.plot$Org, levels = as.character(data.plot[data.plot$Type == "Pathogen", 1][id]))
data.plot$Type <- factor(data.plot$Type, levels = c("Pathogen", "non Pathogen"))
p <- ggbarplot(data.plot, x = "Org", y = "R", 
               fill = "Type",           # change fill color by mpg_level
               group = "Type",
               color = "Type",# jco journal color palett. see ?ggpar         # Sort the value in descending order
               palette = c("red", "blue"),
               sort.by.groups = F,     # Don't sort inside each group
               # x.text.angle = 50,          # Rotate vertically x axis texts
               xlab = "",
               sort.val = "none",
               ylab = expression("R"),
               # legend = "bottom",
               legend.title = "",
               y.text.angle = 0,
               rotate = T,
               font.x = 30, 
               font.y = 20,
               font.tickslab = 20,
               size = 0.9,
               point.size = 0.1,
               font.label = 12,
               stat = "identity",
               theme = theme_pubr(), 
               position = position_dodge(0.8),
)
p1 <- p + geom_hline(yintercept = 0)+ geom_errorbar(aes(ymin = R.lower, ymax = R.upper, group = Type), width = 0.8, 
                                                    position = position_dodge(0.8)) + 
  theme(title = element_text(size = 30, color = "black"), 
        plot.title = element_text(size = 30, color = "black"), 
        axis.title = element_text(size = 30, colour = "black"), 
        legend.text = element_text(size = 25, color = "black"))


id <- order(data.plot[data.plot$Type == "Pathogen", 8], decreasing = F)
data.plot$Org <- factor(data.plot$Org, levels = as.character(data.plot[data.plot$Type == "Pathogen", 1][id]))
data.plot$Type <- factor(data.plot$Type, levels = c("Pathogen", "non Pathogen"))
p <- ggbarplot(data.plot, x = "Org", y = "CR", 
               fill = "Type",           # change fill color by mpg_level
               group = "Type",
               color = "Type",# jco journal color palett. see ?ggpar         # Sort the value in descending order
               palette = c("red", "blue"),
               sort.by.groups = F,     # Don't sort inside each group
               # x.text.angle = 50,          # Rotate vertically x axis texts
               xlab = "",
               sort.val = "none",
               ylab = expression("C"["R"]),
               legend = "none",
               legend.title = "",
               y.text.angle = 0,
               rotate = T,
               font.x = 20, 
               font.y = 20,
               font.tickslab = 20,
               size = 0.9,
               point.size = 0.1,
               font.label = 12,
               stat = "identity",
               theme = theme_pubr(), 
               position = position_dodge(0.8),
)
p2 <- p + geom_hline(yintercept = 0)+ geom_errorbar(aes(ymin = CR.lower, ymax = CR.upper, group = Type), width = 0.8, 
                                                    position = position_dodge(0.8)) + 
  ggtitle(expression(paste("R: Robustness, ", "C"["R"], ": Redundancy", sep = "")))

Plot.5[[2]] <- plot_grid(p1, 
                         p2 + 
                           theme(axis.text.y = element_blank(),
                                 plot.title = element_text(size = 30, color = "black"),
                                 # axis.line.y = element_blank(),
                                 axis.title.y= element_blank(),
                                 axis.ticks.y= element_blank(), 
                                 axis.title.x = element_text(size = 30, colour = "black")),
                         nrow = 1,
                         rel_widths = c(1.2, 1),
                         align = 'h', axis = 'tb')


##Plot Figure 5C
i <- 9
for( j in 1:length(Diseases) ){
  
  ##Print name of the organism.
  message(paste("Disease = ", Diseases[j], sep = ""))
  
  ##Get organism TRN and PPI networks from multiplex network object.
  adj.TRN <- NetworksMultiplex$HINT[[i]]$TRN
  adj.PPI <- NetworksMultiplex$HINT[[i]]$PPI
  id.inter <- NetworksMultiplex$HINT[[i]]$genes
  
  id.match <- match(toupper(id.inter), toupper(Genes.Diseases[[j]]))
  id.disease <- which(!is.na(id.match))
  id.non.disease <- setdiff(1:length(id.inter), id.disease)
  
  
  ##Make organism specific folder to store the attack curves
  dir.write <- paste(getwd(), "/", Organisms_file[i], "_HINT_", sub(" ", "", paste(Diseases[j], sep = "_")), 
                     "_disease_attack_curve", sep = "")
  tmp.path <- rep(0, nboot)
  for( b in 1:nboot ){
    tmp.MCGC <- c(unlist(read.table(paste(dir.write, "/boot_", b, sep = ""), sep = "\t", header = F)))
    id <- which(tmp.MCGC <= 1)[1]
    tmp.path[b] <- abs(sintegral((1:length(tmp.MCGC)/ncol(adj.TRN)), tmp.MCGC/ncol(adj.TRN))$value)
  }
  edge.path <- count_common_edges_among_nodes_complete_iterator(adj.TRN, adj.PPI, id.disease - 1)
  
  dir.write <- paste(getwd(), "/", Organisms_file[i], "_HINT_", sub(" ", "", paste(Diseases[j], sep = "_")), 
                     "_random_attack_curve", sep = "")
  tmp.rand <- rep(0, nboot)
  for( b in 1:nboot ){
    tmp.MCGC <- c(unlist(read.table(paste(dir.write, "/boot_", b, sep = ""), sep = "\t", header = F)))
    id <- which(tmp.MCGC <= 1)[1]
    tmp.rand[b] <- abs(sintegral((1:length(tmp.MCGC)/ncol(adj.TRN)), tmp.MCGC/ncol(adj.TRN))$value)
  }
  dir.write <- paste(getwd(), "/", Organisms_file[i], "_HINT_", sub(" ", "", paste(Diseases[j], sep = "_")), 
                     "_random_ids", sep = "")
  edge.rand <- cor.rand <- rep(0, nboot)
  for( b in 1:nboot ){
    id <- c(unlist(read.table(paste(dir.write, "/boot_", b, sep = ""), sep = "\t", header = F)))
    edge.rand[b] <- count_common_edges_among_nodes_complete_iterator(adj.TRN, adj.PPI, id)
    cor.rand[b] <- cor(rowSums(adj.TRN)[id + 1], colSums(adj.PPI)[id + 1], method = "spearman")
  }
  
  dir.write <- paste(getwd(), "/", Organisms_file[i], "_HINT_", sub(" ", "", paste(Diseases[j], sep = "_")), 
                     "_nondisease_attack_curve", sep = "")
  tmp.nonpath <- rep(0, nboot)
  for( b in 1:nboot ){
    tmp.MCGC <- c(unlist(read.table(paste(dir.write, "/boot_", b, sep = ""), sep = "\t", header = F)))
    id <- which(tmp.MCGC <= 1)[1]
    tmp.nonpath[b] <- abs(sintegral((1:length(tmp.MCGC)/ncol(adj.TRN)), tmp.MCGC/ncol(adj.TRN))$value)
  }
  dir.write <- paste(getwd(), "/", Organisms_file[i], "_HINT_", sub(" ", "", paste(Diseases[j], sep = "_")), 
                     "_nondisease_ids", sep = "")
  edge.nonpath <- cor.nonpath <- rep(0, nboot)
  for( b in 1:nboot ){
    id <- c(unlist(read.table(paste(dir.write, "/boot_", b, sep = ""), sep = "\t", header = F)))
    edge.nonpath[b] <- count_common_edges_among_nodes_complete_iterator(adj.TRN, adj.PPI, id)
    cor.nonpath[b] <- cor(rowSums(adj.TRN)[id + 1], colSums(adj.PPI)[id + 1], method = "spearman")
  }
  
  tmp <- orddom(tmp.rand, tmp.path)
  R.path <- as.numeric(tmp[c(23:25), 2])
  
  tmp <- orddom(tmp.rand, tmp.nonpath)
  R.nonpath <- as.numeric(tmp[c(23:25), 2])
  
  CR.path <- c(edge.path - mean(edge.rand), 
               edge.path - mean(edge.rand) - 2*sd(edge.rand)/10, 
               edge.path - mean(edge.rand) + 2*sd(edge.rand)/10)
  CD.path <- SpearmanRho(rowSums(adj.TRN)[id.disease], colSums(adj.PPI)[id.disease], conf.level = 0.95)
  
  CR.nonpath <- c(mean(edge.nonpath) - mean(edge.rand), 
                  mean(edge.nonpath) - mean(edge.rand) - 2*sqrt(var(edge.rand) + var(edge.nonpath))/10, 
                  mean(edge.nonpath) - mean(edge.rand) + 2*sqrt(var(edge.rand) + var(edge.nonpath))/10)
  CD.nonpath <- mean(cor.nonpath, na.rm = T) + c(0, - 2*sd(cor.nonpath, na.rm = T)/10, + 2*sd(cor.nonpath, na.rm = T)/10)
  
  if( j == 1 ){
    data.plot <- data.frame(Org = rep(paste(Diseases[j], " HINT", sep = ""), 2), 
                            CR = c(CR.path[1], CR.nonpath[1]),
                            CR.lower = c(CR.path[2], CR.nonpath[2]),
                            CR.upper = c(CR.path[3], CR.nonpath[3]),
                            CD = c(CD.path[1], CD.nonpath[1]),
                            CD.lower = c(CD.path[2], CD.nonpath[2]),
                            CD.upper = c(CD.path[3], CD.nonpath[3]),
                            R = c(R.path[1], R.nonpath[1]), 
                            R.lower = c(R.path[2], R.nonpath[2]), 
                            R.upper = c(R.path[3], R.nonpath[3]), 
                            Type = c("Disease", "non Disease"), 
                            size = c(length(id.disease), length(id.non.disease)))
  }else{
    data.tmp <- data.frame(Org = rep(paste(Diseases[j], " HINT", sep = ""), 2), 
                           CR = c(CR.path[1], CR.nonpath[1]),
                           CR.lower = c(CR.path[2], CR.nonpath[2]),
                           CR.upper = c(CR.path[3], CR.nonpath[3]),
                           CD = c(CD.path[1], CD.nonpath[1]),
                           CD.lower = c(CD.path[2], CD.nonpath[2]),
                           CD.upper = c(CD.path[3], CD.nonpath[3]),
                           R = c(R.path[1], R.nonpath[1]), 
                           R.lower = c(R.path[2], R.nonpath[2]), 
                           R.upper = c(R.path[3], R.nonpath[3]), 
                           Type = c("Disease", "non Disease"), 
                           size = c(length(id.disease), length(id.non.disease)))
    data.plot <- rbind(data.plot, data.tmp)
  }
  
  ##BioGRID
  adj.TRN <- NetworksMultiplex$BioGrid[[i]]$TRN
  adj.PPI <- NetworksMultiplex$BioGrid[[i]]$PPI
  id.inter <- NetworksMultiplex$BioGrid[[i]]$genes
  
  id.match <- match(toupper(id.inter), toupper(Genes.Diseases[[j]]))
  id.disease <- which(!is.na(id.match))
  id.non.disease <- setdiff(1:length(id.inter), id.disease)
  
  dir.write <- paste(getwd(), "/", Organisms_file[i], "_BioGrid_", sub(" ", "", paste(Diseases[j], sep = "_")), 
                     "_disease_attack_curve", sep = "")
  tmp.path <- rep(0, nboot)
  for( b in 1:nboot ){
    tmp.MCGC <- c(unlist(read.table(paste(dir.write, "/boot_", b, sep = ""), sep = "\t", header = F)))
    id <- which(tmp.MCGC <= 1)[1]
    tmp.path[b] <- abs(sintegral((1:length(tmp.MCGC)/ncol(adj.TRN)), tmp.MCGC/ncol(adj.TRN))$value)
  }
  edge.path <- count_common_edges_among_nodes_complete_iterator(adj.TRN, adj.PPI, id.disease - 1)
  
  dir.write <- paste(getwd(), "/", Organisms_file[i], "_BioGrid_", sub(" ", "", paste(Diseases[j], sep = "_")), 
                     "_random_attack_curve", sep = "")
  tmp.rand <- rep(0, nboot)
  for( b in 1:nboot ){
    tmp.MCGC <- c(unlist(read.table(paste(dir.write, "/boot_", b, sep = ""), sep = "\t", header = F)))
    id <- which(tmp.MCGC <= 1)[1]
    tmp.rand[b] <- abs(sintegral((1:length(tmp.MCGC)/ncol(adj.TRN)), tmp.MCGC/ncol(adj.TRN))$value)
  }
  dir.write <- paste(getwd(), "/", Organisms_file[i], "_BioGrid_", sub(" ", "", paste(Diseases[j], sep = "_")), 
                     "_random_ids", sep = "")
  edge.rand <- rep(0, nboot)
  for( b in 1:nboot ){
    id <- c(unlist(read.table(paste(dir.write, "/boot_", b, sep = ""), sep = "\t", header = F)))
    edge.rand[b] <- count_common_edges_among_nodes_complete_iterator(adj.TRN, adj.PPI, id)
  }
  
  dir.write <- paste(getwd(), "/", Organisms_file[i], "_BioGrid_", sub(" ", "", paste(Diseases[j], sep = "_")), 
                     "_nondisease_attack_curve", sep = "")
  tmp.nonpath <- rep(0, nboot)
  for( b in 1:nboot ){
    tmp.MCGC <- c(unlist(read.table(paste(dir.write, "/boot_", b, sep = ""), sep = "\t", header = F)))
    id <- which(tmp.MCGC <= 1)[1]
    tmp.nonpath[b] <- abs(sintegral((1:length(tmp.MCGC)/ncol(adj.TRN)), tmp.MCGC/ncol(adj.TRN))$value)
  }
  dir.write <- paste(getwd(), "/", Organisms_file[i], "_BioGrid_", sub(" ", "", paste(Diseases[j], sep = "_")), 
                     "_nondisease_ids", sep = "")
  edge.nonpath <- cor.nonpath <- rep(0, nboot)
  for( b in 1:nboot ){
    id <- c(unlist(read.table(paste(dir.write, "/boot_", b, sep = ""), sep = "\t", header = F)))
    edge.nonpath[b] <- count_common_edges_among_nodes_complete_iterator(adj.TRN, adj.PPI, id)
    cor.nonpath[b] <- cor(rowSums(adj.TRN)[id + 1], colSums(adj.PPI)[id + 1], method = "spearman")
  }
  
  
  tmp <- orddom(tmp.rand, tmp.path)
  R.path <- as.numeric(tmp[c(23:25), 2])
  
  tmp <- orddom(tmp.rand, tmp.nonpath)
  R.nonpath <- as.numeric(tmp[c(23:25), 2])
  
  CR.path <- c(edge.path - mean(edge.rand), 
               edge.path - mean(edge.rand) - 2*sd(edge.rand)/10, 
               edge.path - mean(edge.rand) + 2*sd(edge.rand)/10)
  
  CD.path <- SpearmanRho(rowSums(adj.TRN)[id.disease], colSums(adj.PPI)[id.disease], conf.level = 0.95)
  
  CR.nonpath <- c(mean(edge.nonpath) - mean(edge.rand), 
                  mean(edge.nonpath) - mean(edge.rand) - 2*sqrt(var(edge.rand) + var(edge.nonpath))/10, 
                  mean(edge.nonpath) - mean(edge.rand) + 2*sqrt(var(edge.rand) + var(edge.nonpath))/10)
  CD.nonpath <- mean(cor.nonpath, na.rm = T) + c(0, - 2*sd(cor.nonpath, na.rm = T)/10, + 2*sd(cor.nonpath, na.rm = T)/10)
  
  data.tmp <- data.frame(Org = rep(paste(Diseases[j], " BioGRID", sep = ""), 2), 
                         CR = c(CR.path[1], CR.nonpath[1]),
                         CR.lower = c(CR.path[2], CR.nonpath[2]),
                         CR.upper = c(CR.path[3], CR.nonpath[3]),
                         CD = c(CD.path[1], CD.nonpath[1]),
                         CD.lower = c(CD.path[2], CD.nonpath[2]),
                         CD.upper = c(CD.path[3], CD.nonpath[3]),
                         R = c(R.path[1], R.nonpath[1]), 
                         R.lower = c(R.path[2], R.nonpath[2]), 
                         R.upper = c(R.path[3], R.nonpath[3]), 
                         Type = c("Disease", "non Disease"), 
                         size = c(length(id.disease), length(id.non.disease)))
  data.plot <- rbind(data.plot, data.tmp)
  
}
data.plot.list[[3]] <- data.plot

id <- order(data.plot[data.plot$Type == "Disease", 8], decreasing = F)
data.plot$Org <- factor(data.plot$Org, levels = as.character(data.plot[data.plot$Type == "Disease", 1][id]))
data.plot$Type <- factor(data.plot$Type, levels = c("Disease", "non Disease"))
p <- ggbarplot(data.plot, x = "Org", y = "R", 
               fill = "Type",           # change fill color by mpg_level
               group = "Type",
               color = "Type",# jco journal color palett. see ?ggpar         # Sort the value in descending order
               palette = c("red", "blue"),
               sort.by.groups = F,     # Don't sort inside each group
               # x.text.angle = 50,          # Rotate vertically x axis texts
               xlab = "",
               sort.val = "none",
               ylab = expression("R"),
               legend = "none",
               legend.title = "",
               y.text.angle = 0,
               rotate = T,
               font.x = 30, 
               font.y = 20,
               font.tickslab = 20,
               size = 0.9,
               point.size = 0.1,
               font.label = 12,
               stat = "identity",
               theme = theme_pubr(), 
               position = position_dodge(0.8),
)
p1 <- p + geom_hline(yintercept = 0)+ geom_errorbar(aes(ymin = R.lower, ymax = R.upper, group = Type), width = 0.8, 
                                                    position = position_dodge(0.8)) + 
  theme(title = element_text(size = 20, color = "black"), 
        plot.title = element_text(size = 30, color = "black"), 
        axis.title = element_text(size = 30, colour = "black"), 
        legend.text = element_text(size = 25, color = "black"), legend.position = "top")


id <- order(data.plot[data.plot$Type == "Disease", 8], decreasing = F)
data.plot$Org <- factor(data.plot$Org, levels = as.character(data.plot[data.plot$Type == "Disease", 1][id]))
data.plot$Type <- factor(data.plot$Type, levels = c("Disease", "non Disease"))
p <- ggbarplot(data.plot, x = "Org", y = "CR", 
               fill = "Type",           # change fill color by mpg_level
               group = "Type",
               color = "Type",# jco journal color palett. see ?ggpar         # Sort the value in descending order
               palette = c("red", "blue"),
               sort.by.groups = F,     # Don't sort inside each group
               # x.text.angle = 50,          # Rotate vertically x axis texts
               xlab = "",
               sort.val = "none",
               ylab = expression("C"["R"]),
               legend = "none",
               legend.title = "",
               y.text.angle = 50,
               rotate = T,
               font.x = 20, 
               font.y = 20,
               font.tickslab = 20,
               size = 0.9,
               point.size = 0.1,
               font.label = 12,
               stat = "identity",
               theme = theme_pubr(), 
               position = position_dodge(0.8),
)
p2 <- p + geom_hline(yintercept = 0)+ geom_errorbar(aes(ymin = CR.lower, ymax = CR.upper, group = Type), width = 0.8, 
                                                    position = position_dodge(0.8)) + 
  ggtitle(expression(paste("R: Robustness, ", "C"["R"], ": Redundancy", sep = "")))

Plot.5[[3]] <- plot_grid(p1, 
                         p2 + 
                           theme(axis.text.y = element_blank(),
                                 plot.title = element_text(size = 30, color = "black"),
                                 # axis.line.y = element_blank(),
                                 axis.title.y= element_blank(),
                                 axis.ticks.y= element_blank(), 
                                 axis.title.x = element_text(size = 30, colour = "black")),
                         nrow = 1,
                         rel_widths = c(1.2, 1),
                         align = 'h', axis = 'tb')


##Plot Figure 5D
Cancer.gene.type <- c("Oncogene", "TSG")
for( j in 1:2 ){
  
  ##Print name of the organism.
  message(paste("Cancer = ", Cancer.gene.type[j], sep = ""))
  
  ##Get organism TRN and PPI networks from multiplex network object.
  adj.TRN <- NetworksMultiplex$HINT[[i]]$TRN
  adj.PPI <- NetworksMultiplex$HINT[[i]]$PPI
  id.inter <- NetworksMultiplex$HINT[[i]]$genes
  
  id.match <- match(toupper(id.inter), toupper(Genes.Diseases[[j]]))
  id.disease <- which(!is.na(id.match))
  id.non.disease <- setdiff(1:length(id.inter), id.disease)
  
  
  ##Make organism specific folder to store the attack curves
  if( j == 1 ){
    dir.write <- paste(getwd(), "/", Organisms_file[i], "_HINT_", 
                       "onco_attack_curve", sep = "")
  }else{
    dir.write <- paste(getwd(), "/", Organisms_file[i], "_HINT_", 
                       "TSG_attack_curve", sep = "")
  }
  tmp.path <- rep(0, nboot)
  for( b in 1:nboot ){
    tmp.MCGC <- c(unlist(read.table(paste(dir.write, "/boot_", b, sep = ""), sep = "\t", header = F)))
    id <- which(tmp.MCGC <= 1)[1]
    tmp.path[b] <- abs(sintegral((1:length(tmp.MCGC)/ncol(adj.TRN)), tmp.MCGC/ncol(adj.TRN))$value)
  }
  edge.path <- count_common_edges_among_nodes_complete_iterator(adj.TRN, adj.PPI, id.disease - 1)
  
  if( j == 1 ){
    dir.write <- paste(getwd(), "/", Organisms_file[i], "_HINT_", 
                       "randomonco_attack_curve", sep = "")
  }else{
    dir.write <- paste(getwd(), "/", Organisms_file[i], "_HINT_", 
                       "randomTSG_attack_curve", sep = "")
  }
  tmp.rand <- rep(0, nboot)
  for( b in 1:nboot ){
    tmp.MCGC <- c(unlist(read.table(paste(dir.write, "/boot_", b, sep = ""), sep = "\t", header = F)))
    id <- which(tmp.MCGC <= 1)[1]
    tmp.rand[b] <- abs(sintegral((1:length(tmp.MCGC)/ncol(adj.TRN)), tmp.MCGC/ncol(adj.TRN))$value)
  }
  if( j == 1 ){
    dir.write <- paste(getwd(), "/", Organisms_file[i], "_HINT_", 
                       "randomonco_ids", sep = "")
  }else{
    dir.write <- paste(getwd(), "/", Organisms_file[i], "_HINT_", 
                       "randomTSG_ids", sep = "")
  }
  edge.rand <- cor.rand <- rep(0, nboot)
  for( b in 1:nboot ){
    id <- c(unlist(read.table(paste(dir.write, "/boot_", b, sep = ""), sep = "\t", header = F)))
    edge.rand[b] <- count_common_edges_among_nodes_complete_iterator(adj.TRN, adj.PPI, id)
    cor.rand[b] <- cor(rowSums(adj.TRN)[id + 1], colSums(adj.PPI)[id + 1], method = "spearman")
  }
  
  if( j == 1 ){
    dir.write <- paste(getwd(), "/", Organisms_file[i], "_HINT_", 
                       "nononco_attack_curve", sep = "")
  }else{
    dir.write <- paste(getwd(), "/", Organisms_file[i], "_HINT_", 
                       "nonTSG_attack_curve", sep = "")
  }
  tmp.nonpath <- rep(0, nboot)
  for( b in 1:nboot ){
    tmp.MCGC <- c(unlist(read.table(paste(dir.write, "/boot_", b, sep = ""), sep = "\t", header = F)))
    id <- which(tmp.MCGC <= 1)[1]
    tmp.nonpath[b] <- abs(sintegral((1:length(tmp.MCGC)/ncol(adj.TRN)), tmp.MCGC/ncol(adj.TRN))$value)
  }
  if( j == 1 ){
    dir.write <- paste(getwd(), "/", Organisms_file[i], "_HINT_", 
                       "nononco_ids", sep = "")
  }else{
    dir.write <- paste(getwd(), "/", Organisms_file[i], "_HINT_", 
                       "nonTSG_ids", sep = "")
  }
  edge.nonpath <- cor.nonpath <- rep(0, nboot)
  for( b in 1:nboot ){
    id <- c(unlist(read.table(paste(dir.write, "/boot_", b, sep = ""), sep = "\t", header = F)))
    edge.nonpath[b] <- count_common_edges_among_nodes_complete_iterator(adj.TRN, adj.PPI, id)
    cor.nonpath[b] <- cor(rowSums(adj.TRN)[id + 1], colSums(adj.PPI)[id + 1], method = "spearman")
  }
  
  tmp <- orddom(tmp.rand, tmp.path)
  R.path <- as.numeric(tmp[c(23:25), 2])
  
  tmp <- orddom(tmp.rand, tmp.nonpath)
  R.nonpath <- as.numeric(tmp[c(23:25), 2])
  
  CR.path <- c(edge.path - mean(edge.rand), 
               edge.path - mean(edge.rand) - 2*sd(edge.rand)/10, 
               edge.path - mean(edge.rand) + 2*sd(edge.rand)/10)
  CD.path <- SpearmanRho(rowSums(adj.TRN)[id.disease], colSums(adj.PPI)[id.disease], conf.level = 0.95)
  
  CR.nonpath <- c(mean(edge.nonpath) - mean(edge.rand), 
                  mean(edge.nonpath) - mean(edge.rand) - 2*sqrt(var(edge.rand) + var(edge.nonpath))/10, 
                  mean(edge.nonpath) - mean(edge.rand) + 2*sqrt(var(edge.rand) + var(edge.nonpath))/10)
  CD.nonpath <- mean(cor.nonpath, na.rm = T) + c(0, - 2*sd(cor.nonpath, na.rm = T)/10, + 2*sd(cor.nonpath, na.rm = T)/10)
  
  if( j == 1 ){
    data.plot <- data.frame(Org = rep(paste(Cancer.gene.type[j], " HINT", sep = ""), 2), 
                            CR = c(CR.path[1], CR.nonpath[1]),
                            CR.lower = c(CR.path[2], CR.nonpath[2]),
                            CR.upper = c(CR.path[3], CR.nonpath[3]),
                            CD = c(CD.path[1], CD.nonpath[1]),
                            CD.lower = c(CD.path[2], CD.nonpath[2]),
                            CD.upper = c(CD.path[3], CD.nonpath[3]),
                            R = c(R.path[1], R.nonpath[1]), 
                            R.lower = c(R.path[2], R.nonpath[2]), 
                            R.upper = c(R.path[3], R.nonpath[3]), 
                            Type = c("Cancer", "non Cancer"), 
                            size = c(length(id.disease), length(id.non.disease)))
  }else{
    data.tmp <- data.frame(Org = rep(paste(Cancer.gene.type[j], " HINT", sep = ""), 2), 
                           CR = c(CR.path[1], CR.nonpath[1]),
                           CR.lower = c(CR.path[2], CR.nonpath[2]),
                           CR.upper = c(CR.path[3], CR.nonpath[3]),
                           CD = c(CD.path[1], CD.nonpath[1]),
                           CD.lower = c(CD.path[2], CD.nonpath[2]),
                           CD.upper = c(CD.path[3], CD.nonpath[3]),
                           R = c(R.path[1], R.nonpath[1]), 
                           R.lower = c(R.path[2], R.nonpath[2]), 
                           R.upper = c(R.path[3], R.nonpath[3]), 
                           Type = c("Cancer", "non Cancer"), 
                           size = c(length(id.disease), length(id.non.disease)))
    data.plot <- rbind(data.plot, data.tmp)
  }
  
  ##BioGRID
  adj.TRN <- NetworksMultiplex$BioGrid[[i]]$TRN
  adj.PPI <- NetworksMultiplex$BioGrid[[i]]$PPI
  id.inter <- NetworksMultiplex$BioGrid[[i]]$genes
  
  id.match <- match(toupper(id.inter), toupper(Genes.Diseases[[j]]))
  id.disease <- which(!is.na(id.match))
  id.non.disease <- setdiff(1:length(id.inter), id.disease)
  
  if( j == 1 ){
    dir.write <- paste(getwd(), "/", Organisms_file[i], "_BioGrid_", 
                       "onco_attack_curve", sep = "")
  }else{
    dir.write <- paste(getwd(), "/", Organisms_file[i], "_BioGrid_", 
                       "TSG_attack_curve", sep = "")
  }
  tmp.path <- rep(0, nboot)
  for( b in 1:nboot ){
    tmp.MCGC <- c(unlist(read.table(paste(dir.write, "/boot_", b, sep = ""), sep = "\t", header = F)))
    id <- which(tmp.MCGC <= 1)[1]
    tmp.path[b] <- abs(sintegral((1:length(tmp.MCGC)/ncol(adj.TRN)), tmp.MCGC/ncol(adj.TRN))$value)
  }
  edge.path <- count_common_edges_among_nodes_complete_iterator(adj.TRN, adj.PPI, id.disease - 1)
  
  if( j == 1 ){
    dir.write <- paste(getwd(), "/", Organisms_file[i], "_BioGrid_", 
                       "randomonco_attack_curve", sep = "")
  }else{
    dir.write <- paste(getwd(), "/", Organisms_file[i], "_BioGrid_", 
                       "randomTSG_attack_curve", sep = "")
  }
  tmp.rand <- rep(0, nboot)
  for( b in 1:nboot ){
    tmp.MCGC <- c(unlist(read.table(paste(dir.write, "/boot_", b, sep = ""), sep = "\t", header = F)))
    id <- which(tmp.MCGC <= 1)[1]
    tmp.rand[b] <- abs(sintegral((1:length(tmp.MCGC)/ncol(adj.TRN)), tmp.MCGC/ncol(adj.TRN))$value)
  }
  if( j == 1 ){
    dir.write <- paste(getwd(), "/", Organisms_file[i], "_BioGrid_", 
                       "randomonco_ids", sep = "")
  }else{
    dir.write <- paste(getwd(), "/", Organisms_file[i], "_BioGrid_", 
                       "randomTSG_ids", sep = "")
  }
  edge.rand <- rep(0, nboot)
  for( b in 1:nboot ){
    id <- c(unlist(read.table(paste(dir.write, "/boot_", b, sep = ""), sep = "\t", header = F)))
    edge.rand[b] <- count_common_edges_among_nodes_complete_iterator(adj.TRN, adj.PPI, id)
  }
  
  if( j == 1 ){
    dir.write <- paste(getwd(), "/", Organisms_file[i], "_BioGrid_", 
                       "nononco_attack_curve", sep = "")
  }else{
    dir.write <- paste(getwd(), "/", Organisms_file[i], "_BioGrid_", 
                       "nonTSG_attack_curve", sep = "")
  }
  tmp.nonpath <- rep(0, nboot)
  for( b in 1:nboot ){
    tmp.MCGC <- c(unlist(read.table(paste(dir.write, "/boot_", b, sep = ""), sep = "\t", header = F)))
    id <- which(tmp.MCGC <= 1)[1]
    tmp.nonpath[b] <- abs(sintegral((1:length(tmp.MCGC)/ncol(adj.TRN)), tmp.MCGC/ncol(adj.TRN))$value)
  }
  if( j == 1 ){
    dir.write <- paste(getwd(), "/", Organisms_file[i], "_BioGrid_", 
                       "nononco_ids", sep = "")
  }else{
    dir.write <- paste(getwd(), "/", Organisms_file[i], "_BioGrid_", 
                       "nonTSG_ids", sep = "")
  }
  edge.nonpath <- cor.nonpath <- rep(0, nboot)
  for( b in 1:nboot ){
    id <- c(unlist(read.table(paste(dir.write, "/boot_", b, sep = ""), sep = "\t", header = F)))
    edge.nonpath[b] <- count_common_edges_among_nodes_complete_iterator(adj.TRN, adj.PPI, id)
    cor.nonpath[b] <- cor(rowSums(adj.TRN)[id + 1], colSums(adj.PPI)[id + 1], method = "spearman")
  }
  
  
  tmp <- orddom(tmp.rand, tmp.path)
  R.path <- as.numeric(tmp[c(23:25), 2])
  
  tmp <- orddom(tmp.rand, tmp.nonpath)
  R.nonpath <- as.numeric(tmp[c(23:25), 2])
  
  CR.path <- c(edge.path - mean(edge.rand), 
               edge.path - mean(edge.rand) - 2*sd(edge.rand)/10, 
               edge.path - mean(edge.rand) + 2*sd(edge.rand)/10)
  
  CD.path <- SpearmanRho(rowSums(adj.TRN)[id.disease], colSums(adj.PPI)[id.disease], conf.level = 0.95)
  
  CR.nonpath <- c(mean(edge.nonpath) - mean(edge.rand), 
                  mean(edge.nonpath) - mean(edge.rand) - 2*sqrt(var(edge.rand) + var(edge.nonpath))/10, 
                  mean(edge.nonpath) - mean(edge.rand) + 2*sqrt(var(edge.rand) + var(edge.nonpath))/10)
  CD.nonpath <- mean(cor.nonpath, na.rm = T) + c(0, - 2*sd(cor.nonpath, na.rm = T)/10, + 2*sd(cor.nonpath, na.rm = T)/10)
  
  data.tmp <- data.frame(Org = rep(paste(Cancer.gene.type[j], " BioGRID", sep = ""), 2), 
                         CR = c(CR.path[1], CR.nonpath[1]),
                         CR.lower = c(CR.path[2], CR.nonpath[2]),
                         CR.upper = c(CR.path[3], CR.nonpath[3]),
                         CD = c(CD.path[1], CD.nonpath[1]),
                         CD.lower = c(CD.path[2], CD.nonpath[2]),
                         CD.upper = c(CD.path[3], CD.nonpath[3]),
                         R = c(R.path[1], R.nonpath[1]), 
                         R.lower = c(R.path[2], R.nonpath[2]), 
                         R.upper = c(R.path[3], R.nonpath[3]), 
                         Type = c("Cancer", "non Cancer"), 
                         size = c(length(id.disease), length(id.non.disease)))
  data.plot <- rbind(data.plot, data.tmp)
  
}
data.plot.list[[4]] <- data.plot

id <- order(data.plot[data.plot$Type == "Cancer", 8], decreasing = F)
data.plot$Org <- factor(data.plot$Org, levels = as.character(data.plot[data.plot$Type == "Cancer", 1][id]))
data.plot$Type <- factor(data.plot$Type, levels = c("Cancer", "non Cancer"))
p <- ggbarplot(data.plot, x = "Org", y = "R", 
               fill = "Type",           # change fill color by mpg_level
               group = "Type",
               color = "Type",# jco journal color palett. see ?ggpar         # Sort the value in descending order
               palette = c("red", "blue"),
               sort.by.groups = F,     # Don't sort inside each group
               # x.text.angle = 50,          # Rotate vertically x axis texts
               xlab = "",
               sort.val = "none",
               ylab = expression("R"),
               # legend = "bottom",
               legend.title = "",
               y.text.angle = 50,
               rotate = T,
               font.x = 30, 
               font.y = 20,
               font.tickslab = 20,
               size = 0.9,
               point.size = 0.1,
               font.label = 12,
               stat = "identity",
               theme = theme_pubr(), 
               position = position_dodge(0.8),
)
p1 <- p + geom_hline(yintercept = 0)+ geom_errorbar(aes(ymin = R.lower, ymax = R.upper, group = Type), width = 0.2, 
                                                    position = position_dodge(0.8)) + 
  theme(title = element_text(size = 30, color = "black"), 
        plot.title = element_text(size = 30, color = "black"), 
        axis.title = element_text(size = 30, colour = "black"), 
        legend.text = element_text(size = 25, color = "black"))


id <- order(data.plot[data.plot$Type == "Cancer", 8], decreasing = F)
data.plot$Org <- factor(data.plot$Org, levels = as.character(data.plot[data.plot$Type == "Cancer", 1][id]))
data.plot$Type <- factor(data.plot$Type, levels = c("Cancer", "non Cancer"))
p <- ggbarplot(data.plot, x = "Org", y = "CR", 
               fill = "Type",           # change fill color by mpg_level
               group = "Type",
               color = "Type",# jco journal color palett. see ?ggpar         # Sort the value in descending order
               palette = c("red", "blue"),
               sort.by.groups = F,     # Don't sort inside each group
               # x.text.angle = 50,          # Rotate vertically x axis texts
               xlab = "",
               sort.val = "none",
               ylab = expression("C"["R"]),
               legend = "none",
               legend.title = "",
               y.text.angle = 50,
               rotate = T,
               font.x = 20, 
               font.y = 20,
               font.tickslab = 20,
               size = 0.9,
               point.size = 0.1,
               font.label = 12,
               stat = "identity",
               theme = theme_pubr(), 
               position = position_dodge(0.8),
)
p2 <- p + geom_hline(yintercept = 0)+ geom_errorbar(aes(ymin = CR.lower, ymax = CR.upper, group = Type), width = 0.2, 
                                                    position = position_dodge(0.8)) + 
  ggtitle(expression(paste("R: Robustness, ", "C"["R"], ": Redundancy", sep = "")))

Plot.5[[4]] <- plot_grid(p1, 
                         p2 + 
                           theme(axis.text.y = element_blank(),
                                 plot.title = element_text(size = 30, color = "black"),
                                 # axis.line.y = element_blank(),
                                 axis.title.y= element_blank(),
                                 axis.ticks.y= element_blank(), 
                                 axis.title.x = element_text(size = 30, colour = "black")),
                         nrow = 1,
                         rel_widths = c(1.2, 1),
                         align = 'h', axis = 'tb')

plot1 <- plot_grid(Plot.5[[1]], Plot.5[[2]], nrow = 2, labels = NULL, 
                   align = "v", rel_heights = c(1, 2))
plot2 <- plot_grid(Plot.5[[3]], Plot.5[[4]], nrow = 2, labels = NULL, 
                   align = "v", rel_heights = c(2, 1))
plot1 <- plot_grid(plot1, plot2, ncol = 2, labels = NULL, 
                   align = "h", rel_widths = c(1, 1))
save_plot(paste("Figure_5_paper_grid_new_size.png", sep = ""), 
          plot1, base_height = 26, base_width = 34)
save_plot(paste("Figure_5_paper_grid_new_size.pdf", sep = ""), 
          plot1, base_height = 26, base_width = 34)
```
