## Supplementary material for "Internetwork connectivity of molecular networks across species of life": Code: Simulated_Annealing_Functions.cpp

```
//Simulated Annealing code
#include <RcppArmadillo.h>
#include <iostream>
#include <fstream>
#include <list>
#include <vector>
#include <cmath>
#include <omp.h>
#include <progress.hpp>
#include <progress_bar.hpp>
#include <Attack_functions_new_cpp.cpp>
#define NIL -1 
// #include <otherpackage.h>
// #include "/igraph/igraph.h"
//[[Rcpp::depends(RcppProgress, RcppArmadillo)]]
using namespace Rcpp;
using namespace std;
using namespace arma;


//[[Rcpp::export]]
arma::vec count_common_edges_among_nodes_iterator(
    arma::sp_mat adjTRN, arma::sp_mat adjPPI,
    arma::vec nodes
){
  arma::vec edge_count(nodes.size(), arma::fill::zeros);
  sp_mat::const_iterator it     = adjTRN.begin();
  sp_mat::const_iterator it_end = adjTRN.end();
  for(; it != it_end; ++it){
    if( adjPPI(it.row(), it.col()) == 1 ){
      for( int j = 0; j < nodes.size(); j++ ){
        if( int(nodes(j)) == int(it.row()) || int(nodes(j)) == int(it.col()) ){
          edge_count(j) += 1;
        }
      }
    }
    
  }
  return edge_count;
}

//[[Rcpp::export]]
int count_common_edges_among_nodes_complete_iterator(
    arma::sp_mat adjTRN, arma::sp_mat adjPPI,
    arma::uvec nodes
){
  int edge_count = 0;
  sp_mat::const_iterator it     = adjTRN.begin();
  sp_mat::const_iterator it_end = adjTRN.end();
  for(; it != it_end; ++it){
    if( adjPPI(it.row(), it.col()) == 1 ){
      for( int j = 0; j < nodes.size(); j++ ){
        if( int(nodes(j)) == int(it.row()) || int(nodes(j)) == int(it.col()) ){
          edge_count += 1;
          break;
        }
      }
    }
    
  }
  return edge_count;
}

//[[Rcpp::export]]
int count_red_mult_iterator(
    arma::sp_mat adjTRN, arma::sp_mat adjPPI,
    arma::uvec nodes
){
  int edge_count = 0;
  sp_mat::const_iterator it     = adjTRN.begin();
  sp_mat::const_iterator it_end = adjTRN.end();
  for(; it != it_end; ++it){
    if( adjPPI(it.row(), it.col()) == 1 ){
      int test1 = 0;
      int test2 = 0;
      for( int j = 0; j < nodes.size(); j++ ){
        if( int(nodes(j)) == int(it.row()) ){
          test1 = 1;
        }
        if( int(nodes(j)) == int(it.col()) ){
          test2 = 1;
        }
        if( test1 == 1 && test2 == 1 ){
          edge_count += 1;
          break;
        }
      }
    }
    
  }
  return edge_count;
}

arma::vec rankify(arma::vec & X) { 
  
  int N = X.size(); 
  
  // Rank Vector 
  arma::vec Rank_X(N); 
  
  for( int i = 0; i < N; ++i )  
  { 
    int r = 1, s = 1; 
    
    // Count no of smaller elements 
    // in 0 to i-1 
    for(int j = 0; j < i; j++) { 
      if (X(j) < X(i) ) r++; 
      if (X(j) == X(i) ) s++; 
    } 
    
    // Count no of smaller elements 
    // in i+1 to N-1 
    for (int j = i+1; j < N; j++) { 
      if (X(j) < X(i) ) r++; 
      if (X(j) == X(i) ) s++; 
    } 
    
    // Use Fractional Rank formula 
    // fractional_rank = r + (n-1)/2 
    Rank_X(i) = r + (s-1) * 0.5;         
  } 
  
  // Return Rank Vector 
  return Rank_X; 
} 


double SpearmanRankCorr( arma::vec kin, arma::vec kout ){
  arma::uvec sort_kin = sort_index(kin, "ascend");
  arma::uvec sort_kout = sort_index(kout, "ascend");
  
  arma:vec rank_kin = linspace(0, kin.size() - 1, kin.size());
  vec rank_kout = linspace(0, kout.size() - 1, kout.size());
  
  rank_kin(sort_kin) = rank_kin;
  rank_kout(sort_kout) = rank_kout;
  
  arma::vec kin_un = unique(kin);
  arma::vec kout_un = unique(kout);
  
  for( int i = 0; i < kin_un.size(); ++i ){
    arma::uvec id = find(kin == kin_un(i));
    arma::vec temp(id.size());
    temp.fill(mean(rank_kin(id)));
    rank_kin(id) = temp;
  }
  for( int i = 0; i < kout_un.size(); ++i ){
    arma::uvec id = find(kout == kout_un(i));
    arma::vec temp(id.size());
    temp.fill(mean(rank_kout(id)));
    rank_kout(id) = temp;
  }
  arma::vec cor_vec = cor(rank_kin, rank_kout);
  return cor_vec(0);
}


int RedundancyMultiplex( arma::sp_mat &adjTRN, arma::sp_mat &adjPPI, arma::uvec &idshuffle ){
  int count = 0;
  sp_mat::const_iterator it     = adjPPI.begin();
  sp_mat::const_iterator it_end = adjPPI.end();
  for(; it != it_end; ++it)
  {
    if( adjTRN(idshuffle(it.row()), idshuffle(it.col())) == 1 ){
      count += 1;
    }
  }
  return count;
}


//Simulated annealing (SA) to compute of ids for shuffled TRN genes for shuffled multiplexes
/*Input: 
 * adjTRN - TRN adjacency matrix
 * adjPPI - PPI adjacency matrix
 * indegTRN - indegree for TRN
 * outdegTRN - outdegree for TRN
 * degPPI - degree for PPI
 * redundancy - required redundancy in the shuffled multiplex
 * nsample - number of gene ids to shuffle in each iteration of simulated annealing (SA)
 * Temp0 - intitial temperature for SA
 * lambda - rate for SA
 * rin - required degree-degree coupling with TRN indegree in the shuffled multiplex
 * rout - required degree-degree coupling with TRN outdegree in the shuffled multiplex
 * eps1 - allowed margin of error for rin in the shuffled multiplex compared to the required rin
 * eps - allowed margin of error for rout in the shuffled multiplex compared to the required rout
 * eps2 - allowed margin of error for redundancy in the shuffled multiplex compared to the required redundancy
 * imax - maximum number of SA iterations
 * nboot - number of times to repeat the shuffled TRN id calculation
 * nthreads - number of CPU cores for parallelization
 */
/*Output:
 * shuffled TRN gene ids
 */
//[[Rcpp::plugins(openmp)]]
//[[Rcpp::export]]
arma::uvec SAShuffleMultiplex( arma::sp_mat adjTRN, arma::sp_mat adjPPI, arma::vec indegTRN, arma::vec outdegTRN, 
                               arma::vec degPPI, int redundancy, int nsample, double Temp0, double lambda, 
                               double eps, double eps1, double eps2, int imax, double rin, double rout ){
  int NumNodes = adjTRN.n_rows;
  // srand(seed);
  arma::uvec idprop = arma::randperm(NumNodes);
  arma::uvec idcur = idprop;
  double r1 = SpearmanRankCorr(indegTRN(idprop), degPPI);
  double r2 = SpearmanRankCorr(outdegTRN(idprop), degPPI);
  int redTemp = RedundancyMultiplex(adjTRN, adjPPI, idprop);
  
  double fcur, fprop, fcurin, fcurout, fproprin, fproprout, fcuredge, fpropedge;
  fprop = fcur = 10*abs(double(redTemp - redundancy))/(double(redundancy + 1)) + 
    abs(r1 - rin) + abs(r2 - rout);
  fcurin = fproprin = abs(r1 - rin);
  fcurout = fproprout = abs(r2 - rout);
  fcuredge = fpropedge = abs(double(redTemp - redundancy));
  int i = 0;
  int accept = -1;
  double Temp = Temp0;
  while( (fcurin > eps1 || fcurout > eps || fcuredge > eps2 ) && i < imax ){
    // if( (i) - 1000*floor(i/1000) == 0 ){
    //   cout << "SA loop = " << i << " f.cur.kin = " << fcurin << "  f.cur.kout = " << fcurout <<
    //     " f.cur.edge = " << fcuredge << " f.cur = " << fcur << " Temp = " << Temp << endl;
    // }
    Temp = Temp0*exp(-lambda*i);
    arma::uvec idflip = randperm(NumNodes, nsample);
    arma::uvec idrnd = randperm(nsample);
    idprop = idcur;
    idprop(idflip) = idprop(idflip(idrnd));
    
    r1 = SpearmanRankCorr(indegTRN(idprop), degPPI);
    r2 = SpearmanRankCorr(outdegTRN(idprop), degPPI);
    redTemp = RedundancyMultiplex(adjTRN, adjPPI, idprop);
    
    fprop = 10*abs(double(redTemp - redundancy))/(double(redundancy + 1)) + 
      abs(r1 - rin) + abs(r2 - rout);
    fproprin = abs(r1 - rin);
    fproprout = abs(r2 - rout);
    fpropedge = abs(double(redTemp - redundancy));
    
    double deltaf = fcur - fprop;
    if( deltaf >= 0 ){
      fcur = fprop;
      fcurin = fproprin;
      fcurout = fproprout;
      fcuredge = fpropedge;
      idcur = idprop;
      accept = 1;
    }else{
      double prob = exp(deltaf/Temp);
      int toss = 0;
      if( prob != 0 && prob != 1 ){
        std::default_random_engine generator;
        std::discrete_distribution<int> distribution {prob, 1 - prob};
        toss = distribution(generator);
      }else{
        toss = prob;
      }
      if( toss == 1 ){
        fcur = fprop;
        fcurin = fproprin;
        fcurout = fproprout;
        fcuredge = fpropedge;
        idcur = idprop;
      }
      accept = toss;
      
    }
    i += 1;
  }
  return idcur;
}


//Parallelize computation of ids for shuffled TRN genes for shuffled multiplexes
/*Input: 
 * adjTRN - TRN adjacency matrix
 * adjPPI - PPI adjacency matrix
 * indegTRN - indegree for TRN
 * outdegTRN - outdegree for TRN
 * degPPI - degree for PPI
 * redundancy - required redundancy in the shuffled multiplex
 * nsample - number of gene ids to shuffle in each iteration of simulated annealing (SA)
 * Temp0 - intitial temperature for SA
 * lambda - rate for SA
 * rin - required degree-degree coupling with TRN indegree in the shuffled multiplex
 * rout - required degree-degree coupling with TRN outdegree in the shuffled multiplex
 * eps1 - allowed margin of error for rin in the shuffled multiplex compared to the required rin
 * eps - allowed margin of error for rout in the shuffled multiplex compared to the required rout
 * eps2 - allowed margin of error for redundancy in the shuffled multiplex compared to the required redundancy
 * imax - maximum number of SA iterations
 * seed - random seed
 * nboot - number of times to repeat the shuffled TRN id calculation
 * nthreads - number of CPU cores for parallelization
 * outfile - location to write the shuffled ids for TRN
 */
/*Output:
 * success - success of the code
 */
//[[Rcpp::plugins(openmp)]]
//[[Rcpp::export]]
int SAShuffleMultiplexParallel( arma::sp_mat adjTRN, arma::sp_mat adjPPI, arma::vec indegTRN, arma::vec outdegTRN, 
                                       arma::vec degPPI, int redundancy, int nsample, double Temp0, double lambda, 
                                       double eps, double eps1, double eps2, int imax, double rin, double rout, int seed, int nboot, 
                                       int nthreads, string outfile ){
  // arma::mat AttackCurves(NumNodes, nboot);
  srand(seed);
  Progress p(nboot, true);
  omp_set_num_threads(nthreads);
  int NumNodes = indegTRN.size();
#pragma omp parallel for
  for( int b = 0; b < nboot; ++b ){
    if ( ! Progress::check_abort() ) {
      arma::uvec idshuffle = SAShuffleMultiplex( adjTRN, adjPPI, indegTRN, outdegTRN, 
                                                 degPPI, redundancy, nsample, Temp0, lambda, 
                                                 eps, eps1, eps2, imax, rin, rout);
      ofstream myfile;
      string writefile;
      writefile.append(outfile);
      writefile.append("/boot_");
      writefile.append(to_string(b + 1));
      myfile.open (writefile);
      for( int v = 0; v < NumNodes - 1; ++v ){
        myfile << idshuffle(v) << "\n";
      }
      myfile << idshuffle(NumNodes - 1);
      myfile.close();
      // int tid = omp_get_thread_num();
      // printf("Welcome to GFG from thread = %d\n",
      //        tid);
      p.increment();
    }
   
  }
  return 1;
}

//[[Rcpp::export]]
arma::uvec SASampleGenerateidprop( arma::uvec idprop, int nsample, int NumNodes ){
  arma::vec temp(NumNodes);
  temp.fill(1);
  for ( int i = 0; i < idprop.size(); ++i ){
    temp(idprop(i)) = 0;
  }
  arma::uvec iddiffNodes = find(temp == 1);
  arma::uvec idSample = randperm(iddiffNodes.size(), nsample);
  
  arma::uvec idflip = randperm(idprop.size(), nsample);
  arma::uvec idprop1 = idprop;
  cout << "Error here 1" << endl;
  idprop1(idflip) = iddiffNodes(idSample);
  cout << "Error here 2" << endl;
  return idprop1;
}

//[[Rcpp::export]]
arma::uvec SASampleGenerateidproptest( arma::uvec &idprop, int nsample, int NumNodes ){
  arma::vec temp(NumNodes);
  temp.fill(1);
  for ( int i = 0; i < idprop.size(); ++i ){
    temp(idprop(i)) = 0;
  }
  arma::uvec iddiffNodes = find(temp == 1);
  arma::uvec idSample = randperm(iddiffNodes.size(), nsample);
  
  arma::uvec idflip = randperm(idprop.size(), nsample);
  idprop(idflip) = iddiffNodes(idSample);
  return idprop;
}

//[[Rcpp::plugins(openmp)]]
//[[Rcpp::export]]
arma::uvec SASampleMultiplex( arma::sp_mat adjTRN, arma::sp_mat adjPPI, arma::vec indegTRN, arma::vec outdegTRN, 
                               arma::vec degPPI, int redundancy, int nsample, double Temp0, double lambda, 
                               double eps, double eps1, double eps2, int imax, double rin, double rout, int NumGenes ){
  int NumNodes = adjTRN.n_rows;
  // srand(seed);
  arma::uvec idprop = arma::randperm(NumNodes, NumGenes);
  arma::uvec idcur = idprop;
  double r1 = SpearmanRankCorr(indegTRN(idprop), degPPI(idprop));
  double r2 = SpearmanRankCorr(outdegTRN(idprop), degPPI(idprop));
  // int redTemp = count_common_edges_among_nodes_complete_iterator(adjTRN, adjPPI, idprop);
  int redTemp = count_red_mult_iterator(adjTRN, adjPPI, idprop);
  
  
  double fcur, fprop, fcurin, fcurout, fproprin, fproprout, fcuredge, fpropedge;
  fprop = fcur = 10*abs(double(redTemp - redundancy))/(double(redundancy + 1)) + 
    abs(r1 - rin) + abs(r2 - rout);
  fcurin = fproprin = abs(r1 - rin);
  fcurout = fproprout = abs(r2 - rout);
  fcuredge = fpropedge = abs(double(redTemp - redundancy));
  int i = 0;
  int accept = -1;
  double Temp = Temp0;
  while( (fcurin > eps1 || fcurout > eps || fcuredge > eps2 ) && i < imax ){
    // if( (i) - 1*floor(i/1) == 0 ){
    //   cout << "SA loop = " << i << " f.cur.kin = " << fcurin << "  f.cur.kout = " << fcurout <<
    //     " f.cur.edge = " << fcuredge << " f.cur = " << fcur << " Temp = " << Temp << endl;
    // }
    Temp = Temp0*exp(-lambda*i);
    idprop = idcur;
    
    // cout << "Reached here, " << idprop.max() << " " << idprop.min() << i << endl;
    // idprop = SASampleGenerateidprop(idcur, nsample, NumNodes);
    arma::vec temp(NumNodes);
    temp.fill(1);
    for ( int j = 0; j < idprop.size(); ++j ){
      temp(idprop(j)) = 0;
    }
    arma::uvec iddiffNodes = find(temp == 1);
    arma::uvec idSample = randperm(iddiffNodes.size(), nsample);
    
    arma::uvec idflip = randperm(idprop.size(), nsample);
    // cout << "Error here 1" << endl;
    // arma::uvec idtemp = idprop(idflip);
    
    idprop(idflip) = iddiffNodes(idSample);
    // for( int j = 0; j < nsample; ++j ){
    //   cout << "Error here 2 " << idprop(idflip(j)) << " " << iddiffNodes(idSample(j)) << endl;
    //   idprop(idflip(j)) = iddiffNodes(idSample(j));
    // }
    // int a = max(idprop);
    // int b = min(idprop);
    // if( a + 1 > NumNodes || b < 0 ){
    //   break;
    // }
    // cout << "Error here 3" << endl;
    // cout << "failed here, " << idprop.max() << " " << idprop.min() << i << endl;
    
    r1 = SpearmanRankCorr(indegTRN(idprop), degPPI(idprop));
    r2 = SpearmanRankCorr(outdegTRN(idprop), degPPI(idprop));
    // redTemp = count_common_edges_among_nodes_complete_iterator(adjTRN, adjPPI, idprop);
    redTemp = count_red_mult_iterator(adjTRN, adjPPI, idprop);
    // cout << "Red = " << redTemp << endl;
    
    
    //Red loop
    
    fprop = 10*abs(double(redTemp - redundancy))/(double(redundancy + 1)) + 
      abs(r1 - rin) + abs(r2 - rout);
    fproprin = abs(r1 - rin);
    fproprout = abs(r2 - rout);
    fpropedge = abs(double(redTemp - redundancy));
    
    double deltaf = fcur - fprop;
    if( deltaf >= 0 ){
      fcur = fprop;
      fcurin = fproprin;
      fcurout = fproprout;
      fcuredge = fpropedge;
      idcur = idprop;
      accept = 1;
    }else{
      double prob = exp(deltaf/Temp);
      int toss = 0;
      if( prob != 0 && prob != 1 ){
        std::default_random_engine generator;
        std::discrete_distribution<int> distribution {prob, 1 - prob};
        toss = distribution(generator);
      }else{
        toss = prob;
      }
      if( toss == 1 ){
        fcur = fprop;
        fcurin = fproprin;
        fcurout = fproprout;
        fcuredge = fpropedge;
        idcur = idprop;
      }
      accept = toss;
      
    }
    i += 1;
  }
  return idcur;
}


//[[Rcpp::plugins(openmp)]]
//[[Rcpp::export]]
int SASampleMultiplexParallel( arma::sp_mat adjTRN, arma::sp_mat adjPPI, arma::vec indegTRN, arma::vec outdegTRN, 
                                arma::vec degPPI, int redundancy, int nsample, double Temp0, double lambda, 
                                double eps, double eps1, double eps2, int imax, double rin, double rout, int seed, int nboot, 
                                int nthreads, string outfile, int NumGenes ){
  // arma::mat AttackCurves(NumNodes, nboot);
  srand(seed);
  Progress p(nboot, true);
  omp_set_num_threads(nthreads);
  
#pragma omp parallel for
  for( int b = 0; b < nboot; ++b ){
    if ( ! Progress::check_abort() ) {
      int NumNodes = indegTRN.size();
      arma::uvec idshuffle = SASampleMultiplex( adjTRN, adjPPI, indegTRN, outdegTRN, 
                                                 degPPI, redundancy, nsample, Temp0, lambda, 
                                                 eps, eps1, eps2, imax, rin, rout, NumGenes);
      cout << "success" << idshuffle.size() << endl;
      ofstream myfile;
      string writefile;
      writefile.append(outfile);
      writefile.append("/boot_");
      writefile.append(to_string(b + 1));
      myfile.open (writefile);
      for( int v = 0; v < NumNodes - 1; ++v ){
        myfile << idshuffle(v) << "\n";
      }
      myfile << idshuffle(NumNodes - 1);
      myfile.close();
      // int tid = omp_get_thread_num();
      // printf("Welcome to GFG from thread = %d\n",
      //        tid);
      p.increment();
    }
    
  }
  return 1;
}


//[[Rcpp::plugins(openmp)]]
//[[Rcpp::export]]
arma::uvec SAShuffleMultiplexMCGC( arma::sp_mat adjTRN, arma::sp_mat adjPPI, arma::vec indegTRN, arma::vec outdegTRN, 
                               arma::vec degPPI, int redundancy, int nsample, double Temp0, double lambda, 
                               double eps, double eps1, double eps2, double eps3, int imax, double rin, double rout, 
                               int MCGC_size, arma::mat elTRNorg, arma::mat elPPI){
  int NumNodes = adjTRN.n_rows;
  // srand(seed);
  arma::uvec idprop = arma::randperm(NumNodes);
  arma::uvec idcur = idprop;
  double r1 = SpearmanRankCorr(indegTRN(idprop), degPPI);
  double r2 = SpearmanRankCorr(outdegTRN(idprop), degPPI);
  int redTemp = RedundancyMultiplex(adjTRN, adjPPI, idprop);
  
  //BEGIN:find size of MCGC
  Graph gTRN(NumNodes), gPPI(NumNodes);
  arma::vec TRN_comp(NumNodes), PPI_comp(NumNodes);
  
  for( int i = 0; i < elTRNorg.n_rows; ++i ){
    gTRN.addEdge(elTRNorg(i, 0), elTRNorg(i, 1));
  }
  
  for( int i = 0; i < elPPI.n_rows; ++i ){
    gPPI.addEdge(elPPI(i, 0), elPPI(i, 1));
  }
  
  gTRN.connectedComponents();
  gPPI.connectedComponents();
  
  for( int i = 0; i < NumNodes; ++i ){
    TRN_comp(i) = gTRN.component[i];
    PPI_comp(i) = gPPI.component[i];
  }
  
  MCGCSize(gTRN, gPPI, TRN_comp, PPI_comp, NumNodes);
  int MCGCtemp = sizeTRNPPIreturn(TRN_comp, PPI_comp);
  // cout << "MCGCtemp = " << MCGCtemp << endl;
  //END:find size of MCGC
  
  
  double fcur, fprop, fcurin, fcurout, fproprin, fproprout, fcuredge, fpropedge, fcurMCGC, fpropMCGC;
  fprop = fcur = 10*abs(double(redTemp - redundancy))/(double(redundancy + 1)) + 
    abs(r1 - rin) + abs(r2 - rout) + double(abs(MCGCtemp - MCGC_size))/double(MCGC_size);
  fcurin = fproprin = abs(r1 - rin);
  fcurout = fproprout = abs(r2 - rout);
  fcuredge = fpropedge = abs(double(redTemp - redundancy));
  fcurMCGC = fpropMCGC = double(abs(MCGCtemp - MCGC_size))/double(MCGC_size);
  int i = 0;
  int accept = -1;
  double Temp = Temp0;
  while( (fcurin > eps1 || fcurout > eps || fcuredge > eps2 || fcurMCGC > eps3 ) && i < imax ){
    // if( (i) - 1000*floor(i/1000) == 0 ){
    //   cout << "SA loop = " << i << " f.cur.kin = " << fcurin << "  f.cur.kout = " << fcurout <<
    //     " f.cur.edge = " << fcuredge << " f.cur.MCGC = " << fcurMCGC << " f.cur = " << fcur << " Temp = " << Temp << endl;
    // }
    Temp = Temp0*exp(-lambda*i);
    arma::uvec idflip = randperm(NumNodes, nsample);
    arma::uvec idrnd = randperm(nsample);
    idprop = idcur;
    idprop(idflip) = idprop(idflip(idrnd));
    
    r1 = SpearmanRankCorr(indegTRN(idprop), degPPI);
    r2 = SpearmanRankCorr(outdegTRN(idprop), degPPI);
    redTemp = RedundancyMultiplex(adjTRN, adjPPI, idprop);
    
    //BEGIN:MCGC size for flipped ids
    arma::mat elTRN = elTRNorg;
    arma::uvec idtemp(NumNodes);
    
    arma::uvec idTRN = idprop;
    arma::mat elTRNtemp = elTRN;
    for( int j = 0;j < idTRN.size(); ++j ){
      // cout << "succcess = " << idTRN(j) << endl;
      arma::uvec id1 = find(elTRNtemp.col(0) == idTRN(j));
      if( id1.size() != 0 ){
        for( int k = 0; k < id1.size(); ++k ){
          elTRN(id1(k), 0) = j;
        }
      }
      
      arma::uvec id2 = find(elTRNtemp.col(1) == idTRN(j));
      if( id2.size() != 0 ){
        for( int k = 0; k < id2.size(); ++k ){
          elTRN(id2(k), 1) = j;
        }
      }
    }
    
    Graph gTRN(NumNodes), gPPI(NumNodes);
    arma::vec TRN_comp(NumNodes), PPI_comp(NumNodes);
    
    for( int k = 0; k < elTRN.n_rows; ++k ){
      gTRN.addEdge(elTRN(k, 0), elTRN(k, 1));
    }
    
    for( int k = 0; k < elPPI.n_rows; ++k ){
      gPPI.addEdge(elPPI(k, 0), elPPI(k, 1));
    }
    
    gTRN.connectedComponents();
    gPPI.connectedComponents();
    
    for( int k = 0; k < NumNodes; ++k ){
      TRN_comp(k) = gTRN.component[k];
      PPI_comp(k) = gPPI.component[k];
    }
    
    MCGCSize(gTRN, gPPI, TRN_comp, PPI_comp, NumNodes);
    MCGCtemp = sizeTRNPPIreturn(TRN_comp, PPI_comp);
    
    //END:MCGC size for flipped ids
    
    fprop = 10*abs(double(redTemp - redundancy))/(double(redundancy + 1)) + 
      abs(r1 - rin) + abs(r2 - rout) + double(abs(MCGCtemp - MCGC_size))/double(MCGC_size);
    fproprin = abs(r1 - rin);
    fproprout = abs(r2 - rout);
    fpropedge = abs(double(redTemp - redundancy));
    fcurMCGC = double(abs(MCGCtemp - MCGC_size))/double(MCGC_size);
    
    double deltaf = fcur - fprop;
    if( deltaf >= 0 ){
      fcur = fprop;
      fcurin = fproprin;
      fcurout = fproprout;
      fcuredge = fpropedge;
      fcurMCGC = fpropMCGC;
      idcur = idprop;
      accept = 1;
    }else{
      double prob = exp(deltaf/Temp);
      int toss = 0;
      if( prob != 0 && prob != 1 ){
        std::default_random_engine generator;
        std::discrete_distribution<int> distribution {prob, 1 - prob};
        toss = distribution(generator);
      }else{
        toss = prob;
      }
      if( toss == 1 ){
        fcur = fprop;
        fcurin = fproprin;
        fcurout = fproprout;
        fcuredge = fpropedge;
        fcurMCGC = fpropMCGC;
        idcur = idprop;
      }
      accept = toss;
      
    }
    i += 1;
  }
  return idcur;
}


//[[Rcpp::plugins(openmp)]]
//[[Rcpp::export]]
int SAShuffleMultiplexParallelMCGC( arma::sp_mat adjTRN, arma::sp_mat adjPPI, arma::vec indegTRN, arma::vec outdegTRN, 
                                arma::vec degPPI, int redundancy, int nsample, double Temp0, double lambda, 
                                double eps, double eps1, double eps2, double eps3, int imax, double rin, double rout, 
                                int MCGC_size, arma::mat elTRNorg, arma::mat elPPI, int seed, int nboot, 
                                int nthreads, string outfile ){
  // arma::mat AttackCurves(NumNodes, nboot);
  srand(seed);
  Progress p(nboot, true);
  omp_set_num_threads(nthreads);
  int NumNodes = indegTRN.size();
#pragma omp parallel for
  for( int b = 0; b < nboot; ++b ){
    if ( ! Progress::check_abort() ) {
      arma::uvec idshuffle = SAShuffleMultiplexMCGC( adjTRN, adjPPI, indegTRN, outdegTRN, 
                                                 degPPI, redundancy, nsample, Temp0, lambda, 
                                                 eps, eps1, eps2, eps3, imax, rin, rout, MCGC_size, 
                                                 elTRNorg, elPPI);
      ofstream myfile;
      string writefile;
      writefile.append(outfile);
      writefile.append("/boot_");
      writefile.append(to_string(b + 1));
      myfile.open (writefile);
      for( int v = 0; v < NumNodes - 1; ++v ){
        myfile << idshuffle(v) << "\n";
      }
      myfile << idshuffle(NumNodes - 1);
      myfile.close();
      // int tid = omp_get_thread_num();
      // printf("Welcome to GFG from thread = %d\n",
      //        tid);
      p.increment();
    }
    
  }
  return 1;
}

//[[Rcpp::plugins(openmp)]]
//[[Rcpp::export]]
arma::uvec SASampleMultiplexMaxRed( arma::sp_mat adjTRN, arma::sp_mat adjPPI, arma::vec indegTRN, arma::vec outdegTRN, 
                              arma::vec degPPI, int nsample, double Temp0, double lambda, int imax, int NumGenes ){
  int NumNodes = adjTRN.n_rows;
  // srand(seed);
  arma::uvec idprop = arma::randperm(NumNodes, NumGenes);
  arma::uvec idcur = idprop;
  double r1 = SpearmanRankCorr(indegTRN(idprop), degPPI(idprop));
  double r2 = SpearmanRankCorr(outdegTRN(idprop), degPPI(idprop));
  // int redTemp = count_common_edges_among_nodes_complete_iterator(adjTRN, adjPPI, idprop);
  int redTemp = count_red_mult_iterator(adjTRN, adjPPI, idprop);
  
  
  double fcur, fprop, fcuredge, fpropedge;
  fprop = fcur = abs(double(redTemp));
  fcuredge = fpropedge = abs(double(redTemp));
  int i = 0;
  int accept = -1;
  double Temp = Temp0;
  while( i < imax ){
    // if( (i) - 1*floor(i/1) == 0 ){
    //   cout << "SA loop = " << i << " f.cur.kin = " << fcurin << "  f.cur.kout = " << fcurout <<
    //     " f.cur.edge = " << fcuredge << " f.cur = " << fcur << " Temp = " << Temp << endl;
    // }
    Temp = Temp0*exp(-lambda*i);
    idprop = idcur;
    
    // cout << "Reached here, " << idprop.max() << " " << idprop.min() << i << endl;
    // idprop = SASampleGenerateidprop(idcur, nsample, NumNodes);
    arma::vec temp(NumNodes);
    temp.fill(1);
    for ( int j = 0; j < idprop.size(); ++j ){
      temp(idprop(j)) = 0;
    }
    arma::uvec iddiffNodes = find(temp == 1);
    arma::uvec idSample = randperm(iddiffNodes.size(), nsample);
    
    arma::uvec idflip = randperm(idprop.size(), nsample);
    // cout << "Error here 1" << endl;
    // arma::uvec idtemp = idprop(idflip);
    
    idprop(idflip) = iddiffNodes(idSample);
    // for( int j = 0; j < nsample; ++j ){
    //   cout << "Error here 2 " << idprop(idflip(j)) << " " << iddiffNodes(idSample(j)) << endl;
    //   idprop(idflip(j)) = iddiffNodes(idSample(j));
    // }
    // int a = max(idprop);
    // int b = min(idprop);
    // if( a + 1 > NumNodes || b < 0 ){
    //   break;
    // }
    // cout << "Error here 3" << endl;
    // cout << "failed here, " << idprop.max() << " " << idprop.min() << i << endl;
    
    r1 = SpearmanRankCorr(indegTRN(idprop), degPPI(idprop));
    r2 = SpearmanRankCorr(outdegTRN(idprop), degPPI(idprop));
    // redTemp = count_common_edges_among_nodes_complete_iterator(adjTRN, adjPPI, idprop);
    redTemp = count_red_mult_iterator(adjTRN, adjPPI, idprop);
    // cout << "Red = " << redTemp << endl;
    
    
    //Red loop
    
    fprop = abs(double(redTemp));
    fpropedge = abs(double(redTemp));
    
    double deltaf = - fcur + fprop;
    if( deltaf >= 0 ){
      fcur = fprop;
      fcuredge = fpropedge;
      idcur = idprop;
      accept = 1;
    }else{
      double prob = exp(deltaf/Temp);
      int toss = 0;
      if( prob != 0 && prob != 1 ){
        std::default_random_engine generator;
        std::discrete_distribution<int> distribution {prob, 1 - prob};
        toss = distribution(generator);
      }else{
        toss = prob;
      }
      if( toss == 1 ){
        fcur = fprop;
        fcuredge = fpropedge;
        idcur = idprop;
      }
      accept = toss;
      
    }
    i += 1;
  }
  return idcur;
}
```
