## Supplementary material for "Internetwork connectivity of molecular networks across species of life": Code: utils.cpp

```
#include <RcppArmadillo.h>
// #include <otherpackage.h>
// #include "/igraph/igraph.h"
//[[Rcpp::depends(RcppArmadillo)]]
using namespace Rcpp;
using namespace std;

/*Function to compute edges present in both TRN and PPI and incident on a single gene/protein in the multiplex.*/
//[[Rcpp::export]]
arma::vec count_common_edges_among_nodes(
    arma::sp_mat adj_Prod_sp,
    arma::vec nodes
){
  arma::vec edge_count(nodes.size(), arma::fill::zeros);
  arma::uvec id_nnz = find(adj_Prod_sp);
  for( int i = 0; i < id_nnz.size(); i++ ){
    int id_y = floor((id_nnz(i) + 1)/adj_Prod_sp.n_rows);
    int id_x = adj_Prod_sp.n_rows;
    if( int(id_nnz(i) + 1) - int(id_y*adj_Prod_sp.n_rows) != 0 ){
      id_x = int(id_nnz(i) + 1) - int(id_y*adj_Prod_sp.n_rows);
      id_y += 1;
    }
    for( int j = 0; j < nodes.size(); j++ ){
      if( int(nodes(j)) == int(id_x) || int(nodes(j)) == int(id_y) ){
        edge_count(j) += 1;
      }
    }
  }
  
  return edge_count;
}

//[[Rcpp::export]]
arma::vec count_common_edges_among_nodes_iterator(
    arma::sp_mat adj_Prod_sp,
    arma::vec nodes
){
  arma::vec edge_count(nodes.size(), arma::fill::zeros);
  arma::sp_mat::const_iterator it     = adj_Prod_sp.begin();
  arma::sp_mat::const_iterator it_end = adj_Prod_sp.end();
  for(; it != it_end; ++it){
    for( int j = 0; j < nodes.size(); j++ ){
      if( int(nodes(j)) == int(it.row() + 1) || int(nodes(j)) == int(it.col() + 1) ){
        edge_count(j) += 1;
      }
    }
  }
  return edge_count;
}


/*Function to compute redundancy coupling for a single gene/protein in the multiplex*/
//[[Rcpp::export]]
int count_common_edges_among_nodes_complete(
    arma::sp_mat adj_Prod_sp,
    arma::vec nodes
){
  int edge_count = 0;
  arma::uvec id_nnz = find(adj_Prod_sp);
  for( int i = 0; i < id_nnz.size(); i++ ){
    int id_y = floor((id_nnz(i) + 1)/adj_Prod_sp.n_rows);
    int id_x = adj_Prod_sp.n_rows;
    if( int(id_nnz(i) + 1) - int(id_y*adj_Prod_sp.n_rows) != 0 ){
      id_x = int(id_nnz(i) + 1) - int(id_y*adj_Prod_sp.n_rows);
      id_y += 1;
    }
    
    arma::uvec tmp_x = find(nodes == id_x);
    arma::uvec tmp_y = find(nodes == id_y);
    
    if( tmp_x.size() > 0 || tmp_y.size() > 0){
      edge_count += 1;
    }
  }
  
  return edge_count;
}
```
