## Supplementary material for "Internetwork connectivity of molecular networks across species of life": Latex Files: Mahajan_Dar_2020.bib

@article{barabasi_network_2004,
	title = {Network biology: understanding the cell's functional organization},
	volume = {5},
	shorttitle = {Network biology},
	number = {2},
	journal = {Nature reviews genetics},
	author = {Barabasi, Albert-Laszlo and Oltvai, Zoltan N.},
	year = {2004},
	keywords = {Barabasietal2004},
	pages = {101},
	file = {Full Text:C\:\\Users\\tarunm3\\Zotero\\storage\\I87LHZ5G\\Barabasi and Oltvai - 2004 - Network biology understanding the cell's function.pdf:application/pdf;Snapshot:C\:\\Users\\tarunm3\\Zotero\\storage\\LD9U9U3Y\\nrg1272.html:text/html}
}

@article{szklarczyk_string_2014,
	title = {{STRING} v10: protein–protein interaction networks, integrated over the tree of life},
	volume = {43},
	shorttitle = {{STRING} v10},
	number = {D1},
	journal = {Nucleic acids research},
	author = {Szklarczyk, Damian and Franceschini, Andrea and Wyder, Stefan and Forslund, Kristoffer and Heller, Davide and Huerta-Cepas, Jaime and Simonovic, Milan and Roth, Alexander and Santos, Alberto and Tsafou, Kalliopi P.},
	year = {2014},
	keywords = {STRINGPPI2014, Szklarczyketal2014},
	pages = {D447--D452},
	file = {Full Text:C\:\\Users\\tarunm3\\Zotero\\storage\\UXSIM3IS\\2435295.html:text/html;Snapshot:C\:\\Users\\tarunm3\\Zotero\\storage\\WNW4REQG\\2435295.html:text/html}
}

@article{rual_towards_2005,
	title = {Towards a proteome-scale map of the human protein–protein interaction network},
	volume = {437},
	number = {7062},
	journal = {Nature},
	author = {Rual, Jean-François and Venkatesan, Kavitha and Hao, Tong and Hirozane-Kishikawa, Tomoko and Dricot, Amélie and Li, Ning and Berriz, Gabriel F. and Gibbons, Francis D. and Dreze, Matija and Ayivi-Guedehoussou, Nono},
	year = {2005},
	keywords = {Rualetal2005},
	pages = {1173},
	file = {Full Text:C\:\\Users\\tarunm3\\Zotero\\storage\\KX7X5SK8\\sfxuiu.html:text/html;Snapshot:C\:\\Users\\tarunm3\\Zotero\\storage\\5D9TYQNU\\nature04209.html:text/html}
}

@article{stelzl_human_2005,
	title = {A human protein-protein interaction network: a resource for annotating the proteome},
	volume = {122},
	shorttitle = {A human protein-protein interaction network},
	number = {6},
	journal = {Cell},
	author = {Stelzl, Ulrich and Worm, Uwe and Lalowski, Maciej and Haenig, Christian and Brembeck, Felix H. and Goehler, Heike and Stroedicke, Martin and Zenkner, Martina and Schoenherr, Anke and Koeppen, Susanne},
	year = {2005},
	keywords = {Stelzletal2005},
	pages = {957--968},
	file = {Full Text:C\:\\Users\\tarunm3\\Zotero\\storage\\GHZF2TR6\\S0092867405008664.html:text/html}
}

@article{chatr-aryamontri_biogrid_2017,
	title = {The {BioGRID} interaction database: 2017 update},
	volume = {45},
	shorttitle = {The {BioGRID} interaction database},
	number = {D1},
	journal = {Nucleic acids research},
	author = {Chatr-Aryamontri, Andrew and Oughtred, Rose and Boucher, Lorrie and Rust, Jennifer and Chang, Christie and Kolas, Nadine K. and O'Donnell, Lara and Oster, Sara and Theesfeld, Chandra and Sellam, Adnane},
	year = {2017},
	keywords = {Chatr-Aryamontrietal2017},
	pages = {D369--D379},
	file = {Full Text:C\:\\Users\\tarunm3\\Zotero\\storage\\QKYZB2MW\\2681732.html:text/html;Snapshot:C\:\\Users\\tarunm3\\Zotero\\storage\\JWD4KM9J\\2681732.html:text/html}
}

@article{murali_droid_2010,
	title = {{DroID} 2011: a comprehensive, integrated resource for protein, transcription factor, {RNA} and gene interactions for {Drosophila}},
	volume = {39},
	shorttitle = {{DroID} 2011},
	number = {suppl\_1},
	journal = {Nucleic acids research},
	author = {Murali, Thilakam and Pacifico, Svetlana and Yu, Jingkai and Guest, Stephen and Roberts, George G. and Finley, Russell L.},
	year = {2010},
	keywords = {Muralietal2011},
	pages = {D736--D743},
	file = {Full Text:C\:\\Users\\tarunm3\\Zotero\\storage\\CNAB3AES\\2506628.html:text/html;Snapshot:C\:\\Users\\tarunm3\\Zotero\\storage\\ZV98JWUV\\2506628.html:text/html}
}

@article{yook_functional_2004,
	title = {Functional and topological characterization of protein interaction networks},
	volume = {4},
	number = {4},
	journal = {Proteomics},
	author = {Yook, Soon-Hyung and Oltvai, Zoltán N. and Barabási, Albert-László},
	year = {2004},
	keywords = {Yooketal2004},
	pages = {928--942},
	file = {Snapshot:C\:\\Users\\tarunm3\\Zotero\\storage\\UA938HHQ\\pmic.html:text/html}
}

@article{yu_high-quality_2008,
	title = {High-quality binary protein interaction map of the yeast interactome network},
	volume = {322},
	number = {5898},
	journal = {Science},
	author = {Yu, Haiyuan and Braun, Pascal and Yıldırım, Muhammed A. and Lemmens, Irma and Venkatesan, Kavitha and Sahalie, Julie and Hirozane-Kishikawa, Tomoko and Gebreab, Fana and Li, Na and Simonis, Nicolas},
	year = {2008},
	keywords = {Yuetal2008},
	pages = {104--110},
	file = {Full Text:C\:\\Users\\tarunm3\\Zotero\\storage\\KKNHJSPG\\Yu et al. - 2008 - High-quality binary protein interaction map of the.pdf:application/pdf;Snapshot:C\:\\Users\\tarunm3\\Zotero\\storage\\JYQNJJH5\\104.html:text/html}
}

@article{wang_global_2010,
	title = {Global protein- protein interaction network in the human pathogen {Mycobacterium} tuberculosis {H37Rv}},
	volume = {9},
	number = {12},
	journal = {Journal of proteome research},
	author = {Wang, Yi and Cui, Tao and Zhang, Cong and Yang, Min and Huang, Yuanxia and Li, Weihui and Zhang, Lei and Gao, Chunhui and He, Yang and Li, Yuqing},
	year = {2010},
	keywords = {Wang\_Cuietal2010},
	pages = {6665--6677},
	file = {Full Text:C\:\\Users\\tarunm3\\Zotero\\storage\\8JRBB57K\\pr100808n.html:text/html;Snapshot:C\:\\Users\\tarunm3\\Zotero\\storage\\FG5SB8VI\\pr100808n.html:text/html}
}

@article{hauser_second-generation_2014,
	title = {A second-generation protein–protein interaction network of helicobacter pylori},
	volume = {13},
	number = {5},
	journal = {Molecular \& cellular proteomics},
	author = {Häuser, Roman and Ceol, Arnaud and Rajagopala, Seesandra V. and Mosca, Roberto and Siszler, Gabriella and Wermke, Nadja and Sikorski, Patricia and Schwarz, Frank and Schick, Matthias and Wuchty, Stefan},
	year = {2014},
	keywords = {Hauseretal2014},
	pages = {1318--1329},
	file = {Full Text:C\:\\Users\\tarunm3\\Zotero\\storage\\WUY6P7PE\\Häuser et al. - 2014 - A second-generation protein–protein interaction ne.pdf:application/pdf;Snapshot:C\:\\Users\\tarunm3\\Zotero\\storage\\HYN67UM8\\1318.html:text/html}
}

@article{rajagopala_binary_2014,
	title = {The binary protein-protein interaction landscape of {Escherichia} coli},
	volume = {32},
	number = {3},
	journal = {Nature biotechnology},
	author = {Rajagopala, Seesandra V. and Sikorski, Patricia and Kumar, Ashwani and Mosca, Roberto and Vlasblom, James and Arnold, Roland and Franca-Koh, Jonathan and Pakala, Suman B. and Phanse, Sadhna and Ceol, Arnaud},
	year = {2014},
	keywords = {Rajagopalaetal2014},
	pages = {285},
	file = {Full Text:C\:\\Users\\tarunm3\\Zotero\\storage\\B6VG8YQP\\PMC4123855.html:text/html;Snapshot:C\:\\Users\\tarunm3\\Zotero\\storage\\ANJHQ3BM\\nbt.html:text/html}
}

@article{blais_constructing_2005,
	title = {Constructing transcriptional regulatory networks},
	volume = {19},
	number = {13},
	journal = {Genes \& development},
	author = {Blais, Alexandre and Dynlacht, Brian David},
	year = {2005},
	keywords = {Blaisetal2005},
	pages = {1499--1511},
	file = {Full Text:C\:\\Users\\tarunm3\\Zotero\\storage\\J24P7X6N\\Blais and Dynlacht - 2005 - Constructing transcriptional regulatory networks.pdf:application/pdf;Snapshot:C\:\\Users\\tarunm3\\Zotero\\storage\\K88VPC7S\\1499.html:text/html}
}

@article{guelzim_topological_2002,
	title = {Topological and causal structure of the yeast transcriptional regulatory network},
	volume = {31},
	number = {1},
	journal = {Nature genetics},
	author = {Guelzim, Nabil and Bottani, Samuele and Bourgine, Paul and Képès, François},
	year = {2002},
	keywords = {Guelzimetal2002},
	pages = {60},
	file = {Full Text:C\:\\Users\\tarunm3\\Zotero\\storage\\CQAXLX2R\\Guelzim et al. - 2002 - Topological and causal structure of the yeast tran.pdf:application/pdf;Snapshot:C\:\\Users\\tarunm3\\Zotero\\storage\\KVJ5EZFJ\\ng873z.html:text/html}
}

@article{lee_transcriptional_2002,
	title = {Transcriptional regulatory networks in {Saccharomyces} cerevisiae},
	volume = {298},
	number = {5594},
	journal = {science},
	author = {Lee, Tong Ihn and Rinaldi, Nicola J. and Robert, François and Odom, Duncan T. and Bar-Joseph, Ziv and Gerber, Georg K. and Hannett, Nancy M. and Harbison, Christopher T. and Thompson, Craig M. and Simon, Itamar},
	year = {2002},
	keywords = {Leeetal2002},
	pages = {799--804},
	file = {Full Text:C\:\\Users\\tarunm3\\Zotero\\storage\\LIH6X82Y\\Lee et al. - 2002 - Transcriptional regulatory networks in Saccharomyc.pdf:application/pdf;Snapshot:C\:\\Users\\tarunm3\\Zotero\\storage\\PIVJU8LA\\799.html:text/html}
}

@article{abdulrehman_yeastract:_2010,
	title = {{YEASTRACT}: providing a programmatic access to curated transcriptional regulatory associations in {Saccharomyces} cerevisiae through a web services interface},
	volume = {39},
	shorttitle = {{YEASTRACT}},
	number = {suppl\_1},
	journal = {Nucleic acids research},
	author = {Abdulrehman, Dário and Monteiro, Pedro Tiago and Teixeira, Miguel Cacho and Mira, Nuno Pereira and Lourenco, Artur Bastos and dos Santos, Sandra Costa and Cabrito, Tânia Rodrigues and Francisco, Alexandre Paulo and Madeira, Sara Cordeiro and Aires, Ricardo Santos},
	year = {2010},
	keywords = {Abdulrehmanetal2010},
	pages = {D136--D140},
	file = {Full Text:C\:\\Users\\tarunm3\\Zotero\\storage\\F2X9ZAJ8\\2508954.html:text/html;Snapshot:C\:\\Users\\tarunm3\\Zotero\\storage\\CG2FFWSK\\2508954.html:text/html}
}

@article{shen-orr_network_2002,
	title = {Network motifs in the transcriptional regulation network of {Escherichia} coli},
	volume = {31},
	number = {1},
	journal = {Nature genetics},
	author = {Shen-Orr, Shai S. and Milo, Ron and Mangan, Shmoolik and Alon, Uri},
	year = {2002},
	keywords = {Shen-Orretal2002},
	pages = {64},
	file = {Full Text:C\:\\Users\\tarunm3\\Zotero\\storage\\T6Q45YXM\\Shen-Orr et al. - 2002 - Network motifs in the transcriptional regulation n.pdf:application/pdf;Snapshot:C\:\\Users\\tarunm3\\Zotero\\storage\\VA52UG7A\\ng881z.html:text/html}
}

@article{milo_network_2002,
	title = {Network motifs: simple building blocks of complex networks},
	volume = {298},
	shorttitle = {Network motifs},
	number = {5594},
	journal = {Science},
	author = {Milo, Ron and Shen-Orr, Shai and Itzkovitz, Shalev and Kashtan, Nadav and Chklovskii, Dmitri and Alon, Uri},
	year = {2002},
	keywords = {Miloetal2002},
	pages = {824--827},
	file = {Full Text:C\:\\Users\\tarunm3\\Zotero\\storage\\IZ5EGXA5\\Milo et al. - 2002 - Network motifs simple building blocks of complex .pdf:application/pdf;Snapshot:C\:\\Users\\tarunm3\\Zotero\\storage\\U5MUEP8T\\824.html:text/html}
}

@article{jeong_lethality_2001,
	title = {Lethality and centrality in protein networks},
	volume = {411},
	number = {6833},
	journal = {Nature},
	author = {Jeong, Hawoong and Mason, Sean P. and Barabási, A.-L. and Oltvai, Zoltan N.},
	year = {2001},
	keywords = {Jeongetal2001},
	pages = {41},
	file = {Full Text:C\:\\Users\\tarunm3\\Zotero\\storage\\SEFLMH6A\\Jeong et al. - 2001 - Lethality and centrality in protein networks.pdf:application/pdf;Snapshot:C\:\\Users\\tarunm3\\Zotero\\storage\\T6K96JGE\\35075138.html:text/html}
}

@article{vazquez_global_2003,
	title = {Global protein function prediction from protein-protein interaction networks},
	volume = {21},
	number = {6},
	journal = {Nature biotechnology},
	author = {Vazquez, Alexei and Flammini, Alessandro and Maritan, Amos and Vespignani, Alessandro},
	year = {2003},
	keywords = {Vazquezetal2003},
	pages = {697},
	file = {Full Text:C\:\\Users\\tarunm3\\Zotero\\storage\\ENM2A26U\\Vazquez et al. - 2003 - Global protein function prediction from protein-pr.pdf:application/pdf;Snapshot:C\:\\Users\\tarunm3\\Zotero\\storage\\UL9Z7Q92\\nbt825.html:text/html}
}

@article{han_evidence_2004,
	title = {Evidence for dynamically organized modularity in the yeast protein–protein interaction network},
	volume = {430},
	number = {6995},
	journal = {Nature},
	author = {Han, Jing-Dong J. and Bertin, Nicolas and Hao, Tong and Goldberg, Debra S. and Berriz, Gabriel F. and Zhang, Lan V. and Dupuy, Denis and Walhout, Albertha JM and Cusick, Michael E. and Roth, Frederick P.},
	year = {2004},
	keywords = {Hanetal2004},
	pages = {88},
	file = {Full Text:C\:\\Users\\tarunm3\\Zotero\\storage\\323WFPKC\\Han et al. - 2004 - Evidence for dynamically organized modularity in t.pdf:application/pdf;Snapshot:C\:\\Users\\tarunm3\\Zotero\\storage\\ZX54R7RH\\nature02555.html:text/html}
}

@article{schwikowski_network_2000,
	title = {A network of protein–protein interactions in yeast},
	volume = {18},
	number = {12},
	journal = {Nature biotechnology},
	author = {Schwikowski, Benno and Uetz, Peter and Fields, Stanley},
	year = {2000},
	keywords = {Schwikowskietal2000},
	pages = {1257},
	file = {Full Text:C\:\\Users\\tarunm3\\Zotero\\storage\\3UF6RATX\\Schwikowski et al. - 2000 - A network of protein–protein interactions in yeast.pdf:application/pdf;Snapshot:C\:\\Users\\tarunm3\\Zotero\\storage\\4UY4MHP7\\nbt1200_1257.html:text/html}
}

@article{reece-hoyes_compendium_2005,
	title = {A compendium of {Caenorhabditis} elegans regulatory transcription factors: a resource for mapping transcription regulatory networks},
	volume = {6},
	shorttitle = {A compendium of {Caenorhabditis} elegans regulatory transcription factors},
	number = {13},
	journal = {Genome biology},
	author = {Reece-Hoyes, John S. and Deplancke, Bart and Shingles, Jane and Grove, Christian A. and Hope, Ian A. and Walhout, Albertha JM},
	year = {2005},
	keywords = {Reece-Hoyesetal2005},
	pages = {R110},
	file = {Full Text:C\:\\Users\\tarunm3\\Zotero\\storage\\7D5K957V\\gb-2005-6-13-r110.html:text/html}
}

@article{deplancke_gene-centered_2006,
	title = {A gene-centered {C}. elegans protein-{DNA} interaction network},
	volume = {125},
	number = {6},
	journal = {Cell},
	author = {Deplancke, Bart and Mukhopadhyay, Arnab and Ao, Wanyuan and Elewa, Ahmed M. and Grove, Christian A. and Martinez, Natalia J. and Sequerra, Reynaldo and Doucette-Stamm, Lynn and Reece-Hoyes, John S. and Hope, Ian A.},
	year = {2006},
	keywords = {Deplanckeetal2006},
	pages = {1193--1205},
	file = {Full Text:C\:\\Users\\tarunm3\\Zotero\\storage\\QU2R92LF\\S0092867406006246.html:text/html}
}

@article{sandmann_core_2007,
	title = {A core transcriptional network for early mesoderm development in {Drosophila} melanogaster},
	volume = {21},
	number = {4},
	journal = {Genes \& development},
	author = {Sandmann, Thomas and Girardot, Charles and Brehme, Marc and Tongprasit, Waraporn and Stolc, Viktor and Furlong, Eileen EM},
	year = {2007},
	keywords = {Sandmannetal2007},
	pages = {436--449},
	file = {Full Text:C\:\\Users\\tarunm3\\Zotero\\storage\\2W73US2F\\Sandmann et al. - 2007 - A core transcriptional network for early mesoderm .pdf:application/pdf;Snapshot:C\:\\Users\\tarunm3\\Zotero\\storage\\QVK5WNHF\\436.html:text/html}
}

@article{jin_arabidopsis_2015,
	title = {An {Arabidopsis} transcriptional regulatory map reveals distinct functional and evolutionary features of novel transcription factors},
	volume = {32},
	number = {7},
	journal = {Molecular biology and evolution},
	author = {Jin, Jinpu and He, Kun and Tang, Xing and Li, Zhe and Lv, Le and Zhao, Yi and Luo, Jingchu and Gao, Ge},
	year = {2015},
	pages = {1767--1773},
	file = {Full Text:C\:\\Users\\tarunm3\\Zotero\\storage\\6MY25ZRE\\1017389.html:text/html;Snapshot:C\:\\Users\\tarunm3\\Zotero\\storage\\T46BW4PJ\\1017389.html:text/html}
}

@article{han_trrust_2017,
	title = {{TRRUST} v2: an expanded reference database of human and mouse transcriptional regulatory interactions},
	volume = {46},
	shorttitle = {{TRRUST} v2},
	number = {D1},
	journal = {Nucleic acids research},
	author = {Han, Heonjong and Cho, Jae-Won and Lee, Sangyoung and Yun, Ayoung and Kim, Hyojin and Bae, Dasom and Yang, Sunmo and Kim, Chan Yeong and Lee, Muyoung and Kim, Eunbeen},
	year = {2017},
	keywords = {Hanetal2017},
	pages = {D380--D386},
	file = {Full Text:C\:\\Users\\tarunm3\\Zotero\\storage\\HD5MZYK7\\4566018.html:text/html;Snapshot:C\:\\Users\\tarunm3\\Zotero\\storage\\BZKMZI44\\4566018.html:text/html}
}

@article{jeong_large-scale_2000,
	title = {The large-scale organization of metabolic networks},
	volume = {407},
	number = {6804},
	journal = {Nature},
	author = {Jeong, Hawoong and Tombor, Bálint and Albert, Réka and Oltvai, Zoltan N. and Barabási, A.-L.},
	year = {2000},
	keywords = {Jeongetal2000},
	pages = {651},
	file = {Full Text:C\:\\Users\\tarunm3\\Zotero\\storage\\YV6QVPAF\\Jeong et al. - 2000 - The large-scale organization of metabolic networks.pdf:application/pdf;Snapshot:C\:\\Users\\tarunm3\\Zotero\\storage\\8IR64QLX\\35036627.html:text/html}
}

@article{guimera_functional_2005,
	title = {Functional cartography of complex metabolic networks},
	volume = {433},
	number = {7028},
	journal = {nature},
	author = {Guimera, Roger and Amaral, Luis A. Nunes},
	year = {2005},
	keywords = {Guimeraetal2005},
	pages = {895},
	file = {Full Text:C\:\\Users\\tarunm3\\Zotero\\storage\\TKD5I65U\\PMC2175124.html:text/html;Snapshot:C\:\\Users\\tarunm3\\Zotero\\storage\\YL4TR9QE\\nature03288.html:text/html}
}

@article{stelling_metabolic_2002,
	title = {Metabolic network structure determines key aspects of functionality and regulation},
	volume = {420},
	number = {6912},
	journal = {Nature},
	author = {Stelling, Jörg and Klamt, Steffen and Bettenbrock, Katja and Schuster, Stefan and Gilles, Ernst Dieter},
	year = {2002},
	keywords = {Stellingetal2002},
	pages = {190},
	file = {Full Text:C\:\\Users\\tarunm3\\Zotero\\storage\\35XKRSPG\\Stelling et al. - 2002 - Metabolic network structure determines key aspects.pdf:application/pdf;Snapshot:C\:\\Users\\tarunm3\\Zotero\\storage\\9JUZMT5G\\nature01166.html:text/html}
}

@article{ravasz_hierarchical_2002,
	title = {Hierarchical organization of modularity in metabolic networks},
	volume = {297},
	number = {5586},
	journal = {science},
	author = {Ravasz, Erzsébet and Somera, Anna Lisa and Mongru, Dale A. and Oltvai, Zoltán N. and Barabási, A.-L.},
	year = {2002},
	keywords = {Ravaszetal2002},
	pages = {1551--1555},
	file = {Full Text:C\:\\Users\\tarunm3\\Zotero\\storage\\UQ7XZQUJ\\Ravasz et al. - 2002 - Hierarchical organization of modularity in metabol.pdf:application/pdf;Snapshot:C\:\\Users\\tarunm3\\Zotero\\storage\\JSIUU9R3\\1551.html:text/html}
}

@article{luscombe_genomic_2004,
	title = {Genomic analysis of regulatory network dynamics reveals large topological changes},
	volume = {431},
	number = {7006},
	journal = {Nature},
	author = {Luscombe, Nicholas M. and Babu, M. Madan and Yu, Haiyuan and Snyder, Michael and Teichmann, Sarah A. and Gerstein, Mark},
	year = {2004},
	keywords = {Luscombeelat2004},
	pages = {308},
	file = {Full Text:C\:\\Users\\tarunm3\\Zotero\\storage\\Q72SWGCY\\Luscombe et al. - 2004 - Genomic analysis of regulatory network dynamics re.pdf:application/pdf;Snapshot:C\:\\Users\\tarunm3\\Zotero\\storage\\DM5VGSKN\\nature02782.html:text/html}
}

@article{babu_evolutionary_2006,
	title = {Evolutionary dynamics of prokaryotic transcriptional regulatory networks},
	volume = {358},
	number = {2},
	journal = {Journal of molecular biology},
	author = {Babu, M. Madan and Teichmann, Sarah A. and Aravind, L.},
	year = {2006},
	keywords = {Babuetal2006},
	pages = {614--633},
	file = {Full Text:C\:\\Users\\tarunm3\\Zotero\\storage\\VR4R24LI\\Babu et al. - 2006 - Evolutionary dynamics of prokaryotic transcription.pdf:application/pdf;Snapshot:C\:\\Users\\tarunm3\\Zotero\\storage\\6AC2L7YQ\\S002228360600204X.html:text/html}
}

@article{wang_its_2010,
	title = {It's the machine that matters: predicting gene function and phenotype from protein networks},
	volume = {73},
	shorttitle = {It's the machine that matters},
	number = {11},
	journal = {Journal of proteomics},
	author = {Wang, Peggy I. and Marcotte, Edward M.},
	year = {2010},
	keywords = {Wangetal2010},
	pages = {2277--2289},
	file = {Full Text:C\:\\Users\\tarunm3\\Zotero\\storage\\A39SNDGA\\PMC2953423.html:text/html;Snapshot:C\:\\Users\\tarunm3\\Zotero\\storage\\9BN5KGXN\\S1874391910002095.html:text/html}
}

@article{goh_human_2007,
	title = {The human disease network},
	volume = {104},
	number = {21},
	journal = {Proceedings of the National Academy of Sciences},
	author = {Goh, Kwang-Il and Cusick, Michael E. and Valle, David and Childs, Barton and Vidal, Marc and Barabási, Albert-László},
	year = {2007},
	keywords = {Gohetal2007},
	pages = {8685--8690},
	file = {Full Text:C\:\\Users\\tarunm3\\Zotero\\storage\\F3NK34AL\\8685.html:text/html;Snapshot:C\:\\Users\\tarunm3\\Zotero\\storage\\P6QD4R3K\\8685.html:text/html}
}

@article{barabasi_network_2011,
	title = {Network medicine: a network-based approach to human disease},
	volume = {12},
	shorttitle = {Network medicine},
	number = {1},
	journal = {Nature reviews genetics},
	author = {Barabási, Albert-László and Gulbahce, Natali and Loscalzo, Joseph},
	year = {2011},
	keywords = {Barabásietal2011},
	pages = {56},
	file = {Full Text:C\:\\Users\\tarunm3\\Zotero\\storage\\TV899KXR\\PMC3140052.html:text/html;Snapshot:C\:\\Users\\tarunm3\\Zotero\\storage\\7ECAHXM7\\nrg2918.html:text/html}
}

@article{zhou_human_2014,
	title = {Human symptoms–disease network},
	volume = {5},
	journal = {Nature communications},
	author = {Zhou, XueZhong and Menche, Jörg and Barabási, Albert-László and Sharma, Amitabh},
	year = {2014},
	keywords = {Zhouetal2014},
	pages = {4212},
	file = {Full Text:C\:\\Users\\tarunm3\\Zotero\\storage\\XGC8J68P\\ncomms5212.html:text/html}
}

@article{ay_drugtarget_2007,
	title = {Drug–target network},
	volume = {25},
	number = {10},
	journal = {Nature biotechnology},
	author = {AY, Muhammed and Goh, Kwang-Il and Cusick, Michael E. and Barabasi, Albert-Laszlo and Vidal, Marc},
	year = {2007},
	keywords = {AYetal2007},
	pages = {1119--1127},
	file = {Full Text:C\:\\Users\\tarunm3\\Zotero\\storage\\3KWNHSEZ\\sfxuiu.html:text/html;Snapshot:C\:\\Users\\tarunm3\\Zotero\\storage\\NG89CKXN\\anonymous.html:text/html}
}

@article{hopkins_network_2008,
	title = {Network pharmacology: the next paradigm in drug discovery},
	volume = {4},
	shorttitle = {Network pharmacology},
	number = {11},
	journal = {Nature chemical biology},
	author = {Hopkins, Andrew L.},
	year = {2008},
	keywords = {Hopkinsetal2008},
	pages = {682},
	file = {Snapshot:C\:\\Users\\tarunm3\\Zotero\\storage\\DQCLZN24\\nchembio.html:text/html}
}

@article{kivela_multilayer_2014,
	title = {Multilayer networks},
	volume = {2},
	number = {3},
	journal = {Journal of complex networks},
	author = {Kivelä, Mikko and Arenas, Alex and Barthelemy, Marc and Gleeson, James P. and Moreno, Yamir and Porter, Mason A.},
	year = {2014},
	pages = {203--271},
	file = {Full Text:C\:\\Users\\tarunm3\\Zotero\\storage\\W7LJ8J5X\\2841130.html:text/html;Snapshot:C\:\\Users\\tarunm3\\Zotero\\storage\\LVJWMU85\\2841130.html:text/html}
}

@article{maniatis_extensive_2002,
	title = {An extensive network of coupling among gene expression machines},
	volume = {416},
	number = {6880},
	journal = {Nature},
	author = {Maniatis, Tom and Reed, Robin},
	year = {2002},
	keywords = {Maniatisetal2002},
	pages = {499},
	file = {Full Text:C\:\\Users\\tarunm3\\Zotero\\storage\\QAZ7XNJA\\Maniatis and Reed - 2002 - An extensive network of coupling among gene expres.pdf:application/pdf;Snapshot:C\:\\Users\\tarunm3\\Zotero\\storage\\NNTZ34PD\\416499a.html:text/html}
}

@article{reis_avoiding_2014,
	title = {Avoiding catastrophic failure in correlated networks of networks},
	volume = {10},
	number = {10},
	journal = {Nature Physics},
	author = {Reis, Saulo DS and Hu, Yanqing and Babino, Andrés and Andrade Jr, José S. and Canals, Santiago and Sigman, Mariano and Makse, Hernán A.},
	year = {2014},
	keywords = {Reisetal2014},
	pages = {762},
	file = {Full Text:C\:\\Users\\tarunm3\\Zotero\\storage\\88ITZ3IN\\Reis et al. - 2014 - Avoiding catastrophic failure in correlated networ.pdf:application/pdf;Snapshot:C\:\\Users\\tarunm3\\Zotero\\storage\\PCWBTLYQ\\nphys3081.html:text/html}
}

@article{klosik_interdependent_2017,
	title = {The interdependent network of gene regulation and metabolism is robust where it needs to be},
	volume = {8},
	number = {1},
	journal = {Nature communications},
	author = {Klosik, David F. and Grimbs, Anne and Bornholdt, Stefan and Hütt, Marc-Thorsten},
	year = {2017},
	keywords = {Klosiketal2017},
	pages = {534},
	file = {Full Text:C\:\\Users\\tarunm3\\Zotero\\storage\\JYGR4QPE\\s41467-017-00587-4.html:text/html}
}

@article{nicosia_measuring_2015,
	title = {Measuring and modeling correlations in multiplex networks},
	volume = {92},
	number = {3},
	journal = {Physical Review E},
	author = {Nicosia, Vincenzo and Latora, Vito},
	year = {2015},
	keywords = {Nicosiaetal2015},
	pages = {032805},
	file = {Full Text:C\:\\Users\\tarunm3\\Zotero\\storage\\QR4V2V53\\Nicosia and Latora - 2015 - Measuring and modeling correlations in multiplex n.pdf:application/pdf;Snapshot:C\:\\Users\\tarunm3\\Zotero\\storage\\5AFVWYT3\\PhysRevE.92.html:text/html}
}

@article{de_domenico_structural_2015,
	title = {Structural reducibility of multilayer networks},
	volume = {6},
	journal = {Nature communications},
	author = {De Domenico, Manlio and Nicosia, Vincenzo and Arenas, Alexandre and Latora, Vito},
	year = {2015},
	keywords = {De Domenicoetal2015},
	pages = {6864},
	file = {Full Text:C\:\\Users\\tarunm3\\Zotero\\storage\\25QRBVX2\\ncomms7864.html:text/html}
}

@article{yeger-lotem_network_2004,
	title = {Network motifs in integrated cellular networks of transcription–regulation and protein–protein interaction},
	volume = {101},
	number = {16},
	journal = {Proceedings of the National Academy of Sciences},
	author = {Yeger-Lotem, Esti and Sattath, Shmuel and Kashtan, Nadav and Itzkovitz, Shalev and Milo, Ron and Pinter, Ron Y. and Alon, Uri and Margalit, Hanah},
	year = {2004},
	keywords = {Yeger-Lotemetal2004},
	pages = {5934--5939},
	file = {Full Text:C\:\\Users\\tarunm3\\Zotero\\storage\\9VZLFQJR\\Yeger-Lotem et al. - 2004 - Network motifs in integrated cellular networks of .pdf:application/pdf;Snapshot:C\:\\Users\\tarunm3\\Zotero\\storage\\5WMCKESK\\5934.html:text/html}
}

@article{chen_integration_2008,
	title = {Integration of external signaling pathways with the core transcriptional network in embryonic stem cells},
	volume = {133},
	number = {6},
	journal = {Cell},
	author = {Chen, Xi and Xu, Han and Yuan, Ping and Fang, Fang and Huss, Mikael and Vega, Vinsensius B. and Wong, Eleanor and Orlov, Yuriy L. and Zhang, Weiwei and Jiang, Jianming},
	year = {2008},
	keywords = {Chenetal2008},
	pages = {1106--1117},
	file = {Full Text:C\:\\Users\\tarunm3\\Zotero\\storage\\852AGUN9\\S009286740800617X.html:text/html}
}

@article{ames_modular_2013,
	title = {Modular biological function is most effectively captured by combining molecular interaction data types},
	volume = {8},
	number = {5},
	journal = {PloS one},
	author = {Ames, Ryan M. and MacPherson, Jamie I. and Pinney, John W. and Lovell, Simon C. and Robertson, David L.},
	year = {2013},
	keywords = {Amesetal2013},
	pages = {e62670},
	file = {Full Text:C\:\\Users\\tarunm3\\Zotero\\storage\\5FQ3YIN7\\article.html:text/html}
}

@article{padi_integrating_2015,
	title = {Integrating transcriptional and protein interaction networks to prioritize condition-specific master regulators},
	volume = {9},
	number = {1},
	journal = {BMC systems biology},
	author = {Padi, Megha and Quackenbush, John},
	year = {2015},
	keywords = {Padietal2015},
	pages = {80},
	file = {Full Text:C\:\\Users\\tarunm3\\Zotero\\storage\\AVI9WZN8\\s12918-015-0228-1.html:text/html}
}

@article{parshani_interdependent_2010,
	title = {Interdependent networks: {Reducing} the coupling strength leads to a change from a first to second order percolation transition},
	volume = {105},
	shorttitle = {Interdependent networks},
	number = {4},
	journal = {Physical review letters},
	author = {Parshani, Roni and Buldyrev, Sergey V. and Havlin, Shlomo},
	year = {2010},
	keywords = {Parshanietal2010},
	pages = {048701},
	file = {Full Text:C\:\\Users\\tarunm3\\Zotero\\storage\\6PQR8YT5\\Parshani et al. - 2010 - Interdependent networks Reducing the coupling str.pdf:application/pdf;Snapshot:C\:\\Users\\tarunm3\\Zotero\\storage\\R6FF5T2H\\PhysRevLett.105.html:text/html}
}

@article{lee_correlated_2012,
	title = {Correlated multiplexity and connectivity of multiplex random networks},
	volume = {14},
	number = {3},
	journal = {New Journal of Physics},
	author = {Lee, Kyu-Min and Kim, Jung Yeol and Cho, Won-kuk and Goh, Kwang-Il and Kim, I. M.},
	year = {2012},
	keywords = {Leeetal2012},
	pages = {033027},
	file = {Full Text:C\:\\Users\\tarunm3\\Zotero\\storage\\4V4AZVVE\\Lee et al. - 2012 - Correlated multiplexity and connectivity of multip.pdf:application/pdf;Snapshot:C\:\\Users\\tarunm3\\Zotero\\storage\\6E4FV335\\meta.html:text/html}
}

@article{parshani_critical_2011,
	title = {Critical effect of dependency groups on the function of networks},
	volume = {108},
	number = {3},
	journal = {Proceedings of the National Academy of Sciences},
	author = {Parshani, Roni and Buldyrev, Sergey V. and Havlin, Shlomo},
	year = {2011},
	keywords = {Parshanietal2011},
	pages = {1007--1010},
	file = {Full Text:C\:\\Users\\tarunm3\\Zotero\\storage\\EXVM856D\\Parshani et al. - 2011 - Critical effect of dependency groups on the functi.pdf:application/pdf;Snapshot:C\:\\Users\\tarunm3\\Zotero\\storage\\ABF28EYC\\1007.html:text/html}
}

@article{son_percolation_2012,
	title = {Percolation theory on interdependent networks based on epidemic spreading},
	volume = {97},
	number = {1},
	journal = {EPL (Europhysics Letters)},
	author = {Son, Seung-Woo and Bizhani, Golnoosh and Christensen, Claire and Grassberger, Peter and Paczuski, Maya},
	year = {2012},
	pages = {16006},
	file = {Full Text:C\:\\Users\\tarunm3\\Zotero\\storage\\ISL765L2\\Son et al. - 2012 - Percolation theory on interdependent networks base.pdf:application/pdf;Snapshot:C\:\\Users\\tarunm3\\Zotero\\storage\\JPXSFS9F\\meta.html:text/html}
}

@article{radicchi_abrupt_2013,
	title = {Abrupt transition in the structural formation of interconnected networks},
	volume = {9},
	number = {11},
	journal = {Nature Physics},
	author = {Radicchi, Filippo and Arenas, Alex},
	year = {2013},
	keywords = {Radicchietal2013},
	pages = {717},
	file = {Full Text:C\:\\Users\\tarunm3\\Zotero\\storage\\2BUCVFHI\\Radicchi and Arenas - 2013 - Abrupt transition in the structural formation of i.pdf:application/pdf;Snapshot:C\:\\Users\\tarunm3\\Zotero\\storage\\EDZH4ERY\\nphys2761.html:text/html}
}

@article{zhou_simultaneous_2014,
	title = {Simultaneous first-and second-order percolation transitions in interdependent networks},
	volume = {90},
	number = {1},
	journal = {Physical Review E},
	author = {Zhou, Dong and Bashan, Amir and Cohen, Reuven and Berezin, Yehiel and Shnerb, Nadav and Havlin, Shlomo},
	year = {2014},
	keywords = {Zhouetal2014},
	pages = {012803},
	file = {Full Text:C\:\\Users\\tarunm3\\Zotero\\storage\\R2PKR29L\\Zhou et al. - 2014 - Simultaneous first-and second-order percolation tr.pdf:application/pdf;Snapshot:C\:\\Users\\tarunm3\\Zotero\\storage\\U2YLM9VR\\PhysRevE.90.html:text/html}
}

@article{radicchi_percolation_2015,
	title = {Percolation in real interdependent networks},
	volume = {11},
	number = {7},
	journal = {Nature Physics},
	author = {Radicchi, Filippo},
	year = {2015},
	keywords = {Radicchietal2015},
	pages = {597},
	file = {Full Text:C\:\\Users\\tarunm3\\Zotero\\storage\\P6MX8KYV\\Radicchi - 2015 - Percolation in real interdependent networks.pdf:application/pdf;Snapshot:C\:\\Users\\tarunm3\\Zotero\\storage\\E5D9PJEG\\nphys3374.html:text/html}
}

@article{gao_networks_2012,
	title = {Networks formed from interdependent networks},
	volume = {8},
	number = {1},
	journal = {Nature physics},
	author = {Gao, Jianxi and Buldyrev, Sergey V. and Stanley, H. Eugene and Havlin, Shlomo},
	year = {2012},
	keywords = {Gaoetal2012},
	pages = {40},
	file = {Snapshot:C\:\\Users\\tarunm3\\Zotero\\storage\\5FHXJ3KU\\nphys2180.html:text/html}
}

@article{dong_percolation_2012,
	title = {Percolation of partially interdependent networks under targeted attack},
	volume = {85},
	number = {1},
	journal = {Physical Review E},
	author = {Dong, Gaogao and Gao, Jianxi and Tian, Lixin and Du, Ruijin and He, Yinghuan},
	year = {2012},
	keywords = {Dongetal2012},
	pages = {016112},
	file = {Full Text:C\:\\Users\\tarunm3\\Zotero\\storage\\TBK8P4C4\\Dong et al. - 2012 - Percolation of partially interdependent networks u.pdf:application/pdf;Snapshot:C\:\\Users\\tarunm3\\Zotero\\storage\\DNL6S5I7\\PhysRevE.85.html:text/html}
}

@article{liu_breakdown_2016,
	title = {Breakdown of interdependent directed networks},
	volume = {113},
	number = {5},
	journal = {Proceedings of the National Academy of Sciences},
	author = {Liu, Xueming and Stanley, H. Eugene and Gao, Jianxi},
	year = {2016},
	keywords = {Liuetal2016},
	pages = {1138--1143},
	file = {Full Text:C\:\\Users\\tarunm3\\Zotero\\storage\\N3HGPBU3\\Liu et al. - 2016 - Breakdown of interdependent directed networks.pdf:application/pdf;Snapshot:C\:\\Users\\tarunm3\\Zotero\\storage\\2C5EKJN5\\1138.html:text/html}
}

@book{hahs-vaughn_introduction_2013,
	title = {An introduction to statistical concepts},
	publisher = {Routledge},
	author = {Hahs-Vaughn, Debbie L. and Lomax, Richard G.},
	year = {2013},
	keywords = {Hahs-Vaughnetal2013},
	file = {Snapshot:C\:\\Users\\tarunm3\\Zotero\\storage\\XPED52CL\\9780203137819.html:text/html}
}

@article{benjamini_controlling_1995,
	title = {Controlling the false discovery rate: a practical and powerful approach to multiple testing},
	volume = {57},
	shorttitle = {Controlling the false discovery rate},
	number = {1},
	journal = {Journal of the Royal statistical society: series B (Methodological)},
	author = {Benjamini, Yoav and Hochberg, Yosef},
	year = {1995},
	keywords = {Benjaminietal1995},
	pages = {289--300},
	file = {Full Text:C\:\\Users\\tarunm3\\Zotero\\storage\\HV7P457X\\Benjamini and Hochberg - 1995 - Controlling the false discovery rate a practical .pdf:application/pdf;Snapshot:C\:\\Users\\tarunm3\\Zotero\\storage\\DQMBVWL9\\j.2517-6161.1995.tb02031.html:text/html}
}

@article{kleineberg_geometric_2017,
	title = {Geometric correlations mitigate the extreme vulnerability of multiplex networks against targeted attacks},
	volume = {118},
	number = {21},
	journal = {Physical review letters},
	author = {Kleineberg, Kaj-Kolja and Buzna, Lubos and Papadopoulos, Fragkiskos and Boguñá, Marián and Serrano, M. Ángeles},
	year = {2017},
	keywords = {Kleinebergetal2017},
	pages = {218301},
	file = {Full Text:C\:\\Users\\tarunm3\\Zotero\\storage\\8GBNQGTA\\Kleineberg et al. - 2017 - Geometric correlations mitigate the extreme vulner.pdf:application/pdf;Snapshot:C\:\\Users\\tarunm3\\Zotero\\storage\\4H5IFP6M\\PhysRevLett.118.html:text/html}
}

@article{buldyrev_catastrophic_2010,
	title = {Catastrophic cascade of failures in interdependent networks},
	volume = {464},
	number = {7291},
	journal = {Nature},
	author = {Buldyrev, Sergey V. and Parshani, Roni and Paul, Gerald and Stanley, H. Eugene and Havlin, Shlomo},
	year = {2010},
	keywords = {Buldyrevetal2010},
	pages = {1025},
	file = {Full Text:C\:\\Users\\tarunm3\\Zotero\\storage\\6E5UYZFQ\\Buldyrev et al. - 2010 - Catastrophic cascade of failures in interdependent.pdf:application/pdf;Snapshot:C\:\\Users\\tarunm3\\Zotero\\storage\\FYIXHNZ4\\nature08932.html:text/html}
}

@article{gemmetto_multiplexity_2015,
	title = {Multiplexity versus correlation: the role of local constraints in real multiplexes},
	volume = {5},
	shorttitle = {Multiplexity versus correlation},
	journal = {Scientific reports},
	author = {Gemmetto, Valerio and Garlaschelli, Diego},
	year = {2015},
	keywords = {Gemmettoetal2015},
	pages = {9120},
	file = {Full Text:C\:\\Users\\tarunm3\\Zotero\\storage\\N78RYJEZ\\srep09120.html:text/html}
}

@article{repana_network_2019,
	title = {The {Network} of {Cancer} {Genes} ({NCG}): a comprehensive catalogue of known and candidate cancer genes from cancer sequencing screens},
	volume = {20},
	shorttitle = {The {Network} of {Cancer} {Genes} ({NCG})},
	number = {1},
	journal = {Genome biology},
	author = {Repana, Dimitra and Nulsen, Joel and Dressler, Lisa and Bortolomeazzi, Michele and Venkata, Santhilata Kuppili and Tourna, Aikaterini and Yakovleva, Anna and Palmieri, Tommaso and Ciccarelli, Francesca D.},
	year = {2019},
	keywords = {Repanaetal2019},
	pages = {1},
	file = {Full Text:C\:\\Users\\tarunm3\\Zotero\\storage\\7EF4WPVI\\s13059-018-1612-0.html:text/html}
}

@article{fields_novel_1989,
	title = {A novel genetic system to detect protein–protein interactions},
	volume = {340},
	number = {6230},
	journal = {Nature},
	author = {Fields, Stanley and Song, Ok-kyu},
	year = {1989},
	keywords = {Fieldsetal1989},
	pages = {245},
	file = {Full Text:C\:\\Users\\tarunm3\\Zotero\\storage\\T4RKQ83N\\sfxuiu.html:text/html;Snapshot:C\:\\Users\\tarunm3\\Zotero\\storage\\IXIULN73\\340245a0.html:text/html}
}

@article{bruckner_yeast_2009,
	title = {Yeast two-hybrid, a powerful tool for systems biology},
	volume = {10},
	number = {6},
	journal = {International journal of molecular sciences},
	author = {Brückner, Anna and Polge, Cécile and Lentze, Nicolas and Auerbach, Daniel and Schlattner, Uwe},
	year = {2009},
	keywords = {Brückneretal2009},
	pages = {2763--2788},
	file = {Full Text:C\:\\Users\\tarunm3\\Zotero\\storage\\MQAGTCPE\\Brückner et al. - 2009 - Yeast two-hybrid, a powerful tool for systems biol.pdf:application/pdf;Snapshot:C\:\\Users\\tarunm3\\Zotero\\storage\\LQZXN27Y\\2763.html:text/html}
}

@article{kirkpatrick_optimization_1983,
	title = {Optimization by simulated annealing},
	volume = {220},
	number = {4598},
	journal = {science},
	author = {Kirkpatrick, Scott and Gelatt, C. Daniel and Vecchi, Mario P.},
	year = {1983},
	keywords = {Kirkpatricketal1983},
	pages = {671--680},
	file = {Full Text:C\:\\Users\\tarunm3\\Zotero\\storage\\M6QPBJ2L\\Kirkpatrick et al. - 1983 - Optimization by simulated annealing.pdf:application/pdf;Snapshot:C\:\\Users\\tarunm3\\Zotero\\storage\\IHJ73SZM\\671.html:text/html}
}

@article{sanz_transcriptional_2011,
	title = {The transcriptional regulatory network of {Mycobacterium} tuberculosis},
	volume = {6},
	number = {7},
	journal = {PloS one},
	author = {Sanz, Joaquín and Navarro, Jorge and Arbués, Ainhoa and Martín, Carlos and Marijuán, Pedro C. and Moreno, Yamir},
	year = {2011},
	keywords = {Sanzetal2011},
	pages = {e22178},
	file = {Full Text:C\:\\Users\\tarunm3\\Zotero\\storage\\46ITM89F\\article.html:text/html}
}

@article{danielli_built_2010,
	title = {Built shallow to maintain homeostasis and persistent infection: insight into the transcriptional regulatory network of the gastric human pathogen {Helicobacter} pylori},
	volume = {6},
	shorttitle = {Built shallow to maintain homeostasis and persistent infection},
	number = {6},
	journal = {PLoS pathogens},
	author = {Danielli, Alberto and Amore, Gabriele and Scarlato, Vincenzo},
	year = {2010},
	keywords = {Daniellietal2010},
	pages = {e1000938},
	file = {Full Text:C\:\\Users\\tarunm3\\Zotero\\storage\\XPRVDACQ\\article.html:text/html}
}

@article{salgado_using_2018,
	title = {Using {RegulonDB}, the {Escherichia} coli {K}-12 {Gene} {Regulatory} {Transcriptional} {Network} {Database}},
	volume = {61},
	number = {1},
	journal = {Current protocols in bioinformatics},
	author = {Salgado, Heladia and Martínez-Flores, Irma and Bustamante, Víctor H. and Alquicira-Hernández, Kevin and García-Sotelo, Jair S. and García-Alonso, Delfino and Collado-Vides, Julio},
	year = {2018},
	keywords = {Salgadoetal2018},
	pages = {1--32},
	file = {Snapshot:C\:\\Users\\tarunm3\\Zotero\\storage\\72UK6U8Y\\cpbi.html:text/html}
}

@article{teixeira_yeastract:_2017,
	title = {{YEASTRACT}: an upgraded database for the analysis of transcription regulatory networks in {Saccharomyces} cerevisiae},
	volume = {46},
	shorttitle = {{YEASTRACT}},
	number = {D1},
	journal = {Nucleic acids research},
	author = {Teixeira, Miguel C. and Monteiro, Pedro T. and Palma, Margarida and Costa, Catarina and Godinho, Cláudia P. and Pais, Pedro and Cavalheiro, Mafalda and Antunes, Miguel and Lemos, Alexandre and Pedreira, Tiago},
	year = {2017},
	pages = {D348--D353},
	file = {Full Text:C\:\\Users\\tarunm3\\Zotero\\storage\\VLKRCP9N\\4191333.html:text/html;Snapshot:C\:\\Users\\tarunm3\\Zotero\\storage\\I8QJ6XFV\\4191333.html:text/html}
}

@article{bass_gene-centered_2016,
	title = {A gene-centered {C}. elegans protein–{DNA} interaction network provides a framework for functional predictions},
	volume = {12},
	number = {10},
	journal = {Molecular systems biology},
	author = {Bass, Juan I. Fuxman and Pons, Carles and Kozlowski, Lucie and Reece-Hoyes, John S. and Shrestha, Shaleen and Holdorf, Amy D. and Mori, Akihiro and Myers, Chad L. and Walhout, Albertha JM},
	year = {2016},
	keywords = {Bassetal2016},
	file = {Snapshot:C\:\\Users\\tarunm3\\Zotero\\storage\\9EV2PAQJ\\msb.html:text/html}
}

@article{giot_protein_2003,
	title = {A protein interaction map of {Drosophila} melanogaster},
	volume = {302},
	number = {5651},
	journal = {science},
	author = {Giot, Loic and Bader, Joel S. and Brouwer, C. and Chaudhuri, Amitabha and Kuang, Bing and Li, Y. and Hao, Y. L. and Ooi, C. E. and Godwin, Brian and Vitols, E.},
	year = {2003},
	keywords = {Giotetal2003},
	pages = {1727--1736},
	file = {Full Text:C\:\\Users\\tarunm3\\Zotero\\storage\\U8ST6QDQ\\Giot et al. - 2003 - A protein interaction map of Drosophila melanogast.pdf:application/pdf;Snapshot:C\:\\Users\\tarunm3\\Zotero\\storage\\NE7TXVEC\\1727.html:text/html}
}

@article{min_network_2014,
	title = {Network robustness of multiplex networks with interlayer degree correlations},
	volume = {89},
	number = {4},
	journal = {Physical Review E},
	author = {Min, Byungjoon and Do Yi, Su and Lee, Kyu-Min and Goh, K.-I.},
	year = {2014},
	keywords = {Minetal2014},
	pages = {042811},
	file = {Full Text:C\:\\Users\\tarunm3\\Zotero\\storage\\UL2T2DKB\\Min et al. - 2014 - Network robustness of multiplex networks with inte.pdf:application/pdf;Snapshot:C\:\\Users\\tarunm3\\Zotero\\storage\\4DDUFRYK\\PhysRevE.89.html:text/html}
}

@article{das_hint:_2012,
	title = {{HINT}: {High}-quality protein interactomes and their applications in understanding human disease},
	volume = {6},
	shorttitle = {{HINT}},
	number = {1},
	journal = {BMC systems biology},
	author = {Das, Jishnu and Yu, Haiyuan},
	year = {2012},
	keywords = {Dasetal2012},
	pages = {92},
	file = {Full Text:C\:\\Users\\tarunm3\\Zotero\\storage\\4QKBTHX6\\1752-0509-6-92.html:text/html}
}

@book{bianconi_multilayer_2018,
	title = {Multilayer {Networks}: {Structure} and {Function}},
	shorttitle = {Multilayer {Networks}},
	publisher = {Oxford university press},
	author = {Bianconi, Ginestra},
	year = {2018},
	keywords = {Bianconietal2018},
	file = {Snapshot:C\:\\Users\\tarunm3\\Zotero\\storage\\QP96TU3A\\books.html:text/html}
}

@article{spearman_proof_1961,
	title = {The proof and measurement of association between two things.},
	author = {Spearman, Charles},
	year = {1961},
	keywords = {Spearmanetal1961},
	file = {Full Text:C\:\\Users\\tarunm3\\Zotero\\storage\\5QUM3DGJ\\books.html:text/html;Snapshot:C\:\\Users\\tarunm3\\Zotero\\storage\\7SXF9XY7\\2006-10257-005.html:text/html}
}

@article{chen_ogee:_2011,
	title = {{OGEE}: an online gene essentiality database},
	volume = {40},
	shorttitle = {{OGEE}},
	number = {D1},
	journal = {Nucleic acids research},
	author = {Chen, Wei-Hua and Minguez, Pablo and Lercher, Martin J. and Bork, Peer},
	year = {2011},
	pages = {D901--D906},
	file = {Full Text:C\:\\Users\\tarunm3\\Zotero\\storage\\CMMT7GUS\\2903696.html:text/html;Snapshot:C\:\\Users\\tarunm3\\Zotero\\storage\\ZWVQ3SUN\\2903696.html:text/html}
}

@article{chen_ogee_2016,
	title = {{OGEE} v2: an update of the online gene essentiality database with special focus on differentially essential genes in human cancer cell lines},
	shorttitle = {{OGEE} v2},
	journal = {Nucleic acids research},
	author = {Chen, Wei-Hua and Lu, Guanting and Chen, Xiao and Zhao, Xing-Ming and Bork, Peer},
	year = {2016},
	pages = {gkw1013},
	file = {Full Text:C\:\\Users\\tarunm3\\Zotero\\storage\\KBRYTLR2\\2333904.html:text/html;Snapshot:C\:\\Users\\tarunm3\\Zotero\\storage\\K6V39RM4\\2333904.html:text/html}
}

@article{fisher_014:_1921,
	title = {014: {On} the" {Probable} {Error}" of a {Coefficient} of {Correlation} {Deduced} from a {Small} {Sample}.},
	shorttitle = {014},
	author = {Fisher, Ronald Aylmer},
	year = {1921},
	keywords = {Fisheretal1921},
	file = {Full Text:C\:\\Users\\tarunm3\\Zotero\\storage\\4YKKWGWI\\Fisher - 1921 - 014 On the Probable Error of a Coefficient of C.pdf:application/pdf}
}

@article{myers_spearman_2004,
	title = {Spearman correlation coefficients, differences between},
	volume = {12},
	journal = {Encyclopedia of statistical sciences},
	author = {Myers, Leann and Sirois, Maria J.},
	year = {2004},
	keywords = {Myersetal2004},
	file = {Snapshot:C\:\\Users\\tarunm3\\Zotero\\storage\\BAKVQI23\\0471667196.ess5050.html:text/html}
}

@article{fu_human_2008,
	title = {Human immunodeficiency virus type 1, human protein interaction database at {NCBI}},
	volume = {37},
	number = {suppl\_1},
	journal = {Nucleic acids research},
	author = {Fu, William and Sanders-Beer, Brigitte E. and Katz, Kenneth S. and Maglott, Donna R. and Pruitt, Kim D. and Ptak, Roger G.},
	year = {2008},
	pages = {D417--D422},
	file = {Full Text:C\:\\Users\\tarunm3\\Zotero\\storage\\6ZRNFBW8\\1006818.html:text/html;Snapshot:C\:\\Users\\tarunm3\\Zotero\\storage\\LHVDRWFS\\1006818.html:text/html}
}

@article{ptak_cataloguing_2008,
	title = {Cataloguing the {HIV} type 1 human protein interaction network},
	volume = {24},
	number = {12},
	journal = {AIDS research and human retroviruses},
	author = {Ptak, Roger G. and Fu, William and Sanders-Beer, Brigitte E. and Dickerson, Jonathan E. and Pinney, John W. and Robertson, David L. and Rozanov, Mikhail N. and Katz, Kenneth S. and Maglott, Donna R. and Pruitt, Kim D.},
	year = {2008},
	keywords = {Ptaketal2008},
	pages = {1497--1502},
	file = {Full Text:C\:\\Users\\tarunm3\\Zotero\\storage\\TDQZEH5R\\Ptak et al. - 2008 - Cataloguing the HIV type 1 human protein interacti.pdf:application/pdf;Snapshot:C\:\\Users\\tarunm3\\Zotero\\storage\\W82ISNXD\\aid.2008.html:text/html}
}

@article{pinney_hivhost_2009,
	title = {{HIV}–host interactions: a map of viral perturbation of the host system},
	volume = {23},
	shorttitle = {{HIV}–host interactions},
	number = {5},
	journal = {Aids},
	author = {Pinney, John W. and Dickerson, Jonathan E. and Fu, William and Sanders-Beer, Brigitte E. and Ptak, Roger G. and Robertson, David L.},
	year = {2009},
	keywords = {Pinneyetal2009},
	pages = {549--554},
	file = {Full Text:C\:\\Users\\tarunm3\\Zotero\\storage\\33FFJANV\\hiv_host_interactions__a_map_of_viral_perturbation.1.html:text/html}
}

@article{cellai_percolation_2013,
	title = {Percolation in multiplex networks with overlap},
	volume = {88},
	number = {5},
	journal = {Physical Review E},
	author = {Cellai, Davide and López, Eduardo and Zhou, Jie and Gleeson, James P. and Bianconi, Ginestra},
	year = {2013},
	keywords = {Cellaietal2013},
	pages = {052811},
	file = {Full Text:C\:\\Users\\tarunm3\\Zotero\\storage\\H3TRP9NB\\Cellai et al. - 2013 - Percolation in multiplex networks with overlap.pdf:application/pdf;Snapshot:C\:\\Users\\tarunm3\\Zotero\\storage\\GRAZ3TFZ\\PhysRevE.88.html:text/html}
}

@article{hamming_error_1950,
	title = {Error detecting and error correcting codes},
	volume = {29},
	number = {2},
	journal = {The Bell system technical journal},
	author = {Hamming, Richard W.},
	year = {1950},
	keywords = {Hammingetal1950},
	pages = {147--160},
	file = {Snapshot:C\:\\Users\\tarunm3\\Zotero\\storage\\N33U6ZE9\\6772729.html:text/html}
}

@book{cohen_statistical_2013,
	title = {Statistical power analysis for the behavioral sciences},
	publisher = {Routledge},
	author = {Cohen, Jacob},
	year = {2013},
	keywords = {Cohenetal2013},
	file = {Full Text:C\:\\Users\\tarunm3\\Zotero\\storage\\QFFJSEMJ\\Cohen - 2013 - Statistical power analysis for the behavioral scie.pdf:application/pdf}
}

@article{cliff_dominance_1993,
	title = {Dominance statistics: {Ordinal} analyses to answer ordinal questions.},
	volume = {114},
	shorttitle = {Dominance statistics},
	number = {3},
	journal = {Psychological bulletin},
	author = {Cliff, Norman},
	year = {1993},
	keywords = {Cliffetal1993},
	pages = {494},
	file = {Full Text:C\:\\Users\\tarunm3\\Zotero\\storage\\FFH5I9RV\\sfxuiu.html:text/html;Snapshot:C\:\\Users\\tarunm3\\Zotero\\storage\\MM5AL3LM\\1994-08169-001.html:text/html}
}

@article{parshani_inter-similarity_2011,
	title = {Inter-similarity between coupled networks},
	volume = {92},
	number = {6},
	journal = {EPL (Europhysics Letters)},
	author = {Parshani, Roni and Rozenblat, Celine and Ietri, Daniele and Ducruet, Cesar and Havlin, Shlomo},
	year = {2011},
	pages = {68002},
	file = {Full Text:C\:\\Users\\tarunm3\\Zotero\\storage\\YX4BEBT5\\Parshani et al. - 2011 - Inter-similarity between coupled networks.pdf:application/pdf;Snapshot:C\:\\Users\\tarunm3\\Zotero\\storage\\2PYP4TJA\\meta.html:text/html}
}

@book{gelman_bayesian_2013,
	title = {Bayesian data analysis},
	publisher = {Chapman and Hall/CRC},
	author = {Gelman, Andrew and Carlin, John B. and Stern, Hal S. and Dunson, David B. and Vehtari, Aki and Rubin, Donald B.},
	year = {2013},
	keywords = {Gelmanetal2013},
	file = {Full Text:C\:\\Users\\tarunm3\\Zotero\\storage\\MFI8Y3BB\\Gelman et al. - 2013 - Bayesian data analysis.pdf:application/pdf;Snapshot:C\:\\Users\\tarunm3\\Zotero\\storage\\DIF7PK8K\\9780429113079.html:text/html}
}

@article{kerman_closed-form_2011,
	title = {A closed-form approximation for the median of the beta distribution},
	journal = {arXiv preprint arXiv:1111.0433},
	author = {Kerman, Jouni},
	year = {2011},
	keywords = {Kermanetal2011},
	file = {Full Text:C\:\\Users\\tarunm3\\Zotero\\storage\\US3VEM9L\\Kerman - 2011 - A closed-form approximation for the median of the .pdf:application/pdf;Snapshot:C\:\\Users\\tarunm3\\Zotero\\storage\\LNGF5YC6\\1111.html:text/html}
}

@article{geman_stochastic_1984,
	title = {Stochastic relaxation, {Gibbs} distributions, and the {Bayesian} restoration of images},
	number = {6},
	journal = {IEEE Transactions on pattern analysis and machine intelligence},
	author = {Geman, Stuart and Geman, Donald},
	year = {1984},
	keywords = {Gemanetal1984},
	pages = {721--741},
	file = {Snapshot:C\:\\Users\\tarunm3\\Zotero\\storage\\44A79U5F\\4767596.html:text/html}
}

@article{hastings_monte_1970,
	title = {Monte {Carlo} sampling methods using {Markov} chains and their applications},
	author = {Hastings, W. Keith},
	year = {1970},
	file = {Full Text:C\:\\Users\\tarunm3\\Zotero\\storage\\KM6NW3NF\\Hastings - 1970 - Monte Carlo sampling methods using Markov chains a.pdf:application/pdf;Snapshot:C\:\\Users\\tarunm3\\Zotero\\storage\\QXYVBEAG\\284580.html:text/html}
}

@article{tanner_calculation_1987,
	title = {The calculation of posterior distributions by data augmentation},
	volume = {82},
	number = {398},
	journal = {Journal of the American statistical Association},
	author = {Tanner, Martin A. and Wong, Wing Hung},
	year = {1987},
	keywords = {Tanneretal1987},
	pages = {528--540},
	file = {Full Text:C\:\\Users\\tarunm3\\Zotero\\storage\\DECGNXM3\\Tanner and Wong - 1987 - The calculation of posterior distributions by data.pdf:application/pdf;Snapshot:C\:\\Users\\tarunm3\\Zotero\\storage\\IN949YXW\\01621459.1987.html:text/html}
}

@article{gelfand_sampling-based_1990,
	title = {Sampling-based approaches to calculating marginal densities},
	volume = {85},
	number = {410},
	journal = {Journal of the American statistical association},
	author = {Gelfand, Alan E. and Smith, Adrian FM},
	year = {1990},
	keywords = {Gelfandetal1990},
	pages = {398--409},
	file = {Full Text:C\:\\Users\\tarunm3\\Zotero\\storage\\4CFZB5DH\\Gelfand and Smith - 1990 - Sampling-based approaches to calculating marginal .pdf:application/pdf;Snapshot:C\:\\Users\\tarunm3\\Zotero\\storage\\JCSR52VW\\01621459.1990.html:text/html}
}

@inproceedings{plummer_jags:_2003,
	title = {{JAGS}: {A} program for analysis of {Bayesian} graphical models using {Gibbs} sampling},
	volume = {124},
	shorttitle = {{JAGS}},
	booktitle = {Proceedings of the 3rd international workshop on distributed statistical computing},
	publisher = {Vienna, Austria.},
	author = {Plummer, Martyn},
	year = {2003},
	keywords = {Plummeretal2003},
	pages = {10},
	file = {Full Text:C\:\\Users\\tarunm3\\Zotero\\storage\\NFLPI2PJ\\Plummer - 2003 - JAGS A program for analysis of Bayesian graphical.pdf:application/pdf}
}

@article{battiston_efficient_2016,
	title = {Efficient exploration of multiplex networks},
	volume = {18},
	number = {4},
	journal = {New Journal of Physics},
	author = {Battiston, Federico and Nicosia, Vincenzo and Latora, Vito},
	year = {2016},
	pages = {043035},
	file = {Full Text:C\:\\Users\\tarunm3\\Zotero\\storage\\Q7Z9HE9K\\Battiston et al. - 2016 - Efficient exploration of multiplex networks.pdf:application/pdf}
}

@article{nie_effect_2015,
	title = {Effect of degree correlation on exact controllability of multiplex networks},
	volume = {436},
	journal = {Physica A: Statistical Mechanics and its Applications},
	author = {Nie, Sen and Wang, Xuwen and Wang, Binghong},
	year = {2015},
	keywords = {Nieetal2015},
	pages = {98--102},
	file = {Full Text:C\:\\Users\\tarunm3\\Zotero\\storage\\FA3Q3L5C\\sfxuiu.html:text/html;Snapshot:C\:\\Users\\tarunm3\\Zotero\\storage\\YXSQVKMW\\S0378437115004537.html:text/html}
}

@article{wagner_proliferation_2014,
	title = {Proliferation of cells with {HIV} integrated into cancer genes contributes to persistent infection},
	volume = {345},
	number = {6196},
	journal = {Science},
	author = {Wagner, Thor A. and McLaughlin, Sherry and Garg, Kavita and Cheung, Charles YK and Larsen, Brendan B. and Styrchak, Sheila and Huang, Hannah C. and Edlefsen, Paul T. and Mullins, James I. and Frenkel, Lisa M.},
	year = {2014},
	keywords = {Wagneretal2014},
	pages = {570--573},
	file = {Full Text:C\:\\Users\\tarunm3\\Zotero\\storage\\9EIBDRWM\\570.html:text/html;Snapshot:C\:\\Users\\tarunm3\\Zotero\\storage\\NN2DRNFW\\570.html:text/html}
}

@article{fromentin_pd-1_2019,
	title = {{PD}-1 blockade potentiates {HIV} latency reversal ex vivo in {CD4}+ {T} cells from {ART}-suppressed individuals},
	volume = {10},
	number = {1},
	journal = {Nature communications},
	author = {Fromentin, Rémi and DaFonseca, Sandrina and Costiniuk, Cecilia T. and El-Far, Mohamed and Procopio, Francesco Andrea and Hecht, Frederick M. and Hoh, Rebecca and Deeks, Steven G. and Hazuda, Daria J. and Lewin, Sharon R.},
	year = {2019},
	keywords = {Fromentinetal2019},
	pages = {814},
	file = {Full Text:C\:\\Users\\tarunm3\\Zotero\\storage\\WTBNF8KQ\\s41467-019-08798-7.html:text/html}
}

@article{albert_scale-free_2005,
	title = {Scale-free networks in cell biology},
	volume = {118},
	number = {21},
	journal = {Journal of cell science},
	author = {Albert, Reka},
	year = {2005},
	keywords = {Albertetal2005},
	pages = {4947--4957},
	file = {Full Text:C\:\\Users\\tarunm3\\Zotero\\storage\\FIZPKHU5\\Albert - 2005 - Scale-free networks in cell biology.pdf:application/pdf;Snapshot:C\:\\Users\\tarunm3\\Zotero\\storage\\BSZ9FN3W\\4947.html:text/html}
}

@article{madan_babu_evolution_2003,
	title = {Evolution of transcription factors and the gene regulatory network in {Escherichia} coli},
	volume = {31},
	number = {4},
	journal = {Nucleic acids research},
	author = {Madan Babu, M. and Teichmann, Sarah A.},
	year = {2003},
	keywords = {Babuetal2003},
	pages = {1234--1244},
	file = {Full Text:C\:\\Users\\tarunm3\\Zotero\\storage\\E2LSVJS7\\2375876.html:text/html;Snapshot:C\:\\Users\\tarunm3\\Zotero\\storage\\YWDTN39M\\2375876.html:text/html}
}

@article{babu_structure_2004,
	title = {Structure and evolution of transcriptional regulatory networks},
	volume = {14},
	number = {3},
	journal = {Current opinion in structural biology},
	author = {Babu, M. Madan and Luscombe, Nicholas M. and Aravind, L. and Gerstein, Mark and Teichmann, Sarah A.},
	year = {2004},
	keywords = {Babuetal2004},
	pages = {283--291},
	file = {Full Text:C\:\\Users\\tarunm3\\Zotero\\storage\\X6M7TQK5\\Babu et al. - 2004 - Structure and evolution of transcriptional regulat.pdf:application/pdf;Snapshot:C\:\\Users\\tarunm3\\Zotero\\storage\\BXM5YJAZ\\S0959440X04000788.html:text/html}
}

@article{faith_large-scale_2007,
	title = {Large-scale mapping and validation of {Escherichia} coli transcriptional regulation from a compendium of expression profiles},
	volume = {5},
	number = {1},
	journal = {PLoS biology},
	author = {Faith, Jeremiah J. and Hayete, Boris and Thaden, Joshua T. and Mogno, Ilaria and Wierzbowski, Jamey and Cottarel, Guillaume and Kasif, Simon and Collins, James J. and Gardner, Timothy S.},
	year = {2007},
	keywords = {Faithetal2007},
	pages = {e8},
	file = {Full Text:C\:\\Users\\tarunm3\\Zotero\\storage\\YYW6DJLA\\article.html:text/html}
}

@article{thieffry_specific_1998,
	title = {From specific gene regulation to genomic networks: a global analysis of transcriptional regulation in {Escherichia} coli},
	volume = {20},
	shorttitle = {From specific gene regulation to genomic networks},
	number = {5},
	journal = {Bioessays},
	author = {Thieffry, Denis and Huerta, Araceli M. and Pérez-Rueda, Ernesto and Collado-Vides, Julio},
	year = {1998},
	keywords = {Thieffryetal1998},
	pages = {433--440},
	file = {Snapshot:C\:\\Users\\tarunm3\\Zotero\\storage\\7VYFEVUM\\(SICI)1521-1878(199805)205433AID-BIES103.0.html:text/html}
}

@article{barabasi_scale-free_2003,
	title = {Scale-free networks},
	volume = {288},
	number = {5},
	journal = {Scientific american},
	author = {Barabási, Albert-László and Bonabeau, Eric},
	year = {2003},
	keywords = {Barabásietal2003},
	pages = {60--69},
	file = {Full Text:C\:\\Users\\tarunm3\\Zotero\\storage\\ANN7YES3\\Barabási and Bonabeau - 2003 - Scale-free networks.pdf:application/pdf;Snapshot:C\:\\Users\\tarunm3\\Zotero\\storage\\3XPNJF2S\\26060284.html:text/html}
}

@article{vazquez_large-scale_2002,
	title = {Large-scale topological and dynamical properties of the {Internet}},
	volume = {65},
	number = {6},
	journal = {Physical Review E},
	author = {Vázquez, Alexei and Pastor-Satorras, Romualdo and Vespignani, Alessandro},
	year = {2002},
	keywords = {Vázquezetal2002},
	pages = {066130},
	file = {Full Text:C\:\\Users\\tarunm3\\Zotero\\storage\\EFVEGFHD\\Vázquez et al. - 2002 - Large-scale topological and dynamical properties o.pdf:application/pdf;Snapshot:C\:\\Users\\tarunm3\\Zotero\\storage\\ZDZK8WLU\\PhysRevE.65.html:text/html}
}

@article{zhou_rich-club_2004,
	title = {The rich-club phenomenon in the {Internet} topology},
	volume = {8},
	number = {3},
	journal = {IEEE Communications Letters},
	author = {Zhou, Shi and Mondragón, Raúl J.},
	year = {2004},
	keywords = {Zhouetal2004},
	pages = {180--182},
	file = {Snapshot:C\:\\Users\\tarunm3\\Zotero\\storage\\D9HBZNBT\\1278314.html:text/html}
}

@article{barabasi_scale-free_2009,
	title = {Scale-free networks: a decade and beyond},
	volume = {325},
	shorttitle = {Scale-free networks},
	number = {5939},
	journal = {science},
	author = {Barabási, Albert-László},
	year = {2009},
	keywords = {Barabásietal2009},
	pages = {412--413},
	file = {Full Text:C\:\\Users\\tarunm3\\Zotero\\storage\\44DGHMQI\\Barabási - 2009 - Scale-free networks a decade and beyond.pdf:application/pdf;Snapshot:C\:\\Users\\tarunm3\\Zotero\\storage\\IXPJIG5D\\412.html:text/html}
}

@article{zheng_control_2019,
	title = {Control of multilayer biological networks and applied to target identification of complex diseases},
	volume = {20},
	number = {1},
	journal = {BMC bioinformatics},
	author = {Zheng, Wei and Wang, Dingjie and Zou, Xiufen},
	year = {2019},
	keywords = {Zhengetal2019},
	pages = {271},
	file = {Full Text:C\:\\Users\\tarunm3\\Zotero\\storage\\6U5B6KJH\\s12859-019-2841-2.html:text/html}
}

@article{vogel_protein_2006,
	title = {Protein family expansions and biological complexity},
	volume = {2},
	number = {5},
	journal = {PLoS computational biology},
	author = {Vogel, Christine and Chothia, Cyrus},
	year = {2006},
	keywords = {Vogeletal2006},
	pages = {e48},
	file = {Full Text:C\:\\Users\\tarunm3\\Zotero\\storage\\ZI7AKPY8\\article.html:text/html}
}

@article{weinberger_stochastic_2005,
	title = {Stochastic gene expression in a lentiviral positive-feedback loop: {HIV}-1 {Tat} fluctuations drive phenotypic diversity},
	volume = {122},
	shorttitle = {Stochastic gene expression in a lentiviral positive-feedback loop},
	number = {2},
	journal = {Cell},
	author = {Weinberger, Leor S. and Burnett, John C. and Toettcher, Jared E. and Arkin, Adam P. and Schaffer, David V.},
	year = {2005},
	pages = {169--182},
	file = {Full Text:C\:\\Users\\tarunm3\\Zotero\\storage\\7F3IGZEW\\S0092867405005490.html:text/html}
}

@article{kaern_stochasticity_2005,
	title = {Stochasticity in gene expression: from theories to phenotypes},
	volume = {6},
	issn = {1471-0056, 1471-0064},
	shorttitle = {Stochasticity in gene expression},
	url = {http://www.nature.com/articles/nrg1615},
	doi = {10.1038/nrg1615},
	abstract = {Genetically identical cells exposed to the same environmental conditions can show significant variation in molecular content and marked differences in phenotypic characteristics. This variability is linked to stochasticity in gene expression, which is generally viewed as having detrimental effects on cellular function with potential implications for disease. However, stochasticity in gene expression can also be advantageous. It can provide the flexibility needed by cells to adapt to fluctuating environments or respond to sudden stresses, and a mechanism by which population heterogeneity can be established during cellular differentiation and development.},
	language = {en},
	number = {6},
	urldate = {2019-12-09},
	journal = {Nature Reviews Genetics},
	author = {Kærn, Mads and Elston, Timothy C. and Blake, William J. and Collins, James J.},
	month = jun,
	year = {2005},
	pages = {451--464},
	file = {Kærn et al. - 2005 - Stochasticity in gene expression from theories to.pdf:C\:\\Users\\tarunm3\\Zotero\\storage\\ZGA7QBEK\\Kærn et al. - 2005 - Stochasticity in gene expression from theories to.pdf:application/pdf}
}

@inproceedings{kumar2010hpidb,
  title={HPIDB-a unified resource for host-pathogen interactions},
  author={Kumar, Ranjit and Nanduri, Bindu},
  booktitle={BMC bioinformatics},
  volume={11},
  number={S6},
  pages={S16},
  year={2010},
  organization={Springer}
}

@article{ammari2016hpidb,
  title={HPIDB 2.0: a curated database for host--pathogen interactions},
  author={Ammari, Mais G and Gresham, Cathy R and McCarthy, Fiona M and Nanduri, Bindu},
  journal={Database},
  volume={2016},
  year={2016},
  publisher={Narnia}
}

@article{zhou2020network,
  title={Network-based drug repurposing for novel coronavirus 2019-nCoV/SARS-CoV-2},
  author={Zhou, Yadi and Hou, Yuan and Shen, Jiayu and Huang, Yin and Martin, William and Cheng, Feixiong},
  journal={Cell discovery},
  volume={6},
  number={1},
  pages={1--18},
  year={2020},
  publisher={Nature Publishing Group}
}

@article{pinero2015disgenet,
  title={DisGeNET: a discovery platform for the dynamical exploration of human diseases and their genes},
  author={Pi{\~n}ero, Janet and Queralt-Rosinach, N{\'u}ria and Bravo, Alex and Deu-Pons, Jordi and Bauer-Mehren, Anna and Baron, Martin and Sanz, Ferran and Furlong, Laura I},
  journal={Database},
  volume={2015},
  year={2015},
  publisher={Narnia}
}

@article{pinero2016disgenet,
  title={DisGeNET: a comprehensive platform integrating information on human disease-associated genes and variants},
  author={Pi{\~n}ero, Janet and Bravo, {\`A}lex and Queralt-Rosinach, N{\'u}ria and Guti{\'e}rrez-Sacrist{\'a}n, Alba and Deu-Pons, Jordi and Centeno, Emilio and Garc{\'\i}a-Garc{\'\i}a, Javier and Sanz, Ferran and Furlong, Laura I},
  journal={Nucleic acids research},
  pages={gkw943},
  year={2016},
  publisher={Oxford University Press}
}

@article{pinero2020disgenet,
  title={The DisGeNET knowledge platform for disease genomics: 2019 update},
  author={Pi{\~n}ero, Janet and Ram{\'\i}rez-Anguita, Juan Manuel and Sa{\"u}ch-Pitarch, Josep and Ronzano, Francesco and Centeno, Emilio and Sanz, Ferran and Furlong, Laura I},
  journal={Nucleic acids research},
  volume={48},
  number={D1},
  pages={D845--D855},
  year={2020},
  publisher={Oxford University Press}
}

@article{sondka2018cosmic,
  title={The COSMIC Cancer Gene Census: describing genetic dysfunction across all human cancers},
  author={Sondka, Zbyslaw and Bamford, Sally and Cole, Charlotte G and Ward, Sari A and Dunham, Ian and Forbes, Simon A},
  journal={Nature Reviews Cancer},
  volume={18},
  number={11},
  pages={696--705},
  year={2018},
  publisher={Nature Publishing Group}
}

@article{liu2019robustness,
  title={Robustness and lethality in multilayer biological molecular networks},
  author={Liu, Xueming and Maiorino, Enrico and Halu, Arda and Loscalzo, Joseph and Gao, Jianxi and Sharma, Amitabh},
  journal={bioRxiv},
  pages={818963},
  year={2019},
  publisher={Cold Spring Harbor Laboratory}
}

@article{maslov2002specificity,
  title={Specificity and stability in topology of protein networks},
  author={Maslov, Sergei and Sneppen, Kim},
  journal={Science},
  volume={296},
  number={5569},
  pages={910--913},
  year={2002},
  publisher={American Association for the Advancement of Science}
}

@article{guney2016network,
  title={Network-based in silico drug efficacy screening},
  author={Guney, Emre and Menche, J{\"o}rg and Vidal, Marc and Bar{\'a}basi, Albert-L{\'a}szl{\'o}},
  journal={Nature communications},
  volume={7},
  number={1},
  pages={1--13},
  year={2016},
  publisher={Nature Publishing Group}
}

@article{menche2015uncovering,
  title={Uncovering disease-disease relationships through the incomplete interactome},
  author={Menche, J{\"o}rg and Sharma, Amitabh and Kitsak, Maksim and Ghiassian, Susan Dina and Vidal, Marc and Loscalzo, Joseph and Barab{\'a}si, Albert-L{\'a}szl{\'o}},
  journal={Science},
  volume={347},
  number={6224},
  year={2015},
  publisher={American Association for the Advancement of Science}
}

@article{repana2019network,
  title={The Network of Cancer Genes (NCG): a comprehensive catalogue of known and candidate cancer genes from cancer sequencing screens},
  author={Repana, Dimitra and Nulsen, Joel and Dressler, Lisa and Bortolomeazzi, Michele and Venkata, Santhilata Kuppili and Tourna, Aikaterini and Yakovleva, Anna and Palmieri, Tommaso and Ciccarelli, Francesca D},
  journal={Genome biology},
  volume={20},
  number={1},
  pages={1},
  year={2019},
  publisher={Springer}
}

@Manual{Rlang,
    title = {R: A Language and Environment for Statistical Computing},
    author = {{R Core Team}},
    organization = {R Foundation for Statistical Computing},
    address = {Vienna, Austria},
    year = {2013},
    url = {http://www.R-project.org/},
  }
  
@book{cohen2013statistical,
  title={Statistical power analysis for the behavioral sciences},
  author={Cohen, Jacob},
  year={2013},
  publisher={Academic press}
}
