## Supplementary material for "Internetwork connectivity of molecular networks across species of life": Latex Files: Mahajan_Dar_2020.tex

%% BioMed_Central_Tex_Template_v1.06
%%                                      %
%  bmc_article.tex            ver: 1.06 %
%                                       %

%%IMPORTANT: do not delete the first line of this template
%%It must be present to enable the BMC Submission system to
%%recognise this template!!

%%%%%%%%%%%%%%%%%%%%%%%%%%%%%%%%%%%%%%%%%
%%                                     %%
%%  LaTeX template for BioMed Central  %%
%%     journal article submissions     %%
%%                                     %%
%%          <8 June 2012>              %%
%%                                     %%
%%                                     %%
%%%%%%%%%%%%%%%%%%%%%%%%%%%%%%%%%%%%%%%%%


%%%%%%%%%%%%%%%%%%%%%%%%%%%%%%%%%%%%%%%%%%%%%%%%%%%%%%%%%%%%%%%%%%%%%
%%                                                                 %%
%% For instructions on how to fill out this Tex template           %%
%% document please refer to Readme.html and the instructions for   %%
%% authors page on the biomed central website                      %%
%% http://www.biomedcentral.com/info/authors/                      %%
%%                                                                 %%
%% Please do not use \input{...} to include other tex files.       %%
%% Submit your LaTeX manuscript as one .tex document.              %%
%%                                                                 %%
%% All additional figures and files should be attached             %%
%% separately and not embedded in the \TeX\ document itself.       %%
%%                                                                 %%
%% BioMed Central currently use the MikTex distribution of         %%
%% TeX for Windows) of TeX and LaTeX.  This is available from      %%
%% http://www.miktex.org                                           %%
%%                                                                 %%
%%%%%%%%%%%%%%%%%%%%%%%%%%%%%%%%%%%%%%%%%%%%%%%%%%%%%%%%%%%%%%%%%%%%%

%%% additional documentclass options:
%  [doublespacing]
%  [linenumbers]   - put the line numbers on margins

%%% loading packages, author definitions
%\documentclass[twocolumn]{bmcart}% uncomment this for twocolumn layout and comment line below
\documentclass{bmcart}

%%% Load packages
\usepackage{amsthm,amsmath}
%\RequirePackage{natbib}
%\RequirePackage[authoryear]{natbib}% uncomment this for author-year bibliography
%\RequirePackage{hyperref}
%\usepackage[utf8]{inputenc} %unicode support
%\usepackage[applemac]{inputenc} %applemac support if unicode package fails
%\usepackage[latin1]{inputenc} %UNIX support if unicode package fails
\usepackage{amsfonts}
\usepackage{graphicx}

%%%%%%%%%%%%%%%%%%%%%%%%%%%%%%%%%%%%%%%%%%%%%%%%%
%%                                             %%
%%  If you wish to display your graphics for   %%
%%  your own use using includegraphic or       %%
%%  includegraphics, then comment out the      %%
%%  following two lines of code.               %%
%%  NB: These line *must* be included when     %%
%%  submitting to BMC.                         %%
%%  All figure files must be submitted as      %%
%%  separate graphics through the BMC          %%
%%  submission process, not included in the    %%
%%  submitted article.                         %%
%%                                             %%
%%%%%%%%%%%%%%%%%%%%%%%%%%%%%%%%%%%%%%%%%%%%%%%%%


%\def\includegraphic{}
%\def\includegraphics{}


%%% Put your definitions there:
\startlocaldefs
\endlocaldefs


%%% Begin ...
\begin{document}

%%% Start of article front matter
\begin{frontmatter}

\begin{fmbox}
\dochead{Research}

%%%%%%%%%%%%%%%%%%%%%%%%%%%%%%%%%%%%%%%%%%%%%%
%%                                          %%
%% Enter the title of your article here     %%
%%                                          %%
%%%%%%%%%%%%%%%%%%%%%%%%%%%%%%%%%%%%%%%%%%%%%%

\title{Internetwork connectivity of molecular networks across species of life}

%%%%%%%%%%%%%%%%%%%%%%%%%%%%%%%%%%%%%%%%%%%%%%
%%                                          %%
%% Enter the authors here                   %%
%%                                          %%
%% Specify information, if available,       %%
%% in the form:                             %%
%%   <key>={<id1>,<id2>}                    %%
%%   <key>=                                 %%
%% Comment or delete the keys which are     %%
%% not used. Repeat \author command as much %%
%% as required.                             %%
%%                                          %%
%%%%%%%%%%%%%%%%%%%%%%%%%%%%%%%%%%%%%%%%%%%%%%

\author[
   addressref={aff1},                   % id's of addresses, e.g. {aff1,aff2}
   %corref={aff1},                       % id of corresponding address, if any
   %noteref={n1},                        % id's of article notes, if any
]{\inits{TM}\fnm{Tarun} \snm{Mahajan}}
\author[
   addressref={aff1,aff2, aff3, aff4},
   corref={aff1},
   %noteref={n2},
]{\inits{RDD}\fnm{Roy D} \snm{Dar}}

%%%%%%%%%%%%%%%%%%%%%%%%%%%%%%%%%%%%%%%%%%%%%%
%%                                          %%
%% Enter the authors' addresses here        %%
%%                                          %%
%% Repeat \address commands as much as      %%
%% required.                                %%
%%                                          %%
%%%%%%%%%%%%%%%%%%%%%%%%%%%%%%%%%%%%%%%%%%%%%%

\address[id=aff1]{%                           % unique id
  \orgname{Department of Bioengineering, University of Illinois at Urbana-Champaign}, % university, etc
  \street{1406 W. Green St},                     %
  \postcode{61801}                                % post or zip code
  \city{Urbana, IL},                              % city
  \cny{USA}                                    % country
}
\address[id=aff2]{%
  \orgname{Department of Electrical and Computer Engineering, University of Illinois at Urbana-Champaign},
  \street{306 North Wright St},
  \postcode{61801}
  \city{Urbana, IL},
  \cny{USA}
}
\address[id=aff3]{%
  \orgname{Center for Biophysics and Quantitative Biology, University of Illinois at Urbana-Champaign},
  \street{1110 West Green Street},
  \postcode{61801}
  \city{Urbana, IL},
  \cny{USA}
}
\address[id=aff4]{%
  \orgname{Carl R. Woese Institute for Genomic Biology, University of Illinois at Urbana-Champaign},
  \street{1206 West Gregory Drive},
  \postcode{61801}
  \city{Urbana, IL},
  \cny{USA}
}
%%%%%%%%%%%%%%%%%%%%%%%%%%%%%%%%%%%%%%%%%%%%%%
%%                                          %%
%% Enter short notes here                   %%
%%                                          %%
%% Short notes will be after addresses      %%
%% on first page.                           %%
%%                                          %%
%%%%%%%%%%%%%%%%%%%%%%%%%%%%%%%%%%%%%%%%%%%%%%

%\begin{artnotes}
%\note{Sample of title note}     % note to the article
%\note[id=n1]{Equal contributor} % note, connected to author
%\end{artnotes}

\end{fmbox}% comment this for two column layout

%%%%%%%%%%%%%%%%%%%%%%%%%%%%%%%%%%%%%%%%%%%%%%
%%                                          %%
%% The Abstract begins here                 %%
%%                                          %%
%% Please refer to the Instructions for     %%
%% authors on http://www.biomedcentral.com  %%
%% and include the section headings         %%
%% accordingly for your article type.       %%
%%                                          %%
%%%%%%%%%%%%%%%%%%%%%%%%%%%%%%%%%%%%%%%%%%%%%%

\begin{abstractbox}

\begin{abstract} % abstract
\parttitle{Background} %if any
Molecular interactions have been studied as independent complex networks in systems biology. However, molecular networks don’t exist independently of each other. In a network of networks approach (called multiplex), we uncover the design principles for the joint organization of transcriptional regulatory network (TRN) and protein-protein interaction (PPI) network.

\parttitle{Results} %if any
We find that TRN and PPI networks are non-randomly coupled in the TRN-PPI multiplex across five different eukaryotic species. Gene degrees in TRN (number of downstream genes) are positively correlated with protein degrees in PPI (number of interacting protein partners). Gene-gene interactions in TRN and protein-protein interactions in PPI also non-randomly overlap in the multiplex. These design principles are conserved across the five eukaryotic species. We show that the robustness of the TRN-PPI multiplex is dependent on these design principles. Further, functionally important genes and proteins, such as essential, disease-related and those involved in host-pathogen PPI networks, are preferentially situated in essential parts of the human multiplex with highly overlapping interactions.

\parttitle{Conclusion}
We unveil the multiplex architecture of TRN and PPI networks across different species. Multiplex architecture may thus define a general framework for studying molecular networks across the different species of life. This approach may uncover the building blocks of the hierarchical organization of molecular interactions.
\end{abstract}

%%%%%%%%%%%%%%%%%%%%%%%%%%%%%%%%%%%%%%%%%%%%%%
%%                                          %%
%% The keywords begin here                  %%
%%                                          %%
%% Put each keyword in separate \kwd{}.     %%
%%                                          %%
%%%%%%%%%%%%%%%%%%%%%%%%%%%%%%%%%%%%%%%%%%%%%%

\begin{keyword}
\kwd{Networks}
\kwd{Gene Regulatory Network}
\kwd{Protein-Protein Interaction}
\kwd{Multiplex}
\kwd{Network of Networks}
\kwd{Network Biology}
\kwd{Host-Pathogen Interaction}
\kwd{Disease-Gene Association}
\end{keyword}

% MSC classifications codes, if any
%\begin{keyword}[class=AMS]
%\kwd[Primary ]{}
%\kwd{}
%\kwd[; secondary ]{}
%\end{keyword}

\end{abstractbox}
%
%\end{fmbox}% uncomment this for twcolumn layout

\end{frontmatter}

%%%%%%%%%%%%%%%%%%%%%%%%%%%%%%%%%%%%%%%%%%%%%%
%%                                          %%
%% The Main Body begins here                %%
%%                                          %%
%% Please refer to the instructions for     %%
%% authors on:                              %%
%% http://www.biomedcentral.com/info/authors%%
%% and include the section headings         %%
%% accordingly for your article type.       %%
%%                                          %%
%% See the Results and Discussion section   %%
%% for details on how to create sub-sections%%
%%                                          %%
%% use \cite{...} to cite references        %%
%%  \cite{koon} and                         %%
%%  \cite{oreg,khar,zvai,xjon,schn,pond}    %%
%%  \nocite{smith,marg,hunn,advi,koha,mouse}%%
%%                                          %%
%%%%%%%%%%%%%%%%%%%%%%%%%%%%%%%%%%%%%%%%%%%%%%

%%%%%%%%%%%%%%%%%%%%%%%%% start of article main body
% <put your article body there>

%%%%%%%%%%%%%%%%
%% Background %%
%%
\section*{Background}
Biological functions and characteristics are consequences of complex interactions between numerous components \cite{barabasi_network_2004}. These components can be molecules such as DNA, RNA, proteins and other small molecules or larger units such as cells, tissues, whole organisms or entire ecosystems. These interactions are organized into a hierarchy of networks. Networks at different levels of this hierarchy have been studied extensively. For instance, at the subcellular level, transcriptional regulatory networks (TRN) model protein-DNA interactions \cite{barabasi_network_2004, abdulrehman_yeastract:_2010, blais_constructing_2005, deplancke_gene-centered_2006, guelzim_topological_2002, han_evidence_2004, jin_arabidopsis_2015, lee_transcriptional_2002, milo_network_2002, reece-hoyes_compendium_2005, sandmann_core_2007, shen-orr_network_2002}, protein-protein interaction (PPI) networks capture physical interactions between proteins \cite{han_evidence_2004, babu_structure_2004, chatr-aryamontri_biogrid_2017, hauser_second-generation_2014, jeong_lethality_2001, murali_droid_2010, rajagopala_binary_2014, rual_towards_2005, schwikowski_network_2000, stelzl_human_2005, szklarczyk_string_2014, vazquez_global_2003, wang_global_2010, yook_functional_2004, yu_high-quality_2008} and metabolic networks map interactions between the set of biochemical reactions in an organism \cite{barabasi_network_2004, guimera_functional_2005, jeong_large-scale_2000, stelling_metabolic_2002}. Analysis of individual network layers has answered important biological questions ranging from organization of gene expression \cite{guelzim_topological_2002, lee_transcriptional_2002, stelling_metabolic_2002, babu_evolutionary_2006, luscombe_genomic_2004}, predicting phenotype from molecular interaction networks \cite{jeong_lethality_2001, wang_global_2010}, to understanding disease biology \cite{ay_drugtarget_2007, barabasi_network_2011, goh_human_2007, hopkins_network_2008, zhou_human_2014}.

However, biological networks do not function in isolation. These networks comprise of different types of interactions and even interact with other networks \cite{barabasi_network_2004, maniatis_extensive_2002}. For instance, TRN and PPI networks interact with each other. Proteins are translated from genes in accordance with the regulatory program encoded in the TRN. These translated proteins interact with each other in the PPI layer. Transcription factor proteins interact with other proteins in the PPI layer and also regulate downstream genes in the TRN network. Further, PPI networks can also encode different kinds of physical interactions between proteins, such as the ones revealed by Yeast Two-Hybrid (Y2H) binary, Affinity Purification (AP) protein complexes, synthetic lethality, dosage lethality, genetic interactions, etc, \cite{chatr-aryamontri_biogrid_2017}. Such multilayer networks (comprising multiple networks) can be interdependent when different network layers interact with each other to form a network of networks (NON) architecture \cite{kivela_multilayer_2014}. For instance, the interaction between TRN and PPI networks forms an interdependent NON (Figure 1). Alternatively, multilayer networks can be multiplex with different networks, which encode distinct types of interactions between the same molecular species such as the different types of PPI interactions.

Until recently, network science has focused largely on the study of individual biological networks. Even some of the studies that worked with multiple networks aggregated or integrated the different networks and did not consider a multilayer approach \cite{ames_modular_2013, chen_integration_2008, padi_integrating_2015, yeger-lotem_network_2004}. This could partly be attributed to the fact that multilayer networks have gained popularity only in recent years, especially in statistical physics \cite{kivela_multilayer_2014}. Now, extensive work has been done to study robustness properties of multilayer networks \cite{buldyrev_catastrophic_2010, dong_percolation_2012, lee_correlated_2012, liu_breakdown_2016, parshani_interdependent_2010, parshani_inter-similarity_2011, radicchi_percolation_2015, radicchi_abrupt_2013, reis_avoiding_2014, son_percolation_2012, zhou_simultaneous_2014}. Counter-intuitively, interdependent networks are more fragile to random failure than independent individual networks \cite{buldyrev_catastrophic_2010}. Real interdependent networks mitigate this vulnerability by means of specific intra- and interlayer degree-degree correlation or coupling \cite{liu_breakdown_2016, reis_avoiding_2014}. For a given TRN-PPI interdependent (or multiplex) network (Figure 1A), degree-degree coupling (\(C_D\)) is quantified as the correlation between the connectivity of a protein in the PPI network, \(K\), and the connectivity of its corresponding gene in the TRN, either in-degree (\(k_{in}\), number of regulations incident on a gene from upstream transcription factors), out-degree (\(k_{out}\), number of downstream genes regulated by a transcription factor), or total degree (\(k = k_{in}+k_{out}\)). In this case, \(C_D\) can be negative, positive, or zero. Particularly, positive \(C_D\) makes the multiplex robust to attack (Figure \ref{fig1}A) \cite{bianconi_multilayer_2018, min_network_2014}. With positive \(C_D\), hub nodes are likely to be hub nodes in all the network layers (Figure \ref{fig1}A, top). For negative \(C_D\), hub nodes in one layers are dependent on spokes in the other network layer. For zero \(C_D\), hubs and spokes are randomly dependent on each other across network layers (Figure \ref{fig1}B, bottom). Here, we investigate \(C_D\) across species and assess whether it is positive, negative, or uncoupled, and if there exists a global trend across various species.

Edge overlap or redundancy between network layers also mitigates vulnerability in interdependent networks \cite{cellai_percolation_2013}. Two genes in the TRN network have an interaction between them if one is the transcription factor for the other. While in PPI networks, interaction between two proteins depicts physical or functional interaction between these proteins. We define multiplex redundancy as similarity in interactions between the TRN and PPI networks. We quantify redundancy (\(C_R\)) by counting the number of common interactions simultaneously in both the TRN and PPI networks. If TRN and PPI networks are represented as graphs, then \(C_R\) can be measured by counting the number of edges which are simultaneously present in both the graphs (Figure \ref{fig1}B and Methods). Higher redundancy makes the multiplex more robust to attack (Figure \ref{fig1}B).

We study the multilayer network of TRN and PPI networks in nine different species, namely \textit{H. pylori}, \textit{M. tuberculosis}, \textit{E. coli}, \textit{S. cerevisiae}, \textit{C. elegans}, \textit{D. melanogaster}, \textit{A. thaliana}, \textit{M. musculus} and \textit{H. sapiens}, spanning two domains of life (bacteria and eukaryotes). We collected TRN networks from nine different sources, and 16 different PPI networks from five different sources (see Methods and Table S1, Additional File 2). Two of the PPI sources are publicly curated databases—BioGRID \cite{chatr-aryamontri_biogrid_2017} and HINT \cite{das_hint:_2012}. Interlayer connectivity is defined by one-to-one correspondence between a gene and its corresponding protein. Therefore, this multilayer network can be reduced to an equivalent multiplex network \cite{buldyrev_catastrophic_2010}. Henceforth, we call the TRN-PPI multilayer network a TRN-PPI multiplex. Based on quality control, five (\textit{S. cerevisiae}, \textit{C. elegans}, \textit{D. melanogaster}, \textit{M. musculus} and \textit{H. sapiens}) of the nine species were used for further analysis; multiplexes with visually continuous percolation curves representing second-order like behavior were studied. Here we show that for species TRN-PPI multiplexes, positive \(C_D\) increases robustness to targeted attack on the genes and proteins. Further, increasing \(C_R\) also increases robustness. We find a trade-off between robustness and independence. Independent multiplexes with no degree-degree coupling and redundancy are highly vulnerable, while positively degree-degree coupled and highly redundant multiplexes are highly robust. We show that this trade-off exists for different species individually. Multiplex coupling is also correlated with the distribution of functionally important genes and proteins such as essential, disease and pathogen-interacting genes and proteins. These functionally important genes are selectively situated in redundant and essential parts of the multiplex and consequently, are vulnerable. Interlayer degree-degree coupling and redundancy offer design mechanisms for tuning robustness in molecular multiplex networks.

\section*{Results}
\subsection*{TRN-PPI multiplex is non-randomly coupled across species}
We study coupling between TRN and PPI networks using two multiplex properties---degree-degree coupling (\(C_D\)) and redundancy coupling (\(C_R\)). \(C_D\) is quantified by Pearson's or Spearman's rank correlation between PPI degree, \(K\), and TRN out-degree, \(k_{out}\) (Figures \ref{fig2}A-\ref{fig2}D, Methods). We quantify \(C_R\) by counting the total number of interactions present simultaneously in both TRN and PPI networks (Figure \ref{fig1}B, Methods). We find that \(C_D\) is significantly positive across different eukaryotes (Figures \ref{fig2}A-\ref{fig2}D and Figure S1 Supplementary Information, Additional File 1). \(C_R\) is also non-randomly positive across different eukaryotes (Figures \ref{fig2}E-\ref{fig2}H and Figure S2 Supplementary Information, Additional File 1). Node-specific (for each gene-protein pair in the multiplex) \(C_R\) values are more long-tailed compared to a randomly shuffled null model. Shuffled null model is generated by randomly shuffling labels on genes in TRN, while keeping protein labels fixed in PPI.

\subsection*{Multiplex degree-degree and redundancy couplings modulate robustness}
We study the relationship between multiplex degree-degree (\(C_D\)) and redundancy (\(C_R\)) couplings  and robustness (\(R\)) across species and for individual species. We use a previously reported formalism to study topological robustness of the TRN-PPI multiplex under targeted attack on its nodes \cite{buldyrev_catastrophic_2010, kleineberg_geometric_2017} (Methods). Robustness is related to the size of the mutually connected giant component (MCGC). MCGC is defined as the largest connected component between both layers of the multiplex (Methods). Buldyrev et al. \cite{buldyrev_catastrophic_2010} and Kleinberg et al. \cite{kleineberg_geometric_2017} track the size of MCGC under attack to quantify robustness. We specifically focus on targeted attack. Under targeted attack, at each step of the attack, gene-protein pairs are removed in decreasing order of  multiplex degree, \(K_{mult}(i)=max(K(i), k_{out}(i))\) \cite{kleineberg_geometric_2017}, where \(K_{mult}(i)\) is the multiplex degree for gene-protein pair \(i\), \(K(i)\) is the degree of protein \(i\) in the PPI network and \(k_{out}(i)\) is the out-degree of gene \(i\) in the TRN network (Methods).  Absolute robustness is then measured by tracking the relative size of MCGC (MCGC divided by number of gene-protein pairs in the multiplex) as we successively remove gene-protein pairs from the multiplex. Figure 3 and Figure S3 (Supplementary Information, Additional File 1) show relative size of MCGC as a function of fraction of gene-protein pairs removed during targeted attack for all the species. We call the curve for MCGC the ‘attack curve’. Absolute robustness is quantified by area under the attack curve (we will call this area \(RobustArea\)) (Methods). Large \(RobustArea\) implies large robustness for a given multiplex, and vice versa small \(RobustArea\) means low robustness. Cohen’s d \cite{cohen2013statistical} is used to quantify effect size for robustness by comparing \(RobustArea\) for a given multiplex against an appropriate null model (Methods). This quantity is used as an estimate of robustness (\(R\)) in this work. We use a Zero-Coupling-Zero-Redundancy null model (see Supplementary Information, Additional File 1). Under this null model, we generate multiplexes with \(C_D\) and \(C_R\) fixed to zero.

Attack curves for different eukaryotic species are shown in Figure \ref{fig3}. Visually, we see that organismal \(RobustArea\) values are larger than the null model on average  (except for yeast with HINT PPI network). This is quantified in Figure \ref{fig4}A, where \(C_D\) and \(R\) are positively correlated across different eukaryotic species with Spearman's correlation coefficient of 0.68 (p-value = 0.044), and \(C_R\) and R are positively correlated across different eukaryotic species with Spearman's correlation coefficient of 0.72 (p-value = 0.037). We also quantify the dependence of R on \(C_D\) and \(C_R\) for individual species (Figures \ref{fig4}B and \ref{fig4}C). For each species, we sample a subset of the TRN-PPI multiplex such that the sampled multiplex has specific values of \(C_D\) and \(C_R\) (Supplementary Information, Additional File 1). This sampling is repeated 1000 times. Attack curves and \(RobustArea\) are then computed for the sampled multiplexes. Therefore, for the given combination of \(C_D\) and \(C_R\), we get a distribution of \(RobustArea\), and \(R\) can be calculated. For each species, we explore \(C_D\) and \(C_R\) values over a grid (Figure \ref{fig4}B). Changing both \(C_D\) and \(C_R\) increases \(R\) (Figures \ref{fig4}B and S6 Supplementary Information Additional File 1). For all the species, multiplexes with highest \(C_D\) and \(C_R\) values exhibit maximum \(R\) (Figures \ref{fig4}B and S6 Supplementary Information, Additional File 1). Further, for a fixed value of \(C_R\), \(R\) increases with \(C_D\). Similarly, for fixed \(C_D\), \(R\) also increases with \(C_R\). The effect of \(C_R\) is stronger than \(C_D\), which is evident for human multiplex where \(R\) is high for high \(C_R\) irrespective of \(C_D\). The independent effect of \(C_D\) and \(C_R\) on \(R\) is shown in Figure \ref{fig4}C.

Across species, we find that \(C_D\), \(C_R\) and \(R\) are positively correlated with the number of gene-proteins in the multiplex (Figure S4, Supplementary Information, Additional File 1). To assess whether across species correlation between \(C_D\) (\(C_R\)) and \(R\) is simply an artifact of the difference between the sizes of the multiplexes or not, we sample two subsets of different sizes for yeast, fly and human (Figures S13-S15, Supplementary Information, Additional File 1). For each of these species, we find that the larger sized susbet has a larger robustness while keeping \(C_R\) and \(C_D\) fixed (Figure S13, Supplementary Information, Additional File 1). However, despite this dependence of \(R\) on multiplex size, dependence on \(C_D\) and \(C_R\) can still be assessed by comparing the two subsets (Figures S13 and S14, Supplementary Information, Additional File 1). This suggests that the correlations with \(R\) seen in Figure \ref{fig4}A and \ref{fig4}C are indeed because of \(C_D\) and \(C_R\) in addition to the dependence on the number of gene-protein pairs in the multiplex.

\subsection*{Essential genes and proteins are essential for multiplex robustness}
We collected essential and non-essential genes for three species (yeast, fly and human) from the Online GEne Essentiality (OGEE) database \cite{chen_ogee_2016, chen_ogee:_2011}. We quantify \(C_R\) and \(C_D\) couplings for essential and non-essential genes (Figures \ref{fig5}A and S7 Supplementary Information, Additional File 1, respectively). For human and fly, essential genes have higher \(C_R\) than non-essential genes (Figure \ref{fig5}A). However, there is no significant difference in \(C_D\) (Figure S5 Supplementary Information, Additional File 1). For yeast, \(C_R\) is not significantly different between essential and non-essential, while \(C_D\) for essential genes is higher than non-essential genes. To gauge importance of essential (non-essential) genes for the multiplex, we selectively attack essential (non-essential) genes and proteins in decreasing order of multiplex degree. Partial attack curves are generated by successively attacking essential (non-essential) genes in decreasing order of multiplex degree for the subset. The attack process is halted once all the genes in the subset of essential (non-essential) genes have been removed (Methods). \(RobustArea\) is quantified by computing area under such partial attack curves. \(R\) is calculated by comparing against a randomly selected set of genes. For fly and human, higher \(C_R\) of essential genes compared to non-essential and random genes co-occurs with lower \(R\) (Figure \ref{fig5}A). For yeast with HINT PPI network, higher \(C_D\) of essential genes compared to non-essential and random genes co-occurs with lower \(R\) (Figure S7A Supplementary Information, Additional File 1). Attacking essential genes breaks the multiplex faster than attacking non-essential or a random set of genes, which shows that essential genes and proteins are situated in a highly coupled and important part of the multiplex. We also study the impact of \(C_R\) on robustness by sampling subsets of genes and proteins (size 100) in the human multiplex (Figure S12 Supplementary Information, Additional File 1). As the redundancy of the sampled genes and proteins increases, robustness decreases against a random set of genes. Thus, redundancy controls selective placement of a subset of genes and proteins in important parts of the multiplex.

\subsection*{Pathogen- and disease-related genes and proteins are situated in essential parts of the human multiplex}
\subsubsection*{Pathogen}
We collected human-pathogen protein-protein interaction data for 13 different pathogens. Data for 12 of these 13 pathogens was collected from a publicly available database, HPIDB 3.0 \cite{kumar2010hpidb, ammari2016hpidb}. This is a curated database which currently contains 69,787 unique protein interactions between 66 host and 668 pathogen species. We studied interactions for 12 different human pathogens from HPIDB 3.0 (Figure \ref{fig5}B). Besides these 12 pathogens, we also included human-pathogen protein interactions for various human coronaviruses (HCoVs). We collected a list of 119 human proteins which interact with various HCoVs \cite{zhou2020network}. Therefore, in total we have 13 pathogens in our analysis.

For each pathogen, we divide the human multiplex into two sets of gene-protein pairs: pathogen-related (\(S_P\)) and not pathogen-related (\(S_{NP}\)). We perform targeted attack on these sets of genes and proteins and generate partial attack curves (Methods and Supplementary Information, Additional File 1). The size of \(S_P\) is greater than \(S_{NP}\) for all of the 13 pathogens. Therefore, for an equitable comparison, we randomly sampled genes and proteins from \(S_{NP}\) to match the number in \(S_{P}\). This process is repeated 100 times. For a baseline comparison, we also randomly sample genes and proteins to match the number in \(S_{NP}\). This random sampling was also performed 100 times. \(R\) is computed for pathogen-related and not pathogen-related genes and proteins by comparing partial \(RobustArea\) against the partial \(RobustArea\) for the random set. For all the pathogens, except Zika, targeted attack on the pathogen-related genes and proteins makes the multiplex highly vulnerable (Figure \ref{fig5}B, left). This vulnerability co-occurs with higher \(C_R\) for the pathogen-related genes and proteins compared to not pathogen-related genes and proteins for all the pathogens, except dengue and influenza with BioGRID PPI (Figure \ref{fig5}B, right). \(C_D\) is similar between essential and non-essential genes (Figure S7B, right Supplementary Information, Additional File 1). Further, given our simulations (Figure S12 Supplementary Information, Additional File 1), this suggests that higher redundancy makes the pathogen-related genes and proteins highly important for the human multiplex. \(R\) and \(C_R\) for \(S_P\) are negatively correlated with Spearman's rank correlation of -0.82 (p-value = \(2.1\times10^{-6}\)), (Figure S8C Supplementary Information, Additional File 1). Even after controlling for the different number of \(S_P\) for different pathogens, \(R\) and \(C_R\) are negatively correlated with Spearman's rank correlation of -0.63 (p-value = \(4.6\times10^{-7}\)), (Figure S8D Supplementary Information, Additional File 1). In agreement with this conclusion, pathogen-related genes and proteins are significantly enriched in the human multiplex (Figure S10A Supplementary Information, Additional File 1).

\subsubsection*{Disease}
We collected disease-related genes from a publicly available database, DisGeNET \cite{pinero2015disgenet, pinero2016disgenet, pinero2020disgenet}. The current version (v6.0) contains gene-disease associations between 17,549 genes and 24,166 diseases, disorders, traits, and clinical or abnormal human phenotypes. We collected disease-gene associations for diseases which have at least 100 genes (with HINT PPI network) in the human multiplex considered in this work. After this filtering, we retain 24 diseases in our analysis (Figure \ref{fig5}C).

For each disease, we divide the human multiplex into two sets of genes and proteins: disease-related (\(S_{D}\)) and not disease-related (\(S_{ND}\)). We perform targeted attack on these sets of genes and proteins and generate partial attack curves (Methods and Supplementary Information, Additional File 1). Size of \(S_{D}\) is greater than \(S_{ND}\) for all the 24 diseases. Therefore, for an equitable comparison, we randomly sampled genes and proteins from \(S_{ND}\) to match the number in \(S_{D}\). This process is repeated 100 times. For a baseline comparison, we also randomly sample genes and proteins to match the number in \(S_{D}\). This random sampling was also performed 100 times. \(RobustArea\) and \(R\) are computed as it was done for pathogens previously (Methods and Supplementary Information, Additional File 1). For all the diseases, except Unipolar Depression (with HINT PPI) and Liver Cirrhosis, Experimental (with BioGRID PPI), targeted attack on the disease-related genes and proteins makes the multiplex highly vulnerable (Figure \ref{fig5}C, left). This vulnerability co-occurs with higher \(C_R\) for the disease-related genes and proteins compared to not disease-related genes and proteins for all the diseases, (Figure \ref{fig5}C, right). \(C_D\) is similar between disease and non-disease genes (Figure S7C, right Supplementary Information, Additional File 1). Further, given our simulations (Figure S12 Supplementary Information, Additional File 1), this suggests that higher redundancy makes the disease-related genes and proteins highly important for the human multiplex. \(R\) and \(C_R\) for \(S_D\) are negatively correlated with Spearman's rank correlation of -0.69 (p-value = \(2.1\times10^{-7}\)), Figure S9C (Supplementary Information, Additional File 1). Even after controlling for the different number of \(S_D\) for different diseases, \(R\) and \(C_R\) are negatively correlated with Spearman's rank correlation of -0.59 (p-value = \(2.2\times10^{-10}\)), Figure S9D (Supplementary Information, Additional File 1). In agreement with this conclusion, disease-related genes and proteins are significantly enriched in the human multiplex (Figure S10B Supplementary Information Additional File 1).

We also collected the set of genes that contain mutations which have been causally implicated in cancer from the Network of Cancer Genes (NCG) database \cite{repana2019network}. NCG also includes information on whether a given cancer gene is an oncogene or a tumor suppressor gene (TSG). For the set of oncogenes (TSGs) in NCG, we divide the human multiplex into two sets of genes and proteins: oncogene (TSG) (\(S_{ON}\) (\(S_{TSG}\))) and not oncogene (TSG) (\(S_{NON}\) (\(S_{NTSG}\))). As before, we compare partial robustness between both the sets. We find that targeted attack on oncogenes (TSGs) makes the multiplex highly vulnerable (Figure \ref{fig5}D, left). This vulnerability co-occurs with higher \(C_R\) for the set of oncogenes (TSGs) (Figure \ref{fig5}D, right). \(C_D\) is similar between oncogenes (TSGs) and non-oncogenes (TSGs) (Figure S7D, right Supplementary Information, Additional File 1).

\section*{Discussion}
Recently, robustness properties of different biological multiplex and multilayer networks have been studied. These include brain networks \cite{reis_avoiding_2014, kleineberg_geometric_2017}, multiplex of PPI interactions \cite{kleineberg_geometric_2017}, and a TRN-metabolic multilayer network \cite{klosik_interdependent_2017}. However, these studies have limitations, which are described next. Kleineberg et al. \cite{kleineberg_geometric_2017} do not draw broader conclusions about the organization of PPI multiplex across different species. Further, the different layers encode different types of interactions between proteins. This framework does not capture interaction between different types of molecules. Their conclusions are based on a generative model of network growth. This makes the results contingent on the accuracy of the generative model. This generative model is based on geometric principles and does not incorporate biological motivations or mechanisms. Klosik et al. \cite{klosik_interdependent_2017} study robustness under random failure of a TRN-metabolic multilayer network. The study only focuses on E. coli and there are no species wide comparisons. Moreover, they do not study the dependence of robustness on degree-degree coupling and redundancy. Another recent study focuses on the interdependent or multiplex network of TRN-PPI-metabolic networks in human \cite{liu2019robustness}. Liu et al. 2019 \cite{liu2019robustness} show that this multiplex is more robust than an uncoupled or shuffled multiplex. They also showed that essential and cancer genes are preferentially arranged in essential parts of the multiplex. However, they do not study the dependence of robustness on multiplex properties. Further, there is no cross-species analysis.

This study bridges the gap between theoretical developments and sub-cellular multilayer networks of molecular interactions. The central goal of our work is to investigate the organization and traits of molecular multiplexes. We focus our attention on the multiplex of TRN and PPI networks across five different eukaryotes. Our analysis spans five different TRN and 9 different PPI networks. We show that degree-degree coupling and redundancy are universal principles that shape robustness of the multiplex. Both are independent modulators of robustness. Though maximum robustness is achieved for a degree-degree coupling of 1 and a completely redundant multiplex, the observed species multiplexes have low absolute degree-degree coupling and redundancy. This suggests that robustness is not the only evolutionary pressure shaping the TRN-PPI multiplex. Independence might be a countering force to robustness.  One possible explanation for low absolute degree-degree coupling, redundancy and hence robustness could be an inability to tune degree-degree coupling and redundancy independently. Redundancy and degree-degree coupling are positively correlated across species (Figure S11 Supplementary Information, Additional File 1). Multiplexes in nature might be tuning degree-degree coupling and redundancy in unison. Therefore, increasing robustness would increase redundancy as well. If redundancy were high, both TRN and PPI layers would be encoding similar interactions and the amount of unique information captured by the multiplex would be low \cite{de_domenico_structural_2015}. Species multiplexes might have an upper bound on redundancy which could explain the low absolute values for robustness.

Robustness, independence and redundancy are only some of the pressures which might affect the structure of TRN-PPI multiplex across the domains of life. Other topological factors might possibly be involved. For instance, theoretical studies have established that controllability \cite{nie_effect_2015} and navigability \cite{battiston_efficient_2016} both depend on multiplex structure. Further, these results have been confirmed in macroscale networks \cite{nie_effect_2015, battiston_efficient_2016}. For multiplex networks, with one-to-one correspondence between the nodes in the two layers, controllability decreases with increasing degree-degree coupling \cite{nie_effect_2015}. This means that the TRN-PPI multiplex might become less controllable at high degree-degree coupling, and more genes and proteins will need to be controlled to steer the multiplex towards a desired state. Navigability is negatively affected by redundancy as well. As the number of overlapping edges, and hence redundancy, increases, navigability decreases \cite{battiston_efficient_2016}. Navigability is quantified by two different metrics; maximum entropy of trajectories explored by a random walker over the multiplex, and uniformity in the steady state probability distribution of node occupation under random walks over the multiplex. Maximum entropy decreases and probability distribution of node occupation becomes more heterogeneous with increasing edge overlap. At high redundancy or edge overlap, low maximum entropy will mean that a random walker can only explore a limited set of trajectories, and highly heterogeneous steady-state distributions will lead to unbiased occupancy among nodes. Therefore, controllability and navigability might exert countering pressures to robustness in shaping the structure of the TRN-PPI multiplex. Disassortative mixing is another important property of molecular networks \cite{maslov2002specificity}. Individual network disassortativity might interact with multiplex coupling to create higher order effects, where degrees of neighbors in individual networks might be coupled in the multiplex. Such high order coupling may have additional impact on robustness and other multiplex properties.

We have identified degree-degree coupling and redundancy as two modulators of multiplex robustness. These modulators can potentially be tuned to control robustness in naturally existing TRN-PPI multiplexes. Further, we can even custom design synthetic TRN-PPI multiplexes to have desired robustness values. For instance, if robustness is the desired property for a set of genes, the multiplex could be rewired such that protein hubs are also highly regulated transcriptionally. On the other hand, if independence between TRN and PPI layers is the desired behavior, degree-degree coupling and redundancy can be reduced synthetically. In principle, similar ideas can be extended to multiplexes comprising different types of molecular species, for example, protein coding mRNA, miRNA and protein-binding mRNA. The results of this study can be easily extended to other molecular multiplexes and can inform the design of novel multiplexes with different molecular species to achieve a desired biological function.

Besides the global design principles for multiplex organization, we have also shown that functionally important genes and proteins have a distinct distribution over the TRN-PPI multiplex. Essential, disease- and pathogen-related genes and proteins are preferentially situated in essential parts of the multiplex. This topological placement is dictated by redundancy. Attack on these functionally important genes quickly dismantles the multiplex. For diseases and pathogens, this suggests that these diseases and pathogens have evolved with the human multiplex and preferentially interact with the vulnerable genes and proteins. Thus, multiplex framework can be useful in the study of disease evolution. Network analysis has previously been used for repurposing existing drugs \cite{guney2016network}. We believe that our multiplex approach might help in better identification of drug targets, since a multiplex better captures the complexity of the underlying molecular networks. Therefore, multiplex framework might have application in network medicine.

One limitation of any network analysis of molecular interactions is incomplete data. This problem is further compounded due to partial overlap between the TRN and PPI networks. However, it has been shown previously that if the size of the incomplete network is above a certain threshold such that a giant component exists, incomplete networks are representative of the complete network \cite{menche2015uncovering}. This suggests that our analysis is representative of the complete species multiplexes. Addition of more genes and proteins in the multiplex might change the specific values of \(C_D\), \(C_R\) and \(R\), however the general dependence between robustness and degree-degree coupling and redundancy will hold. Further, we have not incorporated information about isoform proteins in this study. However, it is straightforward to include such information. In the presence of isoforms, the correspondence between TRN genes and PPI proteins will be one-to-many rather than one-to-one.

Quantifying structure and topology in complex biological networks has been actively researched within network biology. Design principles include the universality of scale-free networks \cite{barabasi_network_2004, albert_scale-free_2005} from metabolic networks across species \cite{jeong_large-scale_2000} and gene regulatory networks \cite{guelzim_topological_2002, lee_transcriptional_2002, babu_structure_2004, babu_evolutionary_2006, faith_large-scale_2007, thieffry_specific_1998}, to power grids and the internet \cite{barabasi_scale-free_2003}, lethal deletions in the hubs of yeast \cite{jeong_lethality_2001}, disassortative mixing in molecular networks \cite{maslov2002specificity} [74], and the existence of sub-modules and reoccurring network motifs \cite{milo_network_2002, shen-orr_network_2002}. Network biology has only recently investigated the influence of multilayered multiplex networks in comparison to single network layers in isolation \cite{reis_avoiding_2014, kleineberg_geometric_2017, klosik_interdependent_2017, liu2019robustness, de_domenico_structural_2015, zheng_control_2019}. This study contributes to the understanding of internetwork connectivity in layered molecular interaction networks. It is the first to compare TRN-PPI multiplexes across species covering two domains of life. We discover global trends across species with degree-degree coupling and edge redundancy positively correlated with increased robustness. Robustness is explored in the context of the TRN-PPI multiplex and is proposed as one of the selective pressures by which evolution has shaped internetwork connectivity, degree-degree coupling and redundancy. The design principles presented here may be useful for the future design and understanding of multiplex networks and to improve efficacy for targeting specific gene subgroups, e.g. in disease. This research presents a multiplex framework for additional investigations of design principles in interlayered biological networks.

\section*{Methods}
\subsection*{Data}
\subsubsection*{Networks}
We compiled network data for nine different species. These nine species span two domains of life, namely bacteria and eukaryotes. There are three bacteria---\textit{H. pylori}, \textit{M. tuberculosis} and \textit{E. coli}---and six eukaryotes---\textit{S. cerevisiae}, \textit{C. elegans}, \textit{D. melanogaster}, \textit{A. thaliana}, \textit{M. musculus} and \textit{H. sapiens}. For the eukaryotes, we collected three datasets---one TRN and two PPI networks. Among the bacteria, \textit{E. coli} also has three datasets (one TRN and two independent PPI networks), while \textit{H. pylori} and \textit{M. tuberculosis} have one TRN and one PPI networks each. These datasets have been collected from diverse sources (see Table S1, Supplementary Information Additional File 2). We use PPI data from multiple published sources---species-specific publications \cite{hauser_second-generation_2014, rajagopala_binary_2014, wang_global_2010}, BioGRID database \cite{chatr-aryamontri_biogrid_2017} and HINT database \cite{das_hint:_2012}. Different experimental methods uncover different information about PPI networks \cite{yu_high-quality_2008}. Therefore, we only use PPI networks inferred from Yeast two-hybrid (Y2H) experiments \cite{fields_novel_1989}. Y2H infers binary protein-protein interactions and is a prominent strategy for identifying protein-protein interactions \cite{bruckner_yeast_2009}. BioGRID and HINT do not have data for \textit{H. pylori} and \textit{M. tuberculosis}. PPI networks for these bacteria were collected from individual publications, Häuser et al. \cite{hauser_second-generation_2014} and Wang et al. 2010 \cite{wang_global_2010} respectively. \textit{E. coli} only exists in HINT. We include another published PPI network for \textit{E. coli} \cite{rajagopala_binary_2014}. A consolidated database of TRN networks across species does not exist. Therefore, we collected protein-DNA interactions from different publications. References for all the species are given in Table S1, Supplementary Information Additional File 2. Available TRN and PPI networks are incomplete. Consequently, they only contain a fraction of the total number of possible genes and proteins in the genome and proteome, respectively. Further, TRN and PPI networks used in this study have different numbers of genes and proteins (see Table S1, Supplementary Information Additional File 2). For a given species, we have only considered genes and proteins which are present in both TRN and PPI networks in our analysis.

The following characteristics of the TRN and PPI networks used in this study are included in Table S1, Supplementary Information Additional File 2---Number of genes and proteins, \% proteome coverage in the TRN-PPI multiplex (fraction of the total proteome covered in the multiplex), number of network edges (edges represent connections between genes and proteins in TRN and PPI networks respectively), average degrees (average K in PPI and average kin or kout in TRN) and size of the Largest Connected Component (LCC) (subset of genes/proteins in TRN/PPI where every gene/protein is reachable from every other gene/protein) are shown for all nine species.

\subsubsection*{Essential genes}
List of essential genes was collected for three species (yeast, fly and human) from the Online GEne Essentiality (OGEE) database \cite{chen_ogee_2016, chen_ogee:_2011}. The database has gene essentiality information on 48 species.

\subsubsection*{Pathogen-related genes}
We collected human-pathogen protein-protein interaction data for 12 different pathogens from a publicly available database HPIDB 3.0 \cite{kumar2010hpidb, ammari2016hpidb}. This curated database contains 69,787 unique protein interactions between 66 host and 668 pathogen species. Human-pathogen protein interactions for various human coronaviruses (HCoVs) were collected from a recently published paper \cite{zhou2020network}.

\subsubsection*{Disease-related genes}
We collected disease-related genes from a publicly available database DisGeNET \cite{pinero2015disgenet, pinero2016disgenet, pinero2020disgenet}. The current version (v6.0) contains gene-disease associations between 17,549 genes and 24,166 diseases, disorders, traits, and clinical or abnormal human phenotypes. We collected disease-gene associations for diseases which have at least 100 genes in the human multiplex considered in this work.

\subsubsection*{Oncogenes and tumor suppressor genes}
We collected the set of genes which contain mutations which have been causally implicated in cancer from the Network of Cancer Genes (NCG) \cite{repana2019network}. NCG also includes information on whether a given cancer gene is an oncogene or TSG.
%\nocite{oreg,schn,pond,smith,marg,hunn,advi,koha,mouse}

\subsection*{Multiplex formulation of transcriptional regulatory and protein-protein interaction networks}
TRN and PPI networks are modeled as interdependent networks (Figure \ref{fig1}A). TRN layer encodes the transcriptional program for producing proteins from genes. The proteins translated from the TRN layer participate in protein-protein interactions in the PPI layer. There is one-to-one correspondence between genes and proteins in the TRN and PPI network layers. This specific configuration of interdependent networks can be reduced to a multiplex network \cite{buldyrev_catastrophic_2010}, and we can apply the framework developed by Buldyrev et al. 2010 \cite{buldyrev_catastrophic_2010}.

We use graph theory to model and analyze TRN-PPI multiplex in this work. TRN and PPI networks are modeled as graphs with nodes representing genes and proteins respectively (Figure \ref{fig1}A). Connections between nodes are represented by edges. PPI edges are undirected. Edges in TRN have directionality---transcription factors have edges emanating from them, while downstream genes have incoming edges.  The connectivity pattern of edges is quantified by the concept of degree at each node. In PPI networks, degree (\(K\)) is the number of edges incident on a protein. For TRN, in-degree (\(k_{in}\)) is the number of transcription factors upstream of a gene, and out-degree (\(k_{out}\)) is the number of genes downstream of a transcription factor.

Since TRN and PPI layers have different coverage of the genome and proteome (see Table S1, Supplementary Information Additional File 2), all analysis was done with genes and proteins present in both TRN and PPI networks.

\subsection*{Quantifying multiplex coupling}

\subsubsection*{Degree-degree coupling}
We quantify degree-degree coupling (\(C_D\)) using either Pearson's correlation or Spearman's rank correlation coefficient (Eqn. \ref{eqn1}).
\begin{equation}\label{eqn1}
C_D = cor(k_{out}, K),
\end{equation}
where \(cor()\) is the sample Pearson's correlation or Spearman's rank correlation coefficient, \(k_{out}\) is the TRN out-degree and \(K\) is the PPI degree.

\subsubsection*{Redundancy coupling}
Redundancy coupling (\(C_R\)) is quantified by the number of edges simultaneously present in TRN and PPI. Assume that \(G^1\) and \(G^2\) are graphs representing TRN and PPI networks respectively, and \(V^1\) and \(V^2\) are the corresponding vertex sets. Let \(E(G^1)\) and \(E(G^2)\) be the edge sets for \(G^1\) and \(G^2\) respectively. The elements of the edge sets are vertex pairs. For instance, \((V^1_i, V^1_j)\in G^1 \)  means that gene \(i\) is a transcription factor regulator for gene \(j\), \((V^2_i, V^2_j)\in  G^2\) means that proteins \(i\) and \(j\) interact with each other. Interactions common between TRN and PPI can be mathematically represented by the number of common edges between \(G^1\) and \(G^2\) (Eqn. \ref{eqn2}).
\begin{equation}\label{eqn2}
    Edges_{12} = \rvert\{e \mid e \in E(G^1), e \in E(G^2) \}\lvert,
\end{equation}
where \(Edges_{12}\) is the number of edges common between \(G^1\) and \(G^2\) and \(e\) represents an edge either in \(G^1\) or \(G^2\). In Figure \ref{fig2}, we compute node-specific redundancy coupling. Here, \(C_R\) for each gene-protein is equal to the number of redundant edges incident on that gene-protein pair. \(C_R\) is either calculated as a z-score, which is computed as
 
\(C_R = \frac{Edges_{12} - mean(Edges_{12}^{null})}{sd(Edges_{12}^{null})}\),

where \(Edges_{12}^{null}\) is the number of redundant edges in a null model, or \(C_R = Edges_{12}\) or \(C_R = mean(Edges_{12}) - mean(Edges_{12}^{null})\). The definition of \(C_R\) used is specified in each figure's caption.

\subsection*{Multiplex robustness}
\subsubsection*{Quantifying multiplex robustness}
We use MCGC to quantify multiplex robustness \cite{buldyrev_catastrophic_2010}. MCGC is the set of genes and proteins which are simultaneously connected in both the network layers---every gene/protein in MCGC is reachable from every other gene/protein in MCGC. MCGC is computed by finding the intersection between the largest connected components (LCCs) of the TRN and PPI network layers. To quantify response to targeted attack, we track the size of the largest MCGC at each step of the attack. We simulate attack on the multiplex via the following algorithm.
\begin{enumerate}
    \item Compute multiplex degree for all gene-protein pairs in the multiplex. Multiplex degree is defined as \(K_{mult}(i)=max(K(i), k_{out}(i))\) \cite{kleineberg_geometric_2017}, where \(K_{mult}(i)\) is the multiplex degree for gene-protein pair \(i\), \(K(i)\) is the degree of protein \(i\) in the PPI network and \(k_{out}(i)\) is the out-degree of gene \(i\) in the TRN network. Order multiplex degrees into a list of gene-protein pairs, \(D\), arranged in decreasing order of multiplex degree.
    \item At step \(L\) of the attack, remove the \(L\)th gene-protein pair in \(D\). Removing a gene (protein) from TRN (PPI) layer may lead to the failure of dependent proteins (genes) in the PPI (TRN) layer. This failure may progress recursively, affecting more nodes in the multiplex. This process is called a cascade of failures \cite{buldyrev_catastrophic_2010}.
    \item After removing the attacked gene-protein pair and other failed dependent nodes at step \(L\) (cascade of failures), find the LCC in either TRN or PPI layer. At this stage, MCGC coincides with the LCC. Compute the size of MCGC. For computing size of MCGC, the TRN network is converted to an undirected version. Therefore, we calculate ‘weak’ MCGC, where weak refers to the undirected nature of TRN.
    \item Repeat steps 2 and 3 of this algorithm until MCGC breaks down.
\end{enumerate}

This algorithm will generate a sequence of values, which give the trajectory of the MCGC as the multiplex is successively attacked. If we plot this trajectory as a function of the fraction of gene-protein pairs removed from the multiplex, robustness to attack can be assessed from either the area under the curve or the critical number of nodes removed for which the MCGC is fragmented \cite{liu_breakdown_2016}. We use area under the curve (\(RobustArea\)) as the measure for multiplex robustness. Thus, \(RobustArea\) is given as
\begin{equation}\label{eqn4}
    RobustArea = \int_0^1 RMCGC(f)\, df,
\end{equation}
where \(RobustArea \in (0, 1]\) is the area under the curve, \(f\) is the fraction of gene-protein pairs removed during the attack, and \(RMCGC(f)\) is the size of MCGC relative to the total number of gene-protein pairs in the multiplex (\(n\)) after \(f\) fraction of gene-protein pairs have been removed from the multiplex. \(RobustArea\) quantifies absolute robustness. Relative robustness (\(R\)) is computed by comparing \(RobustArea\) against a null model. Cohen’s d is used to compute effect size of relative robustness (Eqn. \ref{eqn2}).
\begin{equation}\label{eqn5}
R = \frac{{mean(RobustArea_{obs}) - mean(RobustArea_{null})}}{\sqrt{\frac{var(RobustArea_{obs}) + var(RobustArea_{null})}{2}}},
\end{equation}
Where \(R\) is the relative robustness, \(RobustArea_{obs}\) and \(RobustArea_{null}\) are the \(RobustArea\) values for the observed multiplex and null model respectively and \(mean()\) and \(var()\) are the mean and variance functions respectively. We have assumed that \(RobustArea_{obs}\) and \(RobustArea_{null}\) have the same number of samples.

\subsubsection*{Robustness for partial attack curves}
For Figure \ref{fig5}, we estimate robustness for a subset of functionally important gene-protein pairs and compare that against the remaining gene-protein pairs in the multiplex. Here, we explain the strategy to conduct such a comparison. Assume that \(S = \{S_1, S_2,..., S_M\}\) is a collection of sets of gene-protein pairs in a multiplex. Here \(M\) is the total number of sets. Sets \(S_i\), \(i \in \{1, 2,..., M\} \), can be mutually exclusive or not. Let \(M_{min}\) be the size of the smallest set in S. For an equitable comparison, we randomly sample \(M_{min}\) number of gene-protein pairs from all the subsets, except for the smallest subset. We sample each subset 100 times. For each sampled version of a subset \(S_i \in S\), we attack the gene-protein pairs in \(S_i\) in decreasing order of multiplex degree for gene-protein pairs in that set, using the attack algorithm explained previously. We stop the attack once \(M_{min}\) number of gene-protein pairs have been removed from the multiplex. We also perform a similar partial attack on a set of randomly selected gene-protein pairs. This size of the random set is set equal to \(M_{min}\). Robustness can be calculated for each subset based on the obtained partial attack curves. Relative robustness for each subset is calculated by comparing \(RobustArea\) for that subset against the random set.
%%%%%%%%%%%%%%%%%%%%%%%%%%%%%%%%%%%%%%%%%%%%%%
%%                                          %%
%% Backmatter begins here                   %%
%%                                          %%
%%%%%%%%%%%%%%%%%%%%%%%%%%%%%%%%%%%%%%%%%%%%%%

\begin{backmatter}

\section*{Competing interests}
  The authors declare that they have no competing interests.

\section*{Author's contributions}
    R.D.D. and T.M. conceived the analytical and computational work. T.M. carried out the computational work. R.D.D. and T.M. analyzed and interpreted the data. R.D.D. and T.M. wrote the manuscript.

\section*{Acknowledgements}
  We are grateful to the members of the Dar lab for fruitful discussions on the manuscript.

\section*{Availability of data and code}
All TRN and PPI networks are provided as an R programming language \cite{Rlang} data object ("NetworkMultiplex.RData" in Additional File 3). Data for all the pathogens and diseases are also provided as R programming language data objects in Additional file 3.
\linebreak

All the analysis was performed in the R programming language \cite{Rlang}. Custom scripts for reproducing Figures 2-5 are provided in Additional File 3.
%%%%%%%%%%%%%%%%%%%%%%%%%%%%%%%%%%%%%%%%%%%%%%%%%%%%%%%%%%%%%
%%                  The Bibliography                       %%
%%                                                         %%
%%  Bmc_mathpys.bst  will be used to                       %%
%%  create a .BBL file for submission.                     %%
%%  After submission of the .TEX file,                     %%
%%  you will be prompted to submit your .BBL file.         %%
%%                                                         %%
%%                                                         %%
%%  Note that the displayed Bibliography will not          %%
%%  necessarily be rendered by Latex exactly as specified  %%
%%  in the online Instructions for Authors.                %%
%%                                                         %%
%%%%%%%%%%%%%%%%%%%%%%%%%%%%%%%%%%%%%%%%%%%%%%%%%%%%%%%%%%%%%

% if your bibliography is in bibtex format, use those commands:
\bibliographystyle{bmc-mathphys} % Style BST file (bmc-mathphys, vancouver, spbasic).
\bibliography{Mahajan_Dar_2020}      % Bibliography file (usually '*.bib' )
% for author-year bibliography (bmc-mathphys or spbasic)
% a) write to bib file (bmc-mathphys only)
% @settings{label, options="nameyear"}
% b) uncomment next line
%\nocite{label}

% or include bibliography directly:
% \begin{thebibliography}
% \bibitem{b1}
% \end{thebibliography}

%%%%%%%%%%%%%%%%%%%%%%%%%%%%%%%%%%%
%%                               %%
%% Figures                       %%
%%                               %%
%% NB: this is for captions and  %%
%% Titles. All graphics must be  %%
%% submitted separately and NOT  %%
%% included in the Tex document  %%
%%                               %%
%%%%%%%%%%%%%%%%%%%%%%%%%%%%%%%%%%%

%%
%% Do not use \listoffigures as most will included as separate files

\section*{Figures}


%  \begin{figure}[h!]
%  \caption{\csentence{Sample figure title.}
%      A short description of the figure content
%      should go here.}
%      \end{figure}

%\begin{figure}[h!]
%  \caption{\csentence{Sample figure title.}
%      Figure legend text.}
%      \end{figure}

\begin{figure}[ht!]
  %\includegraphics[width=0.9\linewidth]{Figure_1.png}
  \caption{\csentence{Degree-degree coupling and redundancy as potential modulators of robustness in molecular multiplexes.} Degree-degree coupling (\(C_D\)) and redundancy coupling (\(C_R\)) can assume any value in the multiplex of TRN (yellow layer) and PPI (green layer) networks across species. A) (Right, Top) For \(C_D > 0\), highly connected genes in TRN (red spheres) are more likely to produce hubs or highly connected proteins in PPI (red spheres), while sparsely connected genes in TRN (blue spheres) are highly likely to produce spokes or sparsely connected proteins in PPI (blue spheres). (Right, Bottom) For \(C_D = 0\), TRN and PPI will be uncoupled in the multiplex, and the association between genes and proteins would be randomized. Here, highly connected genes produce spoke proteins and vice-versa sparsely connected genes produce hub proteins. (Left) Based on theoretical studies, \(C_D\) is expected to be positively correlated with robustness. Robustness is quantified by area under the attack curve for the Mutually Connected Giant Component (MCGC) (Methods). B) (Left, Top) For \(C_R > 0\), there will be non-zero number of edges which are simultaneously present in both TRN and PPI (edges marked green). (Left, Bottom) For \(C_R = 0\), TRN and PPI will not have common edges (no green edges). (Right) Based on theoretical studies, \(C_R\) is expected to be positively correlated with robustness. Black directed edges represent regulation of downstream genes by transcription factors in TRN. Black undirected edges represent protein-protein interactions in PPI. Dashed edges represent correspondence between a gene and its corresponding protein. }
  \label{fig1}
\end{figure}

\begin{figure}[ht!]
  %\includegraphics[width=0.9\linewidth]{Figure_2.pdf}
  \caption{\csentence{TRN-PPI multiplex is coupled across species.}  Scatter plot for degree (\(K\)) of proteins in the PPI network versus out-degree (\(k_{out}\)) of genes in the TRN for eukaryotes, (A) yeast, B) fly, C) mouse and D) human. Both \(K\) and \(k_{out}\) values have been log-transformed after adding 1. The PPI networks are from the BioGRID database (Methods). Linear interpolated fits between \(K\) and \(k_{out}\) are also shown (green line) with 95\% confidence region shaded in gray. Degree-degree coupling (\(C_D\)) values and corresponding p-values are also shown. Distribution of redundancy (\(C_R\)) for gene-protein pairs for species multiplex and the randomly shuffled null model for eukaryotes, (E) yeast, F) fly, G) mouse and H) human. For each gene-protein pair, \(C_R\) is quantified by the number of redundant edges incident on that gene-protein pair. Shuffled null model is generated by randomly shuffling labels on genes in TRN, while keeping protein labels fixed in PPI. Jensen Shannon divergence (JSD) between distributions of \(C_R\) in organismal and shuffled multiplexes is also given in E-H. }
  \label{fig2}
\end{figure}

\begin{figure}[ht!]
  %\includegraphics[width=0.9\linewidth]{Figure_3.png}
  \caption{\csentence{Multiplex attack curves for eukaryotes.}  Relative size of 
the mutually connected giant component (MCGC) is plotted as a function of the fraction of gene-protein pairs attacked and removed from the multiplex (Methods). Attack curves are shown for five species using PPI networks from either BioGRID or HINT databases (Methods); database for a given panel are annotated next to the species name. Along with the attack curves for the species (red), attack curves for the Zero-Coupling-Zero-Redundancy (Supplementary Information, Additional File 1) null model are also shown (gray). Under this null model, multiplexes have no degree-degree coupling and no redundancy. On average, species attack curves are more robust than the null model. Robustness is quantified by the area under the curve. Attack on the multiplex (organism or null) is repeated 1000 times. }
  \label{fig3}
\end{figure}

\begin{figure}[ht!]
  %\includegraphics[width=0.9\linewidth]{Figure_4.png}
  \caption{\csentence{Degree-degree and redundancy couplings modulate multiplex robustness.} A) Robustness to targeted attack (R) (right) is positively correlated with degree-degree coupling (\(C_D\)) (left) across species with Spearman's rank correlation coefficient of 0.6803 (p-value = 0.04372). Robustness to targeted attack (R) (right) is also positively correlated with redundancy coupling (\(C_R\)) (center) across species with Spearman's rank correlation coefficient of 0.7167 (p-value = 0.03687). \(C_R\) is computed as a z-score against the null model (Methods) B) We sample a subset of gene-protein pairs from the species multiplexes (sizes for the subsets are: \textit{S. cerevisiae}-1000, \textit{D. melanogaster}-2000, \textit{M. musculus}-300, \textit{H. sapiens}-500) with specific \(C_D\) and \(C_R\) values. We repeat the sampling 100 times. We explore \(C_D\) and \(C_R\) over a grid. For each point over the 2D grid, the heatmap shows the robustness (\(R\)) value. \(R\) is computed by comparing \(RobustArea\) of any point over the grid against the lower-left point of the grid (with \(C_D\) = 0, \(C_R\) = 0). For each point over the grid, for each of the sampled subsets, targeted attack is performed. Mean \(R\) values are shown here. Lower and upper 95\% confidence interval (CI) values are shown in Figure S5, Supplementary Information, Additional file 1. \(C_{D}^{out}\): degree-degree coupling between \(k_{out}\) and \(K\),  \(C_{D}^{in}\): degree-degree coupling between \(k_{in}\) and \(K\). C) (Left) Dependence of robustness on degree-degree coupling while redundancy is fixed. For each species curve, \(R\) is calculated by comparison against the sampled multiplex with \(C_D = 0\). (Right) Dependence of robustness on redundancy coupling while degree-degree coupling is fixed to the value in the full multiplex. For each species curve, \(R\) is calculated by comparison against the sampled multiplex with \(C_R = 0\). In all the panels, error bars show 95\% CIs. In panel B, BioGRID PPI networks are used; results with HINT PPI networks are shown in Figure S6B, Supplementary Information, Additional File 1. For panels B and C, \(C_R\) is set equal to the number of redundant edges in the multiplex.}
  \label{fig4}
\end{figure}

\begin{figure}[ht!]
  %\includegraphics[width=0.9\linewidth]{Figure_4.png}
  \caption{\csentence{Functionally important genes and proteins are redundant and essential for human multiplex.}  A) (Left) Essential genes and proteins (red) have lower robustness to targeted attack across species compared to non-essential genes and proteins (blue).  (Right) Lower \(R\) is accompanied by higher redundancy (\(C_R\)) for essential genes and proteins compared to non-essential genes and proteins. B) (Left) Pathogen-related genes and proteins (red) have lower \(R\) to targeted attack across pathogens compared to non-pathogen genes and proteins (blue). (Right) Lower \(R\) is accompanied by higher \(C_R\) for pathogen genes and proteins compared to non-pathogen genes and proteins. C) (Left) Disease-related genes and proteins (red) have lower \(R\) to targeted attack across diseases compared to non-disease genes and proteins (blue). (Right) Lower \(R\) is accompanied by higher \(C_R\) for disease genes and proteins compared to non-disease genes and proteins. D) (Left) oncogenes (tumor suppressor genes (TSGs)) and proteins (red) have lower \(R\) to targeted attack across pathogens compared to non-oncogenes (-TSGs) and proteins (blue). (Right) Lower \(R\) is accompanied by higher \(C_R\) for oncogenes (TSGs) and proteins compared to non-oncogenes (-TSGs) and proteins. In all the panels, database used for PPI networks is annotated on the y-axis (Methods). In all the panels, relative robustness (\(R\)) is measured against a random set of genes and proteins used as the null (Methods), and \(C_R\) values are calculated as the difference in redundancy against the random set of genes. Error bars show 95\% CIs. For all the panels, \(C_R\) is calculated as the mean difference in the number of redundant edges between the species' multiplex and the random set of gene-protein pairs.}
  \label{fig5}
\end{figure}


%%%%%%%%%%%%%%%%%%%%%%%%%%%%%%%%%%%
%%                               %%
%% Tables                        %%
%%                               %%
%%%%%%%%%%%%%%%%%%%%%%%%%%%%%%%%%%%

%% Use of \listoftables is discouraged.
%%
%\section*{Tables}

%%%%%%%%%%%%%%%%%%%%%%%%%%%%%%%%%%%
%%                               %%
%% Additional Files              %%
%%                               %%
%%%%%%%%%%%%%%%%%%%%%%%%%%%%%%%%%%%
\clearpage
\section*{Additional Files}
  \subsection*{Additional File 1 --- Supplementary Information.}
    Supplementary text.

  \subsection*{Additional File 2 --- Table S1.}
    Supplementary table listing the properties and sources for the TRN and PPI networks used in this study.
    
  \subsection*{Additional File 3 --- Code.}
    Custom scripts written in the R programming language \cite{Rlang} for reproducing Figures 2-5.
  
\end{backmatter}
\end{document}
