## Supplementary material for "Internetwork connectivity of molecular networks across species of life": Latex Files: readme.html

BioMed Central TeX template files

### BioMed Central TeX template - Version 0.6 (April 15^th^ 2015)

#### 1 BioMed Central LaTeX template distribution

|  |  |
| --- | --- |
| bmc\_article.tex | The BioMed Central manuscript template. Edit a copy of this file. |
| bmc\_article.pdf | PDF created from The BioMed Central manuscript template. |
| bmc\_article\_2col.pdf | Two column layout PDF created from The BioMed Central manuscript template. |
| bmc\_article.bib | A sample BibTeX bibliography file used to create a .bbl by BibTeX using the bmc-mathphys.bst file. |
| bmc\_article.bbl | A sample bibliography file created by BibTeX using the bmc-mathphys.bst file. |
| bmc-mathphys.bst | The BibTeX bibliography style file for the BioMed Central reference format |
| spbasic.bst | The BibTeX bibliography style file for the Springer Basic reference format |
| vancouver.bst | The BibTeX bibliography style file for the Vancouver reference format |
||  |
| --- | --- |
||  |
| --- | --- |
| bmcart.cls | Page layout style class for for manuscripts. |
| bmcart-biblio.sty | Style for bibliography tags. |
| readme.html | This document. |

  |  |

#### 2 Requirements

### 2.1

The BioMed Central TeX template
should work with TeX distributions on any platform - it has been tested with
TeXShop on Mac OS
X and MiKTeX for Windows.

### 2.2

In order to submit a manuscript
as a .tex file to BioMed Central, you *must*

- use the BioMed Central template
- format your references with BibTeX using
  the bmc-mathphys.bst style file
- not rely on any non-standard macros,
  classes or files

If your TeX manuscript does not
meet the criteria above it will need to be converted to DVI format prior to
submission.

#### 3 Guidelines for creating your manuscript using TeX

### 3.1

Follow the guidelines in the BioMed Central instructions
for authors given at http://www.biomedcentral.com/authors/.

### 3.2

Make sure your manuscript is compiled with LaTeX2e by using \documentclass{...} and not \documentstyle{...}
in the preamble at the top of your .tex  document.

A template manuscript is supplied entitled bmc\_article.tex which
sets up the preferred page layout based on the standard article.cls. Additional styles can be achieved by
\documenclass options: *doublespacing* - for double spaced text, *linenumbers* - for the line numbers on margins,
*twocolumn* - for twocolumn layout. Please note that for twocolumn layout \end{fmbox} position is different,
comment one after \end{artnotes} environment and uncomment one after \end{abstractbox} environment.

### 3.3

Make sure that you only a single .tex  document
for the entire manuscript, as you will need to upload it as a single file (together
with its associated formatted bibliography file). Do not use the \input command
to include other .tex files.

### 3.4

Additional packages can be used in The BioMed Central template: *amsthm* and *amsmath* for
theorems and mathematics respectively, *natbib* for citation style, *hyperref* for url references.

See

http://www.ctan.org/pkg/amsthm  
http://mirrors.ctan.org/macros/latex/required/amslatex/math/amsmath.sty  
http://www.ctan.org/tex-archive/macros/latex/contrib/natbib  
http://www.ctan.org/tex-archive/macros/latex/contrib/hyperref

for these style files if they are not in your local archive.

*natbib.sty* with sort&compress option turns [1,2,3,4] into [1-4].

*hyperref.sty* formats urls so they can be broken down
cleanly when overflowing the right text boundary.

#### 4 BibTeX

References *must* be formatted with BibTeX using
the BioMed Central style file.

I.e. when using the template, choose one of the following: \bibliographystyle{bmc-mathphys},
\bibliographystyle{vancouver} or \bibliographystyle{spbasic} depending on the reference
style that your journal is using.

The bibliography datafile is referred to with \bibliography{datafile1, ..., ...}.

The template makes use of a sample bibliography called bmc\_article.bib -
you should update the \bibliography tag to refer to your own bibliography.

For *author-year* bibliography (bmc-mathphys or spbasic):

1. write to bib file (bmc-mathphys only): `@settings{label, options="nameyear"}`
2. write to tex file: `\nocite{label}`

#### 5 Notes on uploading your manuscript

### 5.1

Make sure you are submitting only one .tex document. Please note that figures,
large tables and any other reference material should be submitted as separate files, not embedded in the
manuscript.

### 5.2

A .bbl file is generated when you use BibTeX
to format your article's reference list. It contains formatted details of all
references used in the manuscript. After uploading a TeX file to BioMed Central,
you will then be prompted to upload the .bbl file which goes with it.

#### 6 The TeX article layout

This is the sectioning for a BMC-series Research Article l
manuscript submission . . .

• Abstract

• Background

• Results

• Conclusions

• Background

• Methods

• Results and Discussion

• Conclusions

• Authors contributions

• Acknowledgements

• References

• Figures

• Tables

• Additional files

#### 7 Further information

The latest version of the BioMed Central TeX
template distribution, and full instructions on its use, are available here:

http://www.biomedcentral.com/ifora/tex/
