## Supplementary material for "Internetwork connectivity of molecular networks across species of life": Latex Files: Reference PDF.pdf

$C_R = \frac{Edges_{12} - mean(Edges_{12}^{null})}{sd(Edges_{12}^{null})}$,

where $Edges_{12}^{null}$ is the number of redundant edges in a null model, or $C_R = Edges_{12}$ or $C_R = mean(Edges_{12}) - mean(Edges_{12}^{null})$. The definition of $C_R$ used is specified in each figure's caption.

$$R = \frac{mean(RobustArea_{obs}) - mean(RobustArea_{null})}{\sqrt{\frac{var(RobustArea_{obs}) + var(RobustArea_{null})}{2}}}, \tag{4}$$

Where $R$ is the relative robustness, $RobustArea_{obs}$ and $RobustArea_{null}$ are the $RobustArea$ values for the observed multiplex and null model respectively and $mean()$ and $var()$ are the mean and variance functions respectively. We have assumed that $RobustArea_{obs}$ and $RobustArea_{null}$ have the same number of samples.

All the analysis was performed in the R programming language [84]. Custom scripts for reproducing Figures 2-5 are provided in Additional File 3.

**Author details**

[1]Department of Bioengineering, University of Illinois at Urbana-Champaign, 1406 W. Green St, 61801 Urbana, IL, USA. [2]Department of Electrical and Computer Engineering, University of Illinois at Urbana-Champaign, 306 North Wright St, 61801 Urbana, IL, USA. [3]Center for Biophysics and Quantitative Biology, University of Illinois at Urbana-Champaign, 1110 West Green Street, 61801 Urbana, IL, USA. [4]Carl R. Woese Institute for Genomic Biology, University of Illinois at Urbana-Champaign, 1206 West Gregory Drive, 61801 Urbana, IL, USA.
